# FPM app: an open-source application for simple and intuitive Fourier ptychographic reconstruction

**DOI:** 10.1101/2021.03.25.436959

**Authors:** Mikołaj Rogalski, Piotr Zdańkowski, Maciej Trusiak

## Abstract

**Summary:** Fourier ptychographic microscopy (FPM) is a computational microscopy technique that enables large field of view and high-resolution microscopic imaging of biological samples. However, the FPM does not yet have an adequately capable open-source software. In order to fill this gap we are presenting novel, simple, universal, semi-automatic and highly intuitive graphical user interface (GUI) open-source application called the *FPM app* enabling wide-scale robust FPM reconstruction. Apart from implementing the FPM in accessible GUI app, we also made several improvements in the FPM image reconstruction process itself, making the FPM more automatic, noise-robust and faster.

**Availability and Implementation:** *FPM app* was implemented in MATLAB and all MATLAB codes along with standalone executable version of the *FPM app* and the online documentation are freely accessible at https://github.com/MRogalski96/FPM-app. Our exemplary FPM datasets may be downloaded at https://bit.ly/2MxNpGb.

## 1 Introduction

One of the main limitations of optical microscopy is the fact that while increasing the resolution of a microscope, due to objective’s mechanical construction, its field of view (FOV) is decreasing – it is not possible to observe the sample with both high resolution and large FOV. Fourier ptychographic microscopy (FPM) (Zheng et al., 2013) is one of a few imaging techniques that can overcome this pivotal limitation. It achieves this by creating a synthetic aperture in the Fourier space through combining the data from many illumination angles to increase the object information content in the final image. As a result, FPM returns an image with the resolution significantly higher than the objective’s diffraction limit, while maintaining the objective’s FOV. Moreover, iterative FPM reconstruction algorithms allow to retrieve object’s quantitative phase information which is particularly significant in the case of label-free high-contrast bioimaging (Ou et al., 2013) and calculate the pupil function of the employed imaging system, which can provide crucial information about system aberrations (Ou et al., 2014). These advantages can be obtained with a relatively low cost, e.g., by modifying classical brightfield microscope with LED array placed instead of the brightfield illumination (every LED will illuminate sample from a different angle).

As a computational imaging technique the FPM is highly dependent on the numerical reconstruction process. However, it lacks simple open-source software that would be appropriate for non-expert users, what significantly reduces its popularity. To make the FPM more approachable and allow a wider audience to benefit from its advantages we are proposing a novel graphical user interface (GUI) application called *FPM app,* which enables to perform FPM reconstruction in a simple and semi-automatic fashion. Moreover, in *FPM app* we included several improvements making the FPM reconstruction process more automatic, noise-robust and faster.

*FPM app* was created in MATLAB 2019a (based on Quasi-Newton algorithm implementation (Tian et al., 2014) and originally developed procedures) - all MATLAB codes (that are free to modify for personal use) along with *FPM app Documentation* (consisting user manual, methods used for processing data and exemplary *FPM app* usage) can be found in Supplementary Data attached to this article.

## 2 Features and methods

### 2.1 Generating synthetic FPM datasets

*FPM app* enables to generate synthetic FPM datasets – a set of low-resolution images determined by the user-specified synthetic FPM system parameters. Each one of these generated images is representing an image recorded by a microscope with a different illumination angle. Reconstructing of these synthetic datasets may be especially useful in learning how the FPM works and what is the influence of the certain input parameters onto the reconstruction. Additionally, simulating the synthetic FPM system may be extremely helpful in planning a real-life experiment. Detailed description of synthetic data generation process may be found in the *FPM app Documentation*, Chapter 4.3.

### 2.2 Reconstructing FPM datasets

Thanks to the simple and intuitive GUI, user can easily define all necessary system parameters (e.g., LED array shape and position, image collecting order, used objective lens) to perform Fourier ptychographic reconstruction. Apart from that, user may also adjust the reconstruction parameters by selecting different options. As a result, *FPM app* returns high resolution images of object phase (**Fig. 1**) and amplitude along with reconstructed complex pupil function of employed microscope system. Finally, *FPM app* allows to postprocess, visualize and save reconstruction results. Detailed description of FPM app reconstruction process may be found in *FPM app Documentation*, Chapter 4.4.

**Fig. 1.**
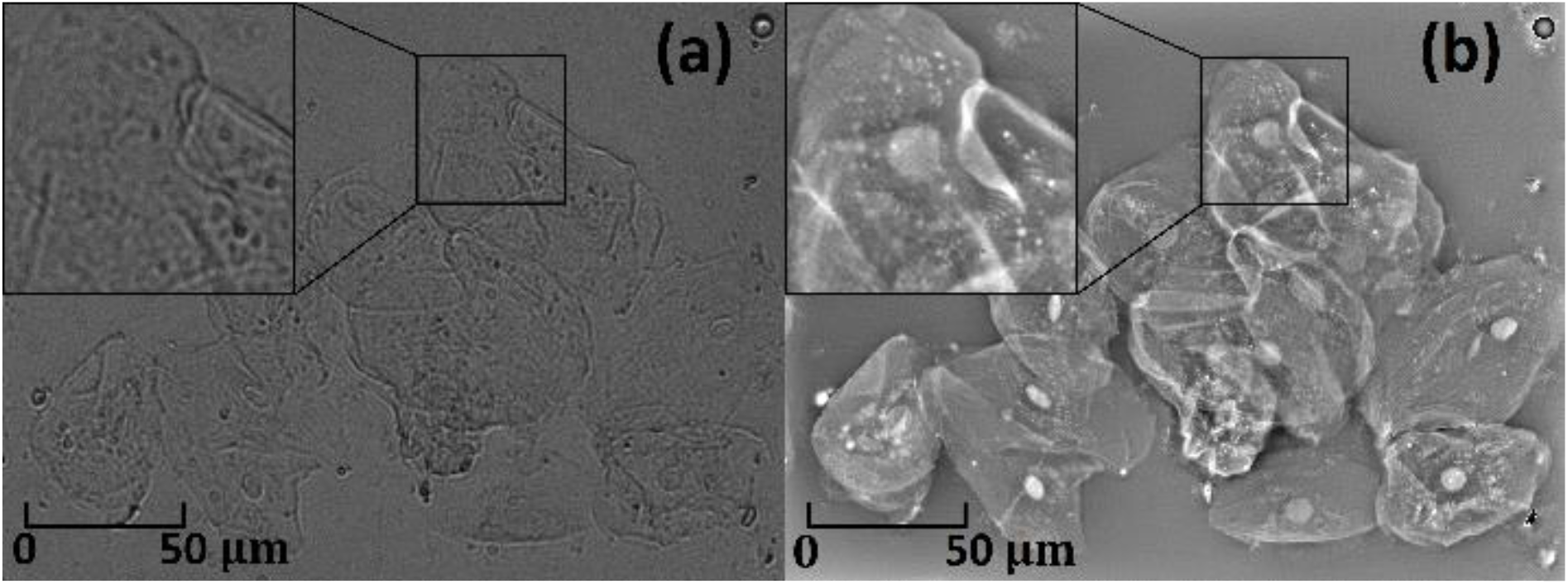
Exemplary FPM app performance. (a) – cheek cell image obtained on our 10×0.25 brightfield microscope. (b) – FPM phase reconstruction obtained on the same microscope (137 input images, synthetic NA = 0.66). Such improvement in cell bio-imaging was achieved solely with the use of 13×13 LED array and the FPM app.

### 2.3 Our modifications in the FPM reconstruction process

Despite the undoubted advantages, FPM is also faced with several problems, i.e., long reconstruction time, high-frequency noise and erroneous reconstructions in poorly aligned systems. To minimize their influence, we have added several improvements into the FPM processing path that are described below.

1. Novel background removing algorithm, that can fully automatically reduce background noise from the input FPM images and therefore remove the high-frequency artifacts from the FPM reconstruction.
2. Adding possibility to perform some of the algorithm operations on the graphical processing unit, which may decrease reconstruction time even several times, especially for large datasets.
3. Applying two simple, fast and robust LED misalignment position correction algorithms, that can enhance reconstruction on poorly aligned systems with similar gain but in up to 5x shorter time than reference solution based on Simulated Annealing algorithm (Sun et al., 2016).
4. Adding possibility to further improve reconstruction results by the state-of-the-art block-matching and 3D filtering (BM3D) (Dabov et al., 2007) denoising algorithm, which significantly reduces high-frequency noise in phase and amplitude reconstructed maps.

We have tested *FPM app* performance by carrying out FPM reconstruction on: (1) synthetic data, (2) real data available on the Internet (at https://www.laurawaller.com/opensource/) and (3) original experimental data. Validation and broader description of all our improvements can be found in the *FPM app Documentation*, Chapter 5.

## 3 Summary

In summary, *FPM app* is a first widely available software, that allows for straightforward and intuitive Fourier ptychographic reconstruction. It should be compatible with most commonly used FPM systems, including the simplest solutions with LED illumination arrays. Thanks to the proposed improvements, FPM reconstruction can be performed faster, in automatized fashion, and provide better results than classical solutions. Moreover, it can be correctly performed in reasonable time even when the FPM system is not perfectly aligned. *FPM app* significantly lowers the time/finance/effort entrance barrier into the FPM imaging and should result in luring more non-expert users to benefit from the FPM advantages and promote its usage worldwide. We believe, that by the dissemination of freely available *FPM app* and by sharing the codes and algorithmic advancements, the stage for democratizing the phase and amplitude contrast microscopy with high space-bandwidth product is set.

## Supporting information

FPM app Documentation

FPM app

## Acknowledgements

The work has been funded, in part, by BIOTECHMED-1 and FOTECH-1 projects granted by Warsaw University of Technology under the program Excellence Initiative: Research University (ID-UB); National Science Center Poland NCN (2017/25/B/ST7/02049) and Foundation for Polish Science FNP (START 2020). The contribution was partially funded by the Warsaw University of Technology statutory funds.

