## Supplementary material for "FPM app: an open-source application for simple and intuitive Fourier ptychographic reconstruction": FPM app Documentation

#### Supplementary documentation

#### and user's guide

Mikołaj Rogalski[1] *, Piotr Zdańkowski[1] *, Maciej Trusiak[1]

*

[1]Warsaw University of Technology, Institute of Micromechanics and Photonics, Boboli 8, 02-525 Warsaw, Poland

#### About this text

This is the supplementary documentation for a *FPM app* – a graphical user interface application which provides tools for processing data acquired by Fourier ptychographic microscopy (FPM). *FPM app* contains methods for generating synthetic FPM data and for reconstructing both: synthetic and real FPM datasets. Several own modification in FPM processing path are also implemented, which improves FPM reconstruction performance.

*FPM app* is distributed under the terms of the GNU GPLv2.0 license and the source code and the associated documentation are available at the project website: https://github.com/MRogalski96/FPM-app.

#### Acknowledgments

The work has been funded, in part, by BIOTECHMED-1 and FOTECH-1 projects granted by Warsaw University of Technology under the program Excellence Initiative: Research University (ID-UB); National Science Center Poland NCN (2017/25/B/ST7/02049) and Foundation for Polish Science FNP (START 2020). The contribution was partially funded by the Warsaw University of Technology statutory funds.

### 1. About Fourier ptychographic microscopy

Microscopy is a field of science that allows people to see things that are invisible to the human eye. One of the most popular types of microscopes and first one that were developed are optical brightfield microscopes. Those microscopes are characterized by using visible light to illuminate the observed sample and using lenses to image sample onto detector. Optical microscopy has been developed over the centuries and many interesting techniques such as fluorescence or super-resolution microscopy has been developed during this time.

Regardless of the type of used optical microscope, there is a principle that the larger is the resolution (which is defined by the objective numerical aperture NA) of used microscope, the smaller is its field of view (FOV). The higher the NA of the objective, the finer structures can be resolved by the imaging system. However, due to objectives mechanical properties (large NA objectives has also large magnifications), increasing the NA also decreases the FOV. In most applications only high resolution or only large FOV of the microscope is sufficient but not ideal, but there are several areas of medicine (like digital pathology, cytology or hematology) where both of these parameters are required to be high/large. There are a few techniques that allow high resolution imaging of a sample over large FOV (motorized scanning and image stitching, structured illumination microscopy [1], with the use of mesolens [2] or with the use of deep learning [3]), but usually they are complex and expensive or have other significant disadvantages. However, there is one method – Fourier ptychographic microscopy (FPM) [4], [5], that is relatively cheap and simple and it is considered by many people as the best for high resolution imaging at the large FOV (high space-bandwidth product imaging).

FPM is a computational microscopy technique that allows obtaining microscopic image of a measured sample with much higher resolution than it would follow from the NA of used microscope objective. Such high resolution is obtained by combining in the Fourier domain information about the measured specimen from different illumination directions. This advantage combined with the fact that low NA objectives are characterized by low magnification, allows obtaining a high-resolution image of a specimen with a large field of view. Moreover, iterative algorithms used in the process of reconstruction, enable obtaining information not only about the amplitude but also about the phase of the sample. The phase distribution is especially significant in the case of bioimaging, as living cells are known to be semi-transparent objects with low amplitude, hence difficult to see under the classical brightfield microscope. More advanced FPM algorithms can also provide information about aberrations of the microscope setup [6] or render 3D sample reconstruction [7].

Another great advantage of the FPM is the fact that the hardware part of the microscope can be quite easily assembled. The simplest way to create FPM system is to modify standard brightfield microscope by replacing illuminator with the LED matrix – every LED will illuminate sample from the different angle. Exemplary simple FPM system is shown in the Figure 1.

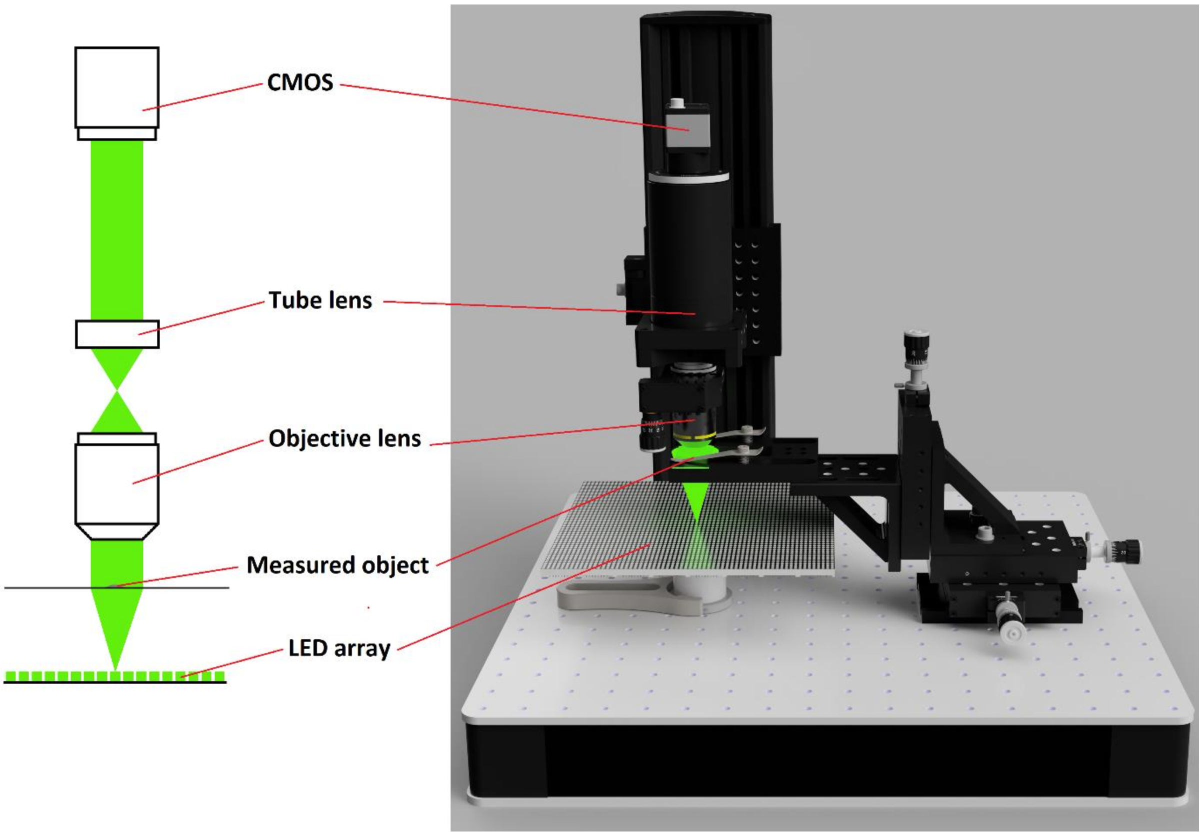

*Figure 1. Scheme and 3D model of the exemplary FPM imaging system.*

### 2. About *FPM app*

Despite the fact that the hardware part of the FPM may be quite simple to assemble, it is not that easy to implement its software part, especially for users which are not proficient in computational microscopy. Up to date, there is no open source FPM software with graphical user interface (GUI), allowing for straightforward ptychographic image reconstruction. The only open source software that can be found in the Internet are a few MATLAB scripts [8]–[11], that may be good to learn FPM, but are cumbersome to adapt to a given experimental setup, mainly due to the fact that every FPM hardware setup may differ from each other on many levels.

To make FPM more accessible, especially for new and non-experts users we are presenting the ***FPM app***: the first simple, intuitive, universal, semi-automatic, easy and open to modifications, GUI open source FPM application. Moreover, we made several improvements in the reconstruction process (see Chapter 5) that make FPM more automatic, noise-robust and faster (hence easier to use).

*FPM app* is released in 2 versions:

- **MATLAB version** – it contains a pack of MATLAB codes (*FPMapp_1.0.zip* file), which are used to open *FPM app* through MATLAB. These codes are open to be modified, to adjust *FPM app* to a given set of preferences or to further improve it.
- **Executable version** – it contains an installer (*FPMapp_Installer_web.exe* file) that installs *FPMapp.exe* executable file along with all the necessary files and *MATLAB Runtime* that is required to run MATLAB standalone applications.

Both of these versions are available at https://github.com/MRogalski96/FPM-app, which will also include future updates of the *FPM app*. Our exemplary datasets are available at https://bit.ly/2MxNpGb.

#### 2.1. Compatibility

To use *FPM app* we recommend to have at least 4GB RAM (or at least 4x more RAM than the size of dataset that you will be reconstructing) and Nvidia's graphics card with current CUDA driver and video memory of at least 2GB (necessary when you want to use GPU acceleration, otherwise integrated GPU is sufficient). The more video memory has your GPU, the larger dataset you can reconstruct with GPU acceleration.

**MATLAB version:**

*FPM app* was created and tested in MATLAB 2019a but should also be compatible with other MATLAB versions.

To use it you need MATLAB with the following toolboxes:

- Signal Processing Toolbox
- Image Processing Toolbox
- Global Optimization Toolbox (only needed for Simulated Annealing LED correction method)
- Parallel Computing Toolbox (only needed for GPU acceleration)

**Executable version:**

*FPMapp.exe* was tested in Windows 7 and 10, but should be also compatible with other Windows 64-bit operating systems.

To run it, you need *MATLAB Runtime* tool. If absent, it will be installed automatically during the *FPM app* installation

#### 2.2. Installation and launching

**MATLAB version:**

- Installation – unpack *FPMapp_1.0.zip* into separate folder.
- Launching – set directory containing *FPM app* as current working directory and run *main.m* script with the MATLAB. It will open *FPM app* main window (Chapter 3.2).

**Executable version:**

- Installation – Open *FPMapp_Installer_web.exe* and follow the instructions on the screen. During the installation, *MATLAB Runtime* tool will be installed (if it is not installed already). On typical desktop computer, installation should not least longer than 20 s (without *MATLAB Runtime* installation). Additional *MATLAB Runtime* installation would also require time needed to download and install 2,8 GB package (approximately several minutes).
- Launching: Open *FPMapp.exe*, it will open FPM app window (Chapter 3.2). *FPMapp.exe* should be installed in *your_directory_path/WUT/FPM app/application* directory.

  WARNING 1: Installation of the *FPM app* in the directory that needs administrator permissions may cause several errors. To avoid them open *FPM app* as administrator or install app in directory that does not require administrator permissions.

  WARNING 2: Choosing an option to create a shortcut on the desktop during installation may cause that this shortcut will not work. Creating shortcut manually (right click on the *FPMapp.exe* and then *create shortcut*) should work properly.

  WARNING 3: Your antivirus might detect that *FPMapp.exe* or *FPMAppInstaller_web.exe* may be dangerous because it comes from an unknown source. Please ignore that message or add the executable files to the exclusions list.

#### 2.3. Inputs and outputs

Inputs – to successfully perform FPM reconstruction you need:

- Input FPM images – images collected by your FPM system (see Chapters 2.4 and 7.2). These images should be stored in separate folder with no other files inside. Image names should be in such an order that sorting them by the name should reflect the order in which they were collected. For example: *Img1, Img2, Img3, …*; *Image_001, Image_002, Image_003, …* or *1,2,3,…*. If you do not have real image data, you can generate synthetic data (Chapter 3.3).
- Knowledge about the following parameters of your real/synthetic imaging system (these parameters should be set manually in main *FPM app* window (see Chapter 3.2)):
    - Spacing between adjacent LEDs in LED array (or other light sources)
    - Distance between LED array and a measured object
    - Pixel size of the camera
    - Central wavelength of the light source
    - Objective lens numerical aperture
    - Magnification of the system
    - LED array layout
    - Order in which images were collected (to associate collected images to appropriate LEDs in LED layout).

**MATLAB version:**

Outputs – after performing reconstruction, outputs will be saved in the MATLAB workspace. Output variables are:

- object – reconstructed complex object
- phase – phase of the object
- pupil – reconstructed complex pupil function
- err – RMS error compared with the input data for each iteration (see Chapter 4.4.3)
- erro – RMS error compared with the known synthetic object for each iteration (if reconstructing synthetic data, see Chapter 4.4.3)
- object_denoised – denoised complex object (if denoising was performed – Chapter 5.4)
- phase_denoised – denoised object phase (if denoising was performed – Chapter 5.4)
- idx_X, idx_Y – positions in X and Y axis in the Fourier domain of collected images (if these positions were corrected, then these corrected positions are saved – Chapter 5.2)

Outputs also can be saved or shown by the *FPM app* functions (Chapter 3.2.3).

**Executable version:**

Outputs – the only access to the outputs variables (MATLAB version description) is to show them through *show results wizard* (Chapter 3.7, object amplitude, phase and both of them denoised can also be exported as .tif images there) or to save them to .mat files (Chapter 3.2.3, access to them through MATLAB).

#### 2.4. How to collect input FPM images

FPM input data consists of multiple microscopic images, each one collected with different illumination angle. The easiest way to collect these images is to use classical microscope system with the LED matrix placed as illuminator (like in Figure 1). With this system in mind we have created our GUI.

To collect data, firstly you need to assemble your hardware. If you plan to build system as in Figure 1, the only thing to consider is a distance between LED matrix and a measured object. This distance should be large enough to achieve around 40-60% overlap [9] of the spectra of adjacent LEDs in the Fourier domain. This overlap can be calculated manually (Chapter 4.2) or you can use *Generate images* window (Chapter 3.3) to simulate your setup and easily check this overlap (exemplary procedure of such simulation is shown in Chapter 7.1).

When you have completed your hardware, you can start collecting images. You need to collect one image per each illuminating diode. It is important that the diodes should illuminate the sample in a proper order – program needs to know in which order you have collected images and has embedded a few basic orders that you can choose from (see *Image collecting order* window – Chapter 3.5). Collected images should also have proper names (Chapter 2.3). Each image should be collected with the same gain and exposure time. It is worth to make sure that no image will be overexposed or underexposed. Exemplary procedure of collecting input FPM images is shown in Chapter 7.2.

### 3. User interface

#### 3.1. How *FPM app* works

To show how the process of working with the *FPM app* should work, the simple graph is depicted in Figure 2. The core of developed app is the certain number of variables (parameters and images) that may be obtained in three different ways. When they are set, the process of reconstruction may begin. After it finishes, reconstruction results are obtained and additional operations on them can be made.

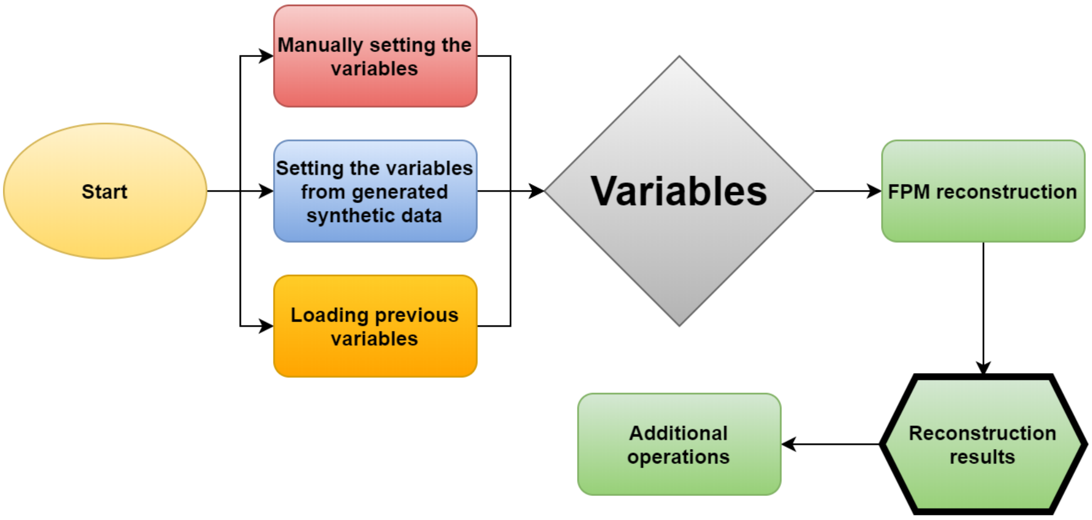

*Figure 2. Simple diagram of the FPM app operation structure.*

*FPM app* has many variables that can be set. These variables are listed below:

- Input FPM images – low-resolution input images: collected by the real or synthetic FPM imaging system.
- Known synthetic object – two images, first representing object amplitude and second representing its phase. This variable is not necessary to run the algorithm. It is only needed to be compared with synthetic data reconstruction result.
- System configuration variables:
    - LED array – matrix that illustrates the shape of the LED array used to collect images.
    - Image collecting order – matrix that illustrates in which order images illuminated by LEDs from the LED array were collected.
    - System setup – configuration of the real/synthetic FPM imaging system:
        - Central wavelength of used illumination source.
        - Objective lens NA.
        - System magnification.
        - Camera pixel size.
        - Spacing between adjacent LEDs in LED array.
        - Distance between LED array and the measured sample.
- Reconstruction variables:
    - Used LEDs – matrix that illustrates which images should be used in the reconstruction (LEDs used to collect these images).
    - Region of interest (ROI) – part of the field of view that will be reconstructed.
    - Reconstruction options:
        - Reconstruction algorithm.
        - Reconstruction order.

- LED correction method.
- Number of iterations.
- Alpha and beta parameters (needed in Quasi-Newton method).
- Usage (or not) GPU to accelerate the reconstruction.

All of these variables are easy to set. For a user that is familiar with the *FPM app*, adjusting all the parameters to a given own setup should take no longer than 2 minutes (not counting the time needed to load few hundreds of input images). *FPM app* is remembering the variables after reconstruction so they do not need to be filled over and over again. Moreover, *FPM app* is even remembering most of the variables (apart from the input FPM images, known synthetic object and ROI) after closing the app which makes using this application more enjoyable.

To make application readable, it was composed of one main window that shows all the variables, and several smaller windows used to perform various operations. Dependences between *FPM app* windows are shown on the graph in Figure 3.

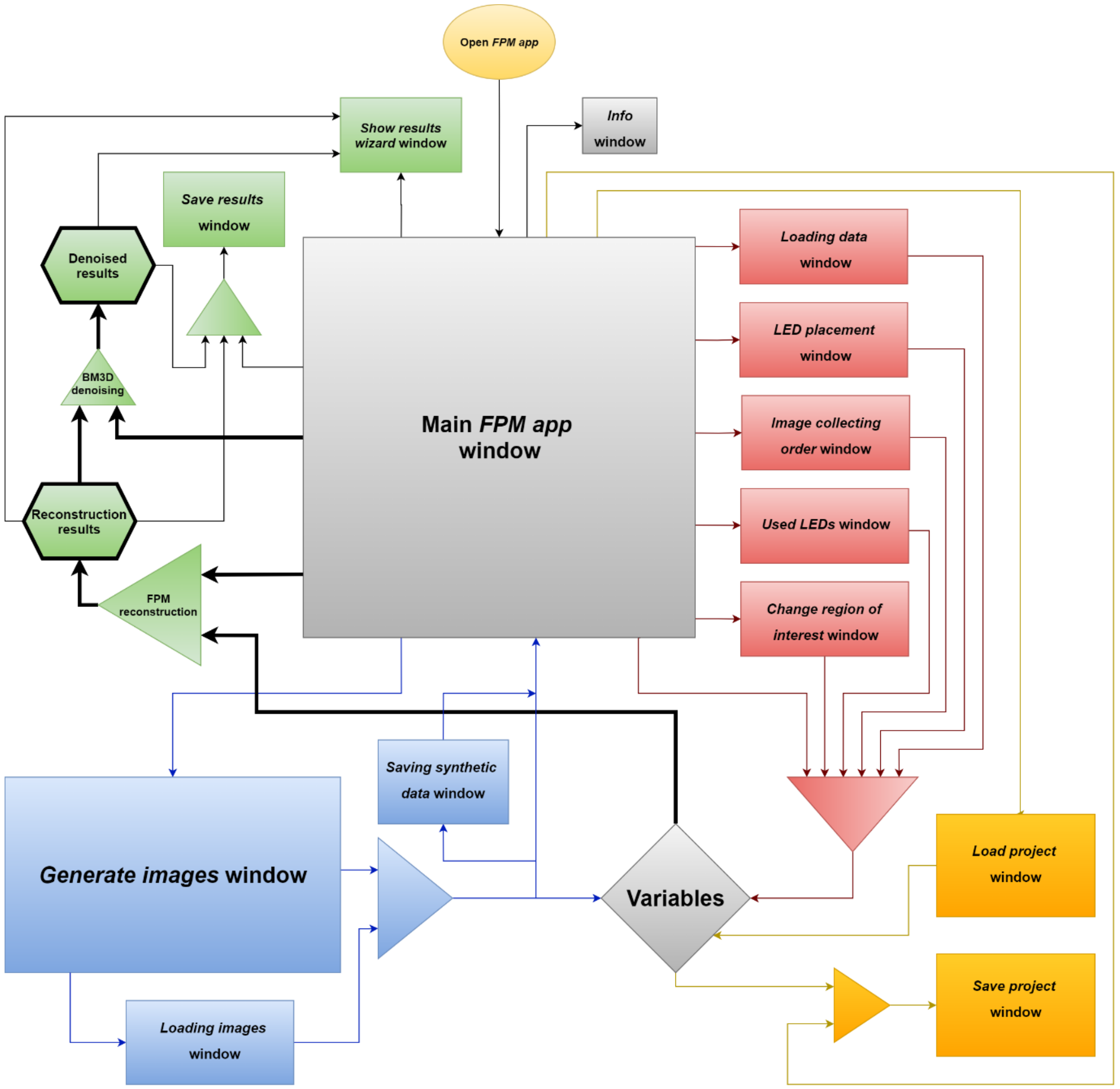

*Figure 3. Dependencies between FPM app windows. Rectangles are representing separate windows generated by the FPM app. Blue path – generating synthetic data; Red path – completing system parameters; Yellow path – saving and loading projects (variables); Green path – reconstruction and output processing.*

There are three ways to obtain *FPM app* variables:

**Manually setting them from the main *FPM app* window (red path in the Figure 2 and Figure 3)** – in this way, firstly, input images need to be loaded. Then imaging system configuration variables (system setup, LED array, image collecting order) need to be changed. At the end, reconstruction variables (ROI, used LEDs, reconstruction options) may be changed to adjust the reconstruction. See Chapter 3.2 for more detailed description of the main *FPM app* window and Chapter 7.3 for exemplary reconstruction process.

**Setting the variables from generated synthetic data window (blue path in the Figure 2 and Figure 3)** – *FPM app* has also possibility to generate synthetic FPM data. In this case, *Generate images* window (Chapter 3.3) may be opened from the main *FPM app* window. In this window firstly, known synthetic object should be simulated by loading two images representing its amplitude and phase. Then synthetic system setup and LED array should be set. After that, input FPM images may be generated. After generation, input images along with the LED array, system setup and image collecting order are set. Reconstruction variables may be still changed in main *FPM app* window.

**Loading previous variables (yellow path in the Figure 2 and Figure 3)** – there is also possibility to load (main *FPM app* window -> File -> load project) previously saved (main *FPM app* window -> File -> save project) variables. In this case all saved variables will be loaded into the app (Chapter 3.2.5).

When all the variables are set, reconstruction may be performed by pressing **Run algorithm** button. During the reconstruction (if the **Show iteration results** checkbox was checked), figure containing currently reconstructed object, pupil and error will pop up, and it will be updating after each iteration.

Once the reconstruction finishes, *Show results wizard* window will open, where some of the reconstruction results may be chosen to be shown in separate figures. Reconstructed object may be further denoised (with the use of the Block-matching and 3D filtering (BM3D) algorithm[12]) and results may be saved – all of this is descripted in-depth in Chapter 3.2.3.

WARNING: Phase that is returned by *FPM app* is wrapped – when imaging samples with large height differences (larger than illumination wavelength) there will be seen phase discontinuities. To remove it you need to use one of the phase unwrapping algorithms.

#### 3.2. Main *FPM app* window

In this window (Figure 4) you can modify algorithm parameters and options, open program sub-windows, load input data and run FPM reconstruction. In this sub-chapter we will describe all *FPM app* window functionalities.

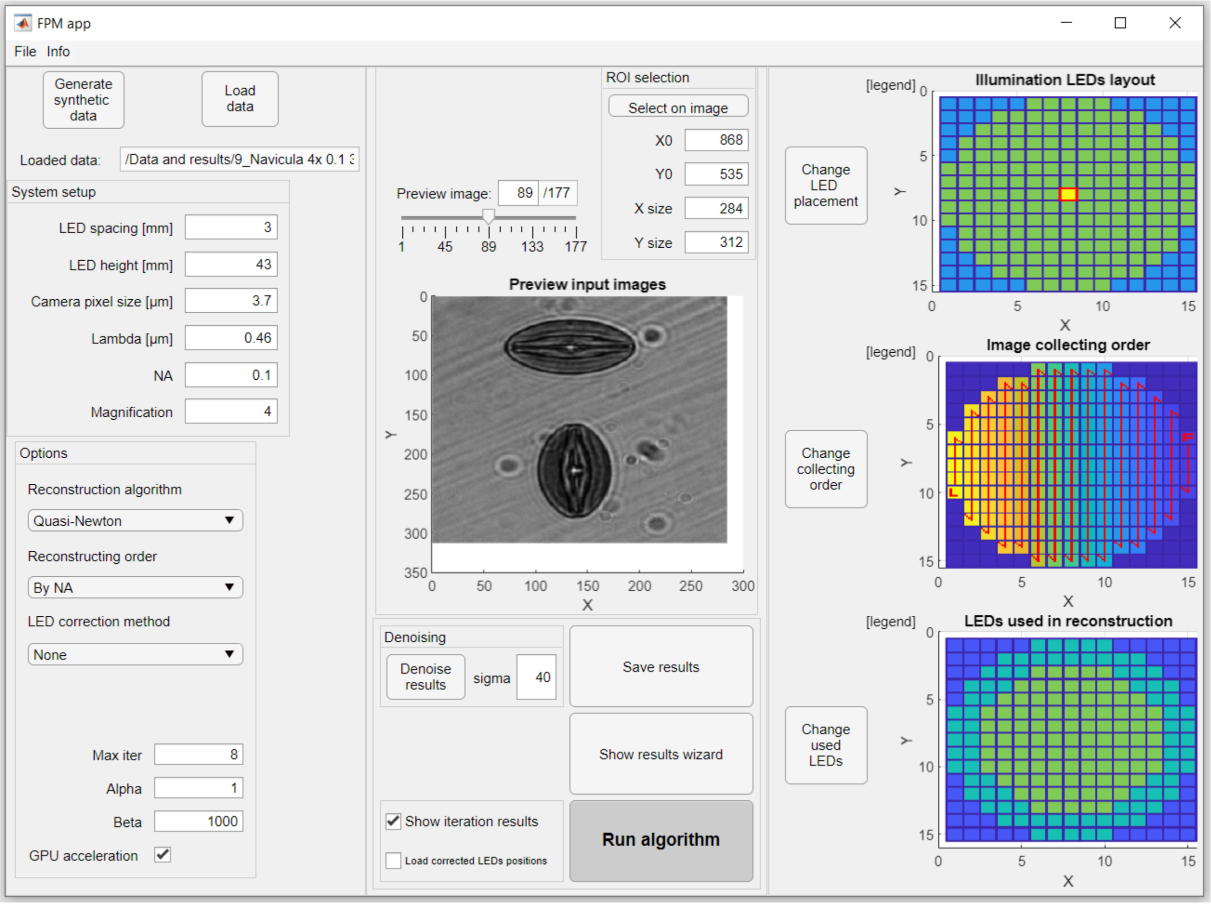

*Figure 4. Main FPM app window.*

##### 3.2.1. Left panel

**Generate synthetic data** button – this button open *Generate images* window (Chapter 3.3) where you can create your own synthetic system and use it to generate synthetic data.

**Load data** button – this button is used to load input FPM images. After pressing it a window will open in which you should select your FPM data folder. It is important that this folder should contain only input images and no other files. When the folder is selected, the images inside it will be loaded into the app. Input images should be named as described in Chapter 2.3. (**MATLAB version** – loaded images may be found in the MATLAB workspace variable *ImagesIn*. You can also check *imageList* workspace variable to be sure that images were loaded in correct order).

**Loaded data** field (below **Load data** button) shows which data was loaded – last 2 parts of the load directory path or the "Generated data" information (Figure 5).

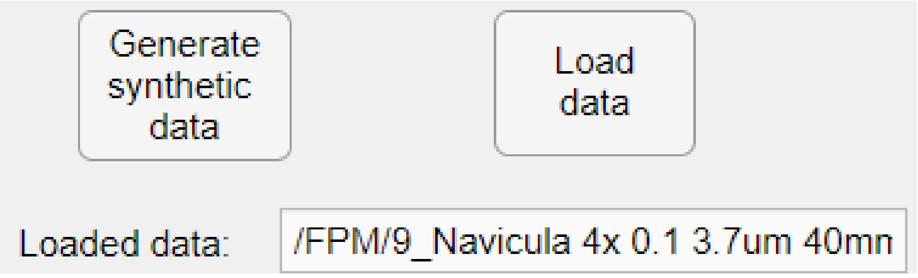

*Figure 5. Information about which data was loaded.*

**System setup** panel (Figure 6) – In this panel you need to write parameters of your real FPM system:

- LED spacing [mm] – spacing between adjacent LEDs in the LED array,
- LED height [mm] – distance between LED array and a probe,
- Camera pixel size [um] – pixel size of the camera,
- Lambda [um] – central wavelength of the LEDs,
- NA – objective lens numerical aperture,
- Magnification – real magnification of the system.

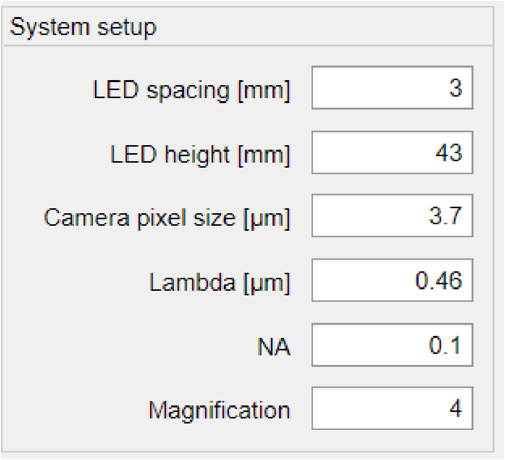

*Figure 6. System setup panel with exemplifying values typed in.*

Detailed explanation of the impact of these parameters onto the reconstruction is described in Chapter 4.1.

**Options** panel (Figure 7) – In the options panel you can select some of the reconstruction options:

- **Reconstruction algorithm** – select one of two implemented reconstruction algorithms
  - Quasi-Newton – implemented as in [13]. Implementation was based on Lei Tian algorithm [11].
  - Gerchberg-Saxton – implemented as in [4].

  Quasi-Newton algorithm should give better results (especially for reconstruction of the real data) and additionally retrieve systems pupil function. Detailed information about the working principle of these algorithms is presented in Chapter 4.4.
- **Reconstructing order** – select the reconstruction order in which object Fourier spectrum will be updated:
  - By NA – from the images collected by the central LED to the images collected by the edge LEDs.
  - By Intensity – from the brightest to the darkest collected images.

  In general, both of these reconstruction orders should give similar results, except for reconstruction with small number of iterations, where intensity order should perform better
- **LED correction method** – there are implemented 4 LED correction algorithms, that can amend for misalignment error (error resulting with reconstructed images disturbed by artifacts in the form of fringes that occurs when illumination angles calculated from system setup differs from real illumination angles). Due to the shortest computing time we recommend **Authors method [fast]** algorithm, although other algorithms may give slightly better results. More information about misalignment error algorithms may be found in Chapter 5.2. Whereas Chapter 7.3 shows exemplary procedure of the usage of the LED correction algorithms.

- **Max iter** – number of iterations. The more iterations, the better is the reconstruction quality but also it takes longer time. The optimal number of iterations is usually between 4 and 10 but it depends on the input data. Error plots that appear during and after the reconstruction may help to set it optimally for the following reconstruction.
- **Alpha** – regularization parameter for object reconstruction in the Quasi-Newton algorithm. Alpha in the range 0.1-10 should give correct result. More information about impact of this parameter can be found in Chapter 4.4.1.
- **Beta** – regularization parameter for pupil reconstruction in the Quasi-Newton algorithm. Beta larger than 0 should give correct result. More information about impact of this parameter can be found in Chapter 4.4.1.
- **GPU acceleration** – when this option is checked, some of the reconstruction algorithm operations are performed on GPU. To use GPU, you need:
    - **MATLAB version** – Nvidia's GPU with CUDA driver version supported by your MATLAB version. See [https://uk.mathworks.com/help/parallel-computing/gpu-support-by-release.html] to know which version of CUDA driver you will need.
    - **MATLAB version** – MATLAB Parallel Computing Toolbox
    - **Executable version** – Nvidia's GPU with CUDA driver version 10.0 or higher

  GPU may significantly increase the speed of the computing algorithm on the large data (approximately for ROI larger than 200x200). For smaller ROIs CPU may be faster than GPU. Using GPU requires storing in the memory a 3D double matrix of all images (images of the size of the ROI). E.g., GPU with the 4GB available video memory can maximally perform reconstruction for data consisting of 300 images with the ROI around 900x900. In the case of not enough memory to perform reconstruction on given ROI, corresponding error will pop up after pushing **Run algorithm** button.

  To reconstruct larger data with the use of the GPU, one of the approaches is to reconstruct several smaller ROIs and then stitch the results. More information about performed GPU optimization may be found in Chapter 5.3.

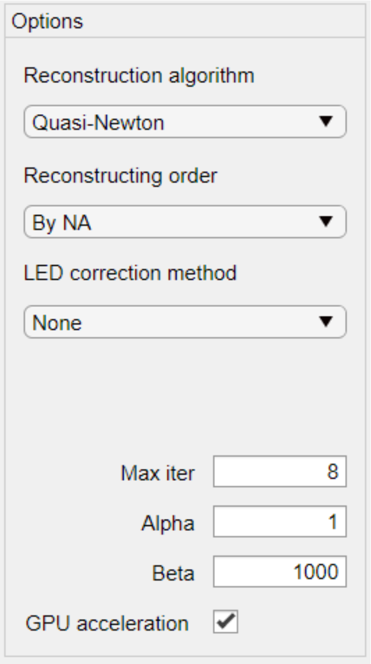

*Figure 7. Options panel*

##### 3.2.2. Top middle panel

In this panel (Figure 8) you can select which of the input images will be shown in the **Preview input images** field. You can select it by typing image number in the **Preview image** field or by moving the slider below it. Image numbering is in the order in which images were loaded into the *FPM app*. It is worth of notice that LED used to collect previewed image is marked with a red rectangle on a **Illumination LED layout** field (Figure 12).

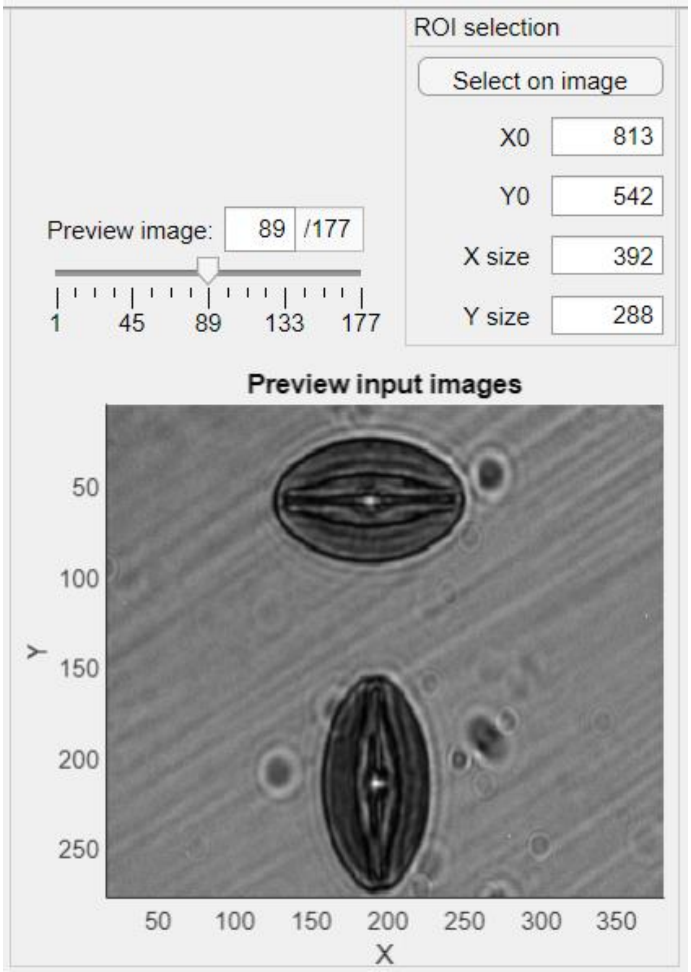

*Figure 8. Top middle panel of the FPM app main window.*

**ROI selection** panel – here you can select Region Of Interest (ROI) you want to reconstruct (also this ROI will be shown in the *Preview input images* image field). When want to perform large FOV reconstruction we recommend to firstly try reconstruction on a smaller ROI to quickly check if there is everything set correctly and outcomes are feasible.

**Select on image** button will open center image (image collected with center diode) in the *ROI selection* window (Figure 9), where you can select ROI with the use of mouse (click on the image and drag a mouse to draw a rectangle, then double click inside rectangle to create ROI). Center image is chosen accordingly to LEDs layout (Chapter 3.4) and Image collecting order (Chapter 3.5) so please do complete these inputs first if you want to select ROI on the center image.

ROI can be also selected by changing **X0**, **Y0**, **X size** and **Y size** fields. **X0** and **Y0** are the coordinates of the top left ROI pixel, **X size** and **Y size** are the ROIs size (Figure 9).

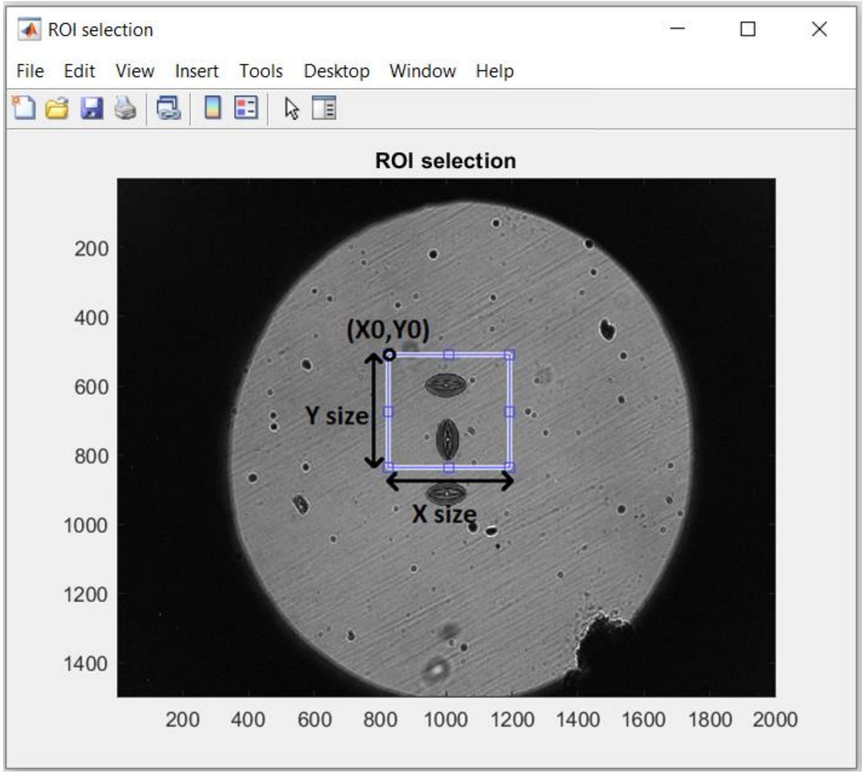

*Figure 9. ROI selection window.*

##### 3.2.3. Bottom middle panel

**Run algorithm** button – push this button to start reconstruction algorithm.

If the **Load corrected LEDs positions** check box is checked, at the beginning of the reconstruction you will need to point previously saved .mat file with reconstruction results (file created with **save results** button). It will make that LEDs positions used to create .mat file results will be used in a current reconstruction (scaled to appropriate ROI size). This function was developed to avoid performing multiple times memory-consuming LED position correction algorithms on the same data (once corrected, LED positions may be used to perform another reconstruction on the same data, e.g., on another ROI).

If the **Show iteration results** check box is checked, after every reconstruction iteration there will be shown *Iteration results* window (Figure 10) with currently reconstructed object amplitude and phase along with pupil function and errors (when reconstructing huge FOV, displaying this window may cause to slow down the reconstruction time).

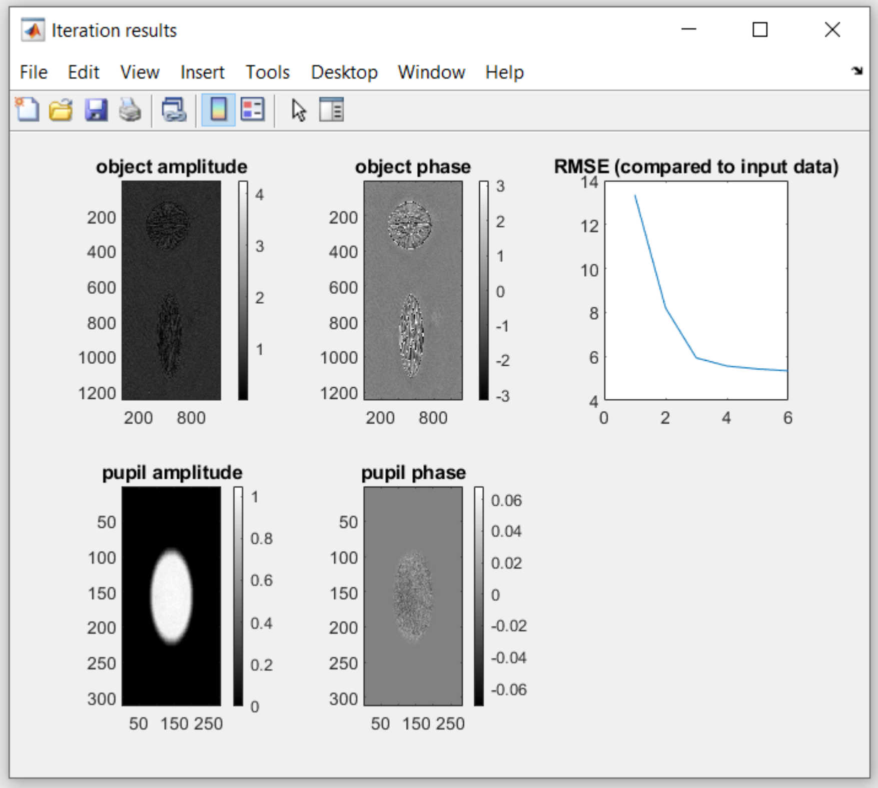

*Figure 10. Iteration results window.*

Once the reconstruction completes, preview figure (if exist) will close and *Show results wizard* window will pop-up, where reconstruction results can be shown. This window may be also opened by pushing **Show results wizard** button (Chapter 3.7).

Reconstruction results may also suffer from some low-level noise. To partially eliminate it, you can try to digitally remove it by pressing **Denoise results** button. It will perform BM3D denoising on the reconstructed object with given sigma value (see Chapter 5.4). Sigma is a denoising parameter (values from 0 to 255). The higher the sigma value is, the stronger the denoising performed. We recommend to set sigma smaller than 50 (start from 5-10 in case of detail-rich objects).

To save the results you need to press **Save result** button, which will open a separate window where you can create the .mat file in which the below results will be saved (if they exist):

- Reconstructed complex object
- Reconstructed phase
- Reconstructed complex pupil
- RMS error compared to input data in function of iteration number
- RMS error compared to known synthetic object in function of iteration number
- Denoised complex object
- Denoised phase
- Corrected LEDs positions

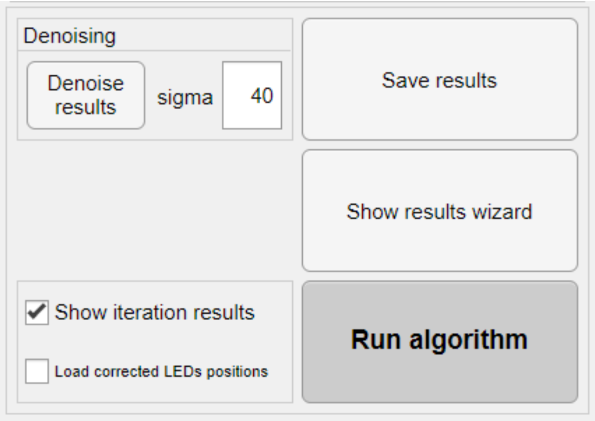

*Figure 11. Bottom middle panel of the FPM app main window.*

##### 3.2.4. Right panel

In the right panel (Figure 12) are placed three buttons which are used to open external windows:

**Change LED placement** button – it opens *LED placement* window (Chapter 3.4) where you can create LED array used to collect images.

**Change collecting order** button – it opens *Image collecting order* window (Chapter 3.5) where you can choose in which order images were collected (to associate correct LED in the LED array to appropriate collected image)

**Change used LEDs** button – it opens *Used LEDs* window (Chapter 3.6) where you can choose which images you want to use in the reconstruction.

Next to each of these buttons there is a small preview image, which illustrates current LED layout/image collecting order/LEDs used in the reconstruction.

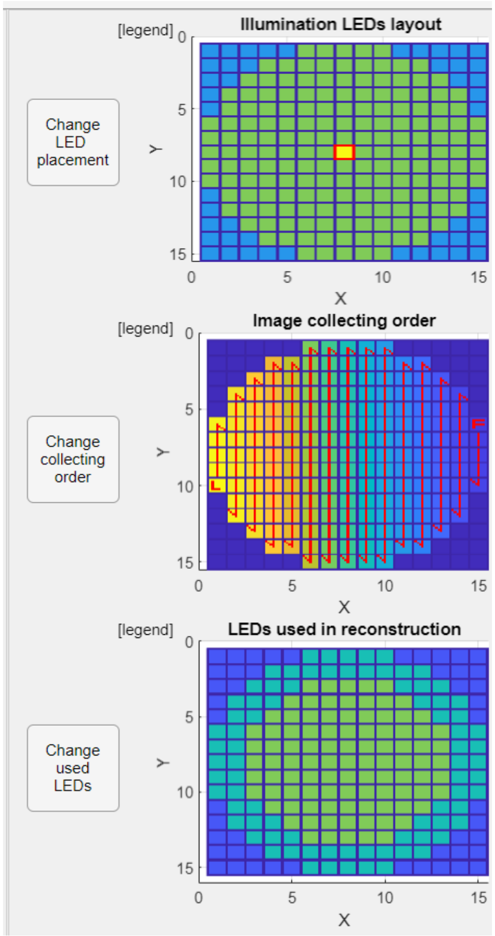

*Figure 12. Right panel of the FPM app main window.*

##### 3.2.5. Menu bar

**File->save project** – save currently used FPM settings into the .mat file. The following *FPM app* variables will be saved:

- System setup
- Reconstruction options
- LED array layout
- Image collecting order
- LEDs used in reconstruction
- Known synthetic object (if exist)
- Data directory (if exist).

**File->load project** – load previously saved project (*FPM app* variables) into the app. The following *FPM app* variables will be loaded:

- System setup
- Reconstruction options
- LED array layout
- Image collecting order
- LEDs used in reconstruction
- Known synthetic object (if exist, input FPM data will be automatically generated)
- Data directory (if exist, input FPM data will be automatically loaded into the app).

#### 3.3. *Generate images* window

*Generate images* window (Figure 13) is used to simulate synthetic FPM system and uses it to generate synthetic FPM data.

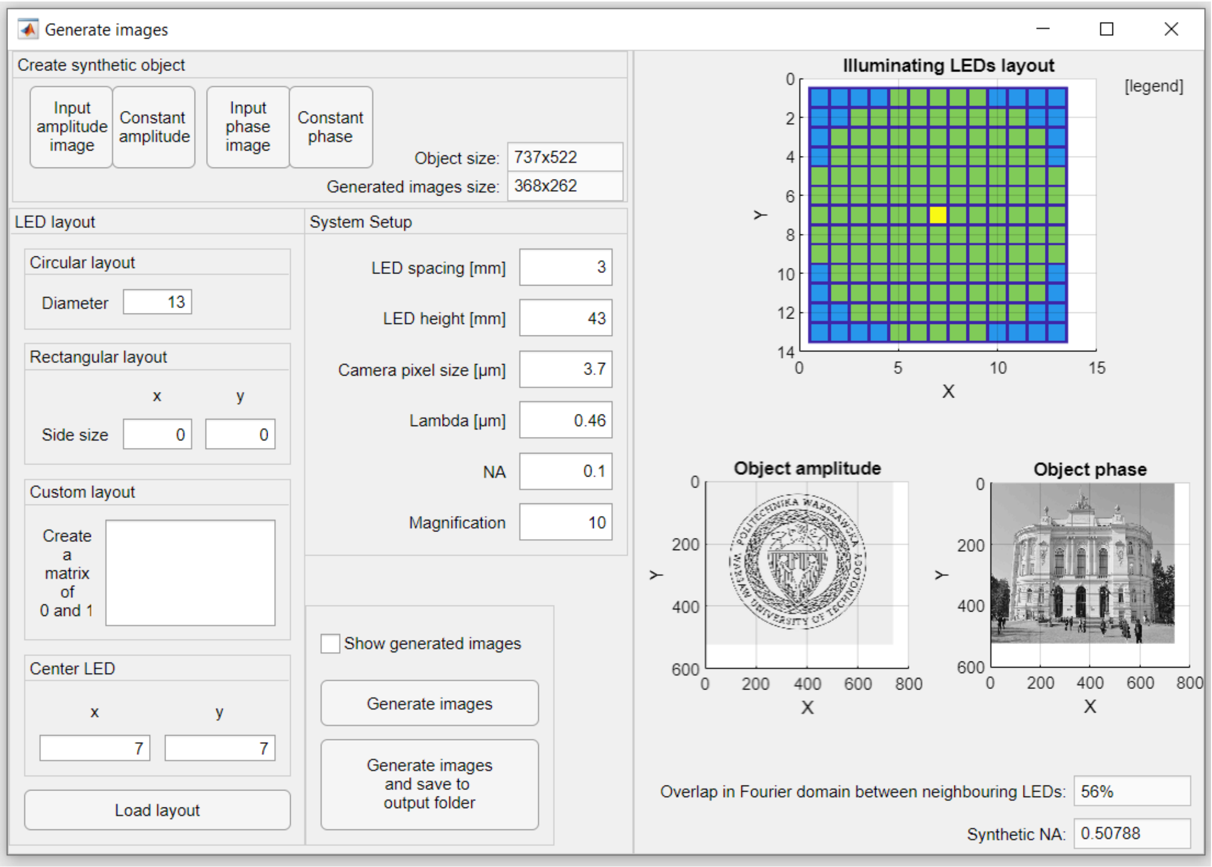

*Figure 13. Generate images window.*

**Create synthetic object** panel – in this panel known synthetic object (ground truth) can be created. This object is simulated merging two images: one that constitutes its amplitude and one that comprise its phase. These are images you need to load by pressing **Input amplitude image** and **Input phase image** buttons. Amplitude and phase images are being resized to match each other size. There are also buttons allowing to quickly place constant value as an amplitude or phase (**Constant amplitude** and **Constant phase** buttons). Next to these buttons, there are fields that show the size of the object and the future size of the generated images.

**LED layout** panel – in this panel you can create your synthetic system illumination layout. Principle of the operation in this panel is the same as in *LED placement* window (see Chapter 3.4).

**System setup** panel – in this panel you can set your synthetic system parameters. Principle of the operation in this panel is the same as in **System setup** panel in main *FPM app* window (see Chapter 3.2.1).

At the bottom of the **left panel** there are buttons responsible for generating input FPM images (**Generate images** button) or generating them and additionally saving them into the output directory (**Generate images and save to output folder** button, output directory may be selected after pressing this button). There is also checkbox that gives the possibility to show or not the images during the reconstruction (Figure 14). Showing the images during their generation will slow down its time.

After image generation, *Generate images* window will close and generated images will be loaded into the *FPM app* along with the LED layout, system setup parameters and image collecting order.

In the **right** panel the LED array used to generate images (see Chapter 3.4) is shown along with images illustrating object amplitude and phase. At the bottom of this panel there are fields displaying the overlap value in Fourier domain between images collected by adjacent LEDs and the synthetic aperture of the system which are useful parameters when designing your own FPM system (see Chapter 4.2). To achieve the best quality results (both in synthetic and real systems) we recommend designing systems with overlap in the range from 40 to 60%.

While working in *Generate images* window, there is also shown *Positions of the images centers* window in which current spectrum of the object along with current positions of the centers of collected images in this spectrum are shown (Figure 15). This window may be helpful in creating your own custom made synthetic system to adjust the LED array and the system setup to obtain optimal sampling in the Fourier domain and desired synthetic numerical aperture. This tool may be also used to plan the setup of real imaging system (see Chapter 7.1).

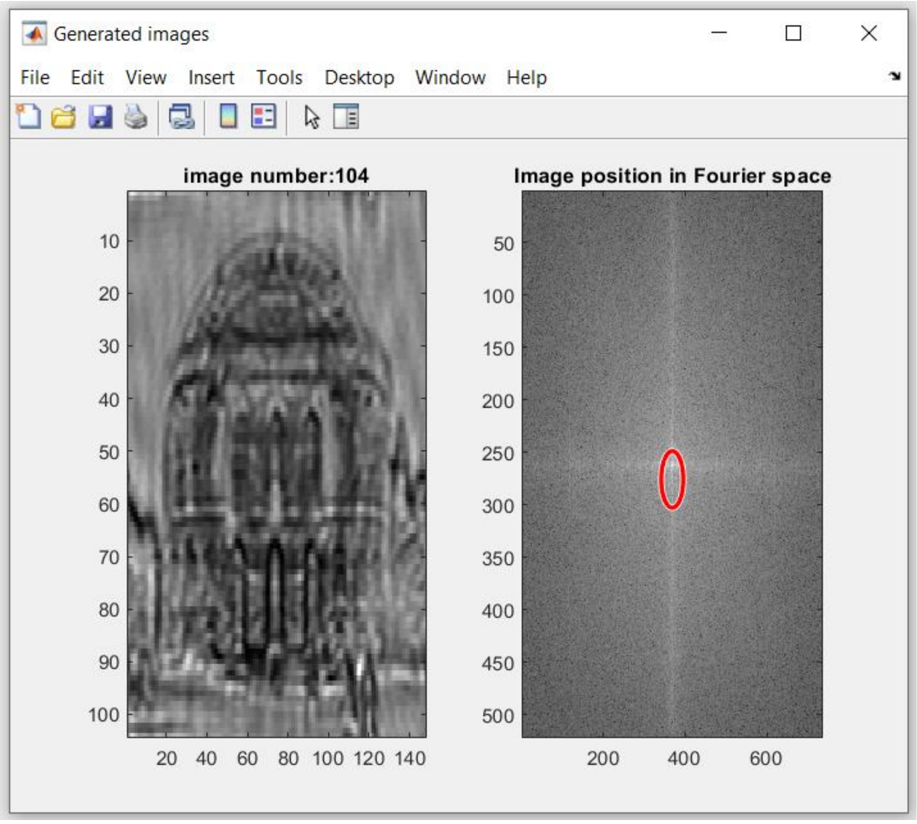

*Figure 14. Generated images window.*

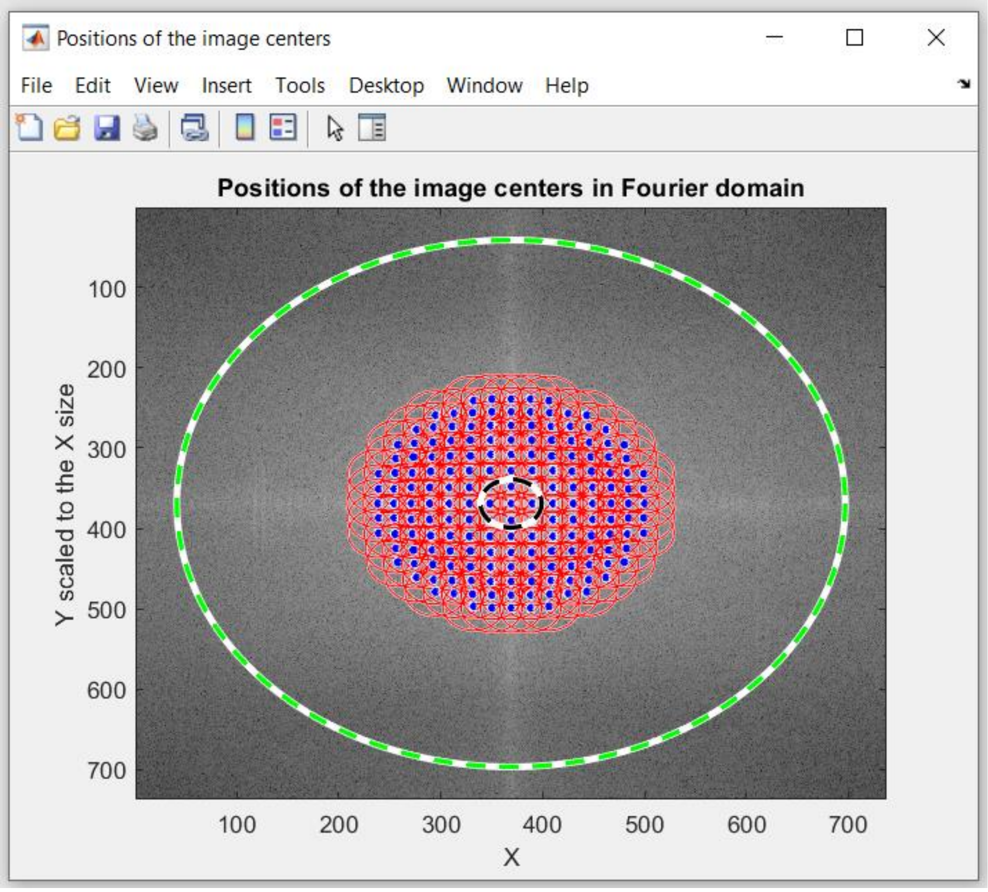

*Figure 15. Position of the image centers window. Blue dots mark image centers, red circles mark spatial frequencies stored in images, black circle mark frequencies stored in central image and green circle mark maximal synthetic NA possible to obtain with this system (when illumination NA equals 1).*

#### 3.4. *LED placement* window

*LED placement* window (Figure 16) is used to create the shape of the LED array used to collect the input FPM images. This window contains two panels:

**Right** panel – in this panel the image containing current LED array is shown. In this image squares separated by dark-blue lines are depicted. Each square is representing one LED spot. Blue LED represents empty space/LED not used to collect images, green square represents LED used to collect images and yellow square represents the central LED – LED that lays on the objective optical axis.

**Left** panel – in this panel LED array can be created in four ways. First way is to create circular array, by typing number of LEDs that lay on the diameter of an array into the **Diameter** edit field. Second way is to create rectangular array by typing number of LEDs constituting both of the rectangle arms into the **x Side size** and **y Side size** edit fields. Third way is to use **Custom layout** panel to create a custom array manually by typing ones and zeros: 0 – selected LED is OFF, 1 – selected LED is ON, 2 – marks center LED. Such matrix can also be loaded to the app using **Load layout** in the form of .mat file. Center LED also can be pointed by typing its coordinates into the **Center LED** panel.

To confirm the LED array generated in this window, press **Set this layout** button, which will load this layout into the *FPM app*.

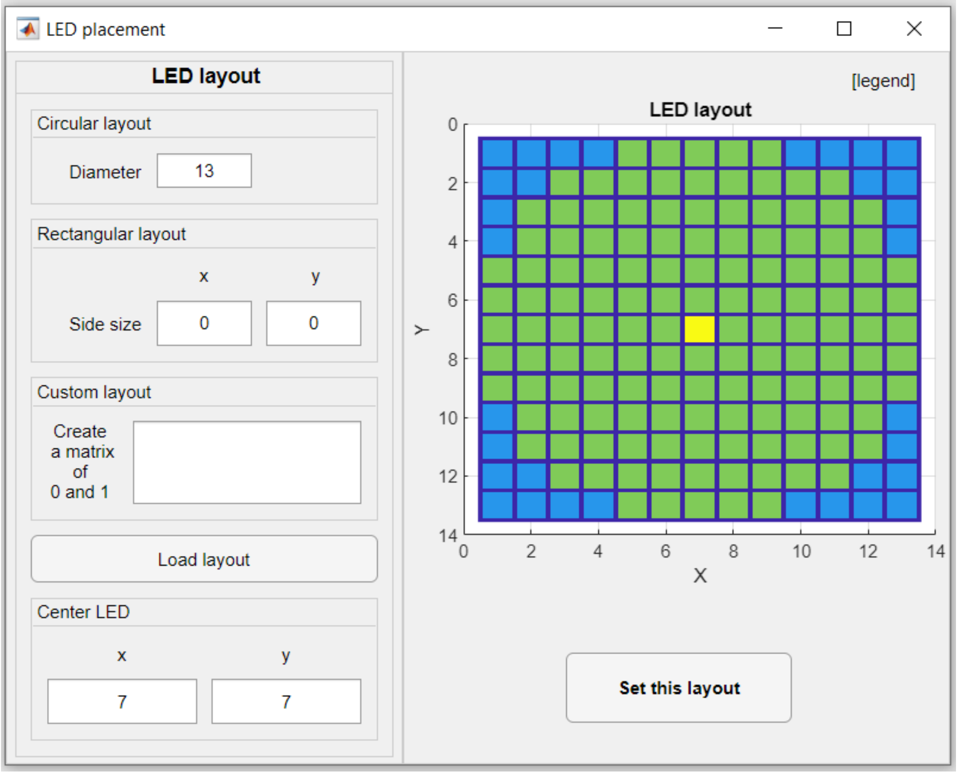

*Figure 16. LED placement window.*

#### 3.5. *Image collecting order* window

This window (Figure 17) is used to select in which order images were collected in your FPM system (in which order LEDs have been turning on in the LED matrix). It is necessary to associate correct image in your images directory to the proper LED in the LED array.

**Right** panel – this panel is showing the LED turning-on order. It is composed of squares separated by dark-blue lines. Each square represents one LED or empty spot in the LED array. Dark-blue squares represent LEDs not used to collect images or empty spots. Blue square represents LED used to collect first image, every next LED is of slightly lighter color than previous, up to the square representing LED used to collect last image, which is yellow. There is also red line that goes through the all squares in selected order. When the LED used to collect current image do not lay next to the LED used to collect next image, red line point in the direction where the square representing next LED should be. Also, the red letters F and L mark the squares representing LEDs used to collect the first and the last image.

**Left** panel – on the left panel image collecting order can be selected. In case of line by line, row by row or snake order, one should start by selecting, first image from the **First image** panel. Then, desired image acquisition order should be selected from **Collecting order** panel which will immediately show current collecting order on the right panel. In case of the spiral order, image collected by the central LED will be set as the first image so first image do not need to be selected. Only in the case of spiral acquisition clockwise or counter-clockwise order should be selected. There is also **Change spiral** button that changes the starting direction of the spiral, e.g. when the spiral starts by going down, by pressing **Change spiral** button it will make spiral starting in right direction. Another click of this button will make this spiral to start in up direction and so on.

To confirm the image collecting order created in this window, press **Set collecting order** button, which will load this order into the *FPM app*.

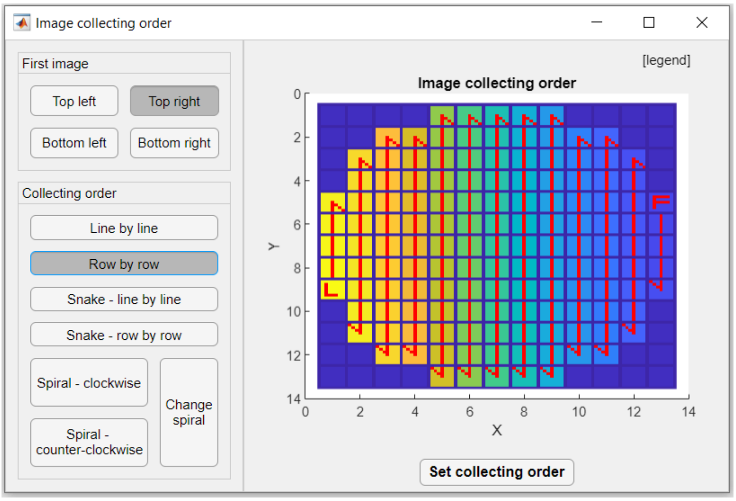

*Figure 17. Image collecting order window.*

#### 3.6. *Used LEDs* window

*Used LEDs* window (Figure 18) is used to choose which images (collected by which diodes) will be used in the reconstruction. This window consists of two panels:

**Right** panel – on this panel image of the LEDs used in the reconstruction is shown. This image is composed of squares separated by dark-blue lines. Each square represents one LED or empty spot in the LED array. Dark-blue squares represent LEDs not used to collect images or empty spots. Light blue squares represent LEDs used to collect images but not to be used in the reconstruction. Green squares represent LEDs used to collect images that will be used in the reconstruction.

**Left** panel – on this panel, images that will be used in the reconstruction may be selected. There are three automatic options – **All**, **Brightfield** and **Darkfield** checkboxes that will automatically select corresponding images (all, only brightfield or only darkfield images). Moreover, these images may be selected manually, by turning two parameters: *R min* and *R max* that corresponds to minimal and maximal number of LEDs from the LED array center respective. Images collected by all LEDs between *R min* and *R max* parameter will be used in reconstruction. The *R min* and *R max* parameters may be changed in **R min** and **R max** fields or by corresponding sliders.

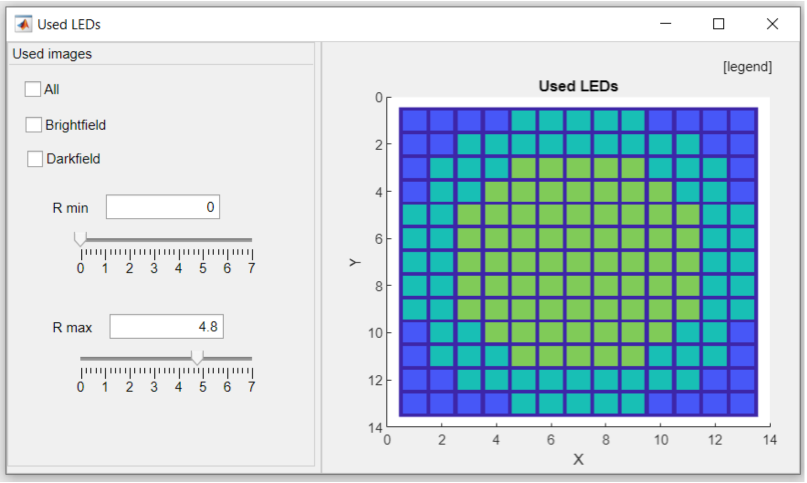

*Figure 18. Used LEDs window*

#### 3.7. *Show results wizard* window

This window (Figure 19) is used to display the reconstruction results. It is automatically opened after the reconstruction or denoising finishes. It also can be opened manually by pressing **Show results wizard** button in main *FPM app* window.

In this window results may be opened or closed by selecting or deselecting corresponding buttons:

- Reconstructed object amplitude
- Reconstructed object phase
- Denoised object amplitude (if denoising was performed)
- Denoised object phase (if denoising was performed)
- Central input image (collected by the central LED)
- Reconstructed pupil function amplitude
- Reconstructed pupil function phase
- Error compared to input images
- Error compared to known synthetic data (if reconstructing synthetic data).

Additionally, reconstructed object and denoised object (both amplitude and phase) have additional options to (Figure 20):

- Narrow down the displaying values range by manipulating the corresponding sliders or fields (this function may be useful when want to increase the contrast of displayed image)
- Change the color map of displayed image
- Display or hide the color bar.

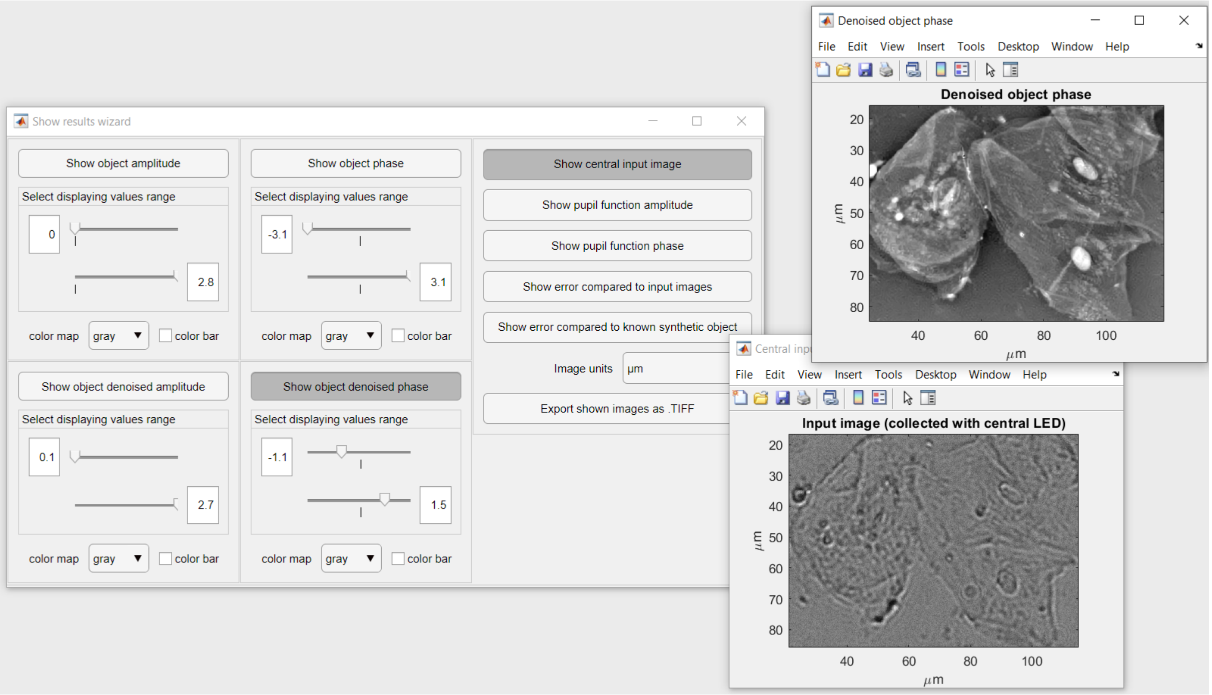

*Figure 19. Show results wizard window with exemplary results displayed by this wizard.*

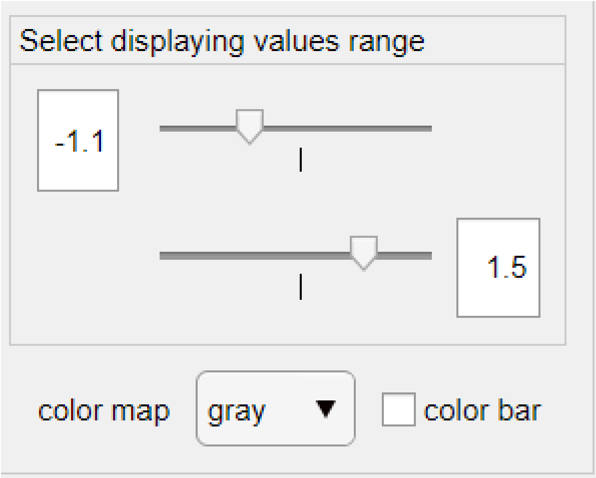

*Figure 20. Additional options in displaying images: narrowing down displayed values range, changing color map and displaying color bar.*

Displayed images units may be adjusted in **Image unit** dropdown menu.

Displayed images (the ones that have their buttons pushed) may be exported as .tif images into the selected directory by pressing **Export shown images as .TIFF** button. Image name will be in the following convention: *DatanameDateType.tif*, where *Dataname* is the last part of data directory path, *Date* is the current date (day-month-year hour-minute) and *Type* is image type (e.g. object amplitude denoised).

### 4. Methods used for the processing of FPM data

In this chapter, all mathematical operations that stands behind *FPM app* are described.

#### 4.1. Impact of System setup parameters

In main *FPM app* window (Chapter 3.2) and *Generate images* (Chapter 3.3) windows the **System setup** panel is placed, where you can set your system settings. They are used to calculate several parameters that FPM algorithm is taking as inputs. Calculating of these input parameters is described in this chapter.

##### 4.1.1. Size of the reconstructed object

Reconstructed object has much better resolution than input images so it's size also must be much bigger.

Size of the reconstructed image in x direction ($S_x$) is calculated as:

$$S_x = ceil\left(\frac{2 \cdot round\left(\frac{2 \cdot MSF_x}{spl_x}\right)}{S_{x0}}\right) \cdot S_{x0}, \tag{1}$$

where $MSF_x$ (equation (2)) is the maximal spatial frequency in x direction (times 2 to take into account negative frequencies), $spl_x$ is the sampling size in the x direction and $S_{x0}$ (equation (3)) is the size of the input images in x direction. Term $ceil\left(\frac{1}{S_{x0}}\right) \cdot S_{x0}$ is to ensure that $S_x$ will be a multiple of $S_{x0}$ to obtain no Fourier transform artefacts.

$$MSF_x = \frac{\max(NA_{ill})}{\lambda} + \frac{NA}{\lambda}, \tag{2}$$

where NA is the objective numerical aperture, $\max(NA_{ill})$ is maximal illumination NA and $\lambda$ is a central wavelength of the used light source.

$$spl_x = \frac{1}{S_{x0} \cdot pixObj}, \tag{3}$$

where $pixObj$ is the pixel size at the object plane:

$$pixObj = \frac{pixCam}{mag}, \tag{4}$$

where $pixCam$ is the pixel size of the camera, $mag$ is the magnification of the system.

Object size in y direction is calculated analogously as in x direction.

##### 4.1.2. LEDs position in the Fourier domain

Position of the image in Fourier domain depends on the illumination angle. For the image illuminated by n-th diode, its position in Fourier domain in x direction ($f_{xn}$) is equal:

$$f_{xn} = \frac{\sin(\theta_{xn})}{\lambda \cdot spl_x}, \tag{5}$$

where $\theta_{xn}$ is the angle of illumination introduced by n-th diode (in x direction):

$$\sin(\theta_{xn}) = \frac{x_n}{\sqrt{{x_n}^2 + h^2}}, \quad (6)$$

where $x_n$ is a distance from the n-th diode to the center diode (in x direction) – directly calculated from *FPM app* LED spacing parameter and LEDs array layout. $h$ is a distance between the LED array and a sample.

LEDs positions in y direction ($f_{yn}$) are calculated analogously as in x direction.

##### 4.1.3. Pupil function size

Pupil function determines which and how good certain spatial frequencies are transported through the system. In x-y plane it has an ellipsoidal shape, defined by two half-axes: $r_x$ and $r_y$ calculated as in the equation (7):

$$r_{x,y} = \frac{NA}{\lambda \cdot spl_{x,y}}. \quad (7)$$

Half-axe in y direction is calculated analogously as in x direction. It is worth of notice that:

$$\frac{r_x}{S_{x0}} = \frac{r_y}{S_{y0}}. \quad (8)$$

In *FPM app*, the cross-section of the pupil function is in a shape of the Tukey window with cosine fraction equal 0.25. This Tukey window (Figure 21) reaches 0.5 of its maximal value in $r_y$ and $r_x$ distance.

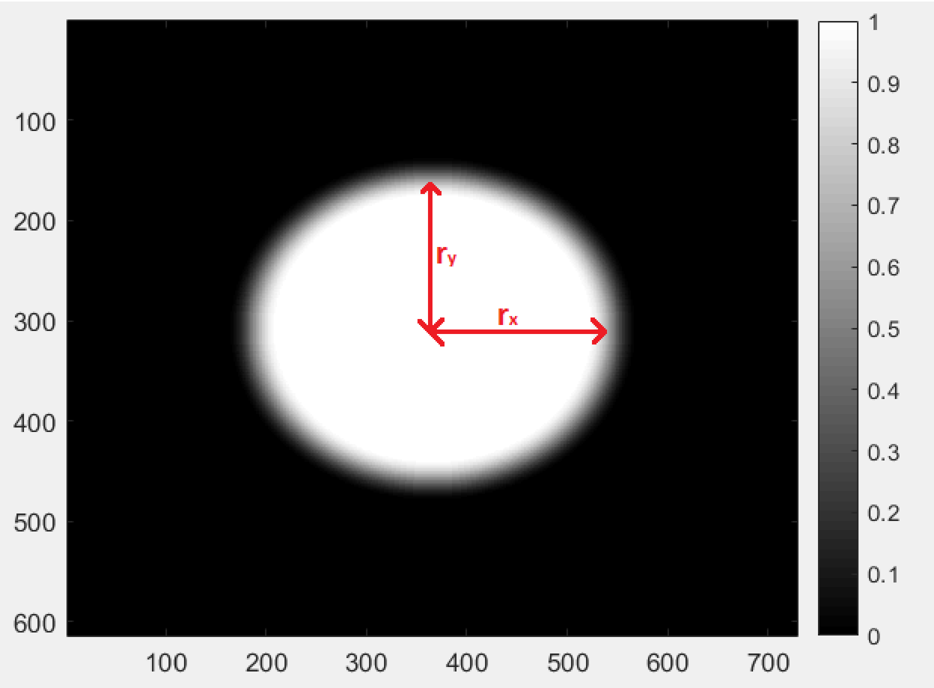

*Figure 21. Amplitude of initial pupil function.*

#### 4.2. Calculation of the overlap and synthetic aperture of the system

While working with FPM it is worth to know the spectrum overlap in the Fourier domain between images collected by adjacent LEDs. Optimal overlap value usually is in a range 40-60%, and when creating your own imaging system it is worth to stick to this value. Spectrum overlap ($ovr$) is calculated by *FPM app* like in equation (9) and this value can be seen in *Generate images* window (Chapter 3.3).

$$ovr = \frac{CA}{PA} \cdot 100\%, \tag{9}$$

where $PA$ is the pupil function area, $CA$ is a common area in the Fourier domain between the pupil function in $(f_{xc}, f_{yc})$ position and pupil function in $(f_{xc+1}, f_{yc+1})$ position, $(f_{xc}, f_{yc})$ is position of center of image collected with a center LED and $(f_{xc+1}, f_{yc+1})$ is position of image collected with LED adjacent to center LED (Figure 22).

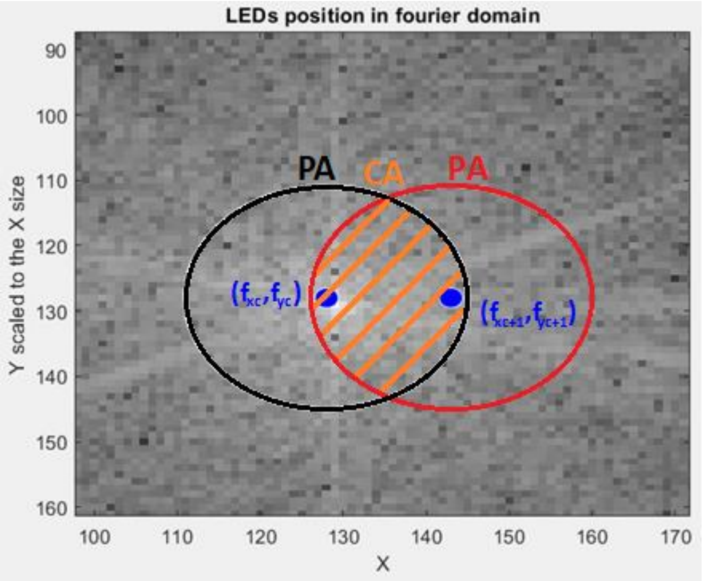

*Figure 22. Illustration of CA and PA areas.*

Another worth to know parameter of FPM system is its synthetic numerical aperture ($NA_{synth}$), which indirectly gives information about reconstructed image resolution. Knowledge about system $NA_{synth}$ may be useful when presenting FPM results. $NA_{synth}$ is also calculated by *FPM app*. This value is shown in *Generate images* window, also in **MATLAB version** it is displayed after reconstruction in the command window. It is calculated as following:

$$NA_{synth} = \max(NA_{ill}) + NA. \tag{10}$$

#### 4.3. Generating synthetic data

Input synthetic FPM images are created from known synthetic object ($obj_{syn}$). This object is simulated by 2 images: one that represents object amplitude ($obj_{amp}$) and another one that represents object phase ($obj_{phs}$):

$$obj_{syn} = obj_{amp} \cdot \exp(i \cdot obj_{phs}). \tag{11}$$

These images are provided by the user. Due to the fact that synthetic object has a finite resolution, it is difficult to determine what the resolution of the synthetic input images should be. Of course, it should be lower than the resolution of the synthetic object, but there is no rule how much lower it should be. In *FPM app*, it was assumed that the input images should have such size that after reconstruction, reconstructed object will have the same size as the known synthetic object – to facilitate comparison of the reconstructed object with the ground truth. Therefore, input image size is calculated with the use of equation (1).

When the input image resolution is known, pupil function ($P$) and LEDs position in the Fourier domain are being calculated. After that, input images ($I_n$) are generated in the way like in the formula below:

$$I_n = \left| F^{-1}\left( crop\left(F\left(obj_{syn}\right)\right)_{S_0 f_{xn} f_{yn}} \cdot P \right) \right|^2, \qquad (12)$$

where term $crop\left(F\left(obj_{syn}\right)\right)_{S_0 f_{xn} f_{yn}}$ represents Fourier transform of the object cropped to the $S_0$ size with the center of the cropped region in $(f_{xn}, f_{yn})$ point, $(f_{xn}, f_{yn})$ positions are calculated as in equation 5.

#### 4.4. FPM reconstructing schemes

Reconstruction process depends on many variables, especially on the type of the reconstruction algorithm. below there is shown a diagram that illustrates the process of the reconstruction:

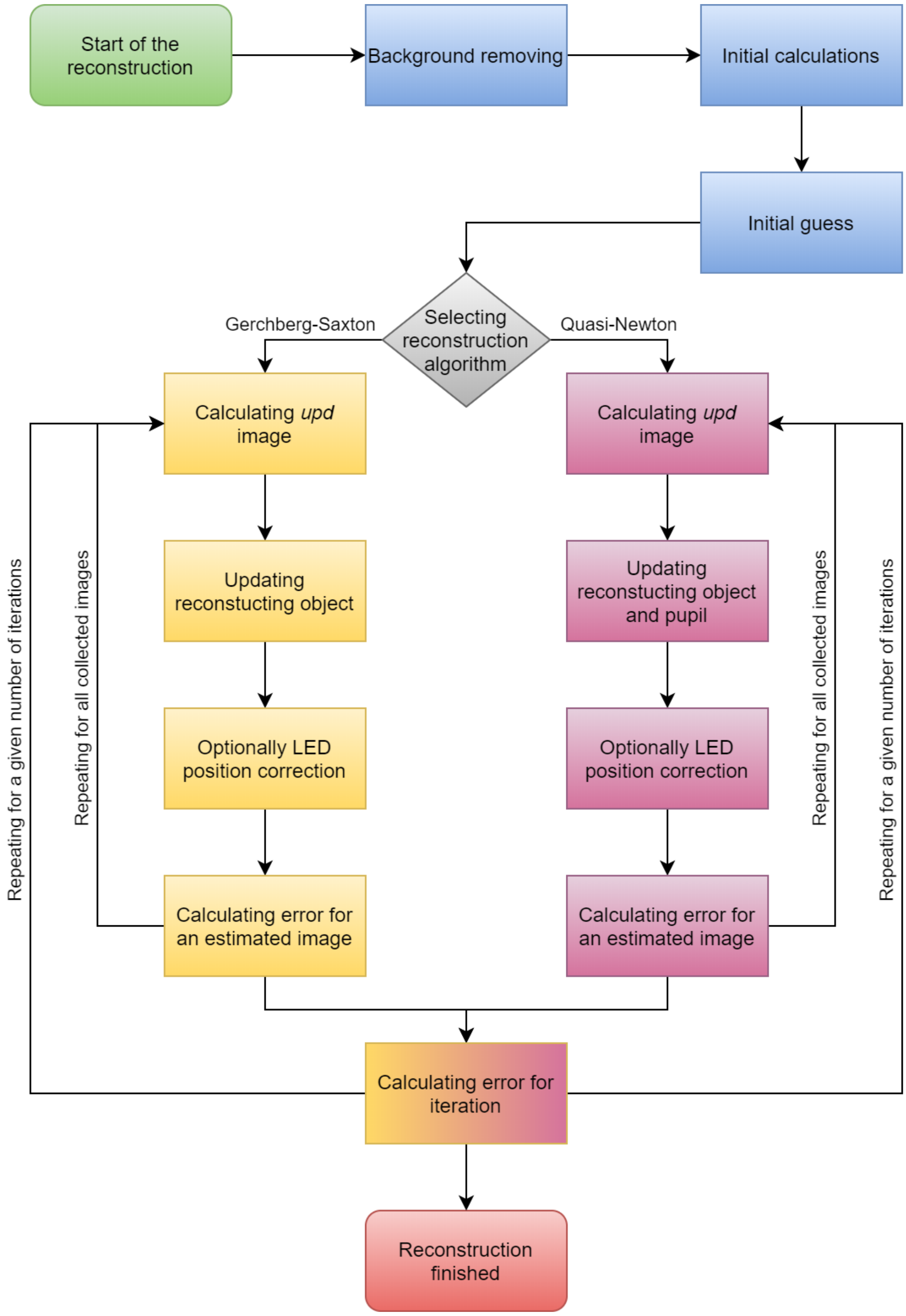

*Figure 23. Diagram illustrating process of reconstruction in FPM app.*

Reconstruction begins with background removal from input FPM images. This process is necessary due to the fact that in real FPM imaging systems, there is always some light hitting the detector which provides non-zero information even when theoretically no light scattered by sample should arrive to the detector. This additional information – background constant – needs to be removed from the input images and in *FPM* app it is performed by an algorithm specifically developed for this purpose, which is described in-depth in Chapter 5.1.

Next step is to perform initial calculations what boils down to calculating: LEDs position in the Fourier domain, system initial pupil function and the size of the reconstructed object, as described in Chapter 4.1.

After that, initial object guess is being established. As an initial guess, image collected with central LED is taken (equation (13)):

$$O = F^{-1}\left(padded\left(F\left(\sqrt{I_1}\right)\cdot P\right)\right), \tag{13}$$

where $O$ is currently reconstructed object (initial guess), $\sqrt{I_1}$ is an amplitude of collected central image, $P$ is an initial pupil function (Chapter 4.1.3). $F$ is Fourier transform and $F^{-1}$ is inverse Fourier transform. Term $F^{-1}(padded(F(\dots)))$ is due to the fact that $O$ has greater resolution (so the greater size too) than $I_n$ – when padding $I_n$ in Fourier domain, the size of the image is being increased to the $O$ size (calculated in equation (1)) but the resolution is remaining the same. In the case when the central image is not taking part in the reconstruction, image from the used images that is collected with the LED closest to the center LED is taken.

When all this is done, iterative loop can begin. This loop depends on the used reconstruction algorithm. In *FPM app* two algorithms have been implemented: Gerchberg-Saxton (G-S) [4] and Quasi-Newton (Q-N) [13], which are described below.

##### 4.4.1. Q-N algorithm

Q-N method begins with creating image ($upd_n$) that will serve to update reconstructed object with the collected image. Firstly, measured image amplitude ($mea_n$) and estimated image ($est_n$) are being calculated:

$$mea_n = \sqrt{I_n}, \tag{14}$$

$$est_n = F^{-1}\left(crop\left(F(O)\right)_{S_0 f_{xn} f_{yn}} \cdot P\right), \tag{15}$$

where $I_n$ is currently measured image, $O$ is the currently reconstructed object and $P$ is the currently reconstructed pupil. Term $crop\left(F(O)\right)_{S_0 f_{xn} f_{yn}} \cdot P$ is to take into consideration only the part of the object spectrum that corresponds to the measured image spectrum.

The $upd_n$ image is created by composing it from amplitude of $mea_n$ and phase of $est_n$ ($\varphi_n$) with additionally subtracted spectrum of $est_n$:

$$F(upd_n) = F\left(mea_n \cdot \mathrm{e}^{i\cdot\varphi_n}\right) - F(est_n). \tag{16}$$

Then $O$ and $P$ are updated:

$$O_{upd} = O_{upd} + \frac{|P|\cdot P^* \cdot F(upd_n)}{\max(|P|)\cdot(|P|^2+\alpha)}, \tag{17}$$

$$P = P + \frac{\left|O_{upd}\right|\cdot {O_{upd}}^* \cdot F(upd_n)}{\max(F(O))\cdot\left(\left|O_{upd}\right|^2+\beta\right)}\cdot P_0, \tag{18}$$

where $\alpha$ and $\beta$ are given by user regularization constants to ensure numerical stability – to avoid division by zero. After empirical verification, we have found that any value of the parameter $\beta$ greater than zero gives practically the same result. Whereas $\alpha$ parameter should be relatively small (smaller than 10), larger values may worsen reconstruction result. $P_0$ is an initial pupil function and $O_{upd}$ is a part of $O$ spectrum that is being updated in this iteration:

$$O_{upd} = crop\big(F(O)\big)_{S_0 f_{xn} f_{yn}} \cdot P_0. \tag{19}$$

After object and pupil update, one of implemented LED positions correction algorithms is being performed (if user has chosen to perform any of it). These algorithms are used to correct misaligned LED positions in the Fourier domain. Their principle of operation is described in more detail in Chapter 5.2.

Image updating process ends with error calculation ($err_n$), which is calculated as root mean square (RMS) of the difference between image estimated and measured:

$$err_n = RMS(I_n - |est_n|^2). \tag{20}$$

All these calculations (violet cells in Figure 23) are being repeated for all images which makes one iteration.

##### 4.4.2. G-S algorithm

G-S method begins with creating image ($upd_n$) that will serve to update reconstructed object with the collected image. Similarly, as in Q-N method, measured image amplitude ($mea_n$) and estimated image ($est_n$) are being calculated:

$$mea_n = \sqrt{I_n}, \tag{21}$$

$$est_n = F^{-1}\left(crop\big(F(O)\big)_{S_0 f_{xn} f_{yn}} \cdot P\right), \tag{22}$$

where $I_n$ is currently measured image, $O$ is the currently reconstructed object and $P$ is the currently reconstructed pupil. Term $crop\big(F(O)\big)_{S_0 f_{xn} f_{yn}} \cdot P$ is to take into consider only part of object spectrum that corresponds to measured image spectrum.

The $upd_n$ image is created by composing it from amplitude of $mea_n$ and phase of $est_n$ ($\varphi_n$):

$$F(upd_n) = F\big(mea_n \cdot \mathrm{e}^{i \cdot \varphi_n}\big) \tag{23}$$

Then $O$ is updated:

$$O_{upd} = F(upd_n) \cdot P_0, \tag{24}$$

where $P_0$ is an initial pupil function and $O_{upd}$ is a part of $O$ spectrum that is being updated in this iteration:

$$O_{upd} = crop\big(F(O)\big)_{S_0 f_{xn} f_{yn}} \cdot P_0. \tag{25}$$

After object update, one of implemented LED positions correction algorithms is being performed (if user has chosen to perform any of it). These algorithms are used to correct misaligned LED position in Fourier domain. Their principle of operation is described in more detail in Chapter 5.2.

Image updating process ends with error calculation ($err_n$), which is calculated as root mean square (RMS) of the difference between image estimated and measured:

$$err_n = RMS(I_n - |est_n|^2). \tag{26}$$

All these calculations (yellow cells in Figure 23) are being repeated for all images which makes one iteration.

##### 4.4.3. Computing errors

After iteration finishes, iteration error is being calculated ($err_{iter}$) as an RMS of vector containing all calculated in this iteration $err_n$ errors ($err_{1-Nimgs}$, calculated in equation 20 or 26):

$$err_{iter} = RMS\left(err_{1-Nimgs}\right). \tag{27}$$

When reconstructing synthetic data generated by the *FPM app*, also the second kind of error ($errS_{iter}$) is being calculated as an RMS between reconstructed object amplitude ($|O_{iter}|$) and known synthetic object amplitude ($obj_{amp}$):

$$errS_{iter} = RMS\left(norm_{0-1}(|O_{iter}|) - norm_{0-1}\left(obj_{amp}\right)\right), \tag{28}$$

where $norm_{0-1}(\dots)$ means image normalization from 0 to 1.

Such iterations are performed in the number indicated by the user. After each of this iteration $err_{iter}$ and $errS_{iter}$ are being calculated. In *FPM app, $err_{iter}$* is called “error compared to input images” and $errS_{iter}$ is called “error compared to known synthetic object”.

### 5. Our innovations into the FPM processing path

During our work with the FPM, we have noticed various areas where FPM can be improved. These improvements are described in this chapter.

#### 5.1. Automatic background removal

Reconstruction in the *FPM app* begins with the background removal from input FPM images. This process is necessary due to the fact that in real-data FPM imaging systems, there is always some light hitting the detector which provides non-zero information even when theoretically no light scattered by sample should arrive to the detector (Figure 24). This additional information – background constant/background noise - is required by the Q-N/G-S algorithm to be removed from the input images, otherwise this noise will be added in reconstruction process. In the original implementation of Q-N algorithm [11], [13] this was resolved by requiring a human interaction to point known region inside the FOV where there is no object. This solution was not automatic, prone to human errors and unrepeatable. Because of that, in the *FPM app* we implemented an algorithm specifically developed for this purpose, which working principle (Figure 25) is similar as original solution but does not require any user interaction.

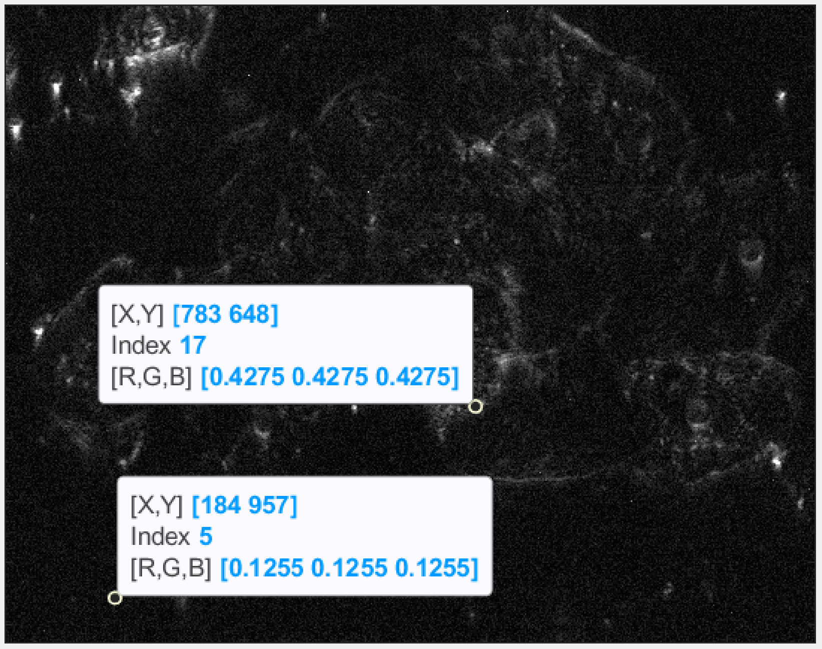

*Figure 24. Darkfield input FPM images always contain some noise that is disturbing the reconstruction. This noise is especially seen in no object regions, which theoretically should be 0 valued. Background noise may be significantly reduced by subtracting background constant value.*

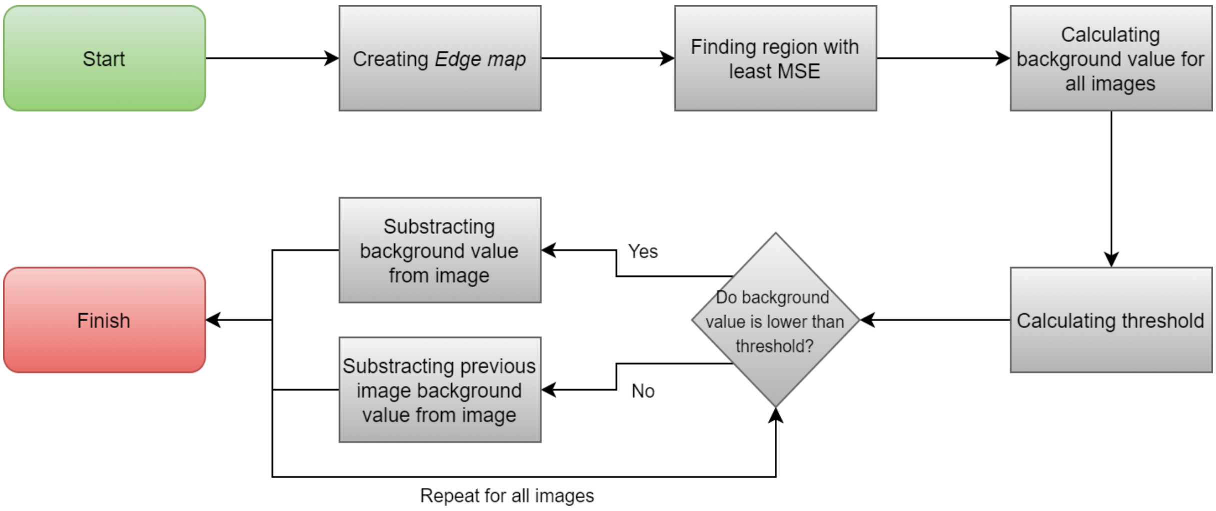

*Figure 25. Principle of operation of our background removing algorithm.*

First step of this algorithm is to create so called *Edge map* which example is shown in the Figure 26b. *Edge map* is an image created by summation of all normalized (by their respective means) input FPM images. Such created image has this advantage that it contains information about each collected image what allows to take into consideration details that are visible only in a few of all images (such details are marked with blue circle in Figure 26b). *Edge map* is also not spoiled by various aberrations that are visible only in singular images (marked with yellow circles in Figure 26a)

After that, *Edge map* is divided into 50x50 pixels regions (or 1% of the image size in case of small images) and in each of these regions, mean squared error (MSE) is calculated. Region with the lowest MSE value is selected as a global background region (GBR). Exemplary found GBR is marked with red circles in Figure 26 (circles are used to visualize fund square region which side size is equal to circle diameter).

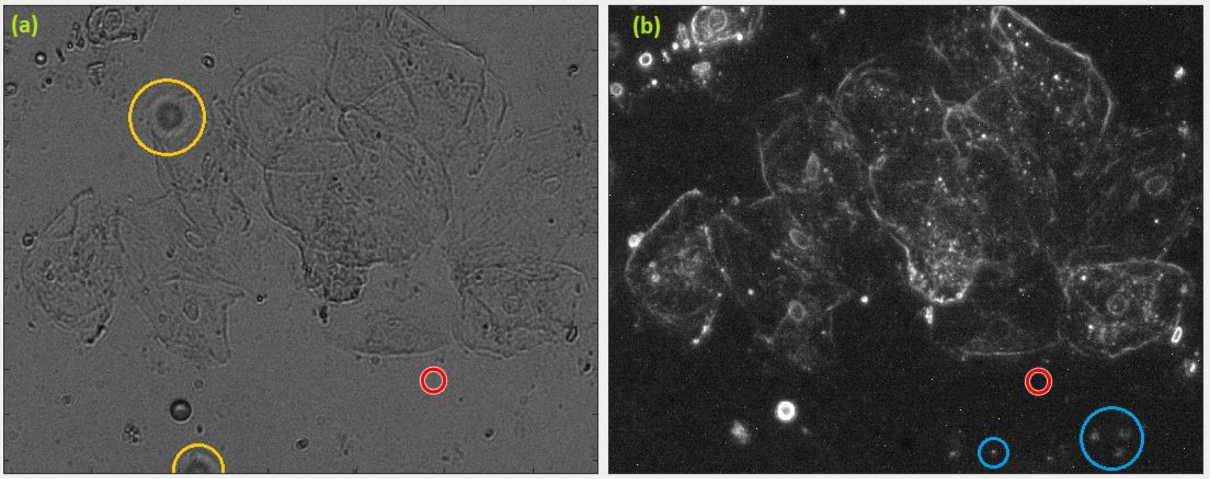

*Figure 26. (a) – input FPM image illuminated with center LED; (b) – Edge map. In Edge map there are not seen various aberrations that are visible only on one of the input images (marked with yellow circles on (a)), but there may be observed real small particles that are not visible on brightfield images (marked with blue circles on (b)). Found GBR is marked with red circle on both shown images.*

Then, for each collected image its background region value (BRV) is calculated as a mean value inside GBR. BRV will be approximately equal to the background constant only in darkfield images. In brightfield images light that is not scattered by the sample also hits the detector, so obtained BRVs will be in fact equal to the background constant plus intensity of non-scattered light. To overcome this problem, brightfield images need to be distinguished from darkfield ones. It is done by determining threshold parameter above which obtained background value belongs to brightfield image and below to darkfield image. Because BRVs for brightfield images are always much larger than for darkfield images, threshold parameter is simply calculated as the average value of the maximal and minimal BRVs.

Last step of background removing algorithm is to subtract the BRVs from input FPM images. When input image BRV is higher than threshold parameter, then previous image BRV is subtracted.

Output of this algorithm are input FPM data with removed background constant and partially removed background noise. Reconstruction of such images results in obtaining much less noisy images than without background removal (Figure 27). Moreover, in the *FPM app*, noise that still remains in reconstruction may be further removed with the use of the additional denoising algorithm (see Chapter 5.4).

Performance of our algorithm is hard to compare with one implemented in the original Q-N method. Both are basing on calculating BRVs in GBR and then subtracting them from input images. The difference is in finding GBR – in our solution it is done automatically whereas in original solution, user needs to point this region, which makes such comparison user-dependent. However, we have tested our algorithm on 20 datasets (10 real datasets and 10 synthetic datasets), and in every time found GBR was a region that for user looked as no object region. Moreover, during our work with *FPM app* we have not encountered any reconstruction error caused by background removing algorithm what should ensure that it will perform correctly in virtually every case.

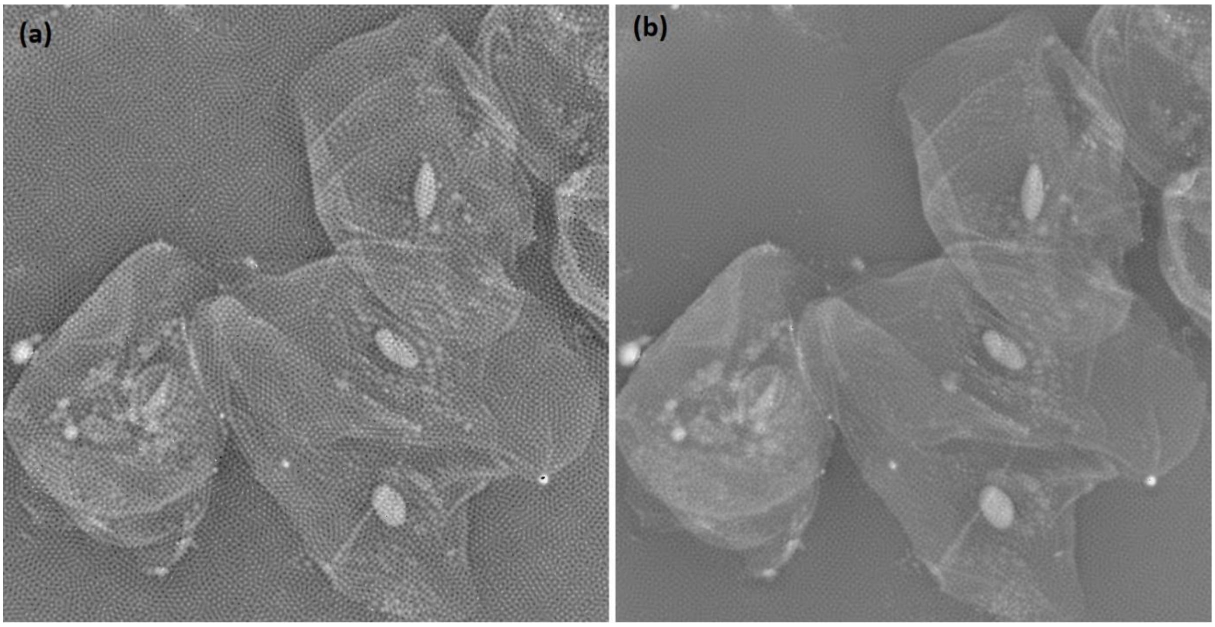

*Figure 27. Cheek cells reconstruction results without background removing (a) and with the use of our background removing algorithm (b).*

#### 5.2. Misalignment error correction

One of the most problematic FPM errors is the misalignment error [5]. When constructing a Fourier ptychographic microscope, LED array (or other light source) position in this microscope has finite accuracy. LED array can be slightly shifted or rotated in any direction, distance between singular LEDs may also not be accurate. All this can cause that real images position in the Fourier domain may differ from positions calculated from system parameters.

Fortunately, these positions can be corrected during reconstruction algorithm. In conventional ptychography, simulated annealing (SA) algorithm [14] was proposed to correct position errors in probe function and this algorithm is also treated as a base in the FPM position correction methods [15] [16]. In FPM, SA algorithm is used to find minimum of a function $\varphi(f_{xn}, f_{yn})$ after every reconstructing object update:

$$\varphi(f_{xn}, f_{yn}) = \sum (est_n(f_{xn}, f_{yn}) - mea_n)^2, \tag{29}$$

where $est_n(f_{xn}, f_{yn})$ is the estimated image with the center position in object spectrum in point $(f_{xn}, f_{yn})$ and $mea_n$ is a measured image.

One of the biggest disadvantages of the SA based method is that it extends the time of the reconstruction more than five-fold. So it is not surprising that other methods were also proposed, e.g., angle self-calibration method [17], method with the use of deep neural networks [18] or recently proposed numerical Multi-Look approach [19].

Because of extremely long time of SA based methods and the fact that other methods are relatively new or hard to implement in the *FPM app*, we proposed two original algorithms (called *Accurate algorithm* and *Fast algorithm*) that should deal with misalignment error in a shorter time than the SA methods. However, in future updates, other methods also may be implemented.

##### 5.2.1. Accurate algorithm

In this method the minimum of the function from equation (29) is sought. This time, no advanced algorithm is being used but all $\varphi(f_{xn}, f_{yn})$ function values inside certain range are calculated (Figure 28) and the position of the smallest calculated value is set as a new image position in Fourier spectrum of the object. Range within the function is calculated and is set as square with a side of 2/3 of the distance between centers of adjacent images in Fourier domain.

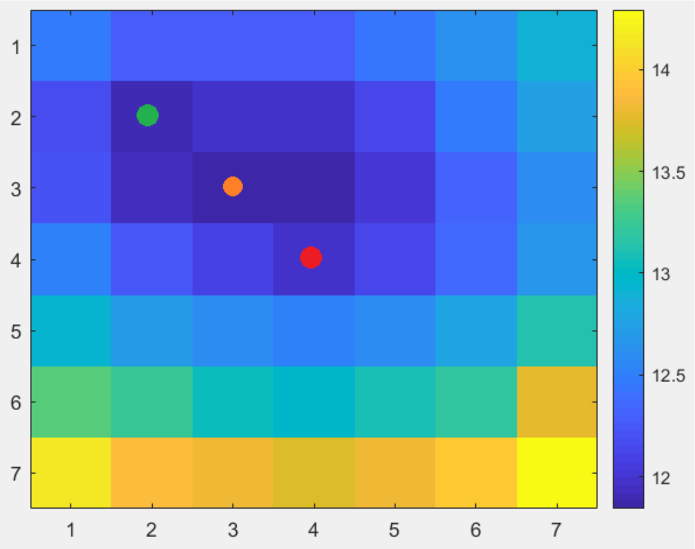

*Figure 28. $F(f_{xn}, f_{yn})$ value for each $(f_{xn}, f_{yn})$ position inside calculated range. $(f_{xn}, f_{yn})$ position calculated from system parameters is marked with red dot. Orange dot marks the position with minimum $F(f_{xn}, f_{yn})$ value. Green dot is the real position of the image in Fourier space.*

##### 5.2.2. Fast algorithm

Principle of operation of fast algorithm is similar to the accurate algorithm but this time, apart from range inside which $\varphi(f_{xn}, f_{yn})$ is being calculated, there is also smaller subrange proposed which currently is rigidly set as square with side of 5 pixels length. Firstly, $\varphi(f_{xn}, f_{yn})$ is being calculated only in that subrange. Then minimum of the function is being found and if it is not in the center of the subrange then $\varphi(f_{xn}, f_{yn})$ is being calculated in the subrange that has center in newly found minimum (Figure 29). This process repeats up to the moment when minimum was found in the center of the subrange.

With this algorithm, $\varphi(f_{xn}, f_{yn})$ is not being calculated in all range, but still the minimum of the function should be found correctly.

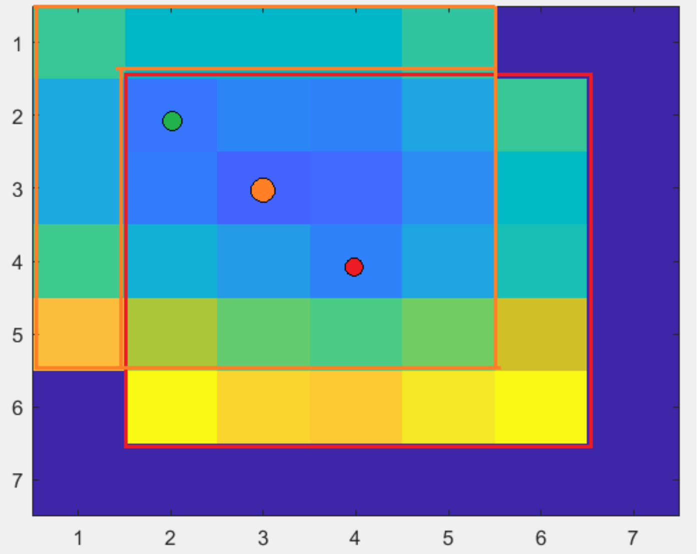

*Figure 29. $F(f_{xn}, f_{yn})$ value for each $(f_{xn}, f_{yn})$ position inside calculated subranges. Firstly, calculated subrange is marked with red square, nest subrange is marked with an orange square. $(f_{xn}, f_{yn})$ position calculated from system parameters is marked with red dot. Orange dot marks the position with minimum $F(f_{xn}, f_{yn})$ value. Green dot is the real position of the image in Fourier space.*

##### 5.2.3. Comparison

In the *FPM app* both developed original algorithms and the SA algorithm were implemented. To test them, synthetic data were created with a random LED position misalignment. Then this data was firstly reconstructed with known corrected misalignment positions (ground truth), and then with all implemented LED correction algorithms and without LED correction. Results are shown in the Figure 30. As can be seen all three methods properly corrected misaligned positions. Root mean squared error (RMSE) was calculated between ground truth and other reconstructions and it was similar for all implemented algorithms (it was slightly smaller for the SA algorithm). However, data reconstruction time of the SA algorithm was almost 2x time longer than second slowest *Accurate algorithm*, what is an important disadvantage of SA algorithm. Also, corrected misaligned positions retrieved by *Fast* and *Accurate* algorithms often overlap with known true positions, which is another clue that this algorithms perform correctly.

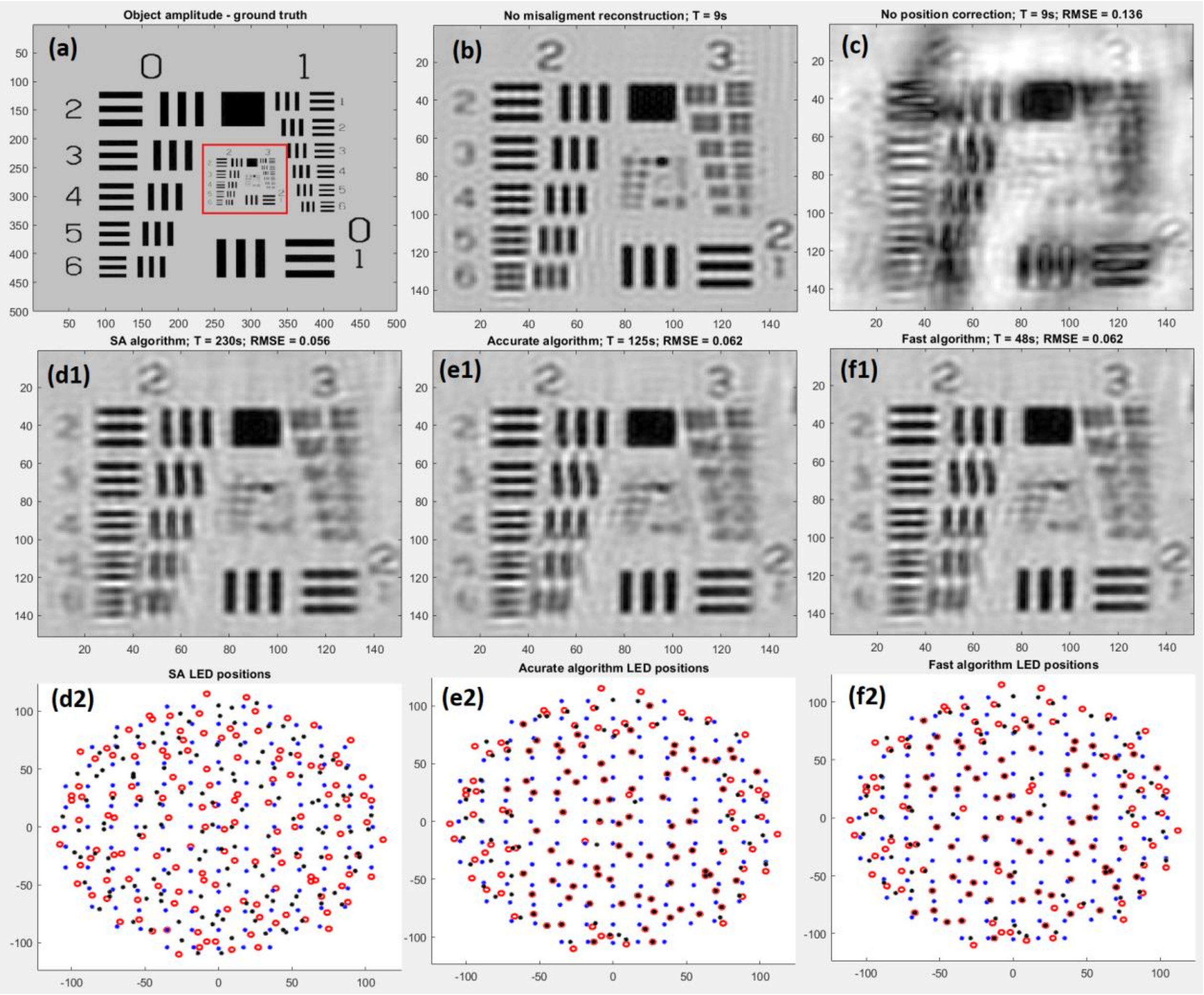

*Figure 30. (a) – synthetic object amplitude; (b) – object reconstruction without introduced misalignment error; (c) – object reconstruction with misalignment error; (d1–f1) – object reconstruction with misalignment LED correction and SA/Accurate/Fast algorithm correction, respectively; (d2–f2) – original LEDs positions (blue dots), corrected LEDs positions (black dots) and known true LEDs positions (red circles) for SA/Accurate/Fast algorithm, respectively.*

#### 5.3. Reconstruction time optimization

One of the biggest problems of the FPM is a long reconstruction time. During the reconstruction tens of thousands of Fourier transforms need to be calculated, usually on large images. Because of that, reconstruction of large FOV may take even several hours. The *FPM app* shares this inherent limitation thus the attempt to optimize reconstruction time was undertaken. Process of optimization were done by giving to the user the possibility to perform some of the calculations on the graphics processing unit (GPU). Results were quite promising, especially for a larger input data size and in comparison to performing all operations on the central processing unit (CPU), reconstruction time was significantly shorter.

Comparison of the reconstruction time in a function of the input data size are shown in the Figure 31. Reconstructions were done on a computer with a: processor 2,3 GHz Intel Core i5-8300H, 8 GB RAM and graphics card Nvidia GeForce GTX 1050 (4 GB video memory).

Using GPU in *FPM app* is not always favorable. For smaller data size, reconstruction performed on the CPU is usually faster. Moreover, usage of the GPU is limited by its memory. To perform reconstruction it must store 3D matrix containing all images in double data type, which for employed GPU (4 GB video memory) did not allow to reconstruct dataset larger than 293 900x900 pixels images. This limit can be circumvented by performing several reconstructions on a smaller ROIs and then stitching the results – with this method, reconstruction of 293 2700x2700 pixels images would take around 25 minutes and will result in obtaining 182 Mpix image. In comparison, analogous CPU reconstruction would take over 2 hours. Another requirement for using the GPU is to have corresponding MATLAB toolboxes (when using *FPM app* **MATLAB version**) and Nvidia graphics card with actualized CUDA driver.

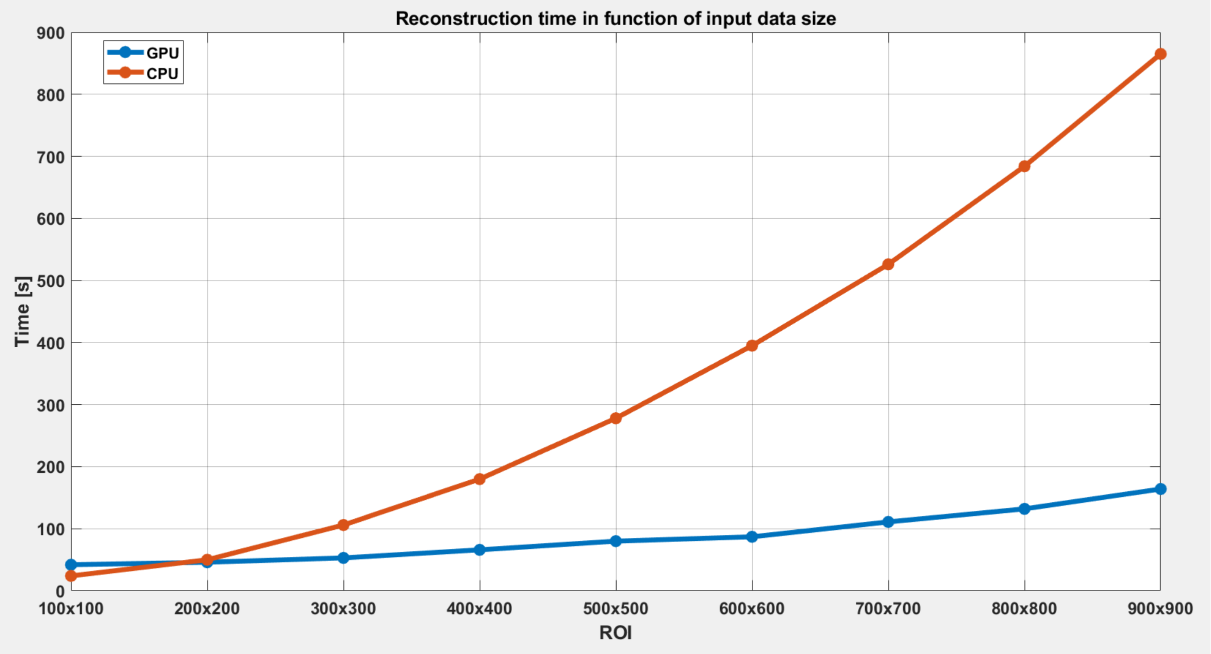

*Figure 31. Comparison of reconstruction time in function of reconstructed input data size. Reconstruction consisted of 8 iterations of 293 ROI size images.*

#### 5.4. Final output denoising

FPM provides high-resolution reconstruction of the object. This reconstruction depends on many parameters which may perturb the reconstruction (e.g., LEDs may have different illumination intensity). Also collected input data may contain various noise instances and reconstruction can cause numerical errors. All this cause deterioration of reconstruction results.

To further improve the results quality, we are proposing one simple improvement – digital denoising of the output. As a denoising method, we have chosen the state-of-the-art block-matching and 3D filtering (BM3D) algorithm [12] due to its simplicity, reliability and good overall performance.

To test the influence of output denoising, reconstructions of real data (cheek cell) and synthetic data were performed. Random 20% noise was added to the synthetic data (to every input image pixel there was added random value from 0 to 20% of maximal image value).

Results of output denoising are shown in the Figure 32. Visual comparison show noticeable improvement of reconstruction quality. Denoised reconstructed objects are much less noisy and did not lose any details. In case of synthetic data we have also calculated RMS error between ground truth and reconstructed object (Table 1), which also confirms positive influence of the BM3D method onto the reconstruction.

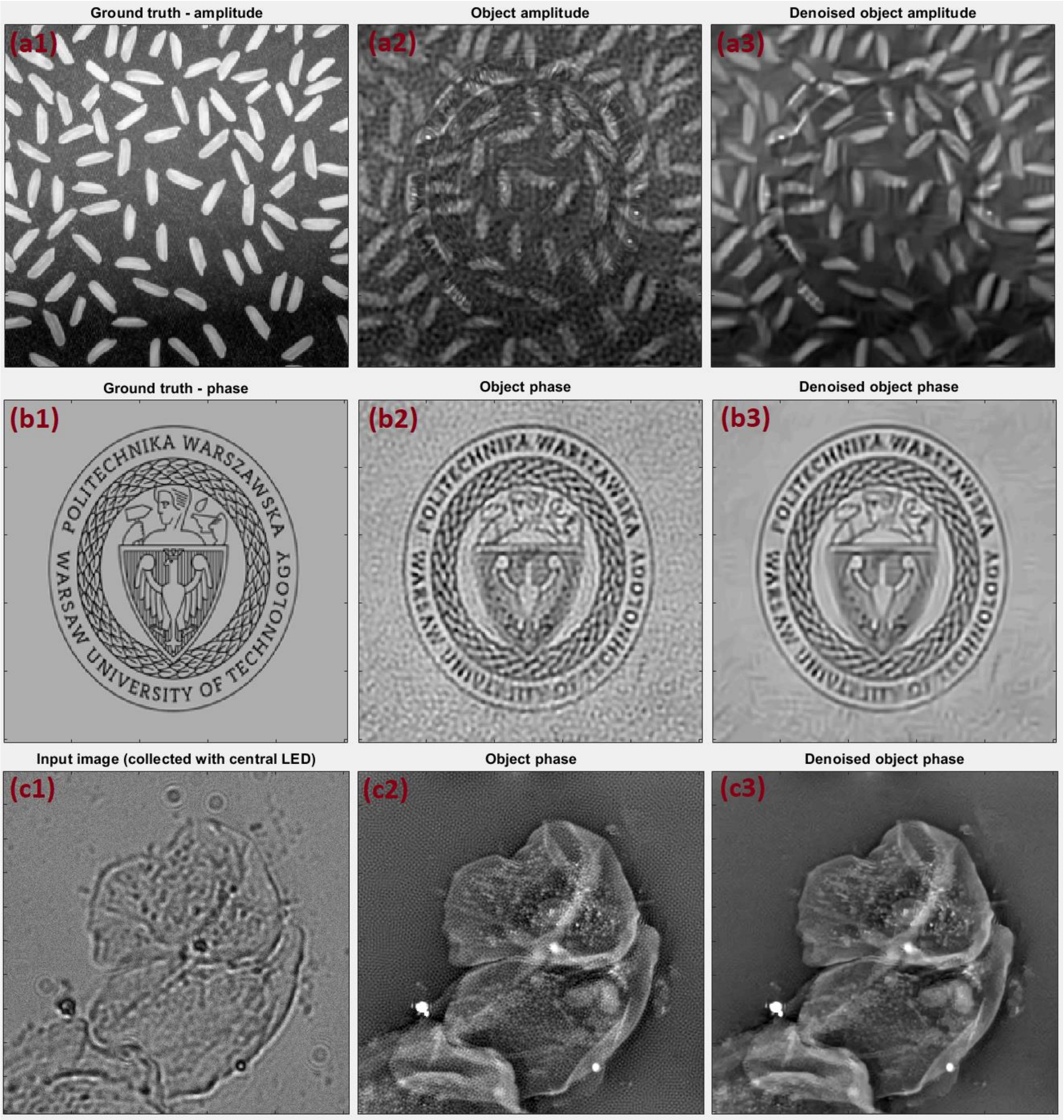

*Figure 32. (a),(b) – synthetic amplitude-phase object reconstruction result; (c) – real data cheek cell reconstruction result*

*Table 1. RMSE calculation of the data presented in Figure 32a and Figure 32b.*

| Image | Reconstructed synthetic amplitude | Reconstructed synthetic phase | Denoised reconstructed synthetic amplitude | Denoised reconstructed synthetic phase |
|---|---|---|---|---|
| RMS error compared to known synthetic object | 0.182 | 0.107 | 0.119 | 0.097 |

### 6. Modifying *FPM app* (MATLAB version)

You are free to modify *FPM app* for your own purposes. However, *FPM app* consists of dozens of scripts, each one consisting of hundreds of codes lines, which may not be the easiest thing to modify. To make it easier, in this chapter some tips that may help you modifying it are highlighted.

All *FPM app* key functions are taking inputs from MATLAB workspace and are returning outputs to it. The easiest way to modify our app is to modify workspace variables manually before a function is taking them as inputs. Most important *FPM app* workspace variables are descripted in Table 2 below.

*Table 2. Most important* FPM app *workspace variables*

| Variable | What does it store | When does it appear | When is it changed | When is it saved * |
|---|---|---|---|---|
| amplitude_synth | Synthetic object amplitude | After opening *Generate images* window | After changing it with **Input amplitude image** or **Constant amplitude** buttons | Manually (**file -> save project**) |
| err | Error compared to input data | After reconstruction finishes | After each next reconstruction | Manually (**Save results** button) |
| erro | Error compared to known synthetic object | After reconstruction finishes | After each next synthetic data reconstruction | Manually (**Save results** button) |
| idx_X | Positions of images centres in Fourier spectrum in x direction | After reconstruction finishes | After each next reconstruction | Manually (**Save results** button) |
| idx_Y | Positions of images centres in Fourier spectrum in y direction | After reconstruction finishes | After each next reconstruction | Manually (**Save results** button) |
| imageColOrder | Images collecting order | After opening *FPM app* | After is changed with **Set collecting order** button in *image collecting order* window | Manually (**file -> save project**) Automatically (after reconstruction finishes) |
| imageList | List of loaded into the app images names | After loading data into the app (**Load data** button) | After loading another data into the app (**Load data** button) | - |
| ImagesIn | Input FPM images | After loading data into the app (**Load data** button) or generating synthetic data in *Generate images* window | After loading another data into the app/generating another data | - |
| LEDs | LED array layout | After opening *FPM app* | After is changed with **Set this layout** button in *LED placement* window | Manually (**file -> save project**) Automatically (after reconstruction finishes) |

| | | | | |
|---|---|---|---|---|
| LEDsUsed | Images (LEDs used to collect them) used in reconstruction | After opening *FPM app* | After is changed in *Used LEDs* window | Manually (**file -> save project**) Automatically (after reconstruction finishes) |
| object | Reconstructed complex object | After reconstruction finishes | After each next reconstruction | Manually (**Save results** button) |
| object_denoised | Denoised reconstructed complex object | After object denoising (**Denoise results** button) | After each next object denoising | Manually (**Save results** button) |
| options | Reconstruction options | After opening *FPM app* | After each reconstruction | Manually (**file -> save project**) Automatically (after reconstruction finishes) |
| phase | Reconstructed object phase | After reconstruction finishes | After each next reconstruction | Manually (**Save results** button) |
| phase_denoised | Denoised reconstructed object phase | After object denoising (**Denoise results** button) | After each next object denoising | Manually (**Save results** button) |
| phase_synth | Synthetic object amplitude | After opening *Generate images* window | After changing it with **Input amplitude image** or **Constant amplitude** buttons | Manually (**file -> save project**) |
| pupil | System pupil function | After reconstruction finishes | After each next reconstruction | Manually (**Save results** button) |
| ROI | Region of interest | After loading data into the app (**Load data** button) or generating synthetic data in *Generate images* window | After loading another data into the app/generating another data or by changing ROI in **ROI selection** panel | - |
| systemSetup | Imaging system parameters | After opening *FPM app* | After each change in **System setup** panels | Manually (**file -> save project**) Automatically (after reconstruction finishes) |

** Some of the variables may be saved manually through the FPM app buttons, some of them are also saved automatically to initialization.mat or initialization2.mat files which are loaded after every FPM app launching.*

*FPM app* scripts also may be modified. To modify existing codes you need to follow the comments inside them to see where should you place your modifications.

One of the modifications that you probably may want to implement is unwrapping of reconstructed phase. In this case we recommend to overwrite phase variable with unwrapped phase (e.g., phase = unwrap_algorithm(phase)). This will cause that BM3D denoising, show images and save images functions will denoise/show/save this unwrapped phase.

### 7. Exemplary process of collecting and reconstructing FPM data (demo)

#### 7.1. Creating FPM system

Simplest FPM imaging system may be created by modifying standard brightfield microscope with a LED array placed instead of microscope illuminator and such system is described in this example.

First thing to consider is which LED array would be suitable for used microscope and where it should be placed. Answer on these questions may be found with the use of *FPM app* like in the following procedure:

1. Open *FPM app.*
2. Push **Generate synthetic data** button.
3. Type your microscope parameters into **System setup** panel.
4. Type spacing between adjacent LEDs in your LED array into **System setup** panel.
5. Set your LED array shape in **LED layout** panel.
6. Manipulate the **LED height** value (distance between LED array and measured sample) in **System setup** panel and observe **Synthetic NA** and **Overlap in Fourier domain between neighbouring LEDs** fields along with *Positions of the image centers* window:
   6.1. Synthetic NA should be as large as possible.
   6.2. Overlap in Fourier domain should be larger than 40%, optimally in range 40-60%.
   6.3. Images positions (blue dots) in *Positions of the image centers* window should be rather evenly sampled.
7. **LED height** value is the distance from measured sample where you should place your LED array.

Exemplary simulated imaging system and **LED height** calculated with above procedure is shown in Figure 33.

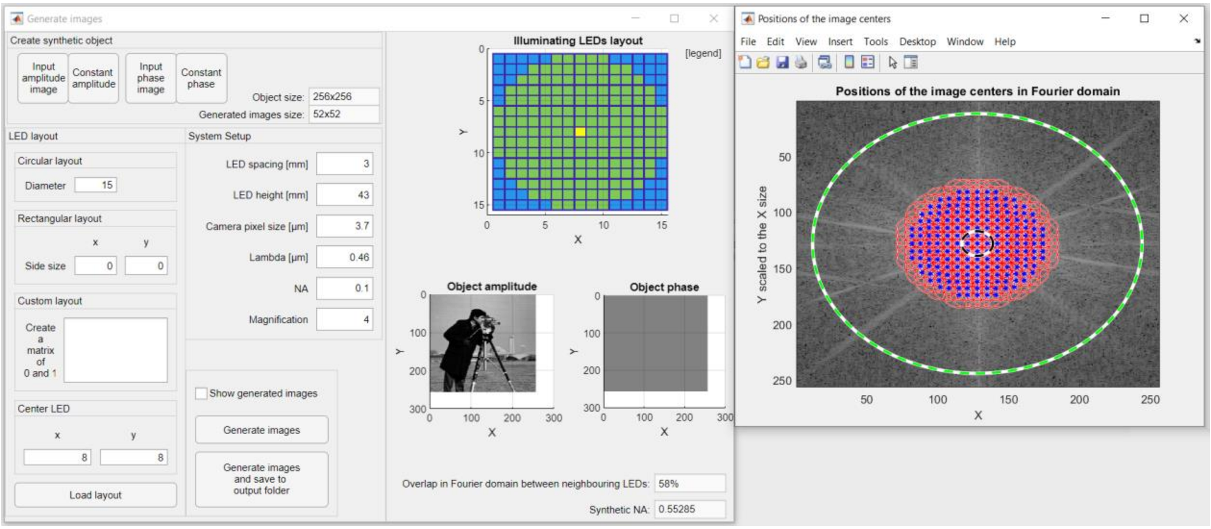

*Figure 33. Exemplary synthetic FPM system.*

When this is known, LED array may be placed in the microscope. When placing the LED array it is important that:

- There should be one center LED in LED array – LED that illuminates sample at 0°. The closer to 0° is this angle, the better should be the reconstruction result. This can be adjusted by observing alternately images illuminated by one LED a few spots to the right and one a few spots to the left from center LED. These images should be as similar to each other as possible (this should be the easiest to determine on partially brightfield partially darkfield images – images observed with illumination angle similar to objective NA). Analogous operation should be performed with LEDs at the bottom and top to the central LED.
- LED array should not be shifted or rotated in any direction. This will cause various errors during the reconstruction, which may be partially corrected but with the unnecessary use of rather time consuming algorithms.

#### 7.2. Collecting FPM images

FPM input data are microscopic images, each one collected at different illumination angle (with a different LED in LED array). From a practical point of view, the easiest way to collect FPM images is to integrate LED array with the microscope camera. Then procedure of collecting images may be as in below pseudo-code:

```
t = 0;
for(x = 1; x <= number_of_LEDs_in_x_direction; x++)
{
        for(y = 1; y <= number_of_LEDs_in_y_direction; y++)
        {
                t++;
                Turn_on_LED(x,y);
                Collect_image;
                Save_image_as(t.tif);
                Turn_off_LED(x,y);
        }
}
```

When collecting images it is important that:

- Each image should be collected with the same gain and exposure time. It is worth to make sure that no image will be overexposed or underexposed.
- LEDs should be turned on in orderly sequence (e.g., line by line), otherwise *FPM app* would not manage to appropriately associate collected images to correct LED.
- Image naming should also be corresponding to order in which they were collected.

Exemplary collected images of the system like in Figure 33 are shown in Figure 34. This dataset may be downloaded at https://bit.ly/2MxNpGb (*Navicula elliptica exemplary dataset.zip* file).

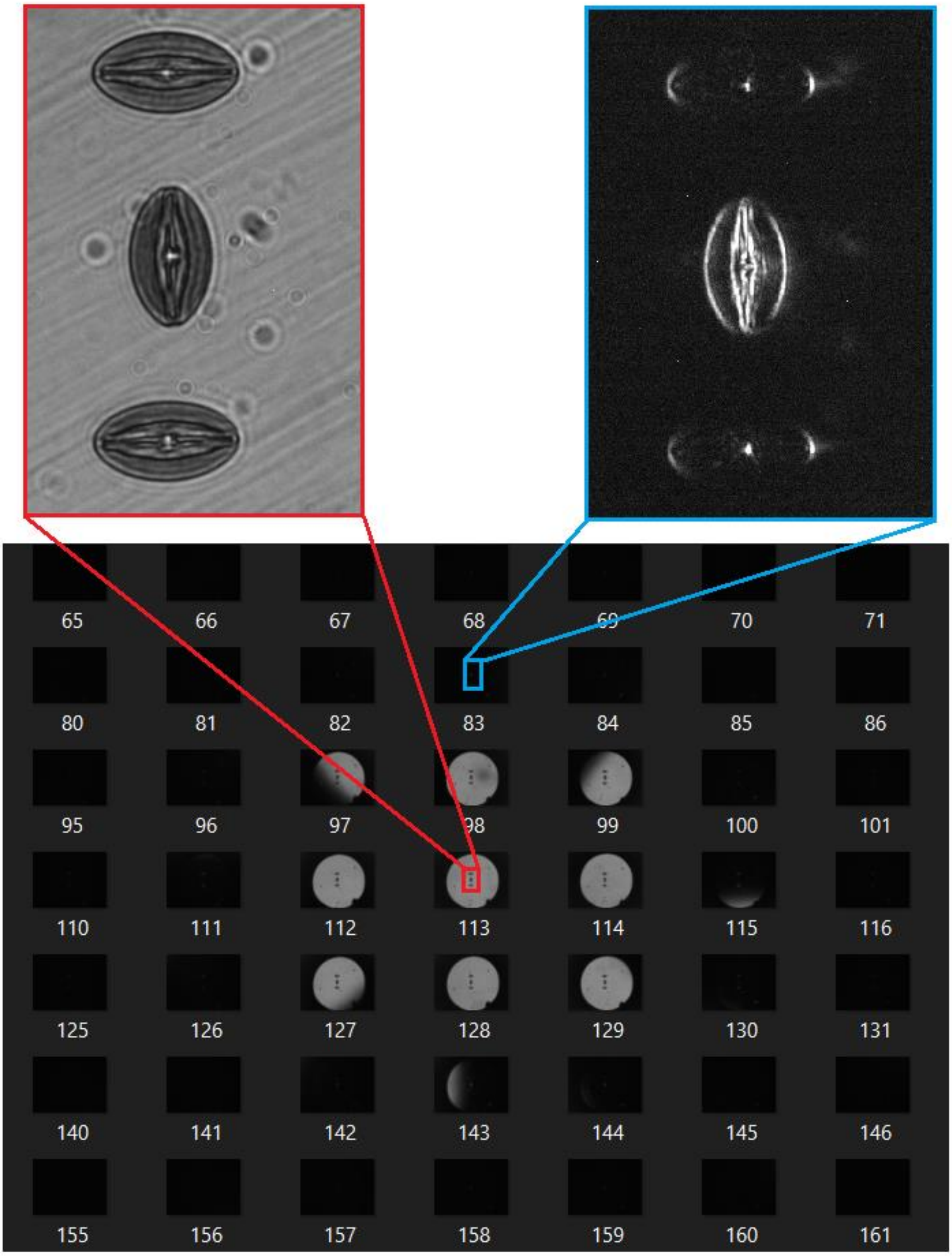

*Figure 34. Some of the FPM input images collected by the exemplary system.*

#### 7.3. Reconstruction of collected dataset

When data is collected, reconstruction with *FPM app* may begin. Exemplary procedure of reconstruction is shown below (in parentheses and images there are parameters for reconstruction of dataset collected in Chapter 7.2):

1. Open *FPM app.*
2. Push **Load data** button and select folder with stored input data.
3. Wait until the data will be loaded into the app.
4. Open *LED placement* window with the use of **Change LED placement** button.
5. In *LED placement* window set the size of LED array that was used to collect images (Rectangular layout 15x15) and push **Set this layout** button to load this layout into the app.

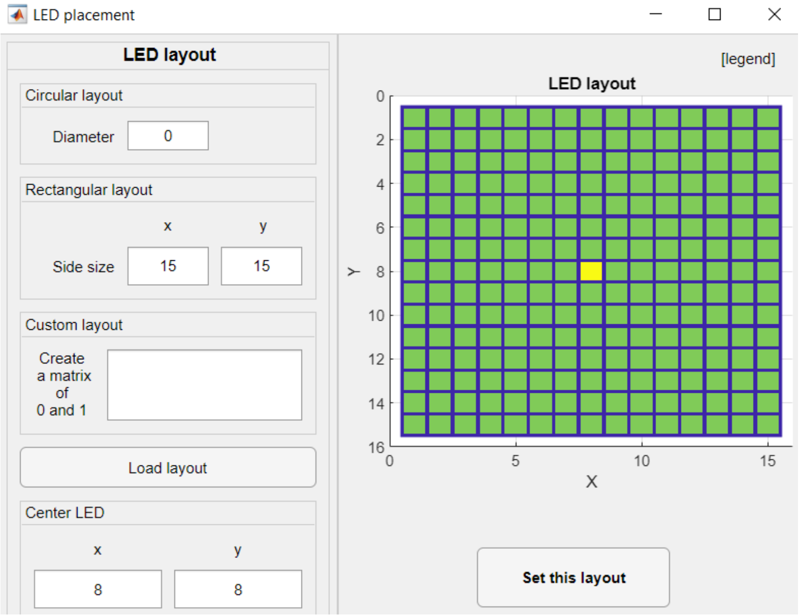

6. Open *Image collecting order* window with the use of **Change collecting order** button.
7. In *Image collecting order* window set the order in which images were collected in relation to position of the microscope camera (row by row order, starting from top right LED) and push **Set collecting order** button to load this order into the app.

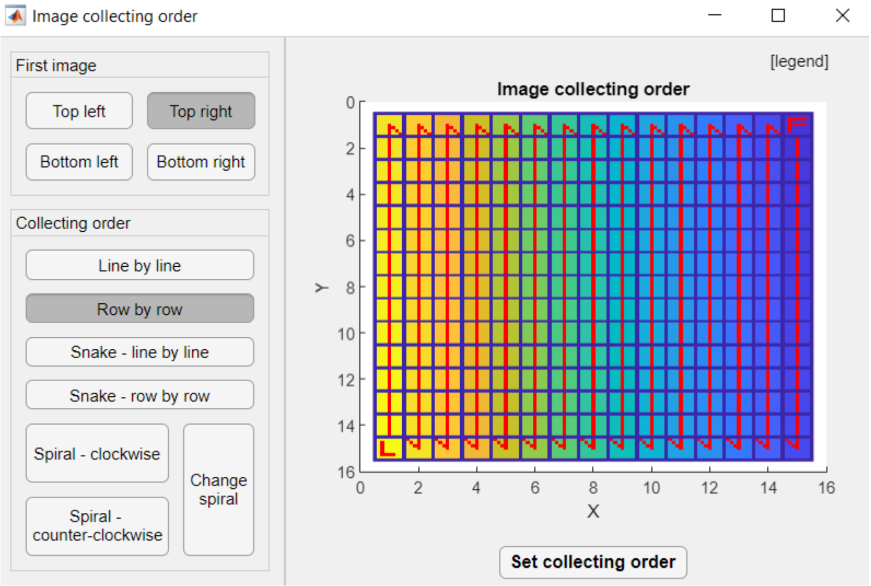

8. Push **Select on image** button to open *ROI selection* window.
9. In *ROI selection* window mark the region of FOV that you want to reconstruct by pressing and holding mouse left button on the image and then moving the mouse to mark the rectangle. Double click inside this rectangle to confirm selected region. If you want to reconstruct full FOV it is also good to firstly try on smaller ROI if the reconstruction will perform correctly.

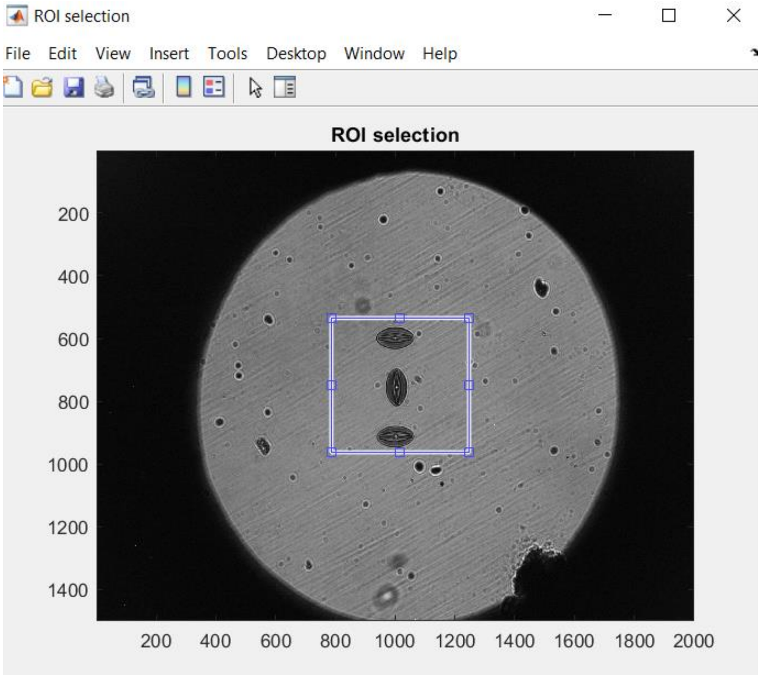

10. In **System setup** panel type your imaging system parameters.

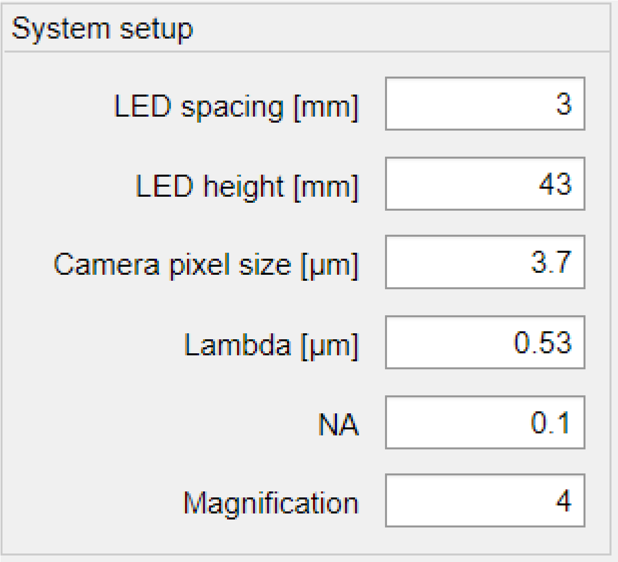

11. Push **Change used LEDs** button to open *Used LEDs* window.
12. In *Used LEDs* window select images that want to use in reconstruction (R min = 0; R max = 7.5). *FPM app* usually performs better with circular LED layouts – in every direction in XY plane there is the same range of spatial frequencies.

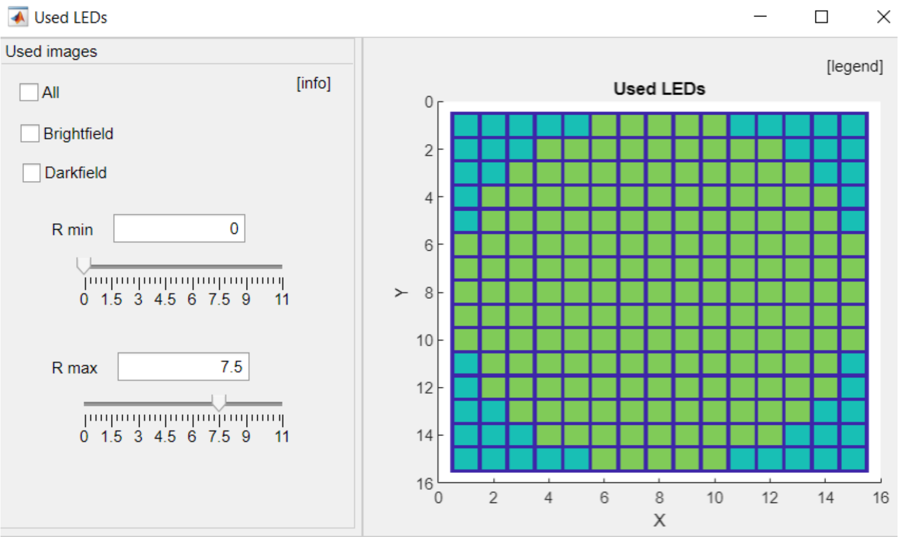

13. Set reconstruction options that want to use in reconstruction. For first reconstruction of new dataset we recommend Q-N algorithm, no LED position correction, number of iterations = 8, alpha = 1 and Beta = 1000.

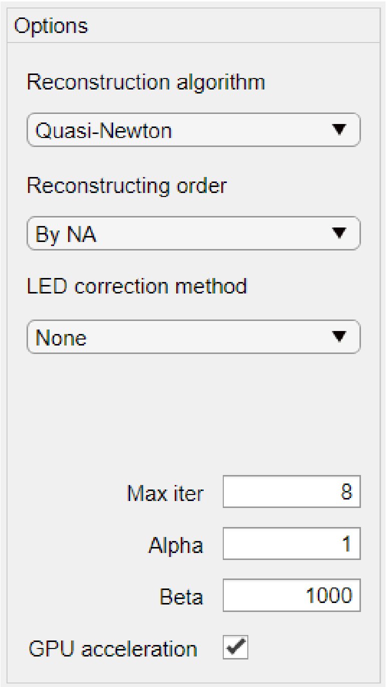

14. Press **Run algorithm** button to begin the reconstruction.
15. When reconstruction finishes, *Show results wizard* window will pop-up, where you can display obtained results. Press **Show object amplitude** and **Show object phase** buttons to display object phase and amplitude. With the use of sliders adjust displaying values range. Press **Export images as .TIFF** button to save results as .TIFF images.

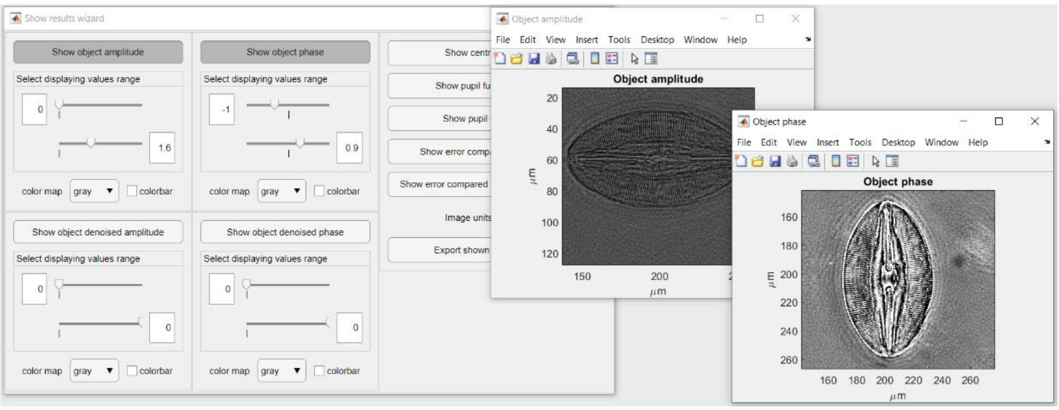

Results of reconstruction data with above procedure is shown in Figure 35. This reconstruction lasted 70 s (all processing times mentioned in this chapter are given for 2.3 GHz, 8 GB RAM and 4 GB GPU video memory laptop).

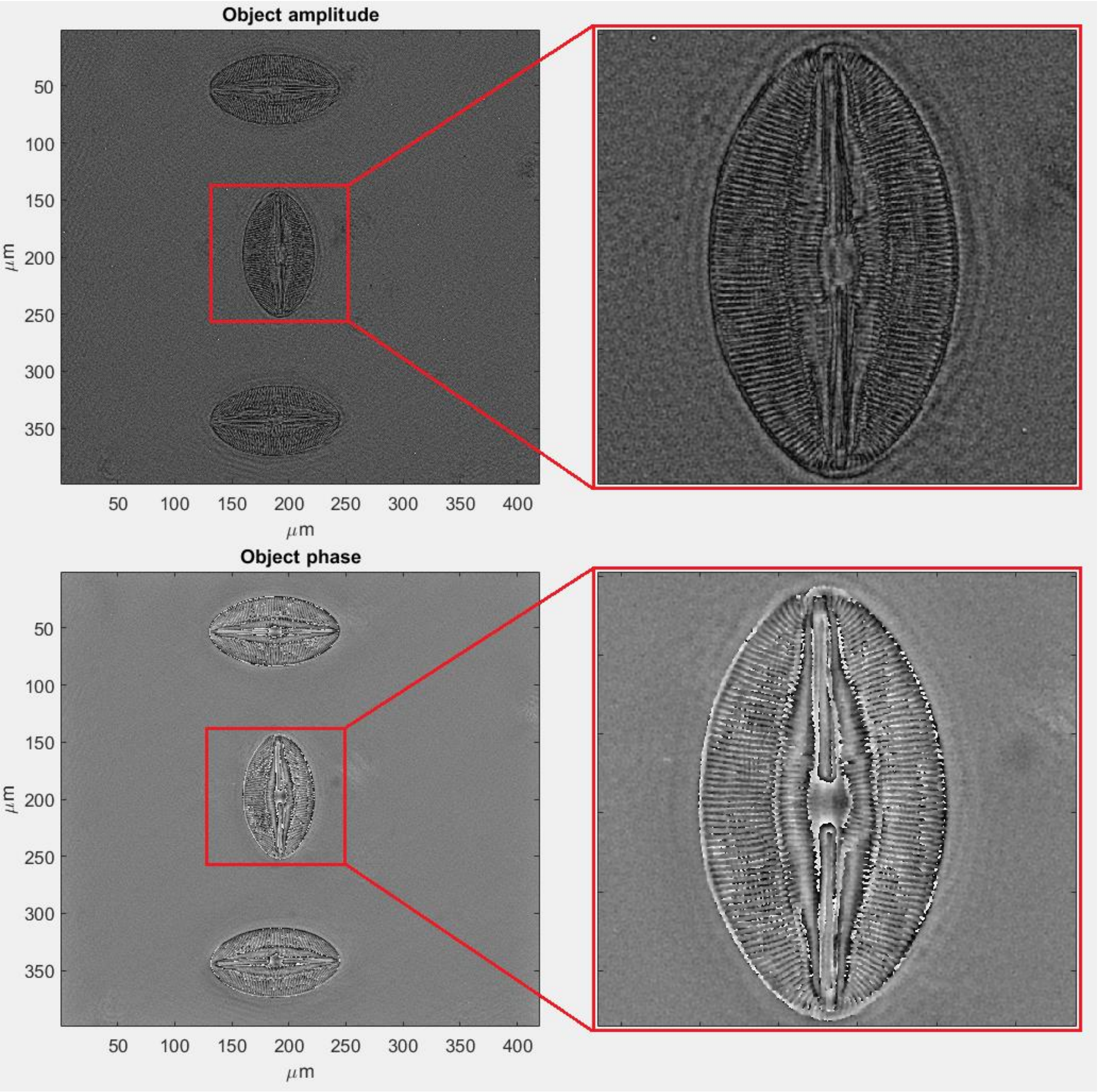

*Figure 35. Reconstructed amplitude and phase obtained with FPM app reconstruction procedure described in this chapter. Amplitude displaying range was narrowed from 0-5.3 to 0-2.3 to increase displaying contrast (all Navicula details fit in this narrowed range).*

**Additional Operations:**

- Output denoising

To minimize the noise, additional output denoising operations may be performed:

1. When reconstruction finishes, set sigma value in **Denoising** panel. The larger is sigma parameter (in range 0-255), the stronger is denoising. We recommend sigma < 50.
2. Push **Denoise results** button to begin denoising. After it finishes, the *Show results wizard* window will pop-up along with displayed denoised object amplitude and phase (in this example, sigma = 40).

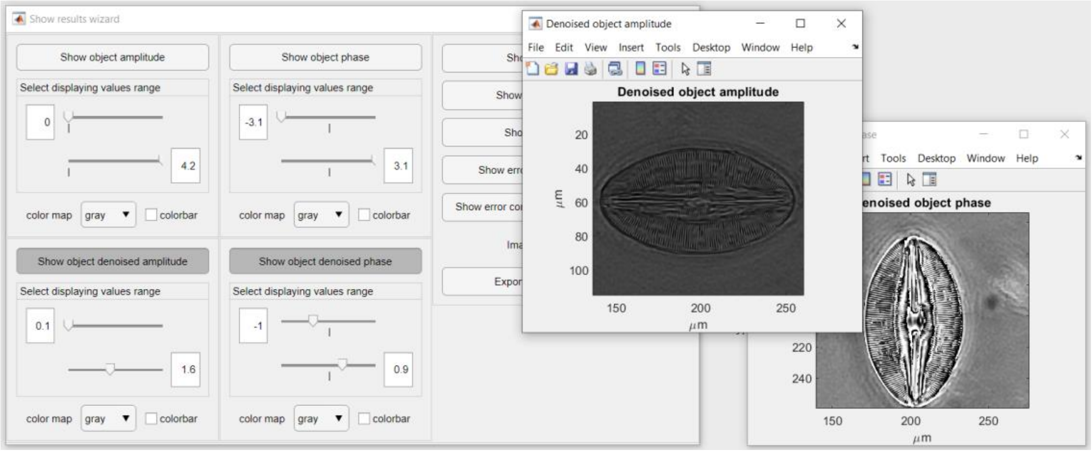

Denoised reconstruction results obtained with above procedure are shown in Figure 36. This denoising lasted 11 s.

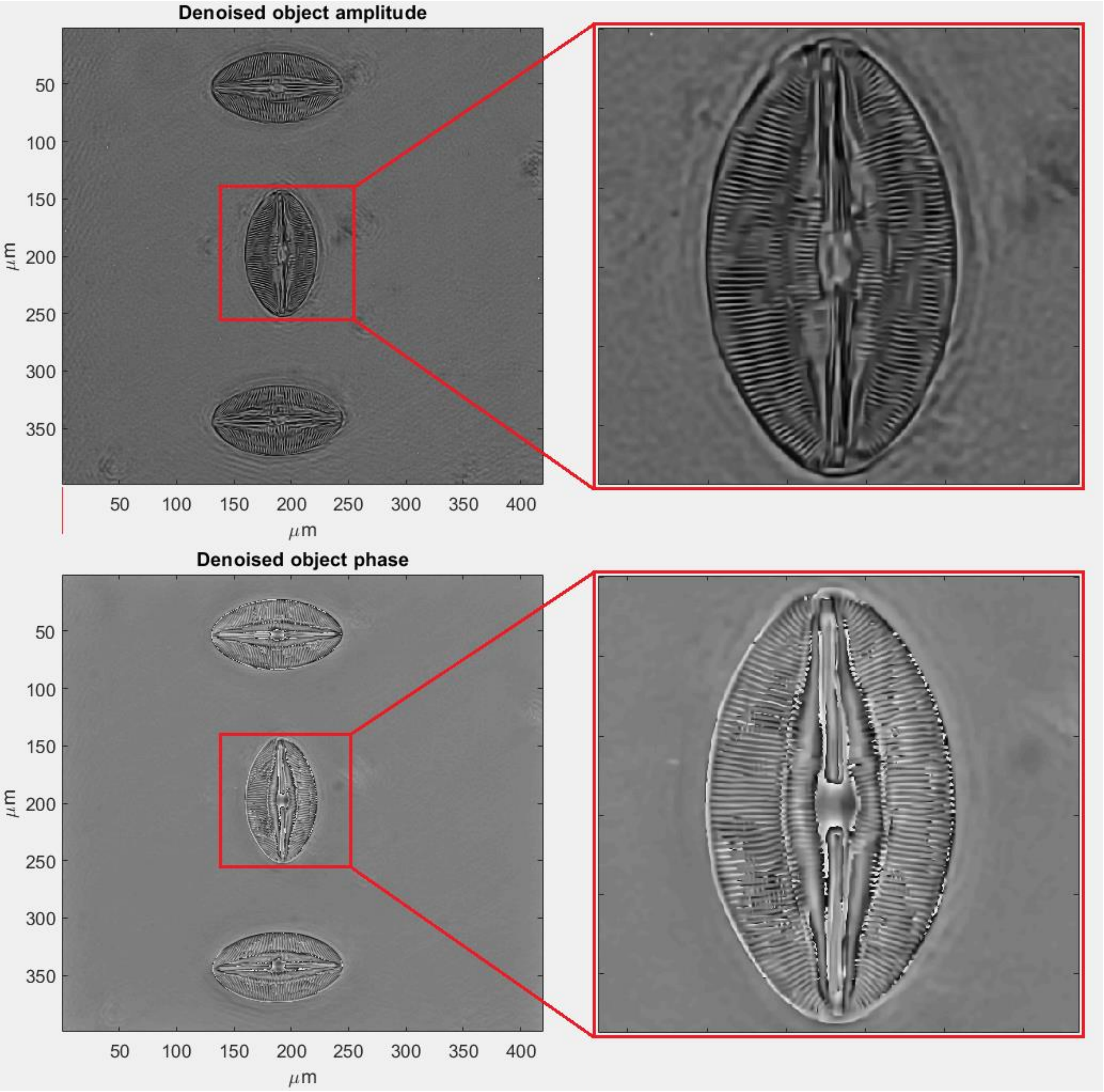

*Figure 36. Denoised amplitude and phase obtained with FPM app denoising procedure described in this chapter. Amplitude displaying range was narrowed from 0.2-4.1 to 0.2-1.6 to increase displaying contrast (all Navicula details fit in this narrowed range).*

- LED position correction

If performed reconstruction is disturbed by unwanted fringes, it may be due to misalignment error (real illumination angles may vary from the angles that result from input system parameters). One of the reasons that causes this error may be wrong distance between LED array and measured object. It is always worth to check if another distance (**LED height** field in **System setup** panel) gives better results (Figure 37).

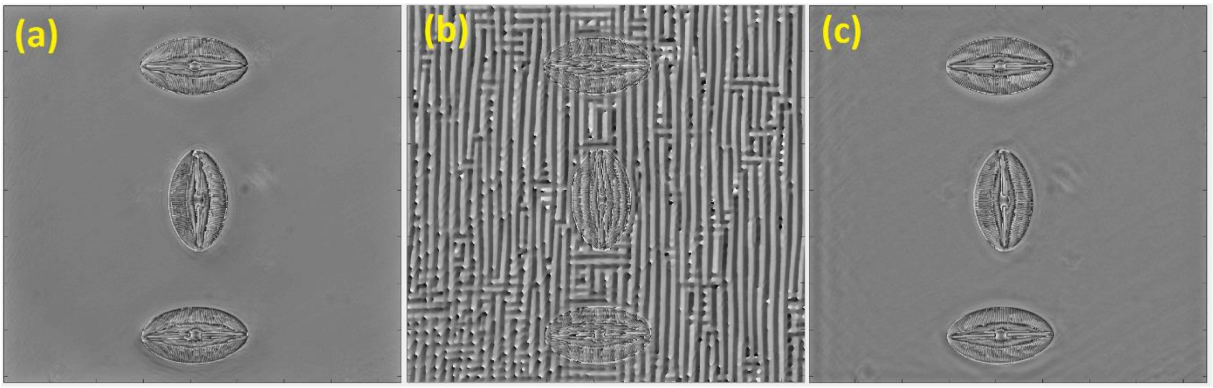

*Figure 37. Reconstructed phase of Navicula elliptica for setting in FPM app different distances LED array – object. (a) – theoretically correct distance = 43 mm; (b) distance = 35 mm; (c) distance = 51 mm.*

There are many other sources of misalignment errors (LED array shift or rotation, distance tolerance between adjacent LEDs), which may be corrected by one of the implemented LED position correction algorithms. This correction may be performed with the use of the procedure listed below (this procedure should be the fastest that gives satisfying misalignment correction in *FPM app*):

1. Perform all steps as in exemplary reconstruction procedure up to point 13.
2. Push **Select on image** button in **ROI selection** panel to open *ROI selection* window.
3. In *ROI selection* window select a small ROI that contains at least part of measured object (in this example ROI size = 200x200).

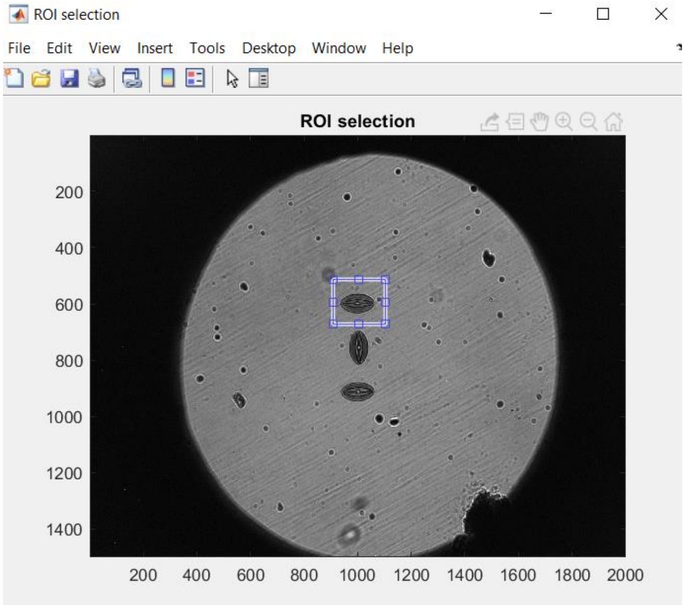

4. In **Options panel** in **LED correction method** dropdown select one of the position correction algorithms (due to shortest computing time we recommend **Authors method [fast]** algorithm).

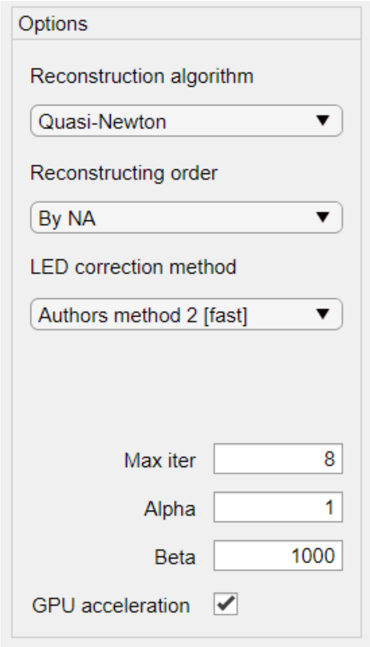

5. Push **Run algorithm** button to perform reconstruction (it may take up to a few minutes).
6. When reconstruction finishes push **Save results** button and save the reconstruction results.

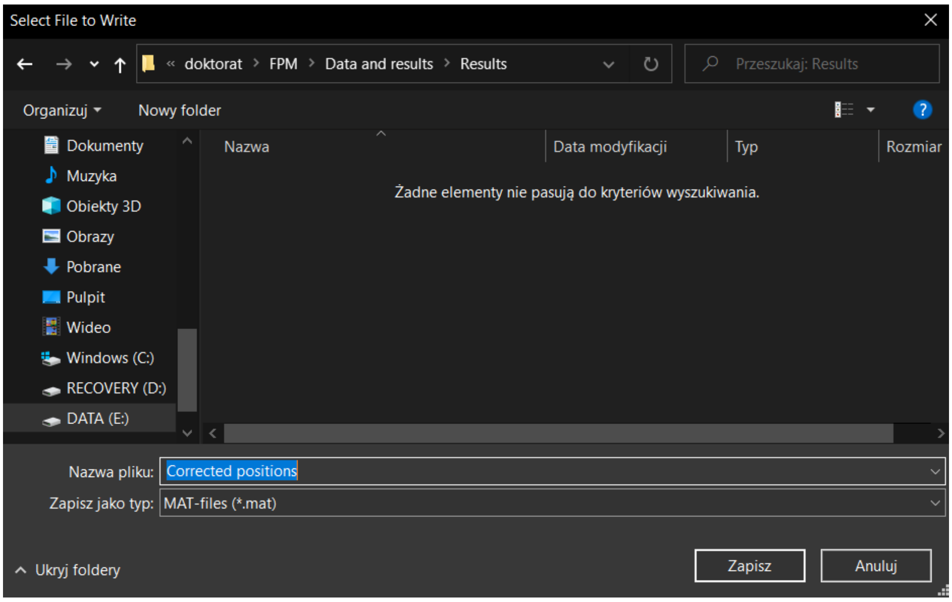

7. Repeat step 3 but this time select ROI that you want to reconstruct.

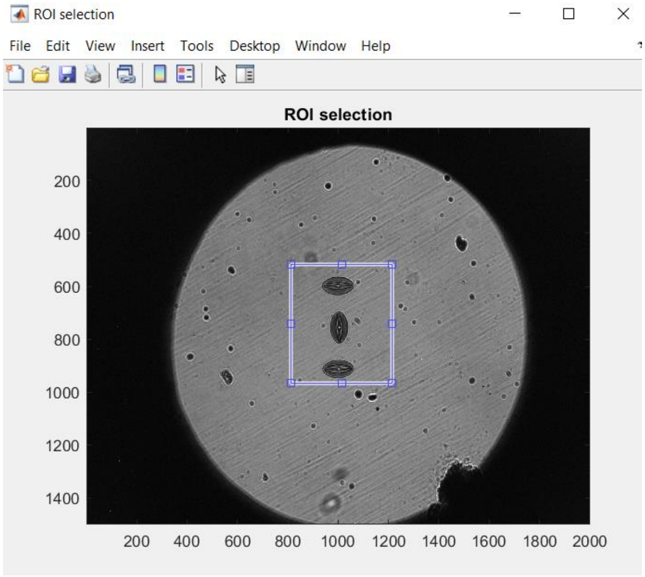

8. In **Options panel** in **LED correction method** dropdown select **None** option.

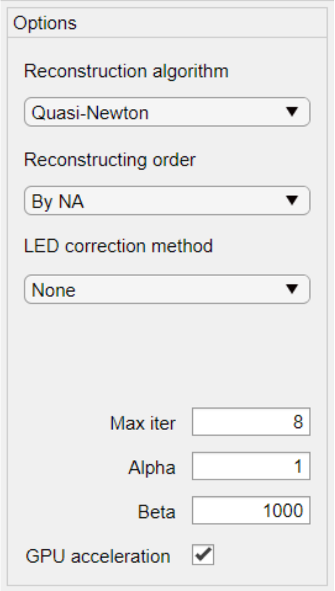

9. Check **Load corrected LEDs positions** checkbox and then push **Run algorithm** button to perform another reconstruction.

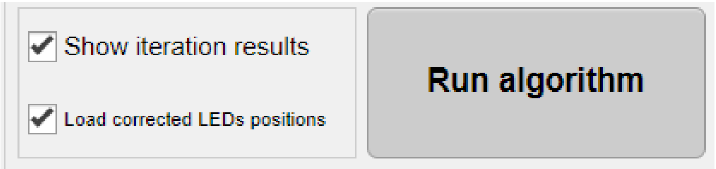

10. At the beginning of reconstruction there will pop-up Select file to open window. Select the file saved in step 6. It will cause that during reconstruction LEDs positions (scaled to current ROI size) will be used that were obtained in step 5 reconstruction with LED position correction.

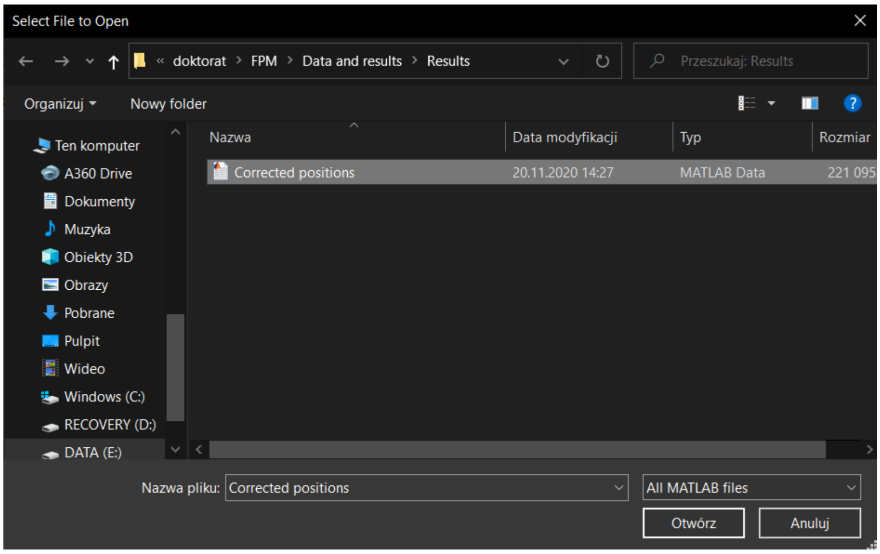

Reconstruction results of misalignment error correction obtained with above procedure are shown in Figure 38 (misalignment is introduced by setting as input incorrect distance LED array –object (35 mm instead of 43 mm)). Correcting misalignment error (step 5) lasted 75 s, reconstructing of the data with corrected positions lasted 70 s.

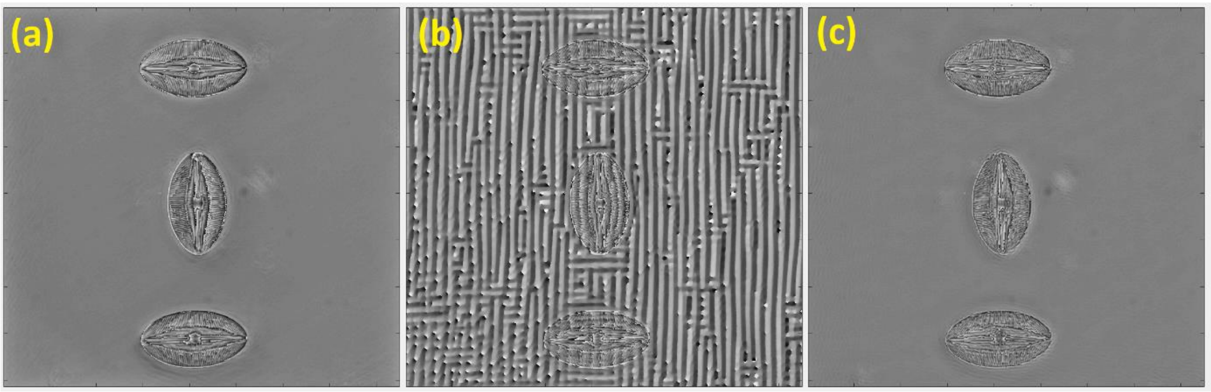

*Figure 38. Reconstructed phase of Navicula elliptica for setting in FPM app different distances LED array – object. (a) – theoretically correct distance = 43 mm; (b) distance = 35 mm; (c) distance = 35 mm but with misalignment error corrected as in procedure described in this chapter.*
