## Supplementary material for "FPM app: an open-source application for simple and intuitive Fourier ptychographic reconstruction": FPM app: Algorithm.m

```
function [o,phase,P,erro,erroS,idx_X,idx_Y] = Algorithm(I, N_obj, idx_X, idx_Y, imageColOrder, recOrder, pupil0, options, cImag, f, showIm, rectype)
% Function that executes FPM algorithms
%   Inputs:
%       I - collected images (3d matrix)
%       N_obj = [N_obj_y,N_obj_y,] - size of the reconstructed image
%       idx_X,idx_Y - LEDs position in Fourier domain
%       imageColOrder - matrix that shows in which order the images were
%                       collected
%           1 - first image; 2 - second image; etc
%           Example:
%               imageColOrder = [0,1,2,3,0
%                                4,5,6,7,8
%                               0,9,10,11,0];
%       recOrder = [first img number, second img number, etc.]' -
%                  reconstruction order
%       pupil0 - initial pupil
%       options - reconstruction options
%           options.alpha - regularization parameter for object reconstruction
%           options.beta - regularization parameter for pupil reconstruction
%           options.maxIter - maximum number of iterations
%           options.algorithm - select reconstruction algorithm
%               1 - Quasi-Newton algorithm
%               2 - Gerchberg-Saxton
%           options.LEDcorrection - LED position correction
%               1 - Angle Self-Calibration
%               2 - simulated annealing
%               3 - genetic algorithm
%           options.useGPU
%               1 - use GPU acceleration
%               0 - don't use
%       cImag - central image number
%   Outputs:
%       o - reconstructed object
%       P - reconstructed pupil
%       erro - RMS error (compared to input data)
%       erroS - RMS error (compared to known synthetic object)

%% Initialization
% Checking if there is created synthetic object
if rectype == 1
    ampl = evalin('base','amplitude_synth');
    phs = evalin('base','phase_synth');
    obj = ampl.*exp(1i.* phs);
end

[Nmy,Nmx,~] = size(I);
Np = [Nmy,Nmx];

cen0 = round((N_obj+1)/2);

FT = @(x) fftshift(fft2(ifftshift(x)));
IFT = @(x) fftshift(ifft2(ifftshift(x)));

% operator to crop region of O from proper location at the O plane
downsamp = @(x,cen) x(cen(1)-floor(Np(1)/2):cen(1)-floor(Np(1)/2)+Np(1)-1,...
    cen(2)-floor(Np(2)/2):cen(2)-floor(Np(2)/2)+Np(2)-1);

%% initialization in FT domain
% initial guess
if options.useGPU == 1
    O = gpuArray(zeros(N_obj));
else
    O = zeros(N_obj);
end

if isempty(cImag) % no central image used in reconstruction
    Os = FT(sqrt(I(:,:,recOrder(1))));  % First image is the initial guess
    [ledsY,ledsX] = find(imageColOrder == recOrder(1));
else % central image used in reconstruction
    Os = FT(sqrt(I(:,:,cImag)));  % Central image is the initial guess
    [ledsY,ledsX] = find(imageColOrder == cImag);
end
nXs = idx_X(ledsY,ledsX);
nYs = idx_Y(ledsY,ledsX);

cen = cen0 + [nYs,nXs];   % Position in Fourier domain of the initial guess
n1 = cen-floor(Np/2);
n2 = n1 + Np-1;

% Reconstruction object matrix (Fourier domain)
O(n1(1):n2(1),n1(2):n2(2)) = Os.*pupil0;

% Pupil matrix
P = pupil0;

% RMS error in compared to input data in function of iteration number
erro = [];
% RMS error in compared to synthetic object in function of iteration
% number
erroS = [];
iter = 0;

% displaying results
if showIm == 1
    o = gather(O);
    o = IFT(o);
    figure(88);
    set(gcf,'Name','Iteration results');
    set(gcf,'NumberTitle','off');
    subplot(2,3,1); imagesc(abs(o)); colormap gray; colorbar;
    title('object amplitude');
    subplot(2,3,2); imagesc(angle(o)); colormap gray; colorbar;
    title('object phase');
    subplot(2,3,4); imagesc(abs(P)); colormap gray; colorbar;
    title('pupil amplitude');
    subplot(2,3,5); imagesc(angle(P).*abs(P)); colorbar;
    title('pupil phase');
    pause(0.1);
end

idx_X2 = idx_X(:,2:end); idx_Y2 = idx_Y(2:end,:);
% max distance in Fourier space between adjacent LEDs
sp0 = max(max(max(abs(idx_Y2-idx_Y(1:end-1,:)))),...
    max(max(abs(idx_X2-idx_X(:,1:end-1)))));
mx = round(sp0/3);
if mx<3; mx = 3; end
%% main algorithm

if options.LEDcorrection == '2'
    warning('off')
end
try
    waitbar(0,f,'iteration...1');
catch
    f = waitbar(0,'iteration...1');
end
if options.useGPU == 1
    err0 = gpuArray(0);
else
    err0 = 0;
end

switch options.algorithm
    case '1'    % Quasi-Newton Algorithm
        
        while iter<options.maxIter
            
            iter = iter+1;
            for mm = 1:length(recOrder)
                % current image number
                nImg = recOrder(mm);
                % current LED position (in LEDs matrix)
                [ledY,ledX] = find(imageColOrder == nImg);
                
                I_mea = I(:,:,nImg); % measured image
                
                nX = idx_X(ledY,ledX);
                nY = idx_Y(ledY,ledX);
                % current LED position in Fourier domain
                cen = cen0+[nY,nX];
                
                Psi0 = downsamp(O,cen).*P;
                psi0 = IFT(Psi0);   % estimated image (amplitude)
                % estimated image
                I_est = abs(psi0).^2;
                
                % projection 1
                Psi = Proj_Fourier_v2(psi0, I_mea, I_est, 1, FT);
                % projection 2
                dPsi = (Psi-Psi0);
                
                Omax = max(max(abs(O)));
                
                P2 = @(O,P,dpsi,Omax,cen)...
                    GDUpdate_Multiplication_rank1(O,P,dpsi,Omax,cen,pupil0,...
                    options.alpha,options.beta);
                [O,P] = P2(O,P,dPsi,Omax,cen);
                err0(mm) = rms(rms(I_mea - I_est));
                
                % % displaying current Fourier space
                % figure(1); imagesc(log(1+abs(O))); colormap gray;
                % title('intensity reconstruction order')
                % pause(0.1)
                
                LEDsPosCorrection;
            end
            
            o = gather(O);
            o = IFT(o); % reconstructed object
            
            % calculating iteration error
            err00 = rms(err0);
            erro = [erro,err00];
            if rectype == 1   % if there is known synthetic object
                obj = imresize(obj, size(o));
                % normalization
                amps = abs(obj) - min(min(abs(obj)));
                amps = amps./max(max(amps));
                amp = abs(o) - min(min(abs(o)));
                amp = amp./(max(max(amp)));
                erroS = [erroS,rms(rms(amp-amps))];
            end
            
            f = ActualizeWaitbar(iter,options.maxIter,f,options);
            if showIm == 1
                DisplayIterationResult(o,P,erro,rectype,erroS)
            end
        end
        phase = angle(o);
        f = ActualizeWaitbar(iter,options.maxIter,f,options);
        
    case '2'    % Gerchberg-Saxton algorithm
        while iter<options.maxIter
            iter = iter+1;
            for mm = 1:length(recOrder)
                nImg = recOrder(mm);
                [ledY,ledX] = find(imageColOrder == nImg);

                I_mea = I(:,:,nImg);
                Psi_mea = FT(sqrt(I_mea)).*P;
                psi_mea = IFT(Psi_mea);
                
                nX = idx_X(ledY,ledX);
                nY = idx_Y(ledY,ledX);
                cen = cen0+[nY,nX];
                
                Psi0 = downsamp(O,cen).*P;
                psi0 = IFT(Psi0);
                % estimated intensity
                I_est = abs(psi0).^2;
                
                imLowRes = abs(psi_mea).*exp(1i.*angle(psi0));
                nl = cen-floor(Np/2);
                nh = nl+Np-1;
                O(nl(1):nh(1),nl(2):nh(2)) = ...
                    (1-pupil0).*O(nl(1):nh(1),nl(2):nh(2)) + ...
                    FT(imLowRes).*P;
                
                err0(mm) = rms(rms(I_mea-I_est));
                
                LEDsPosCorrection;
            end
            
            o = gather(O);
            o = IFT(o); % reconstructed object
            err00 = rms(err0);
            erro = [erro,err00];
            if rectype == 1   % if there is known synthetic object
                obj = imresize(obj, size(o));
                % normalization
                amps = abs(obj) - min(min(abs(obj)));
                amps = amps./max(max(amps));
                amp = abs(o) - min(min(abs(o)));
                amp = amp./(max(max(amp)));
                erroS = [erroS,rms(rms(amp-amps))];
            end
            f = ActualizeWaitbar(iter,options.maxIter,f,options);
            if showIm == 1
                DisplayIterationResult(o,P,erro,rectype,erroS)
            end
        end
        phase = angle(o);
        f = ActualizeWaitbar(iter,options.maxIter,f,options);
        
    case '3'    % Algorithm template
        
        while iter<options.maxIter
            
            iter = iter+1;
            for mm = 1:length(recOrder)
                % current image number
                nImg = recOrder(mm);
                % current LED position (in LEDs matrix)
                [ledY,ledX] = find(imageColOrder == nImg);
                
                I_mea = I(:,:,nImg); % measured image
                
                nX = idx_X(ledY,ledX);
                nY = idx_Y(ledY,ledX);
                % current LED position in Fourier domain
                cen = cen0+[nY,nX];
                
                
                Psi0 = downsamp(O,cen).*P;
                psi0 = IFT(Psi0);   % estimated image (amplitude)
                % estimated image
                I_est = abs(psi0).^2;
                
                %%%%%%%%%%%%%%%%%%%%%%%%%%%%%%%%%%%%%%%%%%%%%%%%%%%%%%%
                % Here place your own algorithm
                %%%%%%%%%%%%%%%%%%%%%%%%%%%%%%%%%%%%%%%%%%%%%%%%%%%%%%%
                
                LEDsPosCorrection;
            end
            
            o = IFT(O);
            err00 = rms(err0);
            erro = [erro,err00];
            if rectype == 1   % if there is known synthetic object
                obj = imresize(obj, size(o));
                % normalization
                amps = abs(obj) - min(min(abs(obj)));
                amps = amps./max(max(amps));
                amp = abs(o) - min(min(abs(o)));
                amp = amp./(max(max(amp)));
                erroS = [erroS,rms(rms(amp-amps))];
            end
            f = ActualizeWaitbar(iter,options.maxIter,f,options);
            if showIm == 1
                DisplayIterationResult(o,P,erro,rectype,erroS)
            end
        end
        phase = angle(o);
        f = ActualizeWaitbar(iter,options.maxIter,f,options);
end
try
    close(f)
end
try
    close(88)
end
end

function DisplayIterationResult(o,P,erro,s,erroS)
% display iteration results
figure(88);
set(gcf,'Name','Iteration results');
set(gcf,'NumberTitle','off');
subplot(2,3,1); imagesc(abs(o)); colormap gray; colorbar;
title('object amplitude');
subplot(2,3,2); imagesc(angle(o)); colormap gray; colorbar;
title('object phase');
subplot(2,3,3); plot(erro); title('RMSE (compared to input data)');
subplot(2,3,4); imagesc(abs(P)); colormap gray; colorbar;
title('pupil amplitude');
subplot(2,3,5); imagesc(angle(P).*abs(P)); colorbar;
title('pupil phase');
if s == 1
    subplot(2,3,6); plot(erroS);
    title('RMSE (compared to known syntetic object)');
end
pause(0.1);
end

function f = ActualizeWaitbar(iter,maxIter,f,options)
% waitbar
if iter < maxIter
    progress = (iter)/maxIter;
    try
        waitbar(progress,f,strcat('iteration...',num2str(iter+1)));
    catch
        f = waitbar(progress,strcat('iteration...',num2str(iter+1)));
    end
else
    try
        waitbar(1,f,'finish');
    catch
        f = waitbar(1,'finish');
    end
end
end
```
