## Supplementary material for "FPM app: an open-source application for simple and intuitive Fourier ptychographic reconstruction": FPM app: AlgorithmManual.m

```
function [rec_object,phase, rec_pupil,err,erro,idx_X2,idx_Y2,svdIdx] = AlgorithmManual(ImagesIn, LEDs, imageColOrder, LEDsUsed, ROI, systemSetup, options, showIm, svdIdx,rectype)
% Function that runs FPM algorithms
%   Inputs:
%       ImagesIn - collected images (3d matrix)
%       LEDs - LED matrix used to collect images
%           0 in place where there is no LED
%           1 in place where there is LED
%           2 in place where there is central LED
%           Example:
%               LEDs = [0,1,1,1,0
%                       1,1,2,1,1
%                       0,1,1,1,0];
%       imageColOrder - matrix that shows in which order the images were 
%                       collected
%           1 - first image; 2 - second image; etc
%           Example:
%               imageColOrder = [0,1,2,3,0
%                                4,5,6,7,8
%                               0,9,10,11,0];
%       LEDsUSED - matrix that shows which images you want to reconstruct
%           1 - LED used in reconstruction
%           0 - LED not used in reconstruction
%           Example:
%               LEDsUsed = [0,1,1,1,0
%                           1,1,0,1,1
%                           0,1,1,1,0];
%       ROI - Region Of Interest
%           ROI = [x0,y0,xSize,ySize];
%       systemSetup
%           systemSetup.NA - NA
%           systemSetup.lambda - wavelength (um)
%           systemSetup.magnification magnification
%           systemSetup.LEDspacing - spacing between neighbour LEDs in LED
%                                    matrix (mm)
%           systemSetup.LEDheight - distance between LED matrix and a sample
%                               (mm)
%           systemSetup.camPizSize - camera pixel size (um)
%       options - reconstruction options
%           options.alpha - regularization parameter for object reconstruction
%           options.beta - regularization parameter for pupil reconstruction
%           options.maxIter - maximum number of iterations
%           options.algorithm - select reconstruction algorithm
%               1 - Quasi-Newton algorithm
%               2 - Gerchberg-Saxton
%           options.recorder - reconstruction order
%               1 - from lowest to highest NA
%               2 - from the lightest to the darkest images
%           options.LEDcorrection - LED position correction
%               1 - Angle Self-Calibration
%               2 - simulated annealing
%               3 - genetic algorithm
%           options.initialPupil - initial pupil
%               1 - ones
%               2 - tukey window
%               3 - gauss window
%           options.useGPU
%               1 - use GPU acceleration
%               0 - don't use
%   Outputs:
%       rec_object - reconstructed object (complex double)
%       rec_pupil - reconstructed pupil (complex double)
%       err - RMS error (compared to input data)
%       erro - RMS error (compared to known synthetic object)

%% initialization
tic
% profile on
f = waitbar(0,'initialization');

[sy,sx,nImgs] = size(ImagesIn);
bck = zeros(nImgs,1);
if sy>100 && sx>100
    for nn = 1:nImgs; bck(nn) = mean2(ImagesIn(1:100,1:100,nn)); end
else
    for nn = 1:nImgs; bck(nn) = mean2(ImagesIn(:,:,nn)); end    
end
thr = (max(bck)+min(bck))/2;

% preparing input images - ROI cropping, background removing, converting to
% GPU array
[ImagesIn,imageColOrder] = InputImagesCrop(ImagesIn,imageColOrder,LEDsUsed,ROI);
[ImagesIn, bck] = BackgroundRemoving(ImagesIn,thr);
if options.useGPU == 1
    ImagesIn = gpuArray(ImagesIn);
end

% reconstruction order
recOrder = img_order(LEDs, imageColOrder, LEDsUsed, options.recorder, bck);

[cledY,cledX] = find(LEDs == 2);    % central LED position
LEDs(LEDs>1) = 1;

if LEDsUsed(cledY,cledX) == 0
    cImag = [];
else
    cImag = imageColOrder(cledY,cledX); % central image number
end

%% system setup
lambda = systemSetup.lambda;
NA = systemSetup.NA; 
um_m = NA/lambda;   % maximum spatial frequency set by NA
mag = systemSetup.magnification; 
pixSizeCam = systemSetup.camPizSize; % pixel size on the sensor plane
pixSizeObj = pixSizeCam/mag; % effective image pixel size on the object plane

% Field of view in the object space
FoVx = ROI(3)*pixSizeObj;
FoVy = ROI(4)*pixSizeObj;

% sampling size in x direction
if mod(ROI(3),2) == 1
    dux = 1/pixSizeObj/(ROI(3)-1);
else
    dux = 1/FoVx;
end
% sampling size in y direction
if mod(ROI(4),2) == 1
    duy = 1/pixSizeObj/(ROI(4)-1);
else
    duy = 1/FoVy;
end


%% low-pass filter diameter set by the NA = bandwidth of a single measurment
m = 1:ROI(3);
n = 1:ROI(4);
[mm,nn] = meshgrid(m-round((ROI(3)+1)/2),n-round((ROI(4)+1)/2));
if ROI(3)>ROI(4)
    nn = nn.*max(max(abs(mm)))./max(max(abs(nn)));
else
    mm = mm.*max(max(abs(nn)))./max(max(abs(mm)));
end
ridx = sqrt(mm.^2+nn.^2);
um_idx = um_m/min(dux,duy);
pupil0 = double(ridx<um_idx);
% figure; imagesc(pupil0); title ones
pupil0 = initialPupil(pupil0,options.initialPupil);
% figure; imagesc(pupil0);  title tukey
%% LEDs position
center = [round(sy/2),round(sx/2)];
img_center(1) = (ROI(2)-center(1)+ROI(4)/2)*pixSizeObj/1000;
img_center(2) = (ROI(1)-center(2)+ROI(3)/2)*pixSizeObj/1000;

[ys,xs] = size(LEDs);
LEDspacing = systemSetup.LEDspacing;    % spacing between adjacent LEDs
LEDheight = systemSetup.LEDheight;  % distance bewteen the LED matrix and the sample
% for LEDheight = 37.5:0.5:42.5
xx = 1:xs; xx = (xx - cledX).*LEDspacing;
yy = 1:ys; yy = (yy - cledY).*LEDspacing;
[LEDsPosX,LEDsPosY] = meshgrid(xx,yy);  % LEDs position in x and y
% LEDsPosX = LEDsPosX - 1;
% distances between LEDs and sample
dist = sqrt((LEDsPosX-img_center(2)).^2+(LEDsPosY-img_center(1)).^2 ...
    + LEDheight.^2);
% corresponding angles for each LEDs
sin_thetaX = LEDsPosX./dist;
sin_thetaY = LEDsPosY./dist;

illuminationNA = sqrt(sin_thetaX.^2+sin_thetaY.^2);
illuminationNA = illuminationNA.*LEDsUsed;

% corresponding spatial freq for each LEDs
xFreq = sin_thetaX/lambda;
yFreq = sin_thetaY/lambda;
% spatial freq index for each plane wave relative to the center
idx_Y = round(yFreq/duy); 
idx_X = round(xFreq/dux);

% loading input LEDs positions
if ~isempty(svdIdx)
    idx_Y = round(svdIdx.idx_Y.*ROI(4)./svdIdx.ROI(4));
    idx_X = round(svdIdx.idx_X.*ROI(3)./svdIdx.ROI(3));
    idx_Y = idx_Y - idx_Y(cledY,cledX);
    idx_X = idx_X - idx_X(cledY,cledX);
end

% % number of brightfield images
% NBF = sum(sum(illuminationNA<NA));

% maxium spatial frequency achievable based on the maximum illumination
% angle from the LED array and NA of the objective
um_p = max(max(illuminationNA))/lambda+um_m;

disp(['synthetic NA is ',num2str(um_p*lambda)]);

% assume the max spatial freq of the original object
% um_obj>um_p
% assume the # of pixels of the original object in x direction
N_objX2 = round(2*um_p/dux)*2;

% need to enforce N_obj/Np = integer to ensure no FT artifacts
N_objX = ceil(N_objX2/ROI(3))*ROI(3);
if N_objX == ROI(3)
    N_objX = N_objX*2;
end
N_objY = ROI(4)*N_objX/ROI(3);

%% reconstruction algorithm

[rec_object,phase,rec_pupil, err, erro,idx_X2,idx_Y2] = ... 
    Algorithm(ImagesIn, [N_objY,N_objX], idx_X, idx_Y, imageColOrder, ...
    recOrder, pupil0, options, cImag, f, showIm, rectype);

% corrected LEDs positions
svdIdx.idx_X = idx_X2;
svdIdx.idx_Y = idx_Y2;
svdIdx.ROI = ROI;

toc
end
```
