## Supplementary material for "FPM app: an open-source application for simple and intuitive Fourier ptychographic reconstruction": FPM app: BackgroundRemoving.m

```
function [I, bck] = BackgroundRemoving(ImagesIn, thr0)
% Function that removes background from FPM data
%   Inputs:
%       ImagesIn - vector of input images
%       thr0 - threshold value to distinguish darkfield images from
%              brightfields images
%   Outputs:
%       I - vector of background removed input images
%       bck - vector of background values

[nY,nX,nImgs] = size(ImagesIn);
% nY = 500; nX = 500;
%% creating edge map
edgeMap = zeros(nY,nX);
for m = 1:nImgs
    edgeMap = edgeMap + double(ImagesIn(:,:,m))./mean2(ImagesIn(:,:,m));
end

%% looking for background region

% default background regions size 50x50
si = 50;
% if small images, background region <= 1% of the image size
if min(nY,nX)/10 < 50
    si = round(min(nY,nX)/10);
end

bckReg = zeros(floor(nY/si),floor(nX/si));

for m1 = 1:si:nY-si+1
    for m2 = 1:si:nX-si+1
        ref(1:si,1:si) = mean2(edgeMap(m1:m1+si-1,m2:m2+si-1));
        bckReg((m1-1)/si+1,(m2-1)/si+1) = immse(double(edgeMap(m1:m1+si-1,m2:m2+si-1)),ref);
    end
end
[y,x] = find(bckReg == min(min(bckReg)));
locy = y; locx = x;

% % visualization
% figure(13);
% imagesc(edgeMap,[100 300]); title('edge map'); colormap gray
% figure(14);
% imagesc(ImagesIn(:,:,69)); colormap gray; title('central image')
% cimag = ImagesIn(:,:,69);
% y = y.*si-round(si/2);
% x = x.*si-round(si/2);
% figure(13);
% viscircles([x,y],round(si/2))
% figure(14);
% viscircles([x,y],round(si/2))
% amp = evalin('base','amplitude');
% figure(1); imagesc(phs); colormap gray;
% viscircles([x*3,y*3],round(si/2*3)); title('real phase')
%% calculating background values

% background value for each image
background = zeros(nImgs,1);

for m = 1:nImgs
    background(m) = mean2(ImagesIn(si*locy-si+1:si*locy,si*locx-si+1:si*locx,m));
end


% figure; plot(background);
bck = background;
if max(max(bck)) > thr0 % if there are brightfield iages
    threshold = (max(background) + min(background)) / 2;
    % if first image is brightfield image
    if background(1) > threshold
        background(1) = mean(background<threshold);
    end
    for m = 2:nImgs
        if background(m) > threshold % if this is brightfield image
            background(m) = background(m-1);
        end
    end
end
% figure; plot(background); title('Calculated background');
% xlabel('image number'); ylabel('background value')

%% removing background
I = double(ImagesIn);
if min(min(bck)) < thr0 % if there are darkfield images
    for m = 1:nImgs
        Itmp = I(:,:,m) - background(m);
        Itmp(Itmp<0) = 0;
        I(:,:,m) = Itmp;
    end
end
end
```
