## Supplementary material for "FPM app: an open-source application for simple and intuitive Fourier ptychographic reconstruction": FPM app: demo_BM3DSAPCA.m

```
% BM3D-SAPCA : BM3D with Shape-Adaptive Principal Component Analysis  (v1.00, 2009)
% (demo script)
%
% BM3D-SAPCA is an algorithm for attenuation of additive white Gaussian noise (AWGN)
% from grayscale images. This algorithm reproduces the results from the article:
%  K. Dabov, A. Foi, V. Katkovnik, and K. Egiazarian, "BM3D Image Denoising with
%  Shape-Adaptive Principal Component Analysis", Proc. Workshop on Signal Processing
%  with Adaptive Sparse Structured Representations (SPARS'09), Saint-Malo, France,
%  April 2009.     (PDF available at  http://www.cs.tut.fi/~foi/GCF-BM3D )
%
%
% SYNTAX:
%
%     y_est = BM3DSAPCA2009(z, sigma)
%
% where  z  is an image corrupted by AWGN with noise standard deviation  sigma
% and  y_est  is an estimate of the noise-free image.
% Signals are assumed on the intensity range [0,1].
%
%
% USAGE EXAMPLE:
%
%     y = im2double(imread('Cameraman256.png'));
%     sigma=25/255;
%     z=y+sigma*randn(size(y));
%     y_est = BM3DSAPCA2009(z,sigma);
%
%
%
% Copyright (c) 2009-2011 Tampere University of Technology.   All rights reserved.
% This work should only be used for nonprofit purposes.
%
% author:  Alessandro Foi,  
%
%%

clear all

y = im2double(imread('Cameraman256.png'));
% y = im2double(imread('Lena512.png'));
randn('seed',0);

sigma=25/255;
z=y+sigma*randn(size(y));

y_est = BM3DSAPCA2009(z,sigma);

PSNR = 10*log10(1/mean((y(:)-y_est(:)).^2));
disp(['PSNR = ',num2str(PSNR)])
if exist('ssim_index')
    [mssim ssim_map] = ssim_index(y*255, y_est*255);
    disp(['SSIM = ',num2str(mssim)])
end
```
