## Supplementary material for "FPM app: an open-source application for simple and intuitive Fourier ptychographic reconstruction": FPM app: function_CreateLPAKernels.m

```
% Creates LPA kernels cell array    (function_CreateLPAKernels)
%
% Alessandro Foi - Tampere University of Technology - 2003-2005
% ---------------------------------------------------------------
%
%  Builds kernels cell arrays kernels{direction,size}
%                  and        kernels_higher_order{direction,size,1:2}
%               kernels_higher_order{direction,size,1}  is the 3D matrix
%                   of all kernels for that particular direction/size
%               kernels_higher_order{direction,size,2}  is the 2D matrix
%                   containing the orders indices for the kernels
%                   contained in kernels_higher_order{direction,size,1}
%
%   ---------------------------------------------------------------------
% 
%   kernels_higher_order{direction,size,1}(:,:,1) is the funcion estimate kernel
%   kernels_higher_order{direction,size,1}(:,:,2) is a first derivative estimate kernel
%
%   kernels_higher_order{direction,size,1}(:,:,n) is a higher order derivative estimate kernel
%   whose orders with respect to x and y are specified in
%   kernels_higher_order{direction,size,2}(n,:)=
%                           =[xorder yorder xorder+yorder]
%   
%%%%%%%%%%%%%%%%%%%%%%%%%%%%%%%%%%%%%%%%%%%%%%%%%%%%%%%%%%%%%%%%%%%%%%%%%%%%%%%%%%%%%%%%%%%%%%%%%%%%%%%%%%%%%%%%%%%%%%%%

function [kernels, kernels_higher_order]=function_createLPAkernels(m,h1,h2,TYPE,window_type,directional_resolution,sig_winds,beta)

%--------------------------------------------------------------------------
% LPA ORDER AND KERNELS SIZES
%--------------------------------------------------------------------------
%    m=[2,0];        % THE VECTOR ORDER OF LPA;

%    h1=[1 2 3 4 5];    %   sizes of the kernel
%    h2=[1 2 3 4 5];    %   row vectors h1 and h2 need to have the same lenght


%--------------------------------------------------------------------------
% WINDOWS PARAMETERS
%--------------------------------------------------------------------------
%    sig_winds=[h1*1 ; h1*1];    % Gaussian parameter
%    beta=1;                     % Parameter of window 6

%    window_type=1 ;  % window_type=1 for uniform, window_type=2 for Gaussian
% window_type=6 for exponentions with beta
% window_type=8 for Interpolation

%    TYPE=00;        % TYPE IS A SYMMETRY OF THE WINDOW
                     % 00 SYMMETRIC
                     % 10 NONSYMMETRIC ON X1 and SYMMETRIC ON X2
                     % 11 NONSYMMETRIC ON X1,X2  (Quadrants)
                     % 
                     % for rotated directional kernels the method that is used for rotation can be specified by adding 
                     % a binary digit in front of these types, as follows:
                     % 
                     % 10 
                     % 11  ARE "STANDARD" USING NN (Nearest Neighb.) (you can think of these numbers with a 0 in front)
                     % 00
                     % 
                     % 110
                     % 111  ARE EXACT SAMPLING OF THE EXACT ROTATED KERNEL
                     % 100
                     % 
                     % 210
                     % 211  ARE WITH BILINEAR INTERP
                     % 200
                     % 
                     % 310
                     % 311  ARE WITH BICUBIC INTERP (not reccomended)
                     % 300

%--------------------------------------------------------------------------
% DIRECTIONAL PARAMETERS
%--------------------------------------------------------------------------
%    directional_resolution=4;       % number of directions


%%%%%%%%%%%%%%%%%%%%%%%%%%%%%%%%%%%%%%%%%%%%%%%%%%%%%%%%%%%%%%%%%%%%%%%%%%%%%%%%%%%%%%%%%%%%%%%%%%%%%%%%%%%%%%%%%%%%%%%%%%%%
%%%%%%%%%%%%%%%%%%%%%%%%%%%%%%%%%%%%%%%%%%%%%%%%%%%%%%%%%%%%%%%%%%%%%%%%%%%%%%%%%%%%%%%%%%%%%%%%%%%%%%%%%%%%%%%%%%%%%%%%%%%%
%%%%%%%%%%%%%%%%%%%%%%%%%%%%%%%%%%%%%%%%%%%%%%%%%%%%%%%%%%%%%%%%%%%%%%%%%%%%%%%%%%%%%%%%%%%%%%%%%%%%%%%%%%%%%%%%%%%%%%%%%%%%
%%%%%%%%%%%%%%%%%%%%%%%%%%%%%%%%%%%%%%%%%%%%%%%%%%%%%%%%%%%%%%%%%%%%%%%%%%%%%%%%%%%%%%%%%%%%%%%%%%%%%%%%%%%%%%%%%%%%%%%%%%%%
%%%%% From this point onwards this file and the create_LPA_kernels.m should be identical %%%%%%%%%%%%%%%%%%%%%%%%%%%%%%%%%%%
%%%%%%%%%%%%%%%%%%%%%%%%%%%%%%%%%%%%%%%%%%%%%%%%%%%%%%%%%%%%%%%%%%%%%%%%%%%%%%%%%%%%%%%%%%%%%%%%%%%%%%%%%%%%%%%%%%%%%%%%%%%%


lenh=max(length(h1),length(h2));
clear kernels
clear kernels_higher_order
kernels=cell(directional_resolution,lenh);
kernels_higher_order=cell(directional_resolution,lenh,2);
THETASTEP=2*pi/directional_resolution;
THETA=[0:THETASTEP:2*pi-THETASTEP];


s1=0;
for theta=THETA,
    s1=s1+1;
    for s=1:lenh,
        
        
        [gh,gh1,gh1degrees]=function_LPAKernelMatrixTheta(ceil(h2(s)),ceil(h1(s)),window_type,[sig_winds(1,s) sig_winds(2,s)],TYPE,theta, m);
        kernels{s1,s}=gh;                          % degree=0 kernel
        kernels_higher_order{s1,s,1}=gh1;          % degree>=0 kernels
        kernels_higher_order{s1,s,2}=gh1degrees;   % polynomial indexes matrix
       
end   % different lengths loop
end   % different directions loop
```
