## Supplementary material for "FPM app: an open-source application for simple and intuitive Fourier ptychographic reconstruction": FPM app: function_LPAKernelMatrixTheta.m

```
% Return the discrete kernels for LPA estimation and their degrees matrix
%
% function [G, G1, index_polynomials]=function_LPAKernelMatrixTheta(h2,h1,window_type,sig_wind,TYPE,theta, m)
%
%
% Outputs:
%
% G  kernel for function estimation
% G1 kernels for function and derivative estimation
%       G1(:,:,j), j=1 for function estimation, j=2 for d/dx, j=3 for d/dy,
%                contains 0 and all higher order kernels (sorted by degree:
%                 1 x y y^2 x^3 x^2y xy^2 y^3 etc...)
% index_polynomials  matrix of degrees first column x powers, second
%                    column y powers, third column total degree
%
%
% Inputs:
%
% h2, h1  size of the kernel (size of the "asymmetrical portion")
% m=[m(1) m(2)] the vector order of the LPA    any order combination should work 
% "theta" is an angle of the directrd window
% "TYPE" is a type of the window support
% "sig_wind" - vector - sigma parameters of the Gaussian wind
% "beta"- parameter of the power in some weights for the window function
%           (these last 3 parameters are fed into function_Window2D function)
%
%
% Alessandro Foi, 6 march 2004

function [G, G1, index_polynomials]=function_LPAKernelMatrixTheta(h2,h1,window_type,sig_wind,TYPE,theta, m)
global beta

%G1=0;
m(1)=min(h1,m(1));
m(2)=min(h2,m(2));

% builds ordered matrix of the monomes powers
number_of_polynomials=(min(m)+1)*(max(m)-min(m)+1)+(min(m)+1)*min(m)/2;   %   =size(index_polynomials,1) 
index_polynomials=zeros(number_of_polynomials,2);
index3=1;
for index1=1:min(m)+1
    for index2=1:max(m)+2-index1
        index_polynomials(index3,:)=[index1-1,index2-1];
        index3=index3+1;
    end
end
if m(1)>m(2)
    index_polynomials=fliplr(index_polynomials);
end
index_polynomials(:,3)=index_polynomials(:,1)+index_polynomials(:,2);    %calculates degrees of polynomials
index_polynomials=sortrows(sortrows(index_polynomials,2),3);             %sorts polynomials by degree (x first)

%=====================================================================================================================================

halfH=max(h1,h2);
H=-halfH+1:halfH-1;


% creates window function and then rotates it
%   win_fun=zeros(halfH-1,halfH-1);
for x1=H
    for x2=H
        if TYPE==00|TYPE==200|TYPE==300 % SYMMETRIC WINDOW
            win_fun1(x2+halfH,x1+halfH)=function_Window2D(x1/h1/(1-1000*eps),x2/h2/(1-1000*eps),window_type,sig_wind,beta,h2/h1); % weight        
        end
        if TYPE==11|TYPE==211|TYPE==311 % NONSYMMETRIC ON X1,X2 WINDOW
            win_fun1(x2+halfH,x1+halfH)=(x1>=-0.05)*(x2>=-0.05)*function_Window2D(x1/h1/(1-1000*eps),x2/h2/(1-1000*eps),window_type,sig_wind,beta,h2/h1); % weight
        end
        if TYPE==10|TYPE==210|TYPE==310 % NONSYMMETRIC ON X1 WINDOW
            win_fun1(x2+halfH,x1+halfH)=(x1>=-0.05)*function_Window2D(x1/h1/(1-1000*eps),x2/h2/(1-1000*eps),window_type,sig_wind,beta,h2/h1); % weight
        end
        
        if TYPE==100|TYPE==110|TYPE==111 % exact sampling
            xt1=x1*cos(-theta)+x2*sin(-theta);
            xt2=x2*cos(-theta)-x1*sin(-theta);
            if TYPE==100 % SYMMETRIC WINDOW
                win_fun1(x2+halfH,x1+halfH)=function_Window2D(xt1/h1/(1-1000*eps),xt2/h2/(1-1000*eps),window_type,sig_wind,beta,h2/h1); % weight        
            end
            if TYPE==111 % NONSYMMETRIC ON X1,X2 WINDOW
                
                win_fun1(x2+halfH,x1+halfH)=(xt1>=-0.05)*(xt2>=-0.05)*function_Window2D(xt1/h1/(1-1000*eps),xt2/h2/(1-1000*eps),window_type,sig_wind,beta,h2/h1); % weight
            end
            if TYPE==110 % NONSYMMETRIC ON X1 WINDOW
                
                win_fun1(x2+halfH,x1+halfH)=(xt1>=-0.05)*function_Window2D(xt1/h1/(1-1000*eps),xt2/h2/(1-1000*eps),window_type,sig_wind,beta,h2/h1); % weight
            end
        end
        
    end
end
win_fun=win_fun1;
if (theta~=0)&(TYPE<100)
    win_fun=imrotate(win_fun1,theta*180/pi,'nearest');     % use 'nearest' or 'bilinear' for different interpolation schemes ('bicubic'...?)
end
if (theta~=0)&(TYPE>=200)&(TYPE<300)
    win_fun=imrotate(win_fun1,theta*180/pi,'bilinear');     % use 'nearest' or 'bilinear' for different interpolation schemes ('bicubic'...?)
end
if (theta~=0)&(TYPE>=300)
    win_fun=imrotate(win_fun1,theta*180/pi,'bicubic');     % use 'nearest' or 'bilinear' for different interpolation schemes ('bicubic'...?)
end


% make the weight support a square
win_fun2=zeros(max(size(win_fun)));
win_fun2((max(size(win_fun))-size(win_fun,1))/2+1:max(size(win_fun))-((max(size(win_fun))-size(win_fun,1))/2),(max(size(win_fun))-size(win_fun,2))/2+1:max(size(win_fun))-((max(size(win_fun))-size(win_fun,2))/2))=win_fun;
win_fun=win_fun2;


%=====================================================================================================================================


%%%%  rotated coordinates
H=-(size(win_fun,1)-1)/2:(size(win_fun,1)-1)/2;
halfH=(size(win_fun,1)+1)/2;
h_radious=halfH;
Hcos=H*cos(theta); Hsin=H*sin(theta);


%%%% Calculation of FI matrix
FI=zeros(number_of_polynomials);
i1=0;
for s1=H
    i1=i1+1;
    i2=0;
    for s2=H
        i2=i2+1;
        x1=Hcos(s1+h_radious)-Hsin(s2+h_radious);
        x2=Hsin(s1+h_radious)+Hcos(s2+h_radious);
        phi=sqrt(win_fun(s2+halfH,s1+halfH))*(prod(((ones(number_of_polynomials,1)*[x1 x2]).^index_polynomials(:,1:2)),2)./prod(gamma(index_polynomials(:,1:2)+1),2).*(-ones(number_of_polynomials,1)).^index_polynomials(:,3));
        FI=FI+phi*phi';
    end % end of s2
end % end of s1

%FI_inv=((FI+1*eps*eye(size(FI)))^(-1));  % invert FI matrix
FI_inv=pinv(FI);   % invert FI matrix (using pseudoinverse)
G1=zeros([size(H,2) size(H,2) number_of_polynomials]);

%%%% Calculation of mask
i1=0;
for s1=H
    i1=i1+1;
    i2=0;
    for s2=H
        i2=i2+1;
        x1=Hcos(s1+h_radious)-Hsin(s2+h_radious);
        x2=Hsin(s1+h_radious)+Hcos(s2+h_radious);
        phi=FI_inv*win_fun(s2+halfH,s1+halfH)*(prod(((ones(number_of_polynomials,1)*[x1 x2]).^index_polynomials(:,1:2)),2)./prod(gamma(index_polynomials(:,1:2)+1),2).*(-ones(number_of_polynomials,1)).^index_polynomials(:,3));
        G(i2,i1,1)=phi(1);              %   Function Est
        G1(i2,i1,:)=phi(:)';  % Function est & Der est on X Y etc...
    end % end of s1
end % end of s2
%keyboard
```
