## Supplementary material for "FPM app: an open-source application for simple and intuitive Fourier ptychographic reconstruction": FPM app: function_Window2D.m

```
% Returns a scalar/matrix weights (window function) for the LPA estimates
% function w=function_Window2D(X,Y,window,sig_wind, beta);
% X,Y scalar/matrix variables
% window - type of the window weight
% sig_wind - std scaling for the Gaussian ro-weight
% beta -parameter of the degree in the weights
%----------------------------------------------------------------------------------
% V. Katkovnik & A. Foi - Tampere University of Technology -  2002-2005


function w=function_Window2D(X,Y,window,sig_wind, beta,ratio);

if nargin == 5
    ratio=1;
end

IND=(abs(X)<=1)&(abs(Y)<=1);
IND2=((X.^2+Y.^2)<=1);
IND3=((X.^2+(Y*ratio).^2)<=1);
   

if window==1           % rectangular symmetric window
w=IND; end

if window==2   %Gaussian
  
X=X/sig_wind(1);
Y=Y/sig_wind(2);
w = IND.*exp(-(X.^2 + Y.^2)/2); %*(abs(Y)<=0.1*abs(X));%.*IND2; %((X.^2+Y.^2)<=1); 
end

if window==3  % Quadratic window
    w=(1-(X.^2+Y.^2)).*((X.^2+Y.^2)<=1); end

if window==4           % triangular symmetric window
 w=(1-abs(X)).*(1-abs(Y)).*((X.^2+Y.^2)<=1); end
  
    
if window==5           % Epanechnikov symmetric window
  w=(1-X.^2).*(1-Y.^2).*((X.^2+Y.^2)<=1); 
end

if window==6   % Generalized Gaussian
  
X=X/sig_wind;
Y=Y/sig_wind;
w = exp(-((X.^2 + Y.^2).^beta)/2).*((X.^2+Y.^2)<=1); end


if window==7
      
X=X/sig_wind;
Y=Y/sig_wind;
w = exp(-abs(X) - abs(Y)).*IND; end

if window==8 % Interpolation
    
w=(1./(abs(X).^4+abs(Y).^4+0.0001)).*IND2; 
end

if window==9 % Interpolation
    
    NORM=(abs(X)).^2+(abs(Y)).^2+0.0001;
w=(1./NORM.*(1-sqrt(NORM)).^2).*(NORM<=1); 
end

if window==10
    w=((X.^2+Y.^2)<=1);
end


if window==11
  
temp=asin(Y./sqrt(X.^2+Y.^2+eps));
temp=temp*0.6; % Width of Beam
temp=(temp>0)*min(temp,1)+(temp<=0)*max(temp,-1);

w=max(0,IND.*cos(pi*temp));   
    
  
end

    

if window==111
  
temp=asin(Y./sqrt(X.^2+Y.^2+eps));
temp=temp*0.8; % Width of Beam
temp=(temp>0)*min(temp,1)+(temp<=0)*max(temp,-1);

w=max(0,IND3.*(cos(pi*temp)>0));   
% w=((X.^2+Y.^2)<=1);    
end

if window==112
  
temp=atan(Y/(X+eps));
%temp=temp*0.8; % Width of Beam
%temp=(temp>0)*min(temp,1)+(temp<=0)*max(temp,-1);
w=max(0,IND3.*((abs(temp))<=pi/4));   
% w=((X.^2+Y.^2)<=1);    

end

    
    
    
    
        
    
    
    


 
```
