## Supplementary material for "FPM app: an open-source application for simple and intuitive Fourier ptychographic reconstruction": FPM app: README-BM3D-SAPCA.txt

--------------------------------------------------------------------

 BM3D-SAPCA : BM3D with Shape-Adaptive Principal Component Analysis
                            v1.00, 2009

--------------------------------------------------------------------

Copyright (c) 2009-2011 Tampere University of Technology. 
All rights reserved.
This work should be used for nonprofit purposes only.

Author:                      Alessandro Foi

BM3D web page:               http://www.cs.tut.fi/~foi/GCF-BM3D


BM3D-SAPCA is an algorithm for attenuation of additive white
Gaussian noise (AWGN) from grayscale images.


This software package reproduces the results from the article:

 K. Dabov, A. Foi, V. Katkovnik, and K. Egiazarian,
 "BM3D Image Denoising with Shape-Adaptive Principal
 Component Analysis", Proc. Workshop on Signal Processing
 with Adaptive Sparse Structured Representations (SPARS'09),
 Saint-Malo, France, April 2009.
 
 ( PDF available at  http://www.cs.tut.fi/~foi/GCF-BM3D )

 
--------------------------------------------------------------------
 
This demo package includes routines from both the
LASIP 2D demobox (http://www.cs.tut.fi/~lasip/2D/)  and the
Pointwise SA-DCT demobox (http://www.cs.tut.fi/~foi/SA-DCT/).

--------------------------------------------------------------------


--------------------------------------------------------------------
 Disclaimer
--------------------------------------------------------------------

Any unauthorized use of these routines for industrial or profit-
oriented activities is expressively prohibited. By downloading 
and/or using any of these files, you implicitly agree to all the 
terms of the TUT limited license, as specified in the document
Legal_Notice.txt (included in this package) and online at
http://www.cs.tut.fi/~foi/GCF-BM3D/legal_notice.html
