## Supplementary material for "FPM app: an open-source application for simple and intuitive Fourier ptychographic reconstruction": FPM app: BM3D_CFA.m

```
function [varargout] = BM3D_CFA(z, sigma)
%%%%%%%%%%%%%%%%%%%%%%%%%%%%%%%%%%%%%%%%%%%%%%%%%%%%%%%%%%%%%%%%%%%%%%%%%%%%%
%
%  BM3D_CFA is the modification of the BM3D algorithm for attenuation of additive white Gaussian noise from 
%  Bayer CFA images. This algorithm reproduces the results from the article:
%
%  [1] A. Danielyan, M. Vehvil�inen, A. Foi, V. Katkovnik, and K. Egiazarian,
%       �Cross-color BM3D filtering of noisy raw data�, 
%       Proc. Int. Workshop on Local and Non-Local Approx. in Image Process.,
%       LNLA 2009, Tuusula, Finland, pp. 125-129, August 2009.
%
%  FUNCTION INTERFACE:
%
%  [y_wiener, y_ht] = BM3D(z, sigma)
%
%  ! The function can work without any of the input arguments, 
%   in which case, the internal default ones are used !

%  INPUT ARGUMENTS (OPTIONAL):
%
%     2) z (matrix M x N): Noisy image (intensities in range [0,1] or [0,255])
%     3) sigma (double)  : Std. dev. of the noise (corresponding to intensities
%                          in range [0,255] even if the range of z is [0,1])
%  OUTPUTS:                                             
%     1) y_wiener (matrix M x N): Final(wiener) estimate (in the range [0,1])
%     2) y_ht (matrix M x N): Basic (hard-thresholding) estimate (in the range [0,1])
%
%
%%%%%%%%%%%%%%%%%%%%%%%%%%%%%%%%%%%%%%%%%%%%%%%%%%%%%%%%%%%%%%%%%%%%%%%%%%
%
% Copyright (c) 2009-2014 Tampere University of Technology.
% All rights reserved.
% This work should only be used for nonprofit purposes.
%
% AUTHORS:
%     Aram Danielyan, email: aram dot danielyan _at_ .tut.fi
%
%%%%%%%%%%%%%%%%%%%%%%%%%%%%%%%%%%%%%%%%%%%%%%%%%%%%%%%%%%%%%%%%%%%%%%%%%%


%%%%%%%%%%%%%%%%%%%%%%%%%%%%%%%%%%%%%%%%%%%%%%%%%%%%%%%%%%%%%%%%%%%%%%%%%%
%%%% In case, a noisy image z is not provided, then use the filename 
%%%%  below to read an original image (might contain path also). Later, 
%%%%  artificial AWGN noise is added and this noisy image is processed 
%%%%  by the BM3D.
%%%%
image_name = [
    'kodim07.png'
%     'kodim08.png'
%     'kodim19.png'
%     'kodim23.png'
    ];

%%%%%%%%%%%%%%%%%%%%%%%%%%%%%%%%%%%%%%%%%%%%%%%%%%%%%%%%%%%%%%%%%%%%%%%%%%
%%%%  Quality/complexity trade-off profile selection
%%%%
%%%%  'np' --> Normal Profile (balanced quality)

if ~exist('profile','var')
    profile         = 'np'; %% default profile
end

%%%%%%%%%%%%%%%%%%%%%%%%%%%%%%%%%%%%%%%%%%%%%%%%%%%%%%%%%%%%%%%%%%%%%%%%%%
%%%%  Specify the std. dev. of the corrupting noise
%%%%
if ~exist('sigma','var')
    sigma               = 25; %% default standard deviation of the AWGN
end

%%%%%%%%%%%%%%%%%%%%%%%%%%%%%%%%%%%%%%%%%%%%%%%%%%%%%%%%%%%%%%%%%%%%%%%%%%
%%%% Following are the parameters for the Normal Profile.
%%%%

%%%% Select transforms ('dct', 'dst', 'hadamard', or anything that is listed by 'help wfilters'):
transform_2D_HT_name     = 'dct'; %% transform used for the HT filt. of size N1 x N1
transform_2D_Wiener_name = 'dct';
transform_3rd_dim_name   = 'haar';    %% transform used in the 3-rd dim, the same for HT and Wiener filt.

%%%% Hard-thresholding (HT) parameters:
N1                  = 5;   %% N1 x N1 is the block size used for the hard-thresholding (HT) filtering
Nstep               = 3;   %% sliding step to process every next reference block
N2                  = 16;  %% maximum number of similar blocks (maximum size of the 3rd dimension of a 3D array)
Ns                  = 39;  %% length of the side of the search neighborhood for full-search block-matching (BM), must be odd
lambda_thr2D        = 0;
tau_match           = 3000;%% threshold for the block-distance (d-distance)
lambda_thr3D        = 2.7; %% threshold parameter for the hard-thresholding in 3D transform domain
beta                = 2.0; %% parameter of the 2D Kaiser window used in the reconstruction

%%%% Step 2: Wiener filtering parameters:
N1_wiener           = 6;
Nstep_wiener        = 3;
N2_wiener           = 32;
Ns_wiener           = 39;
tau_match_wiener    = 400;
beta_wiener         = 2.0;


%%%% Block-matching parameters:
stepFS              = 1;  %% step that forces to switch to full-search BM, "1" implies always full-search
smallLN             = 'not used in np'; %% if stepFS > 1, then this specifies the size of the small local search neighb.
stepFSW             = 1;
smallLNW            = 'not used in np';
thrToIncStep        = 8;  % if the number of non-zero coefficients after HT is less than thrToIncStep,
                          % than the sliding step to the next reference block is incresed to (nm1-1)

decLevel = 0;        %% dec. levels of the dyadic wavelet 2D transform for blocks (0 means full decomposition, higher values decrease the dec. number)
thr_mask = ones(N1); %% N1xN1 mask of threshold scaling coeff. --- by default there is no scaling, however the use of different thresholds for different wavelet decompoistion subbands can be done with this matrix

%%%%%%%%%%%%%%%%%%%%%%%%%%%%%%%%%%%%%%%%%%%%%%%%%%%%%%%%%%%%%%%%%%%%%%%%%%
%%%%%%%%%%%%%%%%%%%%%%%%%%%%%%%%%%%%%%%%%%%%%%%%%%%%%%%%%%%%%%%%%%%%%%%%%%
%%%% Note: touch below this point only if you know what you are doing!
%%%%%%%%%%%%%%%%%%%%%%%%%%%%%%%%%%%%%%%%%%%%%%%%%%%%%%%%%%%%%%%%%%%%%%%%%%


%%%%%%%%%%%%%%%%%%%%%%%%%%%%%%%%%%%%%%%%%%%%%%%%%%%%%%%%%%%%%%%%%%%%%%%%%%
%%%% Create transform matrices, etc.
%%%%
[Tfor, Tinv]   = getTransfMatrix(N1, transform_2D_HT_name, decLevel);     %% get (normalized) forward and inverse transform matrices
[TforW, TinvW] = getTransfMatrix(N1_wiener, transform_2D_Wiener_name, 0); %% get (normalized) forward and inverse transform matrices

if (strcmp(transform_3rd_dim_name, 'haar') == 1) | (strcmp(transform_3rd_dim_name(end-2:end), '1.1') == 1),
    %%% If Haar is used in the 3-rd dimension, then a fast internal transform is used, thus no need to generate transform
    %%% matrices.
    hadper_trans_single_den         = {};
    inverse_hadper_trans_single_den = {};
else
    %%% Create transform matrices. The transforms are later applied by
    %%% matrix-vector multiplication for the 1D case.
    for hpow = 0:ceil(log2(max(N2,N2_wiener))),
        h = 2^hpow;
        [Tfor3rd, Tinv3rd]   = getTransfMatrix(h, transform_3rd_dim_name, 0);
        hadper_trans_single_den{h}         = single(Tfor3rd);
        inverse_hadper_trans_single_den{h} = single(Tinv3rd');
    end
end

%%%%%%%%%%%%%%%%%%%%%%%%%%%%%%%%%%%%%%%%%%%%%%%%%%%%%%%%%%%%%%%%%%%%%%%%%%
%%%% 2D Kaiser windows used in the aggregation of block-wise estimates
%%%%
if beta_wiener==2 & beta==2 & N1_wiener==8 & N1==8 % hardcode the window function so that the signal processing toolbox is not needed by default
    Wwin2D = [ 0.1924    0.2989    0.3846    0.4325    0.4325    0.3846    0.2989    0.1924;
        0.2989    0.4642    0.5974    0.6717    0.6717    0.5974    0.4642    0.2989;
        0.3846    0.5974    0.7688    0.8644    0.8644    0.7688    0.5974    0.3846;
        0.4325    0.6717    0.8644    0.9718    0.9718    0.8644    0.6717    0.4325;
        0.4325    0.6717    0.8644    0.9718    0.9718    0.8644    0.6717    0.4325;
        0.3846    0.5974    0.7688    0.8644    0.8644    0.7688    0.5974    0.3846;
        0.2989    0.4642    0.5974    0.6717    0.6717    0.5974    0.4642    0.2989;
        0.1924    0.2989    0.3846    0.4325    0.4325    0.3846    0.2989    0.1924];
    Wwin2D_wiener = Wwin2D;
else
    Wwin2D           = kaiser(N1, beta) * kaiser(N1, beta)'; % Kaiser window used in the aggregation of the HT part
    Wwin2D_wiener    = kaiser(N1_wiener, beta_wiener) * kaiser(N1_wiener, beta_wiener)'; % Kaiser window used in the aggregation of the Wiener filt. part
end
%%%%%%%%%%%%%%%%%%%%%%%%%%%%%%%%%%%%%%%%%%%%%%%%%%%%%%%%%%%%%%%%%%%%%%%%%%
%%%% If needed, read images, generate noise, or scale the images to the 
%%%% [0,1] interval
%%%%
if ~exist('z','var')
    yRGB        = im2double(imread(image_name));  %% read a noise-free image and put in intensity range [0,1]
    y = zeros(size(yRGB,1), size(yRGB,2));
    y(1:2:end,1:2:end) = yRGB(1:2:end,1:2:end,2);
    y(2:2:end,2:2:end) = yRGB(2:2:end,2:2:end,2);
    y(1:2:end,2:2:end) = yRGB(1:2:end,2:2:end,1);
    y(2:2:end,1:2:end) = yRGB(2:2:end,1:2:end,3);
    
    randn('seed', 0);                          %% generate seed
    z        = y + (sigma/255)*randn(size(y)); %% create a noisy image
    
else  % external images
    
    image_name = 'External image';
    
    % convert z to double precision if needed
    z = double(z);
    y= [];
end

if (size(z,3) ~= 1)
    error('BM3D accepts only grayscale 2D images.');
end

%%% Check whether to dump information to the screen or remain silent
if isempty(y)
    dump_output_information = false;
else
    dump_output_information = true;
end

%%%%%%%%%%%%%%%%%%%%%%%%%%%%%%%%%%%%%%%%%%%%%%%%%%%%%%%%%%%%%%%%%%%%%%%%%%
%%% Print image information to the screen
%%%%
if dump_output_information
    fprintf('Image: %s (%dx%d), sigma: %.1f\n', image_name, size(z,1), size(z,2), sigma);
end

%%%%%%%%%%%%%%%%%%%%%%%%%%%%%%%%%%%%%%%%%%%%%%%%%%%%%%%%%%%%%%%%%%%%%%%%%%
%%%% Step 1. Produce the basic estimate by HT filtering
%%%%
tic;
y_ht = bm3d_CFA_thr(z, hadper_trans_single_den, Nstep, N1, N2, lambda_thr2D,...
	lambda_thr3D, tau_match*N1*N1/(255*255), (Ns-1)/2, (sigma/255), thrToIncStep, single(Tfor), single(Tinv)', inverse_hadper_trans_single_den, single(thr_mask), Wwin2D, smallLN, stepFS );
estimate_elapsed_time = toc;

if dump_output_information
    PSNR_INITIAL_ESTIMATE = 10*log10(1/mean((y(:)-double(y_ht(:))).^2));
    fprintf('BASIC ESTIMATE, PSNR: %.2f dB\n', PSNR_INITIAL_ESTIMATE);
end


%%%%%%%%%%%%%%%%%%%%%%%%%%%%%%%%%%%%%%%%%%%%%%%%%%%%%%%%%%%%%%%%%%%%%%%%%%
%%%% Step 2. Produce the final estimate by Wiener filtering (using the 
%%%%  hard-thresholding initial estimate)
%%%%
tic;
y_wiener = bm3d_CFA_wiener(z, y_ht, hadper_trans_single_den, Nstep_wiener, N1_wiener, N2_wiener, ...
    'unused arg', tau_match_wiener*N1_wiener*N1_wiener/(255*255), (Ns_wiener-1)/2, (sigma/255), 'unused arg', single(TforW), single(TinvW)', inverse_hadper_trans_single_den, Wwin2D_wiener, smallLNW, stepFSW, single(ones(N1_wiener)) );
wiener_elapsed_time = toc;

%%%%%%%%%%%%%%%%%%%%%%%%%%%%%%%%%%%%%%%%%%%%%%%%%%%%%%%%%%%%%%%%%%%%%%%%%%
%%%% Calculate the final estimate's PSNR, print it, and show the
%%%% denoised image next to the noisy one
%%%%
y_wiener = double(y_wiener);


if dump_output_information 
    PSNR = 10*log10(1/mean((y(:)-y_wiener(:)).^2)); % y is valid
    fprintf('FINAL ESTIMATE (total time: %.1f sec), PSNR: %.2f dB\n', ...
        wiener_elapsed_time + estimate_elapsed_time, PSNR);

    figure, imshow(z); title(sprintf('Noisy %s, PSNR: %.3f dB (sigma: %d)', ...
        image_name(1:end-4), 10*log10(1/mean((y(:)-z(:)).^2)), sigma));

    figure, imshow(y_wiener); title(sprintf('Denoised %s, PSNR: %.3f dB', ...
        image_name(1:end-4), PSNR));
    
end

if nargout==0
    varargout={};
else
    varargout{1}=y_wiener;
    varargout{2}=y_ht;
end

return;


%%%%%%%%%%%%%%%%%%%%%%%%%%%%%%%%%%%%%%%%%%%%%%%%%%%%%%%%%%%%%%%%%%%%%%%%%%
%%%%%%%%%%%%%%%%%%%%%%%%%%%%%%%%%%%%%%%%%%%%%%%%%%%%%%%%%%%%%%%%%%%%%%%%%%
% Some auxiliary functions 
%%%%%%%%%%%%%%%%%%%%%%%%%%%%%%%%%%%%%%%%%%%%%%%%%%%%%%%%%%%%%%%%%%%%%%%%%%
%%%%%%%%%%%%%%%%%%%%%%%%%%%%%%%%%%%%%%%%%%%%%%%%%%%%%%%%%%%%%%%%%%%%%%%%%%


function [Tforward, Tinverse] = getTransfMatrix (N, transform_type, dec_levels)
%
% Create forward and inverse transform matrices, which allow for perfect
% reconstruction. The forward transform matrix is normalized so that the 
% l2-norm of each basis element is 1.
%
% [Tforward, Tinverse] = getTransfMatrix (N, transform_type, dec_levels)
%
%  INPUTS:
%
%   N               --> Size of the transform (for wavelets, must be 2^K)
%
%   transform_type  --> 'dct', 'dst', 'hadamard', or anything that is 
%                       listed by 'help wfilters' (bi-orthogonal wavelets)
%                       'DCrand' -- an orthonormal transform with a DC and all
%                       the other basis elements of random nature
%
%   dec_levels      --> If a wavelet transform is generated, this is the
%                       desired decomposition level. Must be in the
%                       range [0, log2(N)-1], where "0" implies
%                       full decomposition.
%
%  OUTPUTS:
%
%   Tforward        --> (N x N) Forward transform matrix
%
%   Tinverse        --> (N x N) Inverse transform matrix
%

if exist('dec_levels') ~= 1,
    dec_levels = 0;
end

if N == 1,
    Tforward = 1;
elseif strcmp(transform_type, 'hadamard') == 1,
    Tforward    = hadamard(N);
elseif (N == 8) & strcmp(transform_type, 'bior1.5')==1 % hardcoded transform so that the wavelet toolbox is not needed to generate it
    Tforward =  [ 0.353553390593274   0.353553390593274   0.353553390593274   0.353553390593274   0.353553390593274   0.353553390593274   0.353553390593274   0.353553390593274;
       0.219417649252501   0.449283757993216   0.449283757993216   0.219417649252501  -0.219417649252501  -0.449283757993216  -0.449283757993216  -0.219417649252501;
       0.569359398342846   0.402347308162278  -0.402347308162278  -0.569359398342846  -0.083506045090284   0.083506045090284  -0.083506045090284   0.083506045090284;
      -0.083506045090284   0.083506045090284  -0.083506045090284   0.083506045090284   0.569359398342846   0.402347308162278  -0.402347308162278  -0.569359398342846;
       0.707106781186547  -0.707106781186547                   0                   0                   0                   0                   0                   0;
                       0                   0   0.707106781186547  -0.707106781186547                   0                   0                   0                   0;
                       0                   0                   0                   0   0.707106781186547  -0.707106781186547                   0                   0;
                       0                   0                   0                   0                   0                   0   0.707106781186547  -0.707106781186547];   
elseif (N == 8) & strcmp(transform_type, 'dct')==1 % hardcoded transform so that the signal processing toolbox is not needed to generate it
    Tforward = [ 0.353553390593274   0.353553390593274   0.353553390593274   0.353553390593274   0.353553390593274   0.353553390593274   0.353553390593274   0.353553390593274;
       0.490392640201615   0.415734806151273   0.277785116509801   0.097545161008064  -0.097545161008064  -0.277785116509801  -0.415734806151273  -0.490392640201615;
       0.461939766255643   0.191341716182545  -0.191341716182545  -0.461939766255643  -0.461939766255643  -0.191341716182545   0.191341716182545   0.461939766255643;
       0.415734806151273  -0.097545161008064  -0.490392640201615  -0.277785116509801   0.277785116509801   0.490392640201615   0.097545161008064  -0.415734806151273;
       0.353553390593274  -0.353553390593274  -0.353553390593274   0.353553390593274   0.353553390593274  -0.353553390593274  -0.353553390593274   0.353553390593274;
       0.277785116509801  -0.490392640201615   0.097545161008064   0.415734806151273  -0.415734806151273  -0.097545161008064   0.490392640201615  -0.277785116509801;
       0.191341716182545  -0.461939766255643   0.461939766255643  -0.191341716182545  -0.191341716182545   0.461939766255643  -0.461939766255643   0.191341716182545;
       0.097545161008064  -0.277785116509801   0.415734806151273  -0.490392640201615   0.490392640201615  -0.415734806151273   0.277785116509801  -0.097545161008064];
elseif (N == 8) & strcmp(transform_type, 'dst')==1 % hardcoded transform so that the PDE toolbox is not needed to generate it
    Tforward = [ 0.161229841765317   0.303012985114696   0.408248290463863   0.464242826880013   0.464242826880013   0.408248290463863   0.303012985114696   0.161229841765317;
       0.303012985114696   0.464242826880013   0.408248290463863   0.161229841765317  -0.161229841765317  -0.408248290463863  -0.464242826880013  -0.303012985114696;
       0.408248290463863   0.408248290463863                   0  -0.408248290463863  -0.408248290463863                   0   0.408248290463863   0.408248290463863;
       0.464242826880013   0.161229841765317  -0.408248290463863  -0.303012985114696   0.303012985114696   0.408248290463863  -0.161229841765317  -0.464242826880013;
       0.464242826880013  -0.161229841765317  -0.408248290463863   0.303012985114696   0.303012985114696  -0.408248290463863  -0.161229841765317   0.464242826880013;
       0.408248290463863  -0.408248290463863                   0   0.408248290463863  -0.408248290463863                   0   0.408248290463863  -0.408248290463863;
       0.303012985114696  -0.464242826880013   0.408248290463863  -0.161229841765317  -0.161229841765317   0.408248290463863  -0.464242826880013   0.303012985114696;
       0.161229841765317  -0.303012985114696   0.408248290463863  -0.464242826880013   0.464242826880013  -0.408248290463863   0.303012985114696  -0.161229841765317];
elseif strcmp(transform_type, 'dct') == 1,
    Tforward    = dct(eye(N));
elseif strcmp(transform_type, 'dst') == 1,
    Tforward    = dst(eye(N));
elseif strcmp(transform_type, 'DCrand') == 1,
    x = randn(N); x(1:end,1) = 1; [Q,R] = qr(x); 
    if (Q(1) < 0), 
        Q = -Q; 
    end;
    Tforward = Q';
else %% a wavelet decomposition supported by 'wavedec'
    %%% Set periodic boundary conditions, to preserve bi-orthogonality
    dwtmode('per','nodisp');  
    
    Tforward = zeros(N,N);
    for i = 1:N
        Tforward(:,i)=wavedec(circshift([1 zeros(1,N-1)],[dec_levels i-1]), log2(N), transform_type);  %% construct transform matrix
    end
end

%%% Normalize the basis elements
Tforward = (Tforward' * diag(sqrt(1./sum(Tforward.^2,2))))'; 

%%% Compute the inverse transform matrix
Tinverse = inv(Tforward);

return;
```
