## Supplementary material for "FPM app: an open-source application for simple and intuitive Fourier ptychographic reconstruction": FPM app: BM3DDEB.m

```
function [ISNR, y_hat_RWI] = BM3DDEB(experiment_number, test_image_name)
%%%%%%%%%%%%%%%%%%%%%%%%%%%%%%%%%%%%%%%%%%%%%%%%%%%%%%%%%%%%%%%%%%%%%%%%%%
%
% Copyright (c) 2008-2014 Tampere University of Technology. All rights reserved.
% This work should only be used for nonprofit purposes.
%
% AUTHORS:
%     Kostadin Dabov
%     Alessandro Foi         email: alessandro.foi _at_ tut.fi
%%%%%%%%%%%%%%%%%%%%%%%%%%%%%%%%%%%%%%%%%%%%%%%%%%%%%%%%%%%%%%%%%%%%%%%%%%
%
%  This function implements the image deblurring method proposed in:
%
%  [1] K. Dabov, A. Foi, V. Katkovnik, and K. Egiazarian, "Image
%   restoration  by sparse 3D transform-domain collaborative filtering,"
%   Proc SPIE Electronic Imaging, January 2008.
%
%  FUNCTION INTERFACE:
%
%  [PSNR, y_hat_RWI] = BM3DDEB(experiment_number, test_image_name)
%
%  INPUT:
%   1) experiment_number: 1 -> PSF 1, sigma^2 = 2
%                         2 -> PSF 1, sigma^2 = 8
%                         3 -> PSF 2, sigma^2 = 0.308
%                         4 -> PSF 3, sigma^2 = 49
%                         5 -> PSF 4, sigma^2 = 4
%                         6 -> PSF 5, sigma^2 = 64
%
%   2) test_image_name:   a valid filename of a grayscale test image
%
%  OUTPUT:
%   1) ISNR:              the output improvement in SNR, dB
%   2) y_hat_RWI:         the restored image
%
%  ! The function can work without any of the input arguments,
%   in which case, the internal default ones are used !
%
%%%%%%%%%%%%%%%%%%%%%%%%%%%%%%%%%%%%%%%%%%%%%%%%%%%%%%%%%%%%%%%%%%%%%%%%%%

%%%% Fixed regularization parameters (obtained empirically after a rough optimization)
Regularization_alpha_RI = 4e-4;
Regularization_alpha_RWI = 5e-3;

%%%% Experiment number (see below for details, e.g. how the blur is generated, etc.)
if (exist('experiment_number') ~= 1)
    experiment_number = 3; % 1 -- 6
end

%%%%%%%%%%%%%%%%%%%%%%%%%%%%%%%%%%%%%%%%%%%%%%%%%%%%%%%%%%%%%%%%%%%%%%%%%%
%%%% Select a single image filename (might contain path)
%%%%
if (exist('test_image_name') ~= 1)
    test_image_name = [
        %        'Lena512.png'
        'Cameraman256.png'
        %        'barbara.png'
        %        'house.png'
        ];
end

%%%% Select 2D transforms ('dct', 'dst', 'hadamard', or anything that is listed by 'help wfilters'):
transform_2D_HT_name      = 'dst'; %% 2D transform (of size N1 x N1) used in Step 1
transform_2D_Wiener_name  = 'dct'; %% 2D transform (of size N1_wiener x N1_wiener) used in Step 2
transform_3rd_dimage_name = 'haar'; %% 1D tranform used in the 3-rd dim, the same for both steps

%%%% Step 1 (BM3D with collaborative hard-thresholding) parameters:
N1                  = 8;   %% N1 x N1 is the block size
Nstep               = 3;   %% sliding step to process every next refernece block
N2                  = 16;  %% maximum number of similar blocks (maximum size of the 3rd dimensiona of a 3D array)
Ns                  = 39;  %% length of the side of the search neighborhood for full-search block-matching (BM)
tau_match           = 6000;%% threshold for the block distance (d-distance)
lambda_thr2D        = 0;   %% threshold for the coarse initial denoising used in the d-distance measure
lambda_thr3D        = 2.9; %% threshold for the hard-thresholding
beta                = 0; %% the beta parameter of the 2D Kaiser window used in the reconstruction

%%%% Step 2 (BM3D with collaborative Wiener filtering) parameters:
N1_wiener           = 8;
Nstep_wiener        = 2;
N2_wiener           = 16;
Ns_wiener           = 39;
tau_match_wiener    = 800;
beta_wiener         = 0;

%%%%  Specify whether to print results and display images
print_to_screen     = 1;


%%%%%%%%%%%%%%%%%%%%%%%%%%%%%%%%%%%%%%%%%%%%%%%%%%%%%%%%%%%%%%%%%%%%%%%%%%
%%%%%%%%%%%%%%%%%%%%%%%%%%%%%%%%%%%%%%%%%%%%%%%%%%%%%%%%%%%%%%%%%%%%%%%%%%
%%%% Note: touch below this point only if you know what you are doing!
%%%%%%%%%%%%%%%%%%%%%%%%%%%%%%%%%%%%%%%%%%%%%%%%%%%%%%%%%%%%%%%%%%%%%%%%%%
%%%%%%%%%%%%%%%%%%%%%%%%%%%%%%%%%%%%%%%%%%%%%%%%%%%%%%%%%%%%%%%%%%%%%%%%%%

%%%%%%%%%%%%%%%%%%%%%%%%%%%%%%%%%%%%%%%%%%%%%%%%%%%%%%%%%%%%%%%%%%%%%%%%%%
%%%% Make parameters compatible with the interface of the mex-functions
%%%%

[Tfor, Tinv]   = getTransfMatrix(N1, transform_2D_HT_name, 0); %% get (normalized) forward and inverse transform matrices
[TforW, TinvW] = getTransfMatrix(N1_wiener, transform_2D_Wiener_name, 0); %% get (normalized) forward and inverse transform matrices

if (strcmp(transform_3rd_dimage_name, 'haar') == 1),
    %%% Fast internal transform is used, no need to generate transform
    %%% matrices.
    hadper_trans_single_den         = {};
    inverse_hadper_trans_single_den = {};
else
    %%% Create transform matrices. The transforms are later applied by
    %%% vector-matrix multiplications
    for hpow = 0:ceil(log2(max(N2,N2_wiener))),
        h = 2^hpow;
        [Tfor3rd, Tinv3rd] = getTransfMatrix(h, transform_3rd_dimage_name, 0);
        hadper_trans_single_den{h}         = single(Tfor3rd);
        inverse_hadper_trans_single_den{h} = single(Tinv3rd');
    end
end

if beta == 0 & beta_wiener == 0
    Wwin2D = ones(N1,N1);
    Wwin2D_wiener = ones(N1_wiener,N1_wiener);
else
    Wwin2D        = kaiser(N1, beta) * kaiser(N1, beta)'; % Kaiser window used in the hard-thresholding part
    Wwin2D_wiener = kaiser(N1_wiener, beta_wiener) * kaiser(N1_wiener, beta_wiener)'; % Kaiser window used in the Wiener filtering part
end

%%%%%%%%%%%%%%%%%%%%%%%%%%%%%%%%%%%%%%%%%%%%%%%%%%%%%%%%%%%%%%%%%%%%%%%%%%
%%%% Read an image and generate a blurred and noisy image
%%%%
y = im2double(imread(test_image_name));

if experiment_number==1
    sigma=sqrt(2)/255;
    for x1=-7:7; for x2=-7:7; v(x1+8,x2+8)=1/(x1^2+x2^2+1); end, end; v=v./sum(v(:));
end
if experiment_number==2
    sigma=sqrt(8)/255;
    s1=0; for a1=-7:7; s1=s1+1; s2=0; for a2=-7:7; s2=s2+1; v(s1,s2)=1/(a1^2+a2^2+1); end, end;  v=v./sum(v(:));
end
if experiment_number==3
    BSNR=40; sigma=-1; % if "sigma=-1", then the value of sigma depends on the BSNR
    v=ones(9); v=v./sum(v(:));
end
if experiment_number==4
    sigma=7/255;
    v=[1 4 6 4 1]'*[1 4 6 4 1]; v=v./sum(v(:));  % PSF
end
if experiment_number==5
    sigma=2/255;
    v=fspecial('gaussian', 25, 1.6);
end
if experiment_number==6
    sigma=8/255;
    v=fspecial('gaussian', 25, .4);
end


[Xv, Xh]  = size(y);
[ghy,ghx] = size(v);
big_v  = zeros(Xv,Xh); big_v(1:ghy,1:ghx)=v; big_v=circshift(big_v,-round([(ghy-1)/2 (ghx-1)/2])); % pad PSF with zeros to whole image domain, and center it
V      = fft2(big_v); % frequency response of the PSF
y_blur = imfilter(y, v(end:-1:1,end:-1:1), 'circular'); % performs blurring (by circular convolution)

randn('seed',0);  %%% fix seed for the random number generator
if sigma == -1;   %% check whether to use BSNR in order to define value of sigma
    sigma=sqrt(norm(y_blur(:)-mean(y_blur(:)),2)^2 /(Xh*Xv*10^(BSNR/10))); % compute sigma from the desired BSNR
end

%%%% Create a blurred and noisy observation
z = y_blur + sigma*randn(Xv,Xh);

tic;
%%%%%%%%%%%%%%%%%%%%%%%%%%%%%%%%%%%%%%%%%%%%%%%%%%%%%%%%%%%%%%%%%%%%%%%%%%
%%%% Step 1: Final estimate by Regularized Inversion (RI) followed by
%%%% BM3D with collaborative hard-thresholding
%%%%

%%%% Step 1.1. Regularized Inversion
RI= conj(V)./( (abs(V).^2) + Regularization_alpha_RI * Xv*Xh*sigma^2); % Transfer Matrix for RI    %% Standard Tikhonov Regularization
zRI=real(ifft2( fft2(z).* RI ));   % Regularized Inverse Estimate (RI OBSERVATION)

stdRI = zeros(N1, N1);
for ii = 1:N1,
    for jj = 1:N1,
        UnitMatrix = zeros(N1,N1); UnitMatrix(ii,jj)=1;
        BasisElementPadded = zeros(Xv, Xh); BasisElementPadded(1:N1,1:N1) = Tinv*UnitMatrix*Tinv';
        TransfBasisElementPadded = fft2(BasisElementPadded);
        stdRI(ii,jj) = sqrt( (1/(Xv*Xh)) * sum(sum(abs(TransfBasisElementPadded.*RI).^2)) )*sigma;
    end,
end

%%%% Step 1.2. Colored noise suppression by BM3D with collaborative hard-
%%%% thresholding

y_hat_RI = bm3d_thr_colored_noise(zRI, hadper_trans_single_den, Nstep, N1, N2, lambda_thr2D,...
    lambda_thr3D, tau_match*N1*N1/(255*255), (Ns-1)/2, sigma, 0, single(Tfor), single(Tinv)',...
    inverse_hadper_trans_single_den, single(stdRI'), Wwin2D, 0, 1 );

PSNR_INITIAL_ESTIMATE = 10*log10(1/mean((y(:)-y_hat_RI(:)).^2));
ISNR_INITIAL_ESTIMATE = PSNR_INITIAL_ESTIMATE - 10*log10(1/mean((y(:)-z(:)).^2));

%%%%%%%%%%%%%%%%%%%%%%%%%%%%%%%%%%%%%%%%%%%%%%%%%%%%%%%%%%%%%%%%%%%%%%%%%%
%%%% Step 2: Final estimate by Regularized Wiener Inversion (RWI) followed
%%%% by BM3D with collaborative Wiener filtering
%%%%

%%%% Step 2.1. Regularized Wiener Inversion
Wiener_Pilot = abs(fft2(double(y_hat_RI)));   %%% Wiener reference estimate
RWI  = conj(V).*Wiener_Pilot.^2./(Wiener_Pilot.^2.*(abs(V).^2) + Regularization_alpha_RWI*Xv*Xh*sigma^2);   % Transfer Matrix for RWI (uses standard regularization 'a-la-Tikhonov')
zRWI = real(ifft2(fft2(z).*RWI));   % RWI OBSERVATION

stdRWI = zeros(N1_wiener, N1_wiener);
for ii = 1:N1_wiener,
    for jj = 1:N1_wiener,
        UnitMatrix = zeros(N1_wiener,N1_wiener); UnitMatrix(ii,jj)=1;
        BasisElementPadded = zeros(Xv, Xh); BasisElementPadded(1:N1_wiener,1:N1_wiener) = TinvW*UnitMatrix*TinvW';
        TransfBasisElementPadded = fft2(BasisElementPadded);
        stdRWI(ii,jj) = sqrt( (1/(Xv*Xh)) * sum(sum(abs(TransfBasisElementPadded.*RWI).^2)) )*sigma;
    end,
end

%%%% Step 2.2. Colored noise suppression by BM3D with collaborative Wiener
%%%% filtering
y_hat_RWI = bm3d_wiener_colored_noise(zRWI, y_hat_RI, hadper_trans_single_den, Nstep_wiener, N1_wiener, N2_wiener, ...
    0, tau_match_wiener*N1_wiener*N1_wiener/(255*255), (Ns_wiener-1)/2, 0, single(stdRWI'), single(TforW), single(TinvW)',...
    inverse_hadper_trans_single_den, Wwin2D_wiener, 0, 1, single(ones(N1_wiener)) );

elapsed_time = toc;


%%%%%%%%%%%%%%%%%%%%%%%%%%%%%%%%%%%%%%%%%%%%%%%%%%%%%%%%%%%%%%%%%%%%%%%%%%
%%%% Calculate the final estimate's PSNR and ISNR, print them, and show the
%%%% restored image
%%%%
PSNR = 10*log10(1/mean((y(:)-y_hat_RWI(:)).^2));
ISNR = PSNR - 10*log10(1/mean((y(:)-z(:)).^2));

if print_to_screen == 1
    fprintf('Image: %s, Exp %d, Time: %.1f sec, PSNR-RI: %.2f dB, PSNR-RWI: %.2f, ISNR-RWI: %.2f dB\n', ...
        test_image_name, experiment_number, elapsed_time, PSNR_INITIAL_ESTIMATE, PSNR, ISNR);
    figure,imshow(z);
    figure,imshow(double(y_hat_RWI));
end

return;


%%%%%%%%%%%%%%%%%%%%%%%%%%%%%%%%%%%%%%%%%%%%%%%%%%%%%%%%%%%%%%%%%%%%%%%%%%
%%%%%%%%%%%%%%%%%%%%%%%%%%%%%%%%%%%%%%%%%%%%%%%%%%%%%%%%%%%%%%%%%%%%%%%%%%
% Some auxiliary functions
%%%%%%%%%%%%%%%%%%%%%%%%%%%%%%%%%%%%%%%%%%%%%%%%%%%%%%%%%%%%%%%%%%%%%%%%%%
%%%%%%%%%%%%%%%%%%%%%%%%%%%%%%%%%%%%%%%%%%%%%%%%%%%%%%%%%%%%%%%%%%%%%%%%%%


function [Tforward, Tinverse] = getTransfMatrix (N, transform_type, dec_levels)
%
% Create forward and inverse transform matrices, which allow for perfect
% reconstruction. The forward transform matrix is normalized so that the
% l2-norm of each basis element is 1.
%
% [Tforward, Tinverse] = getTransfMatrix (N, transform_type, dec_levels)
%
%  INPUTS:
%
%   N               --> Size of the transform (for wavelets, must be 2^K)
%
%   transform_type  --> 'dct', 'dst', 'hadamard', or anything that is
%                       listed by 'help wfilters' (bi-orthogonal wavelets)
%                       'DCrand' -- an orthonormal transform with a DC and all
%                       the other basis elements of random nature
%
%   dec_levels      --> If a wavelet transform is generated, this is the
%                       desired decomposition level. Must be in the
%                       range [0, log2(N)-1], where "0" implies
%                       full decomposition.
%
%  OUTPUTS:
%
%   Tforward        --> (N x N) Forward transform matrix
%
%   Tinverse        --> (N x N) Inverse transform matrix
%

if exist('dec_levels') ~= 1,
    dec_levels = 0;
end

if N == 1,
    Tforward = 1;
elseif strcmp(transform_type, 'hadamard') == 1,
    Tforward    = hadamard(N);
elseif (N == 8) & strcmp(transform_type, 'bior1.5')==1 % hardcoded transform so that the wavelet toolbox is not needed to generate it
    Tforward =  [ 0.353553390593274   0.353553390593274   0.353553390593274   0.353553390593274   0.353553390593274   0.353553390593274   0.353553390593274   0.353553390593274;
        0.219417649252501   0.449283757993216   0.449283757993216   0.219417649252501  -0.219417649252501  -0.449283757993216  -0.449283757993216  -0.219417649252501;
        0.569359398342846   0.402347308162278  -0.402347308162278  -0.569359398342846  -0.083506045090284   0.083506045090284  -0.083506045090284   0.083506045090284;
        -0.083506045090284   0.083506045090284  -0.083506045090284   0.083506045090284   0.569359398342846   0.402347308162278  -0.402347308162278  -0.569359398342846;
        0.707106781186547  -0.707106781186547                   0                   0                   0                   0                   0                   0;
        0                   0   0.707106781186547  -0.707106781186547                   0                   0                   0                   0;
        0                   0                   0                   0   0.707106781186547  -0.707106781186547                   0                   0;
        0                   0                   0                   0                   0                   0   0.707106781186547  -0.707106781186547];
elseif (N == 8) & strcmp(transform_type, 'dct')==1 % hardcoded transform so that the signal processing toolbox is not needed to generate it
    Tforward = [ 0.353553390593274   0.353553390593274   0.353553390593274   0.353553390593274   0.353553390593274   0.353553390593274   0.353553390593274   0.353553390593274;
        0.490392640201615   0.415734806151273   0.277785116509801   0.097545161008064  -0.097545161008064  -0.277785116509801  -0.415734806151273  -0.490392640201615;
        0.461939766255643   0.191341716182545  -0.191341716182545  -0.461939766255643  -0.461939766255643  -0.191341716182545   0.191341716182545   0.461939766255643;
        0.415734806151273  -0.097545161008064  -0.490392640201615  -0.277785116509801   0.277785116509801   0.490392640201615   0.097545161008064  -0.415734806151273;
        0.353553390593274  -0.353553390593274  -0.353553390593274   0.353553390593274   0.353553390593274  -0.353553390593274  -0.353553390593274   0.353553390593274;
        0.277785116509801  -0.490392640201615   0.097545161008064   0.415734806151273  -0.415734806151273  -0.097545161008064   0.490392640201615  -0.277785116509801;
        0.191341716182545  -0.461939766255643   0.461939766255643  -0.191341716182545  -0.191341716182545   0.461939766255643  -0.461939766255643   0.191341716182545;
        0.097545161008064  -0.277785116509801   0.415734806151273  -0.490392640201615   0.490392640201615  -0.415734806151273   0.277785116509801  -0.097545161008064];
elseif (N == 8) & strcmp(transform_type, 'dst')==1 % hardcoded transform so that the PDE toolbox is not needed to generate it
    Tforward = [ 0.161229841765317   0.303012985114696   0.408248290463863   0.464242826880013   0.464242826880013   0.408248290463863   0.303012985114696   0.161229841765317;
        0.303012985114696   0.464242826880013   0.408248290463863   0.161229841765317  -0.161229841765317  -0.408248290463863  -0.464242826880013  -0.303012985114696;
        0.408248290463863   0.408248290463863                   0  -0.408248290463863  -0.408248290463863                   0   0.408248290463863   0.408248290463863;
        0.464242826880013   0.161229841765317  -0.408248290463863  -0.303012985114696   0.303012985114696   0.408248290463863  -0.161229841765317  -0.464242826880013;
        0.464242826880013  -0.161229841765317  -0.408248290463863   0.303012985114696   0.303012985114696  -0.408248290463863  -0.161229841765317   0.464242826880013;
        0.408248290463863  -0.408248290463863                   0   0.408248290463863  -0.408248290463863                   0   0.408248290463863  -0.408248290463863;
        0.303012985114696  -0.464242826880013   0.408248290463863  -0.161229841765317  -0.161229841765317   0.408248290463863  -0.464242826880013   0.303012985114696;
        0.161229841765317  -0.303012985114696   0.408248290463863  -0.464242826880013   0.464242826880013  -0.408248290463863   0.303012985114696  -0.161229841765317];
elseif strcmp(transform_type, 'dct') == 1,
    Tforward    = dct(eye(N));
elseif strcmp(transform_type, 'dst') == 1,
    Tforward    = dst(eye(N));
elseif strcmp(transform_type, 'DCrand') == 1,
    x = randn(N); x(1:end,1) = 1; [Q,R] = qr(x);
    if (Q(1) < 0),
        Q = -Q;
    end;
    Tforward = Q';
else %% a wavelet decomposition supported by 'wavedec'
    %%% Set periodic boundary conditions, to preserve bi-orthogonality
    dwtmode('per','nodisp');
    
    Tforward = zeros(N,N);
    for i = 1:N
        Tforward(:,i)=wavedec(circshift([1 zeros(1,N-1)],[dec_levels i-1]), log2(N), transform_type);  %% construct transform matrix
    end
end

%%% Normalize the basis elements
Tforward = (Tforward' * diag(sqrt(1./sum(Tforward.^2,2))))';

%%% Compute the inverse transform matrix
Tinverse = inv(Tforward);

return;
```
