## Supplementary material for "FPM app: an open-source application for simple and intuitive Fourier ptychographic reconstruction": FPM app: BM3DSHARP.m

```
function [y_hat] = BM3DSHARP(z, sigma, alpha_sharp, profile, print_to_screen)
%
%  Joint sharpening and denoising with BM3D. This is implementation of the 
%  BM3D-SH3D sharpening method that is developed in:
%
%  [1] K. Dabov, A. Foi, V. Katkovnik, and K. Egiazarian, "Joint image
%   sharpening and denoising by 3D transform-domain collaborative filtering," 
%   Proc. 2007 Int. TICSP Workshop Spectral Meth. Multirate Signal Process.,
%   SMMSP 2007, Moscow, Russia, September 2007.
%
%  FUNCTION INTERFACE:
%
%  [ysharp] = BM3DSHARP(z, sigma, alpha_sharp, profile, print_to_screen)
%
%  The function can work without any of the input arguments, hence they are
%  optional!
%
%  INPUTS (OPTIONAL):
%
%        1) z (matrix, size MxN)       : Input image (noisy and with poor contrast)
%        2) sigma (double)             : Noise (IF ANY noise) standard deviation (signal assumed
%                                          in the range [0, 255])
%        3) alpha_sharp (double)       : Sharpening parameter (default: 1.5):
%                                         (1,inf) -> sharpen
%                                          1      -> no sharpening
%                                         (0,1)   -> de-sharpen
%        4) profile (char vector)      : 'lc' --> fast 
%                                        'np' --> normal (default)
%        5) print_to_screen (boolean)  : 0 --> do not print output
%                                          information (and do not plot figures)
%                                        1 --> print figures (default)
%
%   OUTPUTS:
%        1) ysharp (matrix, size MxN)  : Sharpened image (in the range [0,1])
%
%  BASIC USAGE EXAMPLES:     
%     
%     sigma = 10;
%     z = im2double(imread('cameraman.tif'));
%     z = z + (sigma/255)*randn(size(z));
%     alpha_sharp = 1.3;
%     [ysharp] = BM3DSHARP(z, sigma, alpha_sharp);
%
%%%%%%%%%%%%%%%%%%%%%%%%%%%%%%%%%%%%%%%%%%%%%%%%%%%%%%%%%%%%%%%%%%%%%%%%%%
%
% Copyright � 2007 Tampere University of Technology. All rights reserved.
% This work should only be used for nonprofit purposes.
%
% AUTHORS:
%     Kostadin Dabov (2007), email: kostadin.dabov _at_ tut.fi
%
%%%%%%%%%%%%%%%%%%%%%%%%%%%%%%%%%%%%%%%%%%%%%%%%%%%%%%%%%%%%%%%%%%%%%%%%%%


%%%%%%%%%%%%%%%%%%%%%%%%%%%%%%%%%%%%%%%%%%%%%%%%%%%%%%%%%%%%%%%%%%%%%%%%%%
%%%% In case, an input image z is not provided, then use the filename 
%%%%  below to read an original image (might contain path also). Later, 
%%%%  artificial AWGN noise is added and this noisy image is processed 
%%%%  by the BM3D-SH3D.
%%%%
if (exist('image_name') ~= 1)
    image_name = [
    % 
    %%%% Grayscale images
    %     'barco.png'
    %     'pentagon.tif'
         'Cameraman256.png'   
    %     'boat.png'
    %     'Lena512.png'
    %     'house.png'
    %     'barbara.png'
];
end
%%%%%%%%%%%%%%%%%%%%%%%%%%%%%%%%%%%%%%%%%%%%%%%%%%%%%%%%%%%%%%%%%%%%%%%%%%
%%%%  Quality/complexity trade-off profile selection
%%%%
%%%%  'np' --> Normal Profile (balanced quality)
%%%%  'lc' --> Low Complexity Profile (fast, lower quality)
%%%%
%%%%  'high' --> High Profile (high quality, not documented in [1])
%%%%
if (exist('profile') ~= 1)
    profile         = 'np'; %% default profile
end

%%%%%%%%%%%%%%%%%%%%%%%%%%%%%%%%%%%%%%%%%%%%%%%%%%%%%%%%%%%%%%%%%%%%%%%%%%
%%%%  Specify the std. dev. of the corrupting noise
%%%%
if (exist('sigma') ~= 1),
    if (exist('z') ~= 1)
        sigma = 20; %% default standard deviation of the AWGN
    else
        fprintf('Please specify value for the s.t.d. "sigma"\n');
        y_hat = 0;
        return;
    end
end
    
if (exist('alpha_sharp') ~= 1)
    alpha_sharp = 3/2;
end

%%%%%%%%%%%%%%%%%%%%%%%%%%%%%%%%%%%%%%%%%%%%%%%%%%%%%%%%%%%%%%%%%%%%%%%%%%
%%%% Following are the parameters for the Normal Profile.
%%%%

%%%% Select transforms ('dct', 'dst', 'hadamard', or anything that is listed by 'help wfilters'):
transform_2D_HT_name     = 'bior1.5'; %% transform used for the HT filt. of size N1 x N1
transform_3rd_dim_name   = 'haar';    %% transform used in the 3-rd dim, the same for HT and Wiener filt.

%%%% Hard-thresholding (HT) parameters:
N1                  = 8;   %% N1 x N1 is the block size used for the hard-thresholding (HT) filtering
Nstep               = 3;   %% sliding step to process every next reference block
N2                  = 16;  %% maximum number of similar blocks (maximum size of the 3rd dimension of a 3D array)
Ns                  = 39;  %% length of the side of the search neighborhood for full-search block-matching (BM), must be odd
tau_match           = 3000;%% threshold for the block-distance (d-distance)
lambda_thr2D        = 0;   %% threshold parameter for the coarse initial denoising used in the d-distance measure
lambda_thr3D        = 2.7; %% threshold parameter for the hard-thresholding in 3D transform domain
beta                = 2.0; %% parameter of the 2D Kaiser window used in the reconstruction

%%%% Block-matching parameters:
stepFS              = 1;  %% step that forces to switch to full-search BM, "1" implies always full-search
smallLN             = 'not used in np'; %% if stepFS > 1, then this specifies the size of the small local search neighb.
thrToIncStep        = 8;  %% used in the HT filtering to increase the sliding step in uniform regions

if strcmp(profile, 'lc') == 1,

    Nstep               = 6;
    Ns                  = 25;

    thrToIncStep        = 3;
    smallLN             = 3;
    stepFS              = 6*Nstep;

end

if (strcmp(profile, 'vn') == 1) | (sigma > 40),

    transform_2D_HT_name = 'dct'; 
    
    N1                  = 12;
    Nstep               = 4;
 
    lambda_thr3D        = 2.8;
    lambda_thr2D        = 2.0;
    thrToIncStep        = 3;
    tau_match           = 5000;
    
end

decLevel = 0;        %% dec. levels of the dyadic wavelet 2D transform for blocks (0 means full decomposition, higher values decrease the dec. number)
thr_mask = ones(N1); %% N1xN1 mask of threshold scaling coeff. --- by default there is no scaling, however the use of different thresholds for different wavelet decompoistion subbands can be done with this matrix

if strcmp(profile, 'high') == 1, %% this profile is not documented in [1]
    
    decLevel     = 1; 
    Nstep        = 2;
    lambda_thr3D = 2.5;
    vMask = ones(N1,1); vMask((end/4+1):end/2)= 1.01; vMask((end/2+1):end) = 1.07; %% this allows to have different threhsolds for the finest and next-to-the-finest subbands
    thr_mask = vMask * vMask'; 
    beta         = 2.5;
    beta_wiener  = 1.5;
    
end

%%%%%%%%%%%%%%%%%%%%%%%%%%%%%%%%%%%%%%%%%%%%%%%%%%%%%%%%%%%%%%%%%%%%%%%%%%
%%%%%%%%%%%%%%%%%%%%%%%%%%%%%%%%%%%%%%%%%%%%%%%%%%%%%%%%%%%%%%%%%%%%%%%%%%
%%%% Note: touch below this point only if you know what you are doing!
%%%%%%%%%%%%%%%%%%%%%%%%%%%%%%%%%%%%%%%%%%%%%%%%%%%%%%%%%%%%%%%%%%%%%%%%%%

%%% Check whether to dump information to the screen or remain silent
dump_output_information = 1;
if (exist('print_to_screen') == 1) & (print_to_screen == 0),
    dump_output_information = 0;
end

%%%%%%%%%%%%%%%%%%%%%%%%%%%%%%%%%%%%%%%%%%%%%%%%%%%%%%%%%%%%%%%%%%%%%%%%%%
%%%% Create transform matrices, etc.
%%%%
[Tfor, Tinv]   = getTransfMatrix(N1, transform_2D_HT_name, decLevel);     %% get (normalized) forward and inverse transform matrices

if (strcmp(transform_3rd_dim_name, 'haar') == 1) | (strcmp(transform_3rd_dim_name(end-2:end), '1.1') == 1),
    %%% If Haar is used in the 3-rd dimension, then a fast internal transform is used, thus no need to generate transform
    %%% matrices.
    hadper_trans_single_den         = {};
    inverse_hadper_trans_single_den = {};
else
    %%% Create transform matrices. The transforms are later applied by
    %%% matrix-vector multiplication for the 1D case.
    for hpow = 0:ceil(log2(max(N2,N2_wiener))),
        h = 2^hpow;
        [Tfor3rd, Tinv3rd]   = getTransfMatrix(h, transform_3rd_dim_name, 0);
        hadper_trans_single_den{h}         = single(Tfor3rd);
        inverse_hadper_trans_single_den{h} = single(Tinv3rd');
    end
end

%%%%%%%%%%%%%%%%%%%%%%%%%%%%%%%%%%%%%%%%%%%%%%%%%%%%%%%%%%%%%%%%%%%%%%%%%%
%%%% 2D Kaiser windows used in the aggregation of block-wise estimates
%%%%
if beta==2 & N1==8 % hardcode the window function so that the signal processing toolbox is not needed by default
    Wwin2D = [ 0.1924    0.2989    0.3846    0.4325    0.4325    0.3846    0.2989    0.1924;
        0.2989    0.4642    0.5974    0.6717    0.6717    0.5974    0.4642    0.2989;
        0.3846    0.5974    0.7688    0.8644    0.8644    0.7688    0.5974    0.3846;
        0.4325    0.6717    0.8644    0.9718    0.9718    0.8644    0.6717    0.4325;
        0.4325    0.6717    0.8644    0.9718    0.9718    0.8644    0.6717    0.4325;
        0.3846    0.5974    0.7688    0.8644    0.8644    0.7688    0.5974    0.3846;
        0.2989    0.4642    0.5974    0.6717    0.6717    0.5974    0.4642    0.2989;
        0.1924    0.2989    0.3846    0.4325    0.4325    0.3846    0.2989    0.1924];
else
    Wwin2D = kaiser(N1, beta) * kaiser(N1, beta)'; % Kaiser window used in the aggregation of the HT part
end
%%%%%%%%%%%%%%%%%%%%%%%%%%%%%%%%%%%%%%%%%%%%%%%%%%%%%%%%%%%%%%%%%%%%%%%%%%
%%%% If needed, read images, generate noise, or scale the images to the 
%%%% [0,1] interval
%%%%
if (exist('z') ~= 1)
    y        = im2double(imread(image_name));  %% read a noise-free image and put in intensity range [0,1]
    randn('seed', 0);                          %% generate seed
    z        = y + (sigma/255)*randn(size(y)); %% create a noisy image
else  % external images
    
    image_name = 'External image';
    
    % convert z to double precision if needed
    z = double(z);
    
    % if z's range is [0, 255], then convert to [0, 1]
    if (max(z(:)) > 10), % a naive check for intensity range
        z = z / 255;
    end
    
end

if (size(z,3) ~= 1),
    error('BM3D-SH3D accepts only grayscale 2D images.');
end

%%%%%%%%%%%%%%%%%%%%%%%%%%%%%%%%%%%%%%%%%%%%%%%%%%%%%%%%%%%%%%%%%%%%%%%%%%
%%% Print image information to the screen
%%%%
if dump_output_information == 1,
    fprintf('Image: %s (%dx%d), sigma: %.1f\n', image_name, size(z,1), size(z,2), sigma);
end

%%%%%%%%%%%%%%%%%%%%%%%%%%%%%%%%%%%%%%%%%%%%%%%%%%%%%%%%%%%%%%%%%%%%%%%%%%
%%%% Apply the filtering MEX-subroutine
%%%%
tic;
y_hat = bm3d_thr_sharpen_var(z, hadper_trans_single_den, Nstep, N1, N2, lambda_thr2D,...
         lambda_thr3D, tau_match*N1*N1/(255*255), (Ns-1)/2, (sigma/255), thrToIncStep, single(Tfor), single(Tinv)', inverse_hadper_trans_single_den, single(thr_mask), Wwin2D, smallLN, stepFS, 1/alpha_sharp );
estimate_elapsed_time = toc;

if dump_output_information == 1,
    fprintf('SHARPENING COMPLETED (total time: %.1f sec)\n', ...
        estimate_elapsed_time);
    imshow(z); figure, imshow(double(y_hat));
end

return;


%%%%%%%%%%%%%%%%%%%%%%%%%%%%%%%%%%%%%%%%%%%%%%%%%%%%%%%%%%%%%%%%%%%%%%%%%%
%%%%%%%%%%%%%%%%%%%%%%%%%%%%%%%%%%%%%%%%%%%%%%%%%%%%%%%%%%%%%%%%%%%%%%%%%%
% Some auxiliary functions 
%%%%%%%%%%%%%%%%%%%%%%%%%%%%%%%%%%%%%%%%%%%%%%%%%%%%%%%%%%%%%%%%%%%%%%%%%%
%%%%%%%%%%%%%%%%%%%%%%%%%%%%%%%%%%%%%%%%%%%%%%%%%%%%%%%%%%%%%%%%%%%%%%%%%%


function [Tforward, Tinverse] = getTransfMatrix (N, transform_type, dec_levels)
%
% Create forward and inverse transform matrices, which allow for perfect
% reconstruction. The forward transform matrix is normalized so that the 
% l2-norm of each basis element is 1.
%
% [Tforward, Tinverse] = getTransfMatrix (N, transform_type, dec_levels)
%
%  INPUTS:
%
%   N               --> Size of the transform (for wavelets, must be 2^K)
%
%   transform_type  --> 'dct', 'dst', 'hadamard', or anything that is 
%                       listed by 'help wfilters' (bi-orthogonal wavelets)
%                       'DCrand' -- an orthonormal transform with a DC and all
%                       the other basis elements of random nature
%
%   dec_levels      --> If a wavelet transform is generated, this is the
%                       desired decomposition level. Must be in the
%                       range [0, log2(N)-1], where "0" implies
%                       full decomposition.
%
%  OUTPUTS:
%
%   Tforward        --> (N x N) Forward transform matrix
%
%   Tinverse        --> (N x N) Inverse transform matrix
%

if exist('dec_levels') ~= 1,
    dec_levels = 0;
end

if N == 1,
    Tforward = 1;
elseif strcmp(transform_type, 'hadamard') == 1,
    Tforward    = hadamard(N);
elseif (N == 8) & strcmp(transform_type, 'bior1.5')==1 % hardcoded transform so that the wavelet toolbox is not needed to generate it
    Tforward =  [ 0.353553390593274   0.353553390593274   0.353553390593274   0.353553390593274   0.353553390593274   0.353553390593274   0.353553390593274   0.353553390593274;
       0.219417649252501   0.449283757993216   0.449283757993216   0.219417649252501  -0.219417649252501  -0.449283757993216  -0.449283757993216  -0.219417649252501;
       0.569359398342846   0.402347308162278  -0.402347308162278  -0.569359398342846  -0.083506045090284   0.083506045090284  -0.083506045090284   0.083506045090284;
      -0.083506045090284   0.083506045090284  -0.083506045090284   0.083506045090284   0.569359398342846   0.402347308162278  -0.402347308162278  -0.569359398342846;
       0.707106781186547  -0.707106781186547                   0                   0                   0                   0                   0                   0;
                       0                   0   0.707106781186547  -0.707106781186547                   0                   0                   0                   0;
                       0                   0                   0                   0   0.707106781186547  -0.707106781186547                   0                   0;
                       0                   0                   0                   0                   0                   0   0.707106781186547  -0.707106781186547];   
elseif (N == 8) & strcmp(transform_type, 'dct')==1 % hardcoded transform so that the signal processing toolbox is not needed to generate it
    Tforward = [ 0.353553390593274   0.353553390593274   0.353553390593274   0.353553390593274   0.353553390593274   0.353553390593274   0.353553390593274   0.353553390593274;
       0.490392640201615   0.415734806151273   0.277785116509801   0.097545161008064  -0.097545161008064  -0.277785116509801  -0.415734806151273  -0.490392640201615;
       0.461939766255643   0.191341716182545  -0.191341716182545  -0.461939766255643  -0.461939766255643  -0.191341716182545   0.191341716182545   0.461939766255643;
       0.415734806151273  -0.097545161008064  -0.490392640201615  -0.277785116509801   0.277785116509801   0.490392640201615   0.097545161008064  -0.415734806151273;
       0.353553390593274  -0.353553390593274  -0.353553390593274   0.353553390593274   0.353553390593274  -0.353553390593274  -0.353553390593274   0.353553390593274;
       0.277785116509801  -0.490392640201615   0.097545161008064   0.415734806151273  -0.415734806151273  -0.097545161008064   0.490392640201615  -0.277785116509801;
       0.191341716182545  -0.461939766255643   0.461939766255643  -0.191341716182545  -0.191341716182545   0.461939766255643  -0.461939766255643   0.191341716182545;
       0.097545161008064  -0.277785116509801   0.415734806151273  -0.490392640201615   0.490392640201615  -0.415734806151273   0.277785116509801  -0.097545161008064];
elseif (N == 8) & strcmp(transform_type, 'dst')==1 % hardcoded transform so that the PDE toolbox is not needed to generate it
    Tforward = [ 0.161229841765317   0.303012985114696   0.408248290463863   0.464242826880013   0.464242826880013   0.408248290463863   0.303012985114696   0.161229841765317;
       0.303012985114696   0.464242826880013   0.408248290463863   0.161229841765317  -0.161229841765317  -0.408248290463863  -0.464242826880013  -0.303012985114696;
       0.408248290463863   0.408248290463863                   0  -0.408248290463863  -0.408248290463863                   0   0.408248290463863   0.408248290463863;
       0.464242826880013   0.161229841765317  -0.408248290463863  -0.303012985114696   0.303012985114696   0.408248290463863  -0.161229841765317  -0.464242826880013;
       0.464242826880013  -0.161229841765317  -0.408248290463863   0.303012985114696   0.303012985114696  -0.408248290463863  -0.161229841765317   0.464242826880013;
       0.408248290463863  -0.408248290463863                   0   0.408248290463863  -0.408248290463863                   0   0.408248290463863  -0.408248290463863;
       0.303012985114696  -0.464242826880013   0.408248290463863  -0.161229841765317  -0.161229841765317   0.408248290463863  -0.464242826880013   0.303012985114696;
       0.161229841765317  -0.303012985114696   0.408248290463863  -0.464242826880013   0.464242826880013  -0.408248290463863   0.303012985114696  -0.161229841765317];
elseif strcmp(transform_type, 'dct') == 1,
    Tforward    = dct(eye(N));
elseif strcmp(transform_type, 'dst') == 1,
    Tforward    = dst(eye(N));
elseif strcmp(transform_type, 'DCrand') == 1,
    x = randn(N); x(1:end,1) = 1; [Q,R] = qr(x); 
    if (Q(1) < 0), 
        Q = -Q; 
    end;
    Tforward = Q';
else %% a wavelet decomposition supported by 'wavedec'
    %%% Set periodic boundary conditions, to preserve bi-orthogonality
    dwtmode('per','nodisp');  
    
    Tforward = zeros(N,N);
    for i = 1:N
        Tforward(:,i)=wavedec(circshift([1 zeros(1,N-1)],[dec_levels i-1]), log2(N), transform_type);  %% construct transform matrix
    end
end

%%% Normalize the basis elements
Tforward = (Tforward' * diag(sqrt(1./sum(Tforward.^2,2))))'; 

%%% Compute the inverse transform matrix
Tinverse = inv(Tforward);

return;
```
