## Supplementary material for "FPM app: an open-source application for simple and intuitive Fourier ptychographic reconstruction": FPM app: CBM3D.m

```
function [PSNR, yRGB_est] = CBM3D(yRGB, zRGB, sigma, profile, print_to_screen, colorspace)
%
%  CBM3D is algorithm for attenuation of additive white Gaussian noise from 
%  color RGB images. This algorithm reproduces the results from the article:
%
%  [1] K. Dabov, A. Foi, V. Katkovnik, and K. Egiazarian, "Color image
%   denoising via sparse 3D collaborative filtering with grouping constraint in 
%   luminance-chrominance space," submitted to IEEE Int. Conf. Image Process., 
%   January 2007, in review, preprint at http://www.cs.tut.fi/~foi/GCF-BM3D.
%
%  FUNCTION INTERFACE:
%
%  [PSNR, yRGB_est] = CBM3D(yRGB, zRGB, sigma, profile, print_to_screen, colorspace)
%
%  ! The function can work without any of the input arguments, 
%   in which case, the internal default ones are used !
% 
%  BASIC USAGE EXAMPLES:
%
%     Case 1) Using the default parameters (i.e., image name, sigma, etc.)
% 
%      [PSNR, yRGB_est] = CBM3D;
% 
%     Case 2) Using an external noisy image:
%
%      % Read an RGB image and scale its intensities in range [0,1]
%      yRGB = im2double(imread('image_House256rgb.png')); 
%      % Generate the same seed used in the experimental results of [1]
%      randn('seed', 0);
%      % Standard deviation of the noise --- corresponding to intensity 
%      %  range [0,255], despite that the input was scaled in [0,1]
%      sigma = 25;
%      % Add the AWGN with zero mean and standard deviation 'sigma'
%      zRGB = yRGB + (sigma/255)*randn(size(yRGB));
%      % Denoise 'zRGB'. The denoised image is 'yRGB_est', and 'NA = 1'  
%      %  because the true image was not provided
%      [NA, yRGB_est] = CBM3D(1, zRGB, sigma); 
%      % Compute the putput PSNR
%      PSNR = 10*log10(1/mean((yRGB(:)-yRGB_est(:)).^2))
%      % show the noisy image 'zRGB' and the denoised 'yRGB_est'
%      figure; imshow(min(max(zRGB,0),1));   
%      figure; imshow(min(max(yRGB_est,0),1));
% 
%     Case 3) If the original image yRGB is provided as the first input 
%      argument, then some additional information is printed (PSNRs, 
%      figures, etc.). That is, "[NA, yRGB_est] = BM3D(1, zRGB, sigma);" in the
%      above code should be replaced with:
% 
%      [PSNR, yRGB_est] = CBM3D(yRGB, zRGB, sigma);
% 
% 
%  INPUT ARGUMENTS (OPTIONAL):
%     1) yRGB (M x N x 3): Noise-free RGB image (needed for computing PSNR),
%                           replace with the scalar 1 if not available.
%     2) zRGB (M x N x 3): Noisy RGBimage (intensities in range [0,1] or [0,255])
%     3) sigma (double)  : Std. dev. of the noise (corresponding to intensities
%                            in range [0,255] even if the range of zRGB is [0,1])
%     4) profile (char)  : 'np' --> Normal Profile 
%                          'lc' --> Fast Profile
%     5) print_to_screen : 0 --> do not print output information (and do 
%                                not plot figures)
%                          1 --> print information and plot figures
%     6) colorspace (char): 'opp'   --> use opponent colorspace
%                          'yCbCr' --> use yCbCr colorspace
%
%  OUTPUTS:
%     1) PSNR (double)          : Output PSNR (dB), only if the original 
%                                 image is available, otherwise PSNR = 0                                               
%     2) yRGB_est (M x N x 3): Final RGB estimate (in the range [0,1])
%
%
%%%%%%%%%%%%%%%%%%%%%%%%%%%%%%%%%%%%%%%%%%%%%%%%%%%%%%%%%%%%%%%%%%%%%%%%%%
%
% Copyright (c) 2007-2011 Tampere University of Technology.
% All rights reserved.
% This work should only be used for nonprofit purposes.
%
% AUTHORS:
%     Kostadin Dabov, email: dabov _at_ cs.tut.fi
%
%%%%%%%%%%%%%%%%%%%%%%%%%%%%%%%%%%%%%%%%%%%%%%%%%%%%%%%%%%%%%%%%%%%%%%%%%%

%%%%%%%%%%%%%%%%%%%%%%%%%%%%%%%%%%%%%%%%%%%%%%%%%%%%%%%%%%%%%%%%%%%%%%%%%%
%%%% In case, there is no input image (zRGB or yRGB), then use the filename 
%%%%  below to read an original image (might contain path also). Later, 
%%%%  artificial AWGN noise is added and this noisy image is processed 
%%%%  by the CBM3D.
%%%%
image_name = [
%    'kodim12.png'
   'image_Lena512rgb.png'
%     'image_House256rgb.png'
%    'image_Peppers512rgb.png'
%    'image_Baboon512rgb.png'
%    'image_F16_512rgb.png'
   ];

%%%%%%%%%%%%%%%%%%%%%%%%%%%%%%%%%%%%%%%%%%%%%%%%%%%%%%%%%%%%%%%%%%%%%%%%%%
%%%%  Quality/complexity trade-off 
%%%%
%%%%  'np' --> Normal Profile (balanced quality)
%%%%  'lc' --> Low Complexity Profile (fast, lower quality)
%%%%
%%%%  'high' --> High Profile (high quality, not documented in [1])
%%%%
%%%%  'vn' --> This profile is automatically enabled for high noise 
%%%%           when sigma > 40
%%%%
%%%%  'vn_old' --> This is the old 'vn' profile that was used in [1].
%%%%           It gives inferior results than 'vn' in most cases. 
%%%%
if (exist('profile') ~= 1)
    profile         = 'np'; %% default profile
end


%%%%%%%%%%%%%%%%%%%%%%%%%%%%%%%%%%%%%%%%%%%%%%%%%%%%%%%%%%%%%%%%%%%%%%%%%%
%%%%  Specify the std. dev. of the corrupting noise
%%%%
if (exist('sigma') ~= 1),
   sigma                = 50; %% default standard deviation of the AWGN
end

%%%%%%%%%%%%%%%%%%%%%%%%%%%%%%%%%%%%%%%%%%%%%%%%%%%%%%%%%%%%%%%%%%%%%%%%%%
%%%%  Colorspace in which we perform denoising. BM is applied to the first
%%%%  component and the matching information is re-used for the other two.
%%%%
if (exist('colorspace') ~= 1),
    colorspace              = 'opp'; %%% (valid colorspaces are: 'yCbCr' and 'opp')
end

%%%%%%%%%%%%%%%%%%%%%%%%%%%%%%%%%%%%%%%%%%%%%%%%%%%%%%%%%%%%%%%%%%%%%%%%%%
%%%% Following are the parameters for the Normal Profile.
%%%%

%%%% Select transforms ('dct', 'dst', 'hadamard', or anything that is listed by 'help wfilters'):
transform_2D_HT_name     = 'bior1.5'; %% transform used for the HT filt. of size N1 x N1
transform_2D_Wiener_name = 'dct';     %% transform used for the Wiener filt. of size N1_wiener x N1_wiener
transform_3rd_dim_name   = 'haar';    %% transform used in the 3-rd dim, the same for HT and Wiener filt.

%%%% Hard-thresholding (HT) parameters:
N1                  = 8;   %% N1 x N1 is the block size used for the hard-thresholding (HT) filtering
Nstep               = 3;   %% sliding step to process every next reference block
N2                  = 16;  %% maximum number of similar blocks (maximum size of the 3rd dimension of a 3D array)
Ns                  = 39;  %% length of the side of the search neighborhood for full-search block-matching (BM), must be odd
tau_match           = 3000;%% threshold for the block-distance (d-distance)
lambda_thr2D        = 0;   %% threshold parameter for the coarse initial denoising used in the d-distance measure
lambda_thr3D        = 2.7; %% threshold parameter for the hard-thresholding in 3D transform domain
beta                = 2.0; %% parameter of the 2D Kaiser window used in the reconstruction

%%%% Wiener filtering parameters:
N1_wiener           = 8;
Nstep_wiener        = 3;
N2_wiener           = 32;
Ns_wiener           = 39;
tau_match_wiener    = 400;
beta_wiener         = 2.0;

%%%% Block-matching parameters:
stepFS              = 1;  %% step that forces to switch to full-search BM, "1" implies always full-search
smallLN             = 'not used in np'; %% if stepFS > 1, then this specifies the size of the small local search neighb.
stepFSW             = 1;
smallLNW            = 'not used in np';
thrToIncStep        = 8;  %% used in the HT filtering to increase the sliding step in uniform regions

if strcmp(profile, 'lc') == 1,

    Nstep               = 6;
    Ns                  = 25;
    Nstep_wiener        = 5;
    N2_wiener           = 16;
    Ns_wiener           = 25;

    thrToIncStep        = 3;
    smallLN             = 3;
    stepFS              = 6*Nstep;
    smallLNW            = 2;
    stepFSW             = 5*Nstep_wiener;

end

% Profile 'vn' was proposed in 
%  Y. Hou, C. Zhao, D. Yang, and Y. Cheng, 'Comment on "Image Denoising by Sparse 3D Transform-Domain
%  Collaborative Filtering"', accepted for publication, IEEE Trans. on Image Processing, July, 2010.
% as a better alternative to that initially proposed in [1] (which is currently in profile 'vn_old')
if (strcmp(profile, 'vn') == 1) | (sigma > 40),

    N2                  = 32;
    Nstep               = 4;
 
    N1_wiener           = 11;
    Nstep_wiener        = 6;

    lambda_thr3D        = 2.8;
    thrToIncStep        = 3;
    tau_match_wiener    = 3500;
    tau_match           = 25000;
    
    Ns_wiener           = 39;
    
end

% The 'vn_old' profile corresponds to the original parameters for strong noise proposed in [1].
if (strcmp(profile, 'vn_old') == 1) & (sigma > 40),

    transform_2D_HT_name = 'dct'; 
    
    N1                  = 12;
    Nstep               = 4;
 
    N1_wiener           = 11;
    Nstep_wiener        = 6;

    lambda_thr3D        = 2.8;
    lambda_thr2D        = 2.0;
    thrToIncStep        = 3;
    tau_match_wiener    = 3500;
    tau_match           = 5000;
    
    Ns_wiener           = 39;
    
end


decLevel = 0;        %% dec. levels of the dyadic wavelet 2D transform for blocks (0 means full decomposition, higher values decrease the dec. number)
thr_mask = ones(N1); %% N1xN1 mask of threshold scaling coeff. --- by default there is no scaling, however the use of different thresholds for different wavelet decompoistion subbands can be done with this matrix

if strcmp(profile, 'high') == 1,
    
    decLevel     = 1; 
    Nstep        = 2;
    Nstep_wiener = 2;
    lambda_thr3D = 2.5;
    vMask = ones(N1,1); vMask((end/4+1):end/2)= 1.01; vMask((end/2+1):end) = 1.07; %% this allows to have different threhsolds for the finest and next-to-the-finest subbands
    thr_mask = vMask * vMask'; 
    beta         = 2.5;
    beta_wiener  = 1.5;
    
end

%%%%%%%%%%%%%%%%%%%%%%%%%%%%%%%%%%%%%%%%%%%%%%%%%%%%%%%%%%%%%%%%%%%%%%%%%%
%%%%%%%%%%%%%%%%%%%%%%%%%%%%%%%%%%%%%%%%%%%%%%%%%%%%%%%%%%%%%%%%%%%%%%%%%%
%%%% Note: touch below this point only if you know what you are doing!
%%%%%%%%%%%%%%%%%%%%%%%%%%%%%%%%%%%%%%%%%%%%%%%%%%%%%%%%%%%%%%%%%%%%%%%%%%
%%%%%%%%%%%%%%%%%%%%%%%%%%%%%%%%%%%%%%%%%%%%%%%%%%%%%%%%%%%%%%%%%%%%%%%%%%

%%% Check whether to dump information to the screen or reamin silent
dump_output_information = 1;
if (exist('print_to_screen') == 1) & (print_to_screen == 0),
    dump_output_information = 0;
end

%%%%%%%%%%%%%%%%%%%%%%%%%%%%%%%%%%%%%%%%%%%%%%%%%%%%%%%%%%%%%%%%%%%%%%%%%%
%%%% Create transform matrices, etc.
%%%%
[Tfor, Tinv]   = getTransfMatrix(N1, transform_2D_HT_name, decLevel);  %% get (normalized) forward and inverse transform matrices
[TforW, TinvW] = getTransfMatrix(N1_wiener, transform_2D_Wiener_name); %% get (normalized) forward and inverse transform matrices

if (strcmp(transform_3rd_dim_name, 'haar') == 1) | (strcmp(transform_3rd_dim_name(end-2:end), '1.1') == 1),
    %%% If Haar is used in the 3-rd dimension, then a fast internal transform is used, thus no need to generate transform
    %%% matrices.
    hadper_trans_single_den         = {};
    inverse_hadper_trans_single_den = {};
else
    %%% Create transform matrices. The transforms are later applied by
    %%% matrix-vector multiplication for the 1D case.
    for hpow = 0:ceil(log2(max(N2,N2_wiener))),
        h = 2^hpow;
        [Tfor3rd, Tinv3rd]   = getTransfMatrix(h, transform_3rd_dim_name, 0);
        hadper_trans_single_den{h}         = single(Tfor3rd);
        inverse_hadper_trans_single_den{h} = single(Tinv3rd');
    end
end

%%%%%%%%%%%%%%%%%%%%%%%%%%%%%%%%%%%%%%%%%%%%%%%%%%%%%%%%%%%%%%%%%%%%%%%%%%
%%%% 2D Kaiser windows used in the aggregation of block-wise estimates
%%%%
if beta_wiener==2 & beta==2 & N1_wiener==8 & N1==8 % hardcode the window function so that the signal processing toolbox is not needed by default
    Wwin2D = [ 0.1924    0.2989    0.3846    0.4325    0.4325    0.3846    0.2989    0.1924;
        0.2989    0.4642    0.5974    0.6717    0.6717    0.5974    0.4642    0.2989;
        0.3846    0.5974    0.7688    0.8644    0.8644    0.7688    0.5974    0.3846;
        0.4325    0.6717    0.8644    0.9718    0.9718    0.8644    0.6717    0.4325;
        0.4325    0.6717    0.8644    0.9718    0.9718    0.8644    0.6717    0.4325;
        0.3846    0.5974    0.7688    0.8644    0.8644    0.7688    0.5974    0.3846;
        0.2989    0.4642    0.5974    0.6717    0.6717    0.5974    0.4642    0.2989;
        0.1924    0.2989    0.3846    0.4325    0.4325    0.3846    0.2989    0.1924];
    Wwin2D_wiener = Wwin2D;
else
    Wwin2D           = kaiser(N1, beta) * kaiser(N1, beta)'; % Kaiser window used in the aggregation of the HT part
    Wwin2D_wiener    = kaiser(N1_wiener, beta_wiener) * kaiser(N1_wiener, beta_wiener)'; % Kaiser window used in the aggregation of the Wiener filt. part
end

%%%%%%%%%%%%%%%%%%%%%%%%%%%%%%%%%%%%%%%%%%%%%%%%%%%%%%%%%%%%%%%%%%%%%%%%%%
%%%% If needed, read images, generate noise, or scale the images to the 
%%%% [0,1] interval
%%%%
if (exist('yRGB') ~= 1) | (exist('zRGB') ~= 1)
    yRGB        = im2double(imread(image_name));  %% read a noise-free image
    randn('seed', 0);                          %% generate seed
    zRGB        = yRGB + (sigma/255)*randn(size(yRGB)); %% create a noisy image
else % external images
    image_name = 'External image';
    
    % convert zRGB to double precision
    zRGB = double(zRGB);

    % convert yRGB to double precision
    yRGB = double(yRGB);
    
    % if zRGB's range is [0, 255], then convert to [0, 1]
    if (max(zRGB(:)) > 10), % a naive check for intensity range
        zRGB = zRGB / 255;
    end
    
    % if yRGB's range is [0, 255], then convert to [0, 1]
    if (max(yRGB(:)) > 10), % a naive check for intensity range
        yRGB = yRGB / 255;
    end    
end


if (size(zRGB,3) ~= 3) | (size(zRGB,4) ~= 1),
    error('CBM3D accepts only input RGB images (i.e. matrices of size M x N x 3).');
end

% Check if the true image yRGB is a valid one; if not, then we cannot compute PSNR, etc.
yRGB_is_invalid_image = (length(size(zRGB)) ~= length(size(yRGB))) | (size(zRGB,1) ~= size(yRGB,1)) | (size(zRGB,2) ~= size(yRGB,2)) | (size(zRGB,3) ~= size(yRGB,3));
if (yRGB_is_invalid_image),
    dump_output_information = 0;
end


[Xv, Xh, numSlices] = size(zRGB);              %%% obtain image sizes

if numSlices ~= 3
    fprintf('Error, an RGB color image is required!\n');
    return;
end

%%%%%%%%%%%%%%%%%%%%%%%%%%%%%%%%%%%%%%%%%%%%%%%%%%%%%%%%%%%%%%%%%%%%%%%%%%
%%%% Change colorspace, compute the l2-norms of the new color channels
%%%%
[zColSpace l2normLumChrom] = function_rgb2LumChrom(zRGB, colorspace);

%%%%%%%%%%%%%%%%%%%%%%%%%%%%%%%%%%%%%%%%%%%%%%%%%%%%%%%%%%%%%%%%%%%%%%%%%%
%%%% Print image information to the screen
%%%%
if dump_output_information == 1,
    fprintf(sprintf('Image: %s (%dx%dx%d), sigma: %.1f\n', image_name, Xv, Xh, numSlices, sigma));
end


%%%%%%%%%%%%%%%%%%%%%%%%%%%%%%%%%%%%%%%%%%%%%%%%%%%%%%%%%%%%%%%%%%%%%%%%%%
%%%% Step 1. Basic estimate by collaborative hard-thresholding and using
%%%% the grouping constraint on the chrominances.
%%%%
tic;
y_hat = bm3d_thr_color(zColSpace, hadper_trans_single_den, Nstep, N1, N2, lambda_thr2D,...
    lambda_thr3D, tau_match*N1*N1/(255*255), (Ns-1)/2, sigma/255, thrToIncStep, single(Tfor), single(Tinv)', inverse_hadper_trans_single_den, single(thr_mask), 'unused arg', 'unused arg', l2normLumChrom, Wwin2D, smallLN, stepFS );
estimate_elapsed_time = toc;


%%%%%%%%%%%%%%%%%%%%%%%%%%%%%%%%%%%%%%%%%%%%%%%%%%%%%%%%%%%%%%%%%%%%%%%%%%
%%%% Step 2. Final estimate by collaborative Wiener filtering and using
%%%% the grouping constraint on the chrominances.
%%%%
tic;
yRGB_est = bm3d_wiener_color(zColSpace, y_hat, hadper_trans_single_den, Nstep_wiener, N1_wiener, N2_wiener, ...
    'unused_arg', tau_match_wiener*N1_wiener*N1_wiener/(255*255), (Ns_wiener-1)/2, sigma/255, 'unused arg', single(TforW), single(TinvW)', inverse_hadper_trans_single_den, 'unused arg', 'unused arg', l2normLumChrom, Wwin2D_wiener, smallLNW, stepFSW );
wiener_elapsed_time = toc;

yRGB_est = double(yRGB_est);

%%%%%%%%%%%%%%%%%%%%%%%%%%%%%%%%%%%%%%%%%%%%%%%%%%%%%%%%%%%%%%%%%%%%%%%%%%
%%%% Convert back to RGB colorspace
%%%%
yRGB_est = function_LumChrom2rgb(yRGB_est, colorspace);


%%%%%%%%%%%%%%%%%%%%%%%%%%%%%%%%%%%%%%%%%%%%%%%%%%%%%%%%%%%%%%%%%%%%%%%%%%
%%%% Calculate final estimate's PSNR and ISNR, print them, and show the
%%%% denoised image
%%%%
PSNR = 0; %% Remains 0 if the true image yRGB is not available
if (~yRGB_is_invalid_image), % then we assume yRGB is a valid image
    PSNR = 10*log10(1/mean((yRGB(:)-yRGB_est(:)).^2));
end

if dump_output_information == 1,
    fprintf(sprintf('FINAL ESTIMATE (total time: %.1f sec), PSNR: %.2f dB\n', ...
        wiener_elapsed_time + estimate_elapsed_time, PSNR));

    figure, imshow(min(max(zRGB,0),1)); title(sprintf('Noisy %s, PSNR: %.3f dB (sigma: %d)', ...
        image_name(1:end-4), 10*log10(1/mean((yRGB(:)-zRGB(:)).^2)), sigma));

    figure, imshow(min(max(yRGB_est,0),1)); title(sprintf('Denoised %s, PSNR: %.3f dB', ...
        image_name(1:end-4), PSNR));
end

return;


%%%%%%%%%%%%%%%%%%%%%%%%%%%%%%%%%%%%%%%%%%%%%%%%%%%%%%%%%%%%%%%%%%%%%%%%%%
%%%%%%%%%%%%%%%%%%%%%%%%%%%%%%%%%%%%%%%%%%%%%%%%%%%%%%%%%%%%%%%%%%%%%%%%%%
% Some auxiliary functions
%%%%%%%%%%%%%%%%%%%%%%%%%%%%%%%%%%%%%%%%%%%%%%%%%%%%%%%%%%%%%%%%%%%%%%%%%%
%%%%%%%%%%%%%%%%%%%%%%%%%%%%%%%%%%%%%%%%%%%%%%%%%%%%%%%%%%%%%%%%%%%%%%%%%%


function [Tforward, Tinverse] = getTransfMatrix (N, transform_type, dec_levels)
%
% Create forward and inverse transform matrices, which allow for perfect
% reconstruction. The forward transform matrix is normalized so that the 
% l2-norm of each basis element is 1.
%
% [Tforward, Tinverse] = getTransfMatrix (N, transform_type, dec_levels)
%
%  INPUTS:
%
%   N               --> Size of the transform (for wavelets, must be 2^K)
%
%   transform_type  --> 'dct', 'dst', 'hadamard', or anything that is 
%                       listed by 'help wfilters' (bi-orthogonal wavelets)
%                       'DCrand' -- an orthonormal transform with a DC and all
%                       the other basis elements of random nature
%
%   dec_levels      --> If a wavelet transform is generated, this is the
%                       desired decomposition level. Must be in the
%                       range [0, log2(N)-1], where "0" implies
%                       full decomposition.
%
%  OUTPUTS:
%
%   Tforward        --> (N x N) Forward transform matrix
%
%   Tinverse        --> (N x N) Inverse transform matrix
%

if exist('dec_levels') ~= 1,
    dec_levels = 0;
end

if N == 1,
    Tforward = 1;
elseif strcmp(transform_type, 'hadamard') == 1,
    Tforward    = hadamard(N);
elseif (N == 8) & strcmp(transform_type, 'bior1.5')==1 % hardcoded transform so that the wavelet toolbox is not needed to generate it
    Tforward =  [ 0.353553390593274   0.353553390593274   0.353553390593274   0.353553390593274   0.353553390593274   0.353553390593274   0.353553390593274   0.353553390593274;
       0.219417649252501   0.449283757993216   0.449283757993216   0.219417649252501  -0.219417649252501  -0.449283757993216  -0.449283757993216  -0.219417649252501;
       0.569359398342846   0.402347308162278  -0.402347308162278  -0.569359398342846  -0.083506045090284   0.083506045090284  -0.083506045090284   0.083506045090284;
      -0.083506045090284   0.083506045090284  -0.083506045090284   0.083506045090284   0.569359398342846   0.402347308162278  -0.402347308162278  -0.569359398342846;
       0.707106781186547  -0.707106781186547                   0                   0                   0                   0                   0                   0;
                       0                   0   0.707106781186547  -0.707106781186547                   0                   0                   0                   0;
                       0                   0                   0                   0   0.707106781186547  -0.707106781186547                   0                   0;
                       0                   0                   0                   0                   0                   0   0.707106781186547  -0.707106781186547];   
elseif (N == 8) & strcmp(transform_type, 'dct')==1 % hardcoded transform so that the signal processing toolbox is not needed to generate it
    Tforward = [ 0.353553390593274   0.353553390593274   0.353553390593274   0.353553390593274   0.353553390593274   0.353553390593274   0.353553390593274   0.353553390593274;
       0.490392640201615   0.415734806151273   0.277785116509801   0.097545161008064  -0.097545161008064  -0.277785116509801  -0.415734806151273  -0.490392640201615;
       0.461939766255643   0.191341716182545  -0.191341716182545  -0.461939766255643  -0.461939766255643  -0.191341716182545   0.191341716182545   0.461939766255643;
       0.415734806151273  -0.097545161008064  -0.490392640201615  -0.277785116509801   0.277785116509801   0.490392640201615   0.097545161008064  -0.415734806151273;
       0.353553390593274  -0.353553390593274  -0.353553390593274   0.353553390593274   0.353553390593274  -0.353553390593274  -0.353553390593274   0.353553390593274;
       0.277785116509801  -0.490392640201615   0.097545161008064   0.415734806151273  -0.415734806151273  -0.097545161008064   0.490392640201615  -0.277785116509801;
       0.191341716182545  -0.461939766255643   0.461939766255643  -0.191341716182545  -0.191341716182545   0.461939766255643  -0.461939766255643   0.191341716182545;
       0.097545161008064  -0.277785116509801   0.415734806151273  -0.490392640201615   0.490392640201615  -0.415734806151273   0.277785116509801  -0.097545161008064];
elseif (N == 8) & strcmp(transform_type, 'dst')==1 % hardcoded transform so that the PDE toolbox is not needed to generate it
    Tforward = [ 0.161229841765317   0.303012985114696   0.408248290463863   0.464242826880013   0.464242826880013   0.408248290463863   0.303012985114696   0.161229841765317;
       0.303012985114696   0.464242826880013   0.408248290463863   0.161229841765317  -0.161229841765317  -0.408248290463863  -0.464242826880013  -0.303012985114696;
       0.408248290463863   0.408248290463863                   0  -0.408248290463863  -0.408248290463863                   0   0.408248290463863   0.408248290463863;
       0.464242826880013   0.161229841765317  -0.408248290463863  -0.303012985114696   0.303012985114696   0.408248290463863  -0.161229841765317  -0.464242826880013;
       0.464242826880013  -0.161229841765317  -0.408248290463863   0.303012985114696   0.303012985114696  -0.408248290463863  -0.161229841765317   0.464242826880013;
       0.408248290463863  -0.408248290463863                   0   0.408248290463863  -0.408248290463863                   0   0.408248290463863  -0.408248290463863;
       0.303012985114696  -0.464242826880013   0.408248290463863  -0.161229841765317  -0.161229841765317   0.408248290463863  -0.464242826880013   0.303012985114696;
       0.161229841765317  -0.303012985114696   0.408248290463863  -0.464242826880013   0.464242826880013  -0.408248290463863   0.303012985114696  -0.161229841765317];
elseif strcmp(transform_type, 'dct') == 1,
    Tforward    = dct(eye(N));
elseif strcmp(transform_type, 'dst') == 1,
    Tforward    = dst(eye(N));
elseif strcmp(transform_type, 'DCrand') == 1,
    x = randn(N); x(1:end,1) = 1; [Q,R] = qr(x); 
    if (Q(1) < 0), 
        Q = -Q; 
    end;
    Tforward = Q';
else %% a wavelet decomposition supported by 'wavedec'
    %%% Set periodic boundary conditions, to preserve bi-orthogonality
    dwtmode('per','nodisp');  
    
    Tforward = zeros(N,N);
    for i = 1:N
        Tforward(:,i)=wavedec(circshift([1 zeros(1,N-1)],[dec_levels i-1]), log2(N), transform_type);  %% construct transform matrix
    end
end

%%% Normalize the basis elements
Tforward = (Tforward' * diag(sqrt(1./sum(Tforward.^2,2))))'; 

%%% Compute the inverse transform matrix
Tinverse = inv(Tforward);

return;

function [y, A, l2normLumChrom]=function_rgb2LumChrom(xRGB, colormode)
% Forward color-space transformation   ( inverse transformation is function_LumChrom2rgb.m )
%
% Alessandro Foi - Tampere University of Technology - 2005 - 2006   Public release v1.03 (March 2006)
% -----------------------------------------------------------------------------------------------------------------------------------------------
%
% SYNTAX:
%
%   [y A l2normLumChrom] = function_rgb2LumChrom(xRGB, colormode);
%
% INPUTS:
%   xRGB  is RGB image with range [0 1]^3
%
%   colormode = 'opp', 'yCbCr', 'pca', or a custom 3x3 matrix
%
%       'opp'     Opponent color space ('opp' is equirange version)
%       'yCbCr'   The standard yCbCr (e.g. for JPEG images)
%       'pca'     Principal components   (note that this transformation is renormalized to be equirange) 
%
% OUTPUTS:
%   y  is color-transformed image (with range typically included in or equal to [0 1]^3, depending on the transformation matrix)
%
%   l2normLumChrom (optional) l2-norm of the transformation (useful for noise std calculation)
%   A  transformation matrix  (used necessarily if colormode='pca')
%
%   NOTES:  -  If only two outputs are used, then the second output is l2normLumChrom, unless colormode='pca';
%           -  'opp' is used by default if no colormode is specified.
%
%
% USAGE EXAMPLE FOR PCA TRANSFORMATION:
%  %%%%  -- forward color transformation --
%    if colormode=='pca'
%       [zLumChrom colormode] = function_rgb2LumChrom(zRGB,colormode); % 'colormode' is assigned a 3x3 transform matrix
%    else
%       zLumChrom = function_rgb2LumChrom(zRGB,colormode);
%    end
%
%  %%%% [ ... ]  Some processing  [ ... ]
%
%  %%%%  -- inverse color transformation --
%    zRGB=function_LumChrom2rgb(zLumChrom,colormode);
%

if nargin==1
    colormode='opp';
end
change_output=0;
if size(colormode)==[3 3]
    A=colormode;
    l2normLumChrom=sqrt(sum(A.^2,2));
else
    if strcmp(colormode,'opp')
        A=[1/3 1/3 1/3; 0.5  0  -0.5; 0.25  -0.5  0.25];
    end
    if strcmp(colormode,'yCbCr')
        A=[0.299   0.587   0.114;   -0.16873660714285  -0.33126339285715   0.5;   0.5  -0.4186875  -0.0813125];
    end
    if strcmp(colormode,'pca')
        A=princomp(reshape(xRGB,[size(xRGB,1)*size(xRGB,2) 3]))';
        A=A./repmat(sum(A.*(A>0),2)-sum(A.*(A<0),2),[1 3]);  %% ranges are normalized to unitary length;
    else
        if nargout==2
            change_output=1;
        end
    end
end

%%%% Make sure that each channel's intensity range is [0,1]
maxV = sum(A.*(A>0),2);
minV = sum(A.*(A<0),2);
yNormal = (reshape(xRGB,[size(xRGB,1)*size(xRGB,2) 3]) * A' - repmat(minV, [1 size(xRGB,1)*size(xRGB,2)])') * diag(1./(maxV-minV)); % put in range [0,1]
y = reshape(yNormal, [size(xRGB,1) size(xRGB,2) 3]);

%%%% The l2-norm of each of the 3 transform basis elements 
l2normLumChrom = diag(1./(maxV-minV))*sqrt(sum(A.^2,2));

if change_output
    A=l2normLumChrom;
end

return;


function yRGB=function_LumChrom2rgb(x,colormode)
% Inverse color-space transformation   ( forward transformation is function_rgb2LumChrom.m )
%
% Alessandro Foi - Tampere University of Technology - 2005 - 2006   Public release v1.03 (March 2006)
% -----------------------------------------------------------------------------------------------------------------------------------------------
%
% SYNTAX:
%
%   yRGB = function_LumChrom2rgb(x,colormode);
%
% INPUTS:
%  x  is color-transformed image (with range typically included in or equal to [0 1]^3, depending on the transformation matrix)
%
%  colormode = 'opp', 'yCbCr', or a custom 3x3 matrix (e.g. provided by the forward transform when 'pca' is selected)
%
%       'opp'      opponent color space ('opp' is equirange version)
%       'yCbCr'    standard yCbCr (e.g. for JPEG images)
%
% OUTPUTS:
%   x  is RGB image (with range [0 1]^3)
%
%
% NOTE:    'opp' is used by default if no colormode is specified
%

if nargin==1
    colormode='opp';
end
if size(colormode)==[3 3]
    A=colormode;
    B=inv(A);
else
    if strcmp(colormode,'opp')
        A =[1/3 1/3 1/3; 0.5  0  -0.5; 0.25  -0.5  0.25];
        B =[1 1 2/3;1 0 -4/3;1 -1 2/3];
    end
    if strcmp(colormode,'yCbCr')
        A=[0.299   0.587   0.114;   -0.16873660714285  -0.33126339285715   0.5;   0.5  -0.4186875  -0.0813125];
        B=inv(A);
    end
end

%%%% Make sure that each channel's intensity range is [0,1]
maxV = sum(A.*(A>0),2);
minV = sum(A.*(A<0),2);
xNormal = reshape(x,[size(x,1)*size(x,2) 3]) * diag(maxV-minV) +  repmat(minV, [1 size(x,1)*size(x,2)])'; % put in range [0,1]
yRGB = reshape(xNormal * B', [ size(x,1) size(x,2) 3]);

return;
```
