## Supplementary material for "FPM app: an open-source application for simple and intuitive Fourier ptychographic reconstruction": FPM app: CVBM3D.m

```
function [Xdenoised] = CVBM3D(Xnoisy, sigma, Xorig)
%  CVBM3D denoising of RGB videos corrupted with AWGN.
%
%
%  [Xdenoised] = CVBM3D(Xnoisy, sigma, Xorig)
%
%  INPUTS:
%
%   1)  Xnoisy --> Either a filename of a noisy AVI RGB uncompressed video (e.g. 'SMg20.avi') 
%                  or a 4-D matrix of dimensions (M x N x 3 x NumberOfFrames)
%                  The intensity range is [0,255]!
%   2)  Sigma -->  Noise standard deviation (assumed intensity range is [0,255])
%
%   3)  Xorig     (optional parameter) --> Filename of the original video
%
%  OUTPUT: .avi files are written to the current matlab folder
%
%   1) Xdenoised --> A 4-D matrix with the denoised RGB-video
%
%  USAGE EXAMPLES:
%   1) To denoise a video:
%      CVBM3D('SMg20.avi', 20)
%
%   2) To denoise a video and print PSNR:
%      CVBM3D('SMg20.avi', 20, 'SM.avi')
%
%   1) To denoise a 4-D matrix representing a noisy RGB video:
%      CVBM3D(X_4D_matrix, 20)
%%%%%%%%%%%%%%%%%%%%%%%%%%%%%%%%%%%%%%%%%%%%%%%%%%%%%%%%%%%%%%%%%%%%%%%%%%
%
% Copyright � 2009 Tampere University of Technology. All rights reserved.
% This work should only be used for nonprofit purposes.
%
%%%%%%%%%%%%%%%%%%%%%%%%%%%%%%%%%%%%%%%%%%%%%%%%%%%%%%%%%%%%%%%%%%%%%%%%%%

% If no input argument is provided, then use the internal ones from below:
if exist('sigma', 'var') ~= 1,
    Xnoisy = 'SMg20.avi';  sigma = 20;  ;
end

% Whether or not to print information to the screen
dump_information = 1;

% If the input is a 4-D matrix, then save it as AVI file that is used as
% input to the denoising
if ischar(Xnoisy) == 0;
    NumberOfFrames = size(Xnoisy,4);

    if NumberOfFrames <= 1
        error('The input RGB video should be a 4-D matrix (M x N x 3 x NumberOfFrames)');
    end
    avi_filename = sprintf('ExternalMatrix_%.6d.avi', round(rand*50000));
    if exist(avi_filename, 'file') == 2,
        delete(avi_filename);
    end
    mov = avifile(avi_filename, 'Colormap', gray(256), 'compression', 'None', 'fps', 30);
    if mean2(Xnoisy) <= 1
        fprintf('Possible error: the input RGB-videos should be in range [0,255] and not in [0,1]!\n');
    else
        for ii = [1:NumberOfFrames],
            mov = addframe(mov, uint8(Xnoisy(:,:,:,ii)));
        end        
    end
    mov = close(mov);
    
    if dump_information == 1
        fprintf('The input 4-D matrix was written to: %s.\n', avi_filename);
    end

    clear Xnoisy
    Xnoisy = avi_filename;
end

% Read some properties of the noisy RGB video
noi_avi_file_info = aviinfo(Xnoisy);
NumberOfFrames = noi_avi_file_info.NumFrames;

%%% Read Xorig video --- needed if one wants to compute PSNR and ISNR
if exist('Xorig', 'var') == 1,
    if ischar(Xorig) == 1;    
        org_avi_file_info = aviinfo(Xorig);
        mo = aviread(Xorig);
        Xorig = zeros([size(mo(1).cdata), NumberOfFrames], 'single');
        for cf = 1:NumberOfFrames
            Xorig(:,:,:,cf) = single(mo(cf).cdata(:,:,:));
        end
        clear mo;

        if (org_avi_file_info.NumFrames == noi_avi_file_info.NumFrames && org_avi_file_info.FramesPerSecond == noi_avi_file_info.FramesPerSecond && ...
                org_avi_file_info.Width == noi_avi_file_info.Width && org_avi_file_info.Height == noi_avi_file_info.Height)
            dump_information = 1;
        end 
    else
        Xorig = single(Xorig);
        if mean2(Xorig) <= 1
            fprintf('Possible error: the input RGB-videos should be in range [0,255] and not in [0,1]!\n');
        end

    end
end

denoiseFrames  = min(9, NumberOfFrames);
denoiseFramesW = min(9, NumberOfFrames);

%%%%%%%%%%%%%%%%%%%%%%%%%%%%%%%%%%%%%%%%%%%%%%%%%%%%%%%%%%%%%%%%%%%%%%%%%%
%%%%  Quality/complexity trade-off
%%%%
%%%%  'np' --> Normal Profile (balanced quality)
%%%%  'lc' --> Low Complexity Profile (fast, lower quality)
%%%%
if (exist('bm3dProfile') ~= 1)
    bm3dProfile         = 'np';
end

%%%%%%%%%%%%%%%%%%%%%%%%%%%%%%%%%%%%%%%%%%%%%%%%%%%%%%%%%%%%%%%%%%%%%%%%%%
%%%% Following are the parameters for the Normal Profile.
%%%%

%%%% Select transforms ('dct', 'dst', 'hadamard', or anything that is listed by 'help wfilters'):
transform_2D_HT_name     = 'bior1.5'; %% transform used for the HT filt. of size N1 x N1
transform_2D_Wiener_name = 'dct';     %% transform used for the Wiener filt. of size N1_wiener x N1_wiener
transform_3rd_dim_name   = 'haar'; %% tranform used in the 3-rd dim, the same for HT and Wiener filt.

%%%% Step 1: Hard-thresholding (HT) parameters:
N1                  = 8;  %% N1 x N1 is the block size used for the hard-thresholding (HT) filtering
Nstep               = 5;  %% sliding step to process every next refernece block
N2                  = 8;  %% maximum number of similar blocks (maximum size of the 3rd dimension of the 3D groups)
Ns                  = 7;  %% length of the side of the search neighborhood for full-search block-matching (BM)
Npr                 = 3;  %% length of the side of the motion-adaptive search neighborhood, use din the predictive-search BM
tau_match           = 3000; %% threshold for the block distance (d-distance)
lambda_thr3D        = 2.7; %% threshold parameter for the hard-thresholding in 3D DFT domain
dsub                = 13;  %% a small value subtracted from the distnce of blocks with the same spatial coordinate as the reference one
Nb                  = 2;  %% number of blocks to follow in each next frame, used in the predictive-search BM
beta                = 2.0; %% the beta parameter of the 2D Kaiser window used in the reconstruction


%%%% Step 2: Wiener filtering parameters:
N1_wiener           = 7;
Nstep_wiener        = 4;
N2_wiener           = 8;
Ns_wiener           = 7;
Npr_wiener          = 3;
tau_match_wiener    = 1000;
beta_wiener         = 2.0;
dsub_wiener         = 1.5;
Nb_wiener           = 2;

%%%% Block-matching parameters:
stepFS              = 1; %% step that firces to switch to full-search BM, "1" implies always full-search
stepFSW             = 1;
thrToIncStep        = 8;  %% used in the HT filtering to increase the sliding step in uniform regions


%%%%%%%%%%%%%%%%%%%%%%%%%%%%%%%%%%%%%%%%%%%%%%%%%%%%%%%%%%%%%%%%%%%%%%%%%%
%%%% Following are the parameters for the Low Complexity Profile.
%%%%
if strcmp(bm3dProfile, 'lc') == 1,
    lambda_thr3D = 2.8;
    denoiseFrames  = min(5, NumberOfFrames);
    denoiseFramesW = min(5, NumberOfFrames);
    N2_wiener = 4;
    N2 = 4;
    Ns = 3;
    Ns_wiener = 3;
    Nb = 1;
    Nb_wiener = 1;
end

if strcmp(bm3dProfile, 'hi') == 1,
    Nstep        = 3;
    Nstep_wiener = 3;
end

if sigma > 30,
    N1_wiener = 8;
    tau_match    = 4500;
    tau_match_wiener    = 3000;
end

%%%%%%%%%%%%%%%%%%%%%%%%%%%%%%%%%%%%%%%%%%%%%%%%%%%%%%%%%%%%%%%%%%%%%%%%%%
%%%%%%%%%%%%%%%%%%%%%%%%%%%%%%%%%%%%%%%%%%%%%%%%%%%%%%%%%%%%%%%%%%%%%%%%%%
%%%% Note: touch below this point only if you know what you are doing!
%%%%%%%%%%%%%%%%%%%%%%%%%%%%%%%%%%%%%%%%%%%%%%%%%%%%%%%%%%%%%%%%%%%%%%%%%%
%%%%%%%%%%%%%%%%%%%%%%%%%%%%%%%%%%%%%%%%%%%%%%%%%%%%%%%%%%%%%%%%%%%%%%%%%%


%%%%%%%%%%%%%%%%%%%%%%%%%%%%%%%%%%%%%%%%%%%%%%%%%%%%%%%%%%%%%%%%%%%%%%%%%%
%%%% Create transform matrices, etc.
%%%%
decLevel                     = 0;    %% dec. levels of the dyadic wavelet 2D transform for blocks (0 means full decomposition, higher values decrease the dec. number)
decLevel3                    = 0;    %% dec. level for the wavelet transform in the 3rd dimension

[Tfor, Tinv]   = getTransfMatrix(N1, transform_2D_HT_name, decLevel); %% get (normalized) forward and inverse transform matrices
[TforW, TinvW] = getTransfMatrix(N1_wiener, transform_2D_Wiener_name); %% get (normalized) forward and inverse transform matrices

if (strcmp(transform_3rd_dim_name, 'haar') == 1 || strcmp(transform_3rd_dim_name(end-2:end), '1.1') == 1),
    %%% Fast internal transform is used, no need to generate transform
    %%% matrices.
    hadper_trans_single_den         = {};
    inverse_hadper_trans_single_den = {};
else
    %%% Create transform matrices. The transforms are later computed by
    %%% matrix multiplication with them
    for hh = [1 2 4 8 16 32];
        [Tfor3rd, Tinv3rd]   = getTransfMatrix(hh, transform_3rd_dim_name, decLevel3);
        hadper_trans_single_den{hh}         = single(Tfor3rd);
        inverse_hadper_trans_single_den{hh} = single(Tinv3rd');
    end
end


%%%%%%%%%%%%%%%%%%%%%%%%%%%%%%%%%%%%%%%%%%%%%%%%%%%%%%%%%%%%%%%%%%%%%%%%%%
%%%% 2D Kaiser windows that scale the reconstructed blocks
%%%%
if beta_wiener==2 & beta==2 & N1_wiener==7 & N1==8 % hardcode the window function so that the signal processing toolbox is not needed by default
    Wwin2D = [ 0.1924    0.2989    0.3846    0.4325    0.4325    0.3846    0.2989    0.1924;
        0.2989    0.4642    0.5974    0.6717    0.6717    0.5974    0.4642    0.2989;
        0.3846    0.5974    0.7688    0.8644    0.8644    0.7688    0.5974    0.3846;
        0.4325    0.6717    0.8644    0.9718    0.9718    0.8644    0.6717    0.4325;
        0.4325    0.6717    0.8644    0.9718    0.9718    0.8644    0.6717    0.4325;
        0.3846    0.5974    0.7688    0.8644    0.8644    0.7688    0.5974    0.3846;
        0.2989    0.4642    0.5974    0.6717    0.6717    0.5974    0.4642    0.2989;
        0.1924    0.2989    0.3846    0.4325    0.4325    0.3846    0.2989    0.1924 ];
    Wwin2D_wiener = [ 0.1924    0.3151    0.4055    0.4387    0.4055    0.3151    0.1924;
        0.3151    0.5161    0.6640    0.7184    0.6640    0.5161    0.3151;
        0.4055    0.6640    0.8544    0.9243    0.8544    0.6640    0.4055;
        0.4387    0.7184    0.9243    1.0000    0.9243    0.7184    0.4387;
        0.4055    0.6640    0.8544    0.9243    0.8544    0.6640    0.4055;
        0.3151    0.5161    0.6640    0.7184    0.6640    0.5161    0.3151;
        0.1924    0.3151    0.4055    0.4387    0.4055    0.3151    0.1924 ];
else
    Wwin2D           = kaiser(N1, beta) * kaiser(N1, beta)'; % Kaiser window used in the aggregation of the HT part
    Wwin2D_wiener    = kaiser(N1_wiener, beta_wiener) * kaiser(N1_wiener, beta_wiener)'; % Kaiser window used in the aggregation of the Wiener filt. part
end
%%%%%%%%%%%%%%%%%%%%%%%%%%%%%%%%%%%%%%%%%%%%%%%%%%%%%%%%%%%%%%%%%%%%%%%%%%
%%%% Read an image, generate noise and add it to the image
%%%%

if dump_information == 1
    fprintf('Input video: %s, sigma: %.1f\n', Xnoisy, sigma);
end

%%%%%%%%%%%%%%%%%%%%%%%%%%%%%%%%%%%%%%%%%%%%%%%%%%%%%%%%%%%%%%%%%%%%%%%%%%
%%%% Determine unique filenames of intermediate avi files
%%%%

HT_avi_file = sprintf('%s_cvbm3d_step1_0.avi', Xnoisy(1:end-4));
Denoised_avi_file = sprintf('%s_cvbm3d_0.avi', Xnoisy(1:end-4));
i = 1;
while (exist(['./' HT_avi_file], 'file') ~= 0) | (exist(['./' Denoised_avi_file],'file') ~= 0)
    HT_avi_file = sprintf('%s_cvbm3d_step1_%d.avi', Xnoisy(1:end-4),i);
    Denoised_avi_file = sprintf('%s_cvbm3d_%d.avi', Xnoisy(1:end-4),i);
    i = i + 1;
end

%%%%%%%%%%%%%%%%%%%%%%%%%%%%%%%%%%%%%%%%%%%%%%%%%%%%%%%%%%%%%%%%%%%%%%%%%%
%%%% Initial estimate by hard-thresholding filtering
HT_IO = {which(Xnoisy), HT_avi_file};

tic;
bm3d_thr_video_c(HT_IO, hadper_trans_single_den, Nstep, N1, N2, 0,...
    lambda_thr3D, tau_match*N1*N1/(255*255), (Ns-1)/2, sigma/255, thrToIncStep,...
    single(Tfor), single(Tinv)', inverse_hadper_trans_single_den, single(ones(N1)),...
    'unused arg', dsub*dsub/255 * (sigma^2 / 255), ones(NumberOfFrames,1), Wwin2D,...
    (Npr-1)/2, stepFS, denoiseFrames, Nb, 0 );
estimate_elapsed_time = toc;

if dump_information == 1
%     mo = aviread(HT_avi_file);
%     y_hat = zeros([size(mo(1).cdata(:,:,1)), 3, NumberOfFrames], 'single');
%     for cf = 1:NumberOfFrames
%         y_hat(:,:,:,cf) = single(mo(cf).cdata(:,:,:))/255;
%     end
%     clear  mo
% 
%     PSNR_HT_ESTIMATE = 10*log10(1/mean2((Xorig-y_hat).^2));
%     fprintf('HT ESTIMATE, PSNR: %.3f dB\n', PSNR_HT_ESTIMATE);
%     clear y_hat;
     fprintf('STEP1 completed!\n');
end

%%
%%%%%%%%%%%%%%%%%%%%%%%%%%%%%%%%%%%%%%%%%%%%%%%%%%%%%%%%%%%%%%%%%%%%%%%%%%
% %%%% Final estimate by Wiener filtering (using the hard-thresholding
% initial estimate)

lut_ic = ClipComp16b(sigma/255);

WIE_IO = {which(Xnoisy), HT_avi_file, Denoised_avi_file};

tic;
bm3d_wiener_video_c(WIE_IO, 'unused', hadper_trans_single_den, Nstep_wiener, N1_wiener, N2_wiener, ...
    'unused_arg', tau_match_wiener*N1_wiener*N1_wiener/(255*255), (Ns_wiener-1)/2, sigma/255, 'unused arg',...
    single(TforW), single(TinvW)', inverse_hadper_trans_single_den, 'unused arg', dsub_wiener*dsub_wiener/255*(sigma^2 / 255),...
    ones(NumberOfFrames,1), Wwin2D_wiener, (Npr_wiener-1)/2, stepFSW, denoiseFramesW, Nb_wiener, 0, lut_ic);

wiener_elapsed_time = toc;

if nargout == 1
    mo = aviread(Denoised_avi_file);
    Xdenoised = zeros([size(mo(1).cdata(:,:,1)), 3, NumberOfFrames], 'single');
    for cf = 1:NumberOfFrames
        Xdenoised(:,:,:,cf) = single(mo(cf).cdata(:,:,:));
    end
    clear  mo
end

if dump_information == 1
    if nargout ~= 1
        mo = aviread(Denoised_avi_file);
        Xdenoised = zeros([size(mo(1).cdata(:,:,1)), 3, NumberOfFrames], 'single');
        for cf = 1:NumberOfFrames
            Xdenoised(:,:,:,cf) = single(mo(cf).cdata(:,:,:));
        end
        clear  mo
    end
    
    PSNR_TEXT='';
    if exist('Xorig', 'var') == 1
        PSNR = 10*log10(255*255/mean((Xorig(:)-Xdenoised(:)).^2));
        PSNR_TEXT=sprintf(' PSNR: %.3f dB,', PSNR);
        New_Denoised_avi_file = sprintf('%s_PSNR%.2f.avi',Denoised_avi_file(1:end-4),PSNR);
        movefile(Denoised_avi_file, New_Denoised_avi_file);
        Denoised_avi_file = New_Denoised_avi_file;
    end

%     PSNRs = zeros(NumberOfFrames,1);
%     for ii = 1:NumberOfFrames,
%         PSNRs(ii) = 10*log10(1/mean2( (Xorig(:,:,:,ii)-Xdenoised(:,:,:,ii)).^2));
%         fprintf('Frame: %d, PSNR: %.2f\n', ii, PSNRs(ii));
%     end
    if nargout == 0
        clear Xdenoised
    end

    fprintf('FILTERING COMPLETED (frames/sec: %.2f,%s denoised video saved as %s)\n', ...
        NumberOfFrames/(wiener_elapsed_time + estimate_elapsed_time), PSNR_TEXT, Denoised_avi_file);
    
end


return;


function [Tforward, Tinverse] = getTransfMatrix (N, transform_type, dec_levels)
%
% Create forward and inverse transform matrices, which allow for perfect
% reconstruction. The forward transform matrix is normalized so that the 
% l2-norm of each basis element is 1.
%
% [Tforward, Tinverse] = getTransfMatrix (N, transform_type, dec_levels)
%
%  INPUTS:
%
%   N               --> Size of the transform (for wavelets, must be 2^K)
%
%   transform_type  --> 'dct', 'dst', 'hadamard', or anything that is 
%                       listed by 'help wfilters' (bi-orthogonal wavelets)
%                       'DCrand' -- an orthonormal transform with a DC and all
%                       the other basis elements of random nature
%
%   dec_levels      --> If a wavelet transform is generated, this is the
%                       desired decomposition level. Must be in the
%                       range [0, log2(N)-1], where "0" implies
%                       full decomposition.
%
%  OUTPUTS:
%
%   Tforward        --> (N x N) Forward transform matrix
%
%   Tinverse        --> (N x N) Inverse transform matrix
%

if exist('dec_levels') ~= 1,
    dec_levels = 0;
end

if N == 1,
    Tforward = 1;
elseif strcmp(transform_type, 'hadamard') == 1,
    Tforward    = hadamard(N);
elseif (N == 8) & strcmp(transform_type, 'bior1.5')==1 % hardcoded transform so that the wavelet toolbox is not needed to generate it
    Tforward =  [ 0.353553390593274   0.353553390593274   0.353553390593274   0.353553390593274   0.353553390593274   0.353553390593274   0.353553390593274   0.353553390593274;
       0.219417649252501   0.449283757993216   0.449283757993216   0.219417649252501  -0.219417649252501  -0.449283757993216  -0.449283757993216  -0.219417649252501;
       0.569359398342846   0.402347308162278  -0.402347308162278  -0.569359398342846  -0.083506045090284   0.083506045090284  -0.083506045090284   0.083506045090284;
      -0.083506045090284   0.083506045090284  -0.083506045090284   0.083506045090284   0.569359398342846   0.402347308162278  -0.402347308162278  -0.569359398342846;
       0.707106781186547  -0.707106781186547                   0                   0                   0                   0                   0                   0;
                       0                   0   0.707106781186547  -0.707106781186547                   0                   0                   0                   0;
                       0                   0                   0                   0   0.707106781186547  -0.707106781186547                   0                   0;
                       0                   0                   0                   0                   0                   0   0.707106781186547  -0.707106781186547];   
elseif (N == 8) & strcmp(transform_type, 'dct')==1 % hardcoded transform so that the signal processing toolbox is not needed to generate it
    Tforward = [ 0.353553390593274   0.353553390593274   0.353553390593274   0.353553390593274   0.353553390593274   0.353553390593274   0.353553390593274   0.353553390593274;
       0.490392640201615   0.415734806151273   0.277785116509801   0.097545161008064  -0.097545161008064  -0.277785116509801  -0.415734806151273  -0.490392640201615;
       0.461939766255643   0.191341716182545  -0.191341716182545  -0.461939766255643  -0.461939766255643  -0.191341716182545   0.191341716182545   0.461939766255643;
       0.415734806151273  -0.097545161008064  -0.490392640201615  -0.277785116509801   0.277785116509801   0.490392640201615   0.097545161008064  -0.415734806151273;
       0.353553390593274  -0.353553390593274  -0.353553390593274   0.353553390593274   0.353553390593274  -0.353553390593274  -0.353553390593274   0.353553390593274;
       0.277785116509801  -0.490392640201615   0.097545161008064   0.415734806151273  -0.415734806151273  -0.097545161008064   0.490392640201615  -0.277785116509801;
       0.191341716182545  -0.461939766255643   0.461939766255643  -0.191341716182545  -0.191341716182545   0.461939766255643  -0.461939766255643   0.191341716182545;
       0.097545161008064  -0.277785116509801   0.415734806151273  -0.490392640201615   0.490392640201615  -0.415734806151273   0.277785116509801  -0.097545161008064];
elseif (N == 8) & strcmp(transform_type, 'dst')==1 % hardcoded transform so that the PDE toolbox is not needed to generate it
    Tforward = [ 0.161229841765317   0.303012985114696   0.408248290463863   0.464242826880013   0.464242826880013   0.408248290463863   0.303012985114696   0.161229841765317;
       0.303012985114696   0.464242826880013   0.408248290463863   0.161229841765317  -0.161229841765317  -0.408248290463863  -0.464242826880013  -0.303012985114696;
       0.408248290463863   0.408248290463863                   0  -0.408248290463863  -0.408248290463863                   0   0.408248290463863   0.408248290463863;
       0.464242826880013   0.161229841765317  -0.408248290463863  -0.303012985114696   0.303012985114696   0.408248290463863  -0.161229841765317  -0.464242826880013;
       0.464242826880013  -0.161229841765317  -0.408248290463863   0.303012985114696   0.303012985114696  -0.408248290463863  -0.161229841765317   0.464242826880013;
       0.408248290463863  -0.408248290463863                   0   0.408248290463863  -0.408248290463863                   0   0.408248290463863  -0.408248290463863;
       0.303012985114696  -0.464242826880013   0.408248290463863  -0.161229841765317  -0.161229841765317   0.408248290463863  -0.464242826880013   0.303012985114696;
       0.161229841765317  -0.303012985114696   0.408248290463863  -0.464242826880013   0.464242826880013  -0.408248290463863   0.303012985114696  -0.161229841765317];
elseif (N == 7) & strcmp(transform_type, 'dct')==1 % hardcoded transform so that the signal processing toolbox is not needed to generate it
    Tforward =[ 0.377964473009227   0.377964473009227   0.377964473009227   0.377964473009227   0.377964473009227   0.377964473009227   0.377964473009227;
       0.521120889169602   0.417906505941275   0.231920613924330                   0  -0.231920613924330  -0.417906505941275  -0.521120889169602;
       0.481588117120063   0.118942442321354  -0.333269317528993  -0.534522483824849  -0.333269317528993   0.118942442321354   0.481588117120063;
       0.417906505941275  -0.231920613924330  -0.521120889169602                   0   0.521120889169602   0.231920613924330  -0.417906505941275;
       0.333269317528993  -0.481588117120063  -0.118942442321354   0.534522483824849  -0.118942442321354  -0.481588117120063   0.333269317528993;
       0.231920613924330  -0.521120889169602   0.417906505941275                   0  -0.417906505941275   0.521120889169602  -0.231920613924330;
       0.118942442321354  -0.333269317528993   0.481588117120063  -0.534522483824849   0.481588117120063  -0.333269317528993   0.118942442321354];   
elseif strcmp(transform_type, 'dct') == 1,
    Tforward    = dct(eye(N));
elseif strcmp(transform_type, 'dst') == 1,
    Tforward    = dst(eye(N));
elseif strcmp(transform_type, 'DCrand') == 1,
    x = randn(N); x(1:end,1) = 1; [Q,R] = qr(x); 
    if (Q(1) < 0), 
        Q = -Q; 
    end;
    Tforward = Q';
else %% a wavelet decomposition supported by 'wavedec'
    %%% Set periodic boundary conditions, to preserve bi-orthogonality
    dwtmode('per','nodisp');  
    
    Tforward = zeros(N,N);
    for i = 1:N
        Tforward(:,i)=wavedec(circshift([1 zeros(1,N-1)],[dec_levels i-1]), log2(N), transform_type);  %% construct transform matrix
    end
end

%%% Normalize the basis elements
Tforward = (Tforward' * diag(sqrt(1./sum(Tforward.^2,2))))'; 

%%% Compute the inverse transform matrix
Tinverse = inv(Tforward);

return;
```
