## Supplementary material for "FPM app: an open-source application for simple and intuitive Fourier ptychographic reconstruction": FPM app: Demo_IDDBM3D.m

```
function  [isnr, y_hat] = Demo_IDDBM3D(experiment_number, test_image_name)
% ------------------------------------------------------------------------------------------
%
%     Demo software for BM3D-frame based image deblurring
%               Public release ver. 0.8 (beta) (June 03, 2011)
%
% ------------------------------------------------------------------------------------------
%
%  This function implements the IDDBM3D image deblurring algorithm proposed in:
%
%  [1] A.Danielyan, V. Katkovnik, and K. Egiazarian, "BM3D frames and 
%   variational image deblurring," submitted to IEEE TIP, May 2011 
%
% ------------------------------------------------------------------------------------------
%
% authors:               Aram Danielyan
%                        Vladimir Katkovnik
%
% web page:              http://www.cs.tut.fi/~foi/GCF-BM3D/
%
% contact:              
%
% ------------------------------------------------------------------------------------------
% Copyright (c) 2011 Tampere University of Technology.
% All rights reserved.
% This work should be used for nonprofit purposes only.
% ------------------------------------------------------------------------------------------
%
% Disclaimer
% ----------
%
% Any unauthorized use of these routines for industrial or profit-oriented activities is
% expressively prohibited. By downloading and/or using any of these files, you implicitly
% agree to all the terms of the TUT limited license (included in the file Legal_Notice.txt).
% ------------------------------------------------------------------------------------------


%%%%%%%%%%%%%%%%%%%%%%%%%%%%%%%%%%%%%%%%%%%%%%%%%%%%%%%%%%%%%%%%%%%%%%%%%%
%  FUNCTION INTERFACE:
%
%  [psnr, y_hat] = Demo_IDDBM3D(experiment_number, test_image_name)
%  
%  INPUT:
%   1) experiment_number: 1 -> PSF 1, sigma^2 = 2
%                         2 -> PSF 1, sigma^2 = 8
%                         3 -> PSF 2, sigma^2 = 0.308
%                         4 -> PSF 3, sigma^2 = 49
%                         5 -> PSF 4, sigma^2 = 4
%                         6 -> PSF 5, sigma^2 = 64
%                         7-13 -> experiments 7-13 are not described in [1].
%                         see this file for the blur and noise parameters.
%   2) test_image_name:   a valid filename of a grayscale test image
%
%  OUTPUT:
%   1) isnr           the output improvement in SNR, dB
%   2) y_hat:         the restored image
%
%  ! The function can work without any of the input arguments, 
%   in which case, the internal default ones are used !
%   
%   To run this demo functions within the BM3D package should be accessible to Matlab 
%%%%%%%%%%%%%%%%%%%%%%%%%%%%%%%%%%%%%%%%%%%%%%%%%%%%%%%%%%%%%%%%%%%%%%%%%%
addpath('../')

if ~exist('experiment_number','var'), experiment_number=3; end
if ~exist('test_image_name','var'), test_image_name='Cameraman256.png'; end

filename=test_image_name;

if 1 % 
    initType = 'bm3ddeb'; %use output of the BM3DDEB to initialize the algorithm
else
	initType = 'zeros'; %use zero image to initialize the algorithm
end

matchType = 'bm3ddeb'; %build groups using output of the BM3DDEB algorithm
numIt = 200;

fprintf('Experiment number: %d\n', experiment_number);
fprintf('Image: %s\n', filename);

%% ------- Generating bservation ---------------------------------------------
disp('--- Generating observation ----');
y=im2double(imread(filename));

[yN,xN]=size(y);

switch experiment_number
    case 1
        sigma=sqrt(2)/255; 
        for x1=-7:7; for x2=-7:7; h(x1+8,x2+8)=1/(x1^2+x2^2+1); end, end; h=h./sum(h(:));
    case 2
        sigma=sqrt(8)/255;
        s1=0; for a1=-7:7; s1=s1+1; s2=0; for a2=-7:7; s2=s2+1; h(s1,s2)=1/(a1^2+a2^2+1); end, end;  h=h./sum(h(:));
    case 3 
        BSNR=40;
        sigma=-1; % if "sigma=-1", then the value of sigma depends on the BSNR
        h=ones(9); h=h./sum(h(:));
    case 4
        sigma=7/255;
        h=[1 4 6 4 1]'*[1 4 6 4 1]; h=h./sum(h(:));  % PSF
    case 5
        sigma=2/255;
        h=fspecial('gaussian', 25, 1.6);
    case 6
        sigma=8/255;
        h=fspecial('gaussian', 25, .4);
    %extra experiments
    case 7
        BSNR=30;
        sigma=-1;
        h=ones(9); h=h./sum(h(:));            
    case 8
        BSNR=20;
        sigma=-1;
        h=ones(9); h=h./sum(h(:));  
    case 9
        BSNR=40;
        sigma=-1;
        h=fspecial('gaussian', 25, 1.6);    
    case 10
        BSNR=20;
        sigma=-1;
        h=fspecial('gaussian', 25, 1.6);            
    case 11
        BSNR=15;
        sigma=-1; 
        h=fspecial('gaussian', 25, 1.6);    
    case 12
        BSNR=40;
        sigma=-1; % if "sigma=-1", then the value of sigma depends on the BSNR
        h=ones(19); h=h./sum(h(:));            
    case 13
        BSNR=25;
        sigma=-1; % if "sigma=-1", then the value of sigma depends on the BSNR
        h=ones(19); h=h./sum(h(:));  
end

y_blur = imfilter(y, h, 'circular'); % performs blurring (by circular convolution)

if sigma == -1;   %% check whether to use BSNR in order to define value of sigma
    sigma=sqrt(norm(y_blur(:)-mean(y_blur(:)),2)^2 /(yN*xN*10^(BSNR/10)));
    %     Xv% compute sigma from the desired BSNR
end

%%%% Create a blurred and noisy observation
randn('seed',0);
z = y_blur + sigma*randn(yN, xN);

bsnr=10*log10(norm(y_blur(:)-mean(y_blur(:)),2)^2 /sigma^2/yN/xN);
psnr_z =PSNR(y,z,1,0);
%%%%%%%%%%%%%%%%%%%%%%%%%%%%%%%%%%%%%%%%%%%%%%%%%%%%%%%%%%%%%%%%%%%%%%%

fprintf('Observation BSNR: %4.2f, PSNR: %4.2f\n', bsnr, psnr_z);

%% ----- Computing initial estimate ---------------------
disp('--- Computing initial estimate  ----');

[dummy, y_hat_RI,y_hat_RWI,zRI] = BM3DDEB_init(experiment_number, y, z, h, sigma);

switch lower(initType)
    case 'zeros'
        y_hat_init=zeros(size(z));
    case 'zri'
        y_hat_init=zRI;
    case 'ri'
        y_hat_init=y_hat_RI;
    case 'bm3ddeb'
        y_hat_init=y_hat_RWI;

end

switch lower(matchType)
    case 'z'
        match_im = z;
    case 'y'
        match_im = y;
    case 'zri'
        match_im = zRI;
    case 'ri'
        match_im = y_hat_RI;
    case 'bm3ddeb'
        match_im = y_hat_RWI;   
end

psnr_init = PSNR(y, y_hat_init,1,0);

fprintf('Initialization method: %s\n', initType);
fprintf('Initial estimate ISNR: %4.2f, PSNR: %4.2f\n', psnr_init-psnr_z, psnr_init);

%% ------- Core algorithm ---------------------
%------ Description of the parameters of the IDDBM3D function ----------
%y - true image (use [] if true image is unavaliable)
%z - observed
%h - blurring PSF
%y_hat_init - initial estimate y_0
%match_im - image used to constuct groups and calculate weights g_r
%sigma - standard deviation of the noise
%threshType = 'h'; %use 's' for soft thresholding
%numIt - number of iterations
%gamma - regularization parameter see [1]
%tau - regularization parameter see [1] (thresholding level)
%xi - regularization parameter see [1], it is always set to 1 in this implementation
%showFigure - set to True to display figure with current estimate
%--------------------------------------------------------------------

threshType = 'h';
showFigure = true;

switch threshType
    case {'s'}
        gamma_tau_xi_inits= [
            0.0004509 0.70 1;%1
            0.0006803 0.78 1;%2
            0.0003485 0.65 1;%3
            0.0005259 0.72 1;%4
            0.0005327 0.82 1;%5
            7.632e-05 0.25 1;%6
            0.0005818 0.81 1;%7
            0.001149  1.18 1;%8
            0.0004155 0.74 1;%9
            0.0005591 0.74 1;%10
            0.0007989 0.82 1;%11
            0.0006702 0.75 1;%12
            0.001931  1.83 1;%13 
        ];
    case {'h'}
        gamma_tau_xi_inits= [ 
            0.00051   3.13 1;%1
            0.0006004 2.75 1;%2
            0.0004573 2.91 1;%3
            0.0005959 2.82 1;%4
            0.0006018 3.63 1;%5
            0.0001726 2.24 1;%6
            0.00062   2.98 1;%7
            0.001047  3.80 1;%8
            0.0005125 3.00 1;%9
            0.0005685 2.80 1;%10
            0.0005716 2.75 1;%11
            0.0005938 2.55 1;%12
            0.001602  4.16 1;%13
        ];
end

gamma = gamma_tau_xi_inits(experiment_number,1);
tau   = gamma_tau_xi_inits(experiment_number,2)/255*2.7;
xi    = gamma_tau_xi_inits(experiment_number,3);

disp('-------- Start ----------');
fprintf('Number of iterations to perform: %d\n', numIt);
fprintf('Thresholding type: %s\n', threshType);

y_hat = IDDBM3D(y, h, z, y_hat_init, match_im, sigma, threshType, numIt, gamma, tau, xi, showFigure);
psnr = PSNR(y,y_hat,1,0);
isnr = psnr-psnr_z;

disp('-------- Results --------');
fprintf('Final estimate ISNR: %4.2f, PSNR: %4.2f\n', isnr, psnr);
return;

end

function PSNRdb = PSNR(x, y, maxval, borders)
    if ~exist('borders', 'var'), borders = 0; end
    if ~exist('maxval', 'var'), maxval = 255; end
    
    xx=borders+1:size(x,1)-borders;
    yy=borders+1:size(x,2)-borders;
            
    PSNRdb = zeros(1,size(x,3));
    for fr=1:size(x,3) 
        err = x(xx,yy,fr) - y(xx,yy,fr);
        PSNRdb(fr) = 10 * log10((maxval^2)/mean2(err.^2));    
    end
end
```
