## Supplementary material for "FPM app: an open-source application for simple and intuitive Fourier ptychographic reconstruction": FPM app: LEGAL_NOTICE.txt

Legal Notice

By accessing these World Wide Web pages you agree to the following terms. If you do not agree to the following terms, please notice that you are not allowed to use the site.

Copyright, author rights, trademarks and other intellectual property rights

This website and its contents are protected by copyright, author rights and/or other intellectual property rights which are the property of Tampere University of Technology ("TUT"), its researchers and/or third parties. Reproduction, modification, and use of the materials (or any information incorporated thereto such as but not limited to reports, publications, software, pictures, diagrams, video material) published on this website are hereby authorized provided that:

(i) reproduction, use, and modification are for informational and non-commercial or personal use only and will not be copied or posted on any network computer or broadcast in any media; and

(ii) any reproduction or modification retains all original notices including proprietary or copyright notices; and

(iii) reference to the original authors is given whenever results, which arise from the use of the provided material or any modification of it, are made public.

No other use of the materials and of any information incorporated thereto is hereby authorized.

In addition, be informed that some names are protected by trademarks which are the property of TUT, its researchers and/or other third parties whether a specific mention in that respect is made or not.

Disclaimers

The material, which is found on this website, is provided for general information only and should not be relied upon or used as the basis for making any transactions of any kind whatsoever. All the information and any part thereof provided on this website are provided � AS IS � without warranty of any kind either expressed or implied including, without limitation, warranties of merchantability, fitness for a particular purpose or non infringement of intellectual property rights.
TUT makes no representations or warranties as to the accuracy or completeness of any materials and information incorporated thereto and contained on this website. TUT makes no representations or warranties that access to this website will be uninterrupted or error-free, that this website (the materials and/or any information incorporated thereto) will be secure and free of virus or other harmful components.

The use of the materials (or any information incorporated thereto), in whole or in part, contained in this website is your sole responsibility. TUT disclaims any liability for any damages whatsoever including without limitation direct, indirect, incidental and/or consequential damages resulting from access to the website and use of the materials provided therein.

This website may contain links to third party sites. The links are provided to you only as a convenience and the inclusion of any link do not imply either an endorsement by TUT of the linked sites or any warranty from TUT on said sites. Access to said linked sites is at your own risk.

Transmission of user information

Any and all information or request for information you may direct to TUT through this website or through e-mail as may be linked to this website is to be considered as not confidential.

You may also address your information or request through mail to TUT's registered office for the attention of the department identified in the relevant part of this website.

Modifications

TUT reserves the right to revise the site or withdraw access to them at any time. 
