## Supplementary material for "FPM app: an open-source application for simple and intuitive Fourier ptychographic reconstruction": FPM app: README.txt

-------------------------------------------------------------------

  BM3D demo software for image/video restoration and enhancement  
                   Public release v2.00 (30 January 2014) 

-------------------------------------------------------------------

Copyright (c) 2006-2014 Tampere University of Technology. 
All rights reserved.
This work should be used for nonprofit purposes only.

Authors:                     Kostadin Dabov
                             Aram Danieyan
                             Alessandro Foi


BM3D web page:               http://www.cs.tut.fi/~foi/GCF-BM3D


-------------------------------------------------------------------
 Contents
-------------------------------------------------------------------

The package comprises these functions

*) BM3D.m        : BM3D grayscale-image denoising [1]
*) CBM3D.m       : CBM3D RGB-image denoising [2]
*) VBM3D.m       : VBM3D grayscale-video denoising [3]
*) CVBM3D.m      : CVBM3D RGB-video denoising
*) BM3DSHARP.m   : BM3D-SHARP grayscale-image sharepening & 
                   denoising [4]
*) BM3DDEB.m     : BM3D-DEB grayscale-image deblurring [5]
*) IDDBM3D\Demo_IDDBM3D : IDDBM3D grayscale-image deblurring [8]
*) BM3D-SAPCA\BM3DSAPCA2009 : BM3D-SAPCA grayscale-image denoising [9]
*) BM3D_CFA.m    : BM3D denoising of Bayer data [10]

For help on how to use these scripts, you can e.g. use "help BM3D"
or "help CBM3D".

Each demo calls MEX-functions that allow to change all possible 
parameters used in the algorithm from within the corresponding 
M-file.


-------------------------------------------------------------------
 Installation
-------------------------------------------------------------------

Unzip both BM3D.zip (contains codes) and BM3D_images.zip (contains 
test images) in a folder that is in the MATLAB path.


-------------------------------------------------------------------
 Requirements
-------------------------------------------------------------------

*) MS Windows (32 or 64 bit), Linux (32 bit or 64 bit)
   or Mac OS X (32 or 64 bit)
*) Matlab v.7.1 or later with installed:
   -- Image Processing Toolbox (for visualization with "imshow")
*) CVBM3D currently supports only 32-bit and 64-bit Windows.
*) IDDBM3D currently supports only 32-bit and 64-bit Windows and
   requires Microsoft Visual C++ 2008 SP1 Redistributable Package
   to be installed. It can be downloaded from:
    (x86) http://www.microsoft.com/downloads/en/details.aspx?FamilyID=A5C84275-3B97-4AB7-A40D-3802B2AF5FC2
    (x64) http://www.microsoft.com/downloads/en/details.aspx?FamilyID=BA9257CA-337F-4B40-8C14-157CFDFFEE4E


-------------------------------------------------------------------
 Change log
-------------------------------------------------------------------
v2.00   (30 January 2014)
 + Added BM3D_CFA denoising algorithm for Bayer data [10].
 ! Various fixes in BM3DDEB main script: now works correctly with 
   asymmetric PSFs; corrected several typos which caused first or
   second collaborative filtering stages to fail whenever the block
   sizes and 2-D transforms differed from the default ones.

v1.9    (26 August 2011)
 + Added BM3D-SAPCA denoising algorithm [9].

v1.8    (4 July 2011)
 + Added IDDBM3D deblurring algorithm [8].
 ! Improved float precision of BM3D, CBM3D, and BM3DDEB mex-files.
 
v1.7.6  (4 February 2011) 
 + Added support for Matlab running on Mac OSX 32-bit
 . Changed the strong-noise parameters ("vn" profile) in CBM3D.m,
   as proposed in [6].

v1.7.5  (7 July 2010)
 . Changed the strong-noise parameters ("vn" profile) in BM3D.m,
   as proposed in [6].

v1.7.4  (3 May 2010)
 + Added support for Matlab running on Mac OSX 64-bit

v1.7.3  (15 March 2010)
 ! Fixed a problem with writing to AVI files in CVBM3D
 ! Fixed a problem with VBM3D when the input is a 3-D matrix

v1.7.2  (8 Dec 2009)
 ! Fixed the output of CVBM3D to be in range [0,255] instead of 
   in range [0,1]

v1.7.1  (2 Dec 2009)
 ! Fixed a bug in VBM3D.m introduced in v1.7 that concerns the
   declipping

v1.7  (12 Nov 2009)
 + Added CVBM3D.m script that performs denoising on RGB-videos with
   AWGN
 ! Fixed VBM3D.m to use declipping in the case when noisy AVI file
   is provided

v1.6  (17 June 2009)
 ! Made few fixes to the "getTransfMatrix" internal function.
   If used with default parameters, BM3D no longer requires
   neither Wavelet, PDE, nor Signal Processing toolbox.
 + Added support for x86_64 Linux

v1.5.1  (20 Nov 2008)
 ! Fixed bugs for older versions of Matlab
 + Added support for 32-bit Linux
 + improved the structure of the VBM3D.m script

v1.5  (18 Oct 2008)
 + Added x86_64 version of the MEX-files that run on 64-bit Matlab 
   under Windows
 + Added a missing function in BM3DDEB.m
 + Improves some of the comments in the codes
 ! Fixed a bug in VBM3D when only a input noisy video is provided

v1.4.1  (26 Feb 2008)
 ! Fixed a bug in the grayscale-image deblurring codes and made
   these codes compatible with Matlab 7 or newer versions.

v1.4  (1 Feb 2008)
 + Added grayscale-image deblurring

v1.3  (12 Oct 2007)
 + Added grayscale-image joint sharpening and denoising

v1.2.1  (4 Sept 2007)
 ! Fixed the output of the VBM3D to be the final Wiener estimate 
   rather than the intermediate basic estimate
 ! Fixed a problem when the original video is provided as a 3D
   matrix

v1.2  (11 June 2007) 
 + Added grayscale-video denoising files

v1.1.3  (4 May 2007)
 + Added support for Linux x86-compatible platforms

v1.1.2 
 ! Fixed bugs related with Matlab v.6.1

v1.1.1  (8 March 2007)
 ! Fixed bugs related with Matlab v.6 (e.g., "isfloat" was not 
   available and "imshow" did not work with single precision)
 + Improved the usage examples shown by executing "help BM3D"
   or "help CBM3D" MATLAB commands

v1.1  (6 March 2007)
 ! Fixed a bug in comparisons of the image sizes, which was
   causing problems when executing "CBM3D(1,z,sigma);"
 ! Fixed a bug that was causing a crash when the input images are
   of type "uint8"
 ! Fixed a problem that has caused some versions of imshow to 
   report an error
 ! Fixed few typos in the comments of the functions
 . Made the parameters of the BM3D and the C-BM3D the same

v1.0  (9 December 2006)
 + Initial version, based on BM3D-DFT [7] package (November 2005)


-------------------------------------------------------------------
 References
-------------------------------------------------------------------

[1] K. Dabov, A. Foi, V. Katkovnik, and K. Egiazarian, "Image 
denoising by sparse 3D transform-domain collaborative filtering," 
IEEE Trans. Image Process., vol. 16, no. 8, August 2007.

[2] K. Dabov, A. Foi, V. Katkovnik, and K. Egiazarian, "Color 
image denoising via sparse 3D collaborative filtering with 
grouping constraint in luminance-chrominance space," Proc. IEEE
Int. Conf. Image Process., ICIP 2007, San Antonio (TX), USA, 
September 2007.

[3] K. Dabov, A. Foi, and K. Egiazarian, "Video denoising by 
sparse 3D transform-domain collaborative filtering," Proc.
European Signal Process. Conf., EUSIPCO 2007, Poznan, Poland,
September 2007.

[4] K. Dabov, A. Foi, V. Katkovnik, and K. Egiazarian, "Joint 
image sharpening and denoising by 3D transform-domain 
collaborative filtering," Proc. 2007 Int. TICSP Workshop Spectral 
Meth. Multirate Signal Process., SMMSP 2007, Moscow, Russia, 
September 2007.

[5] K. Dabov, A. Foi, and K. Egiazarian, "Image restoration by 
sparse 3D transform-domain collaborative filtering," Proc. SPIE
Electronic Imaging '08, vol. 6812, no. 6812-1D, San Jose (CA),
USA, January 2008.

[6] Y. Hou, C. Zhao, D. Yang, and Y. Cheng, 'Comment on "Image 
Denoising by Sparse 3D Transform-Domain Collaborative Filtering"'
accepted for publication, IEEE Trans. Image Process., July, 2010.

[7] K. Dabov, A. Foi, V. Katkovnik, and K. Egiazarian, "Image
denoising with block-matching and 3D filtering," Proc. SPIE
Electronic Imaging '06, vol. 6064, no. 6064A-30, San Jose (CA),
USA, January 2006.

[8] A.Danielyan, V. Katkovnik, and K. Egiazarian, "BM3D frames and 
variational image deblurring," accepted for publication in IEEE
Trans. Image Process.
Preprint online at http://www.cs.tut.fi/~foi/GCF-BM3D

[9] K. Dabov, A. Foi, V. Katkovnik, and K. Egiazarian, "BM3D Image
Denoising with Shape-Adaptive Principal Component Analysis", Proc.
Workshop on Signal Processing with Adaptive Sparse Structured
Representations (SPARS'09), Saint-Malo, France, April 2009.

[10] A. Danielyan, M. Vehvil�inen, A. Foi, V. Katkovnik, and
K. Egiazarian, "Cross-color BM3D filtering of noisy raw data", 
Proc. Int. Workshop on Local and Non-Local Approx. in Image Process.,
LNLA 2009, Tuusula, Finland, pp. 125-129, August 2009.

 
-------------------------------------------------------------------
 Disclaimer
-------------------------------------------------------------------

Any unauthorized use of these routines for industrial or profit-
oriented activities is expressively prohibited. By downloading 
and/or using any of these files, you implicitly agree to all the 
terms of the TUT limited license:
http://www.cs.tut.fi/~foi/GCF-BM3D/legal_notice.html


-------------------------------------------------------------------
 Feedback
-------------------------------------------------------------------

If you have any comment, suggestion, or question, please do
contact    Alessandro Foi   at 
