## Supplementary material for "FPM app: an open-source application for simple and intuitive Fourier ptychographic reconstruction": FPM app: VBM3D.m

```
function [PSNR_FINAL_ESTIMATE, y_hat_wi] = VBM3D(Xnoisy, sigma, NumberOfFrames, dump_information, Xorig, bm3dProfile)
%%%%%%%%%%%%%%%%%%%%%%%%%%%%%%%%%%%%%%%%%%%%%%%%%%%%%%%%%%%%%%%%%%%%%%%%%%
%
%  VBM3D is a Matlab function for attenuation of additive white Gaussian 
%  noise from grayscale videos. This algorithm reproduces the results from the article:
%
%  [1] K. Dabov, A. Foi, and K. Egiazarian, "Video denoising by sparse 3D
%  transform-domain collaborative filtering," European Signal Processing
%  Conference (EUSIPCO-2007), September 2007. (accepted)
%
%  INTERFACE:
%
%  [PSNR, Xest] = VBM3D(Xnoisy, Sigma, NFrames, PrintInfo, Xorig)
%
%  INPUTS:
%   1)  Xnoisy     --> A filename of a noisy .avi video, e.g. Xnoisy = 'gstennisg20.avi'
%        OR
%       Xnoisy     --> A 3D matrix of a noisy video in a  (floating point data in range [0,1],
%                                                     or in [0,255])
%   2)  Sigma --> Noise standard deviation (assumed range is [0,255], no matter what is
%                                           the input's range)
%
%   3)  NFrames   (optional paremter!) --> Number of frames to process. If set to 0 or 
%                                          ommited, then process all frames (default: 0).
%
%   4)  PrintInfo (optional paremter!) --> If non-zero, then print to screen and save 
%                                          the denoised video in .AVI
%                                          format. (default: 1)
%
%   5)  Xorig     (optional paremter!) --> Original video's filename or 3D matrix 
%                                          If provided, PSNR, ISNR will be computed.
%
%   NOTE: If Xorig == Xnoisy, then artificial noise is added internally and the
%   obtained noisy video is denoised.
%
%  OUTPUTS:
%  
%   1) PSNR --> If Xorig is valid video, then this contains the PSNR of the
%                denoised one
%
%   1) Xest --> Final video estimate in a 3D matrix (intensities in range [0,1])
%
%   *) If "PrintInfo" is non-zero, then save the denoised video in the current 
%       MATLAB folder.
%
%  USAGE EXAMPLES:
%
%     1) Denoise a noisy (clipped in [0,255] range) video sequence, e.g. 
%        'gsalesmang20.avi' corrupted with AWGN with std. dev. 20:
%          
%          Xest = VBM3D('gsalesmang20.avi', 20, 0, 1); 
%     
%     2) The same, but also print PSNR, ISNR numbers.
%        
%          Xest = VBM3D('gsalesmang20.avi', 20, 0, 1, 'gsalesman.avi');
%
%     3) Add artificial noise to a video, then denoise it (without 
%        considering clipping in [0,255]):
%        
%          Xest = VBM3D('gsalesman.avi', 20, 0, 1, 'gsalesman.avi');
%  
%
%  RESTRICTIONS:
%
%     Since the video sequences are read into memory as 3D matrices,
%     there apply restrictions on the input video size, which are thus
%     proportional to the maximum memory allocatable by Matlab.
%
%%%%%%%%%%%%%%%%%%%%%%%%%%%%%%%%%%%%%%%%%%%%%%%%%%%%%%%%%%%%%%%%%%%%%%%%%%
%
% Copyright � 2007 Tampere University of Technology. All rights reserved.
% This work should only be used for nonprofit purposes.
%
% AUTHORS:
%     Kostadin Dabov, email: dabov _at_ cs.tut.fi
%
%%%%%%%%%%%%%%%%%%%%%%%%%%%%%%%%%%%%%%%%%%%%%%%%%%%%%%%%%%%%%%%%%%%%%%%%%%

% If no input argument is provided, then use these internal ones:
if exist('sigma', 'var') ~= 1,
    Xnoisy = 'gsalesmang20.avi'; Xorig = 'gsalesman.avi'; sigma = 20;
    %Xnoisy = 'gstennisg20.avi';  Xorig = 'gstennis.avi';  sigma = 20;
    %Xnoisy = 'gflowersg20.avi';   Xorig = 'gflower.avi';   sigma = 20;
    
    %Xnoisy = 'gsalesman.avi'; Xorig = Xnoisy; sigma = 20;
    
    NumberOfFrames = 0; %% 0 means process ALL frames.
end


if exist('dump_information', 'var') ~= 1,
    dump_information = 1; % 1 -> print informaion to the screen and save the processed video as an AVI file
end

%%%%%%%%%%%%%%%%%%%%%%%%%%%%%%%%%%%%%%%%%%%%%%%%%%%%%%%%%%%%%%%%%%%%%%%%%%
%%%% Obtain infromation about the input noisy video
%%%%
if (ischar(Xnoisy) == 1), % if the input is a video filename
    isCharacterName = 1;
    Xnoisy_name = Xnoisy;
    videoInfo = aviinfo(Xnoisy);
    videoHeight = videoInfo.Height;
    videoWidth = videoInfo.Width;
    TotalFrames = videoInfo.NumFrames;
elseif length(size(Xnoisy)) == 3% the input argument is a 3D video (spatio-temporal) matrix
    Xnoisy_name = 'Input 3D matrix';
    isCharacterName = 0;
    [videoHeight, videoWidth, TotalFrames] = size(Xnoisy);
else
    fprintf('Oops! The input argument Xnoisy should be either a filename or a 3D matrix!\n');
    PSNR_FINAL_ESTIMATE = 0;
    y_hat_wi = 0;
    return;
end

%%%%%%%%%%%%%%%%%%%%%%%%%%%%%%%%%%%%%%%%%%%%%%%%%%%%%%%%%%%%%%%%%%%%%%%%%%
%%%% Check if we want to process all frames, and save as 'NumberOfFrames' 
%%%% the desired number of frames to process
%%%%
if exist('NumberOfFrames', 'var') == 1,
    if NumberOfFrames <= 0,
        NumberOfFrames = TotalFrames;
    else
        NumberOfFrames = max(min(NumberOfFrames, TotalFrames), 1);
    end    
else
    NumberOfFrames = TotalFrames;
end

%%%%%%%%%%%%%%%%%%%%%%%%%%%%%%%%%%%%%%%%%%%%%%%%%%%%%%%%%%%%%%%%%%%%%%%%%%
%%%%  Quality/complexity trade-off
%%%%
%%%%  'np' --> Normal Profile (balanced quality)
%%%%  'lc' --> Low Complexity Profile (fast, lower quality)
%%%%
if (exist('bm3dProfile', 'var') ~= 1)
    bm3dProfile         = 'np';
end

%%%%%%%%%%%%%%%%%%%%%%%%%%%%%%%%%%%%%%%%%%%%%%%%%%%%%%%%%%%%%%%%%%%%%%%%%%
%%%% Parameters for the Normal Profile.
%%%%
%%%% Select transforms ('dct', 'dst', 'hadamard', or anything that is listed by 'help wfilters'):
transform_2D_HT_name     = 'bior1.5'; %% transform used for the HT filt. of size N1 x N1
transform_2D_Wiener_name = 'dct';     %% transform used for the Wiener filt. of size N1_wiener x N1_wiener
transform_3rd_dim_name   = 'haar'; %% tranform used in the 3-rd dim, the same for HT and Wiener filt.

%%%% Step 1: Hard-thresholding (HT) parameters:
denoiseFrames       = min(9, NumberOfFrames); % number of frames in the temporalwindow (should not exceed the total number of frames 'NumberOfFrames')
N1                  = 8;  %% N1 x N1 is the block size used for the hard-thresholding (HT) filtering
Nstep               = 6;  %% sliding step to process every next refernece block
N2                  = 8;  %% maximum number of similar blocks (maximum size of the 3rd dimension of the 3D groups)
Ns                  = 7;  %% length of the side of the search neighborhood for full-search block-matching (BM)
Npr                 = 5;  %% length of the side of the motion-adaptive search neighborhood, use din the predictive-search BM
tau_match           = 3000; %% threshold for the block distance (d-distance)
lambda_thr3D        = 2.7; %% threshold parameter for the hard-thresholding in 3D DFT domain
dsub                = 7;  %% a small value subtracted from the distnce of blocks with the same spatial coordinate as the reference one 
Nb                  = 2;  %% number of blocks to follow in each next frame, used in the predictive-search BM
beta                = 2.0; %% the beta parameter of the 2D Kaiser window used in the reconstruction

%%%% Step 2: Wiener filtering parameters:
denoiseFramesW      = min(9, NumberOfFrames);
N1_wiener           = 7;
Nstep_wiener        = 4;
N2_wiener           = 8;
Ns_wiener           = 7;
Npr_wiener          = 5;
tau_match_wiener    = 1500;
beta_wiener         = 2.0;
dsub_wiener         = 3;
Nb_wiener           = 2;

%%%% Block-matching parameters:
stepFS              = 1; %% step that forces to switch to full-search BM, "1" implies always full-search
smallLN             = 3; %% if stepFS > 1, then this specifies the size of the small local search neighb.
stepFSW             = 1;
smallLNW            = 3;
thrToIncStep        = 8;  %% used in the HT filtering to increase the sliding step in uniform regions


%%%%%%%%%%%%%%%%%%%%%%%%%%%%%%%%%%%%%%%%%%%%%%%%%%%%%%%%%%%%%%%%%%%%%%%%%%
%%%% Parameters for the Low Complexity Profile.
%%%%
if strcmp(bm3dProfile, 'lc') == 1,
    lambda_thr3D = 2.8;
    smallLN   = 2;
    smallLNW  = 2;
    denoiseFrames  = min(5, NumberOfFrames);
    denoiseFramesW = min(5, NumberOfFrames);
    N2_wiener = 4;
    N2 = 4;
    Ns = 3;
    Ns_wiener = 3;
    NB = 1;
    Nb_wiener = 1;
end

%%%%%%%%%%%%%%%%%%%%%%%%%%%%%%%%%%%%%%%%%%%%%%%%%%%%%%%%%%%%%%%%%%%%%%%%%%
%%%% Parameters for the High Profile.
%%%%
if strcmp(bm3dProfile, 'hi') == 1,
    Nstep        = 3;
    Nstep_wiener = 3;
end

%%%%%%%%%%%%%%%%%%%%%%%%%%%%%%%%%%%%%%%%%%%%%%%%%%%%%%%%%%%%%%%%%%%%%%%%%%
%%%% Parameters for the "Very Noisy" Profile.
%%%%
if sigma > 30,
    N1 = 8;
    N1_wiener = 8;
    Nstep = 6;
    tau_match    = 4500;
    tau_match_wiener    = 3000;
end


%%%%%%%%%%%%%%%%%%%%%%%%%%%%%%%%%%%%%%%%%%%%%%%%%%%%%%%%%%%%%%%%%%%%%%%%%%
%%%%%%%%%%%%%%%%%%%%%%%%%%%%%%%%%%%%%%%%%%%%%%%%%%%%%%%%%%%%%%%%%%%%%%%%%%
%%%% Note: touch below this point only if you know what you are doing!
%%%%%%%%%%%%%%%%%%%%%%%%%%%%%%%%%%%%%%%%%%%%%%%%%%%%%%%%%%%%%%%%%%%%%%%%%%
%%%%%%%%%%%%%%%%%%%%%%%%%%%%%%%%%%%%%%%%%%%%%%%%%%%%%%%%%%%%%%%%%%%%%%%%%%

%%%%%%%%%%%%%%%%%%%%%%%%%%%%%%%%%%%%%%%%%%%%%%%%%%%%%%%%%%%%%%%%%%%%%%%%%%
%%%% Extract the input noisy video and make sure intensities are in [0,1]
%%%% interval, using single-precision float
if isCharacterName,
    mno = aviread(Xnoisy_name);
    z = zeros([videoHeight, videoWidth, NumberOfFrames], 'single');
    for cf = 1:NumberOfFrames
        z(:,:,cf) = single(mno(cf).cdata(:,:,1)) * 0.0039216; % 1/255 = 0.0039216
    end
    clear  mno
else
    if isinteger(Xnoisy) == 1,
        z = single(Xnoisy) * 0.0039216; % 1/255 = 0.0039216
    elseif isfloat(Xnoisy) == 0,
        fprintf('Unknown format of "Xnoisy"! Must be a filename (array of char) or a 3D array of either floating point data (range [0,1]) or integer data (range [0,255]). \n');
        return;
    else        
        z = single(Xnoisy);
    end
end

clear Xnoisy;

%%%% If the original video is provided, then extract it to 'Xorig' 
%%%% which is later used to compute PSNR and ISNR
if exist('Xorig', 'var') == 1,
    randn('seed', 0);
    if ischar(Xorig) == 0,
        if isinteger(Xorig) == 1,
            y = single(Xorig) * 0.0039216; % 1/255 = 0.0039216
        elseif isfloat(Xorig) == 0,
            fprintf('Unknown format of "Xorig"! Must be a filename (array of char) or a 3D array of either floating point data (range [0,1]) or integer data (range [0,255]). \n');
            return;            
        else
            y = single(Xorig);
        end
    else        
        if strcmp(Xorig, Xnoisy_name) == 1, %% special case, noise is aritifically added
            y = z;
            z = z + (sigma/255) * randn(size(z));
        else
            mo = aviread(Xorig);
            y = zeros([videoHeight, videoWidth, NumberOfFrames], 'single');
            for cf = 1:NumberOfFrames
                y(:,:,cf) = single(mo(cf).cdata(:,:,1)) * 0.0039216; % 1/255 = 0.0039216
            end
            clear mo
        end
    end
end

%%%%%%%%%%%%%%%%%%%%%%%%%%%%%%%%%%%%%%%%%%%%%%%%%%%%%%%%%%%%%%%%%%%%%%%%%%
%%%% Create transform matrices, etc.
%%%%
decLevel           = 0;    %% dec. levels of the dyadic wavelet 2D transform for blocks (0 means full decomposition, higher values decrease the dec. number)
decLevel3          = 0;    %% dec. level for the wavelet transform in the 3rd dimension

[Tfor, Tinv]       = getTransfMatrix(N1, transform_2D_HT_name, decLevel); %% get (normalized) forward and inverse transform matrices
[TforW, TinvW]     = getTransfMatrix(N1_wiener, transform_2D_Wiener_name); %% get (normalized) forward and inverse transform matrices
thr_mask           = ones(N1); %% N1xN1 mask of threshold scaling coeff. --- by default there is no scaling, however the use of different thresholds for different wavelet decompoistion subbands can be done with this matrix

if (strcmp(transform_3rd_dim_name, 'haar') == 1 || strcmp(transform_3rd_dim_name(end-2:end), '1.1') == 1),
    %%% Fast internal transform is used, no need to generate transform
    %%% matrices.
    hadper_trans_single_den         = {};
    inverse_hadper_trans_single_den = {};
else
    %%% Create transform matrices. The transforms are later computed by
    %%% matrix multiplication with them
    for hh = [1 2 4 8 16 32];
        [Tfor3rd, Tinv3rd]   = getTransfMatrix(hh, transform_3rd_dim_name, decLevel3);
        hadper_trans_single_den{hh}         = single(Tfor3rd);
        inverse_hadper_trans_single_den{hh} = single(Tinv3rd');
    end
end

%%%%%%%%%%%%%%%%%%%%%%%%%%%%%%%%%%%%%%%%%%%%%%%%%%%%%%%%%%%%%%%%%%%%%%%%%%
%%%% 2D Kaiser windows that scale the reconstructed blocks
%%%%
if beta_wiener==2 & beta==2 & N1_wiener==7 & N1==8 % hardcode the window function so that the signal processing toolbox is not needed by default
    Wwin2D = [ 0.1924    0.2989    0.3846    0.4325    0.4325    0.3846    0.2989    0.1924;
        0.2989    0.4642    0.5974    0.6717    0.6717    0.5974    0.4642    0.2989;
        0.3846    0.5974    0.7688    0.8644    0.8644    0.7688    0.5974    0.3846;
        0.4325    0.6717    0.8644    0.9718    0.9718    0.8644    0.6717    0.4325;
        0.4325    0.6717    0.8644    0.9718    0.9718    0.8644    0.6717    0.4325;
        0.3846    0.5974    0.7688    0.8644    0.8644    0.7688    0.5974    0.3846;
        0.2989    0.4642    0.5974    0.6717    0.6717    0.5974    0.4642    0.2989;
        0.1924    0.2989    0.3846    0.4325    0.4325    0.3846    0.2989    0.1924 ];
    Wwin2D_wiener = [ 0.1924    0.3151    0.4055    0.4387    0.4055    0.3151    0.1924;
        0.3151    0.5161    0.6640    0.7184    0.6640    0.5161    0.3151;
        0.4055    0.6640    0.8544    0.9243    0.8544    0.6640    0.4055;
        0.4387    0.7184    0.9243    1.0000    0.9243    0.7184    0.4387;
        0.4055    0.6640    0.8544    0.9243    0.8544    0.6640    0.4055;
        0.3151    0.5161    0.6640    0.7184    0.6640    0.5161    0.3151;
        0.1924    0.3151    0.4055    0.4387    0.4055    0.3151    0.1924 ];
else
    Wwin2D           = kaiser(N1, beta) * kaiser(N1, beta)'; % Kaiser window used in the aggregation of the HT part
    Wwin2D_wiener    = kaiser(N1_wiener, beta_wiener) * kaiser(N1_wiener, beta_wiener)'; % Kaiser window used in the aggregation of the Wiener filt. part
end

%%%%%%%%%%%%%%%%%%%%%%%%%%%%%%%%%%%%%%%%%%%%%%%%%%%%%%%%%%%%%%%%%%%%%%%%%%
%%%% Read an image, generate noise and add it to the image
%%%%

l2normLumChrom = ones(NumberOfFrames,1); %%% NumberOfFrames == nSl !

if dump_information == 1,
    fprintf('Video: %s (%dx%dx%d), sigma: %.1f\n', Xnoisy_name, videoHeight, videoWidth, NumberOfFrames, sigma);
end

%%%%%%%%%%%%%%%%%%%%%%%%%%%%%%%%%%%%%%%%%%%%%%%%%%%%%%%%%%%%%%%%%%%%%%%%%%
%%%% Initial estimate by hard-thresholding filtering
tic;
y_hat = bm3d_thr_video(z, hadper_trans_single_den, Nstep, N1, N2, 0,...
    lambda_thr3D, tau_match*N1*N1/(255*255), (Ns-1)/2, sigma/255, thrToIncStep, single(Tfor), single(Tinv)', inverse_hadper_trans_single_den, single(thr_mask), 'unused arg', dsub*dsub/255, l2normLumChrom, Wwin2D, (Npr-1)/2, stepFS, denoiseFrames, Nb );
estimate_elapsed_time = toc;

if exist('Xorig', 'var') == 1,
    PSNR_INITIAL_ESTIMATE = 10*log10(1/mean((double(y(:))-double(y_hat(:))).^2));
    PSNR_NOISE = 10*log10(1/mean((double(y(:))-double(z(:))).^2));
    ISNR_INITIAL_ESTIMATE = PSNR_INITIAL_ESTIMATE - PSNR_NOISE;

    if dump_information == 1,    
        fprintf('BASIC ESTIMATE (time: %.1f sec), PSNR: %.3f dB, ISNR: %.3f dB\n', ...
            estimate_elapsed_time, PSNR_INITIAL_ESTIMATE, ISNR_INITIAL_ESTIMATE);
    end
end
      

%%%%%%%%%%%%%%%%%%%%%%%%%%%%%%%%%%%%%%%%%%%%%%%%%%%%%%%%%%%%%%%%%%%%%%%%%%
% %%%% Final estimate by Wiener filtering (using the hard-thresholding
% initial estimate)
tic;
y_hat_wi = bm3d_wiener_video(z, y_hat, hadper_trans_single_den, Nstep_wiener, N1_wiener, N2_wiener, ...
    'unused_arg', tau_match_wiener*N1_wiener*N1_wiener/(255*255), (Ns_wiener-1)/2, sigma/255, 'unused arg', single(TforW), single(TinvW)', inverse_hadper_trans_single_den, 'unused arg', dsub_wiener*dsub_wiener/255, l2normLumChrom, Wwin2D_wiener, (Npr_wiener-1)/2, stepFSW, denoiseFramesW, Nb_wiener );

% In case the input noisy video is clipped in [0,1], then apply declipping  
if isCharacterName
    if exist('Xorig', 'var') == 1
        if ~strcmp(Xorig, Xnoisy_name)
            [y_hat_wi] = ClipComp16b(sigma/255, y_hat_wi);
        end
    else
        [y_hat_wi] = ClipComp16b(sigma/255, y_hat_wi);
    end
end

wiener_elapsed_time = toc;


PSNR_FINAL_ESTIMATE = 0;
if exist('Xorig', 'var') == 1,
    PSNR_FINAL_ESTIMATE = 10*log10(1/mean((double(y(:))-double(y_hat_wi(:))).^2)); 
    ISNR_FINAL_ESTIMATE = PSNR_FINAL_ESTIMATE - 10*log10(1/mean((double(y(:))-double(z(:))).^2));
end

if dump_information == 1,

    text_psnr = '';
    if exist('Xorig', 'var') == 1

        
        %%%% Un-comment the following to print the PSNR of each frame
        %
        %     PSNRs = zeros(NumberOfFrames,1);
        %     for ii = [1:NumberOfFrames],
        %         PSNRs(ii) = 10*log10(1/mean2((y(:,:,ii)-y_hat_wi(:,:,ii)).^2));
        %         fprintf(['Frame: ' sprintf('%d',ii) ', PSNR: ' sprintf('%.2f',PSNRs(ii)) '\n']);
        %     end
        %

        fprintf('FINAL ESTIMATE, PSNR: %.3f dB, ISNR: %.3f dB\n', ...
             PSNR_FINAL_ESTIMATE, ISNR_FINAL_ESTIMATE);

        figure, imshow(double(z(:,:,ceil(NumberOfFrames/2)))); % show the central frame
        title(sprintf('Noisy frame #%d',ceil(NumberOfFrames/2)));           
        
        figure, imshow(double(y_hat_wi(:,:,ceil(NumberOfFrames/2)))); % show the central frame
        title(sprintf('Denoised frame #%d',ceil(NumberOfFrames/2)));
        
        text_psnr = sprintf('_PSNR%.2f', PSNR_FINAL_ESTIMATE);
    end
    
    fprintf('The denoising took: %.1f sec (%.4f sec/frame). ', ...
        wiener_elapsed_time+estimate_elapsed_time, (wiener_elapsed_time+estimate_elapsed_time)/NumberOfFrames);

    
    text_vid = 'Denoised';
    FRATE = 30; % default value
    if isCharacterName,
        text_vid = Xnoisy_name(1:end-4);
        ainfo = aviinfo(Xnoisy_name);
        FRATE = ainfo.FramesPerSecond;
    end

    avi_filename = sprintf('%s%s_%s_BM3D.avi', text_vid, text_psnr, bm3dProfile);
    
    if exist(avi_filename, 'file') ~= 0,
        delete(avi_filename);
    end
    mov = avifile(avi_filename, 'Colormap', gray(256), 'compression', 'None', 'fps', FRATE);
    for ii = [1:NumberOfFrames],
        mov = addframe(mov, uint8(round(255*double(y_hat_wi(:,:,ii)))));
    end
    mov = close(mov);
    fprintf('The denoised video written to: %s.\n\n', avi_filename);
    
end

return;


%%%%%%%%%%%%%%%%%%%%%%%%%%%%%%%%%%%%%%%%%%%%%%%%%%%%%%%%%%%%%%%%%%%%%%%%%%
%%%%%%%%%%%%%%%%%%%%%%%%%%%%%%%%%%%%%%%%%%%%%%%%%%%%%%%%%%%%%%%%%%%%%%%%%%
% Some auxiliary functions 
%%%%%%%%%%%%%%%%%%%%%%%%%%%%%%%%%%%%%%%%%%%%%%%%%%%%%%%%%%%%%%%%%%%%%%%%%%
%%%%%%%%%%%%%%%%%%%%%%%%%%%%%%%%%%%%%%%%%%%%%%%%%%%%%%%%%%%%%%%%%%%%%%%%%%


function [Tforward, Tinverse] = getTransfMatrix (N, transform_type, dec_levels)
%
% Create forward and inverse transform matrices, which allow for perfect
% reconstruction. The forward transform matrix is normalized so that the 
% l2-norm of each basis element is 1.
%
% [Tforward, Tinverse] = getTransfMatrix (N, transform_type, dec_levels)
%
%  INPUTS:
%
%   N               --> Size of the transform (for wavelets, must be 2^K)
%
%   transform_type  --> 'dct', 'dst', 'hadamard', or anything that is 
%                       listed by 'help wfilters' (bi-orthogonal wavelets)
%                       'DCrand' -- an orthonormal transform with a DC and all
%                       the other basis elements of random nature
%
%   dec_levels      --> If a wavelet transform is generated, this is the
%                       desired decomposition level. Must be in the
%                       range [0, log2(N)-1], where "0" implies
%                       full decomposition.
%
%  OUTPUTS:
%
%   Tforward        --> (N x N) Forward transform matrix
%
%   Tinverse        --> (N x N) Inverse transform matrix
%

if exist('dec_levels', 'var') ~= 1,
    dec_levels = 0;
end

if N == 1,
    Tforward = 1;
elseif strcmp(transform_type, 'hadamard') == 1,
    Tforward    = hadamard(N);
elseif (N == 8) & strcmp(transform_type, 'bior1.5')==1 % hardcoded transform so that the wavelet toolbox is not needed to generate it
    Tforward =  [ 0.353553390593274   0.353553390593274   0.353553390593274   0.353553390593274   0.353553390593274   0.353553390593274   0.353553390593274   0.353553390593274;
       0.219417649252501   0.449283757993216   0.449283757993216   0.219417649252501  -0.219417649252501  -0.449283757993216  -0.449283757993216  -0.219417649252501;
       0.569359398342846   0.402347308162278  -0.402347308162278  -0.569359398342846  -0.083506045090284   0.083506045090284  -0.083506045090284   0.083506045090284;
      -0.083506045090284   0.083506045090284  -0.083506045090284   0.083506045090284   0.569359398342846   0.402347308162278  -0.402347308162278  -0.569359398342846;
       0.707106781186547  -0.707106781186547                   0                   0                   0                   0                   0                   0;
                       0                   0   0.707106781186547  -0.707106781186547                   0                   0                   0                   0;
                       0                   0                   0                   0   0.707106781186547  -0.707106781186547                   0                   0;
                       0                   0                   0                   0                   0                   0   0.707106781186547  -0.707106781186547];   
elseif (N == 8) & strcmp(transform_type, 'dct')==1 % hardcoded transform so that the signal processing toolbox is not needed to generate it
    Tforward = [ 0.353553390593274   0.353553390593274   0.353553390593274   0.353553390593274   0.353553390593274   0.353553390593274   0.353553390593274   0.353553390593274;
       0.490392640201615   0.415734806151273   0.277785116509801   0.097545161008064  -0.097545161008064  -0.277785116509801  -0.415734806151273  -0.490392640201615;
       0.461939766255643   0.191341716182545  -0.191341716182545  -0.461939766255643  -0.461939766255643  -0.191341716182545   0.191341716182545   0.461939766255643;
       0.415734806151273  -0.097545161008064  -0.490392640201615  -0.277785116509801   0.277785116509801   0.490392640201615   0.097545161008064  -0.415734806151273;
       0.353553390593274  -0.353553390593274  -0.353553390593274   0.353553390593274   0.353553390593274  -0.353553390593274  -0.353553390593274   0.353553390593274;
       0.277785116509801  -0.490392640201615   0.097545161008064   0.415734806151273  -0.415734806151273  -0.097545161008064   0.490392640201615  -0.277785116509801;
       0.191341716182545  -0.461939766255643   0.461939766255643  -0.191341716182545  -0.191341716182545   0.461939766255643  -0.461939766255643   0.191341716182545;
       0.097545161008064  -0.277785116509801   0.415734806151273  -0.490392640201615   0.490392640201615  -0.415734806151273   0.277785116509801  -0.097545161008064];
elseif (N == 8) & strcmp(transform_type, 'dst')==1 % hardcoded transform so that the PDE toolbox is not needed to generate it
    Tforward = [ 0.161229841765317   0.303012985114696   0.408248290463863   0.464242826880013   0.464242826880013   0.408248290463863   0.303012985114696   0.161229841765317;
       0.303012985114696   0.464242826880013   0.408248290463863   0.161229841765317  -0.161229841765317  -0.408248290463863  -0.464242826880013  -0.303012985114696;
       0.408248290463863   0.408248290463863                   0  -0.408248290463863  -0.408248290463863                   0   0.408248290463863   0.408248290463863;
       0.464242826880013   0.161229841765317  -0.408248290463863  -0.303012985114696   0.303012985114696   0.408248290463863  -0.161229841765317  -0.464242826880013;
       0.464242826880013  -0.161229841765317  -0.408248290463863   0.303012985114696   0.303012985114696  -0.408248290463863  -0.161229841765317   0.464242826880013;
       0.408248290463863  -0.408248290463863                   0   0.408248290463863  -0.408248290463863                   0   0.408248290463863  -0.408248290463863;
       0.303012985114696  -0.464242826880013   0.408248290463863  -0.161229841765317  -0.161229841765317   0.408248290463863  -0.464242826880013   0.303012985114696;
       0.161229841765317  -0.303012985114696   0.408248290463863  -0.464242826880013   0.464242826880013  -0.408248290463863   0.303012985114696  -0.161229841765317];
elseif (N == 7) & strcmp(transform_type, 'dct')==1 % hardcoded transform so that the signal processing toolbox is not needed to generate it
    Tforward =[ 0.377964473009227   0.377964473009227   0.377964473009227   0.377964473009227   0.377964473009227   0.377964473009227   0.377964473009227;
       0.521120889169602   0.417906505941275   0.231920613924330                   0  -0.231920613924330  -0.417906505941275  -0.521120889169602;
       0.481588117120063   0.118942442321354  -0.333269317528993  -0.534522483824849  -0.333269317528993   0.118942442321354   0.481588117120063;
       0.417906505941275  -0.231920613924330  -0.521120889169602                   0   0.521120889169602   0.231920613924330  -0.417906505941275;
       0.333269317528993  -0.481588117120063  -0.118942442321354   0.534522483824849  -0.118942442321354  -0.481588117120063   0.333269317528993;
       0.231920613924330  -0.521120889169602   0.417906505941275                   0  -0.417906505941275   0.521120889169602  -0.231920613924330;
       0.118942442321354  -0.333269317528993   0.481588117120063  -0.534522483824849   0.481588117120063  -0.333269317528993   0.118942442321354];   
elseif strcmp(transform_type, 'dct') == 1,
    Tforward    = dct(eye(N));
elseif strcmp(transform_type, 'dst') == 1,
    Tforward    = dst(eye(N));
elseif strcmp(transform_type, 'DCrand') == 1,
    x = randn(N); x(1:end,1) = 1; [Q,R] = qr(x); 
    if (Q(1) < 0), 
        Q = -Q; 
    end;
    Tforward = Q';
else %% a wavelet decomposition supported by 'wavedec'
    %%% Set periodic boundary conditions, to preserve bi-orthogonality
    dwtmode('per','nodisp');  
    
    Tforward = zeros(N,N);
    for i = 1:N
        Tforward(:,i)=wavedec(circshift([1 zeros(1,N-1)],[dec_levels i-1]), log2(N), transform_type);  %% construct transform matrix
    end
end

%%% Normalize the basis elements
Tforward = (Tforward' * diag(sqrt(1./sum(Tforward.^2,2))))'; 

%%% Compute the inverse transform matrix
Tinverse = inv(Tforward);

return;
```
