## Supplementary material for "FPM app: an open-source application for simple and intuitive Fourier ptychographic reconstruction": FPM app: BM3Ddenoising.m

```
function [denoised_object,phase_denoised] = BM3Ddenoising(object, phase, sigma)
% Function that denoises given complex object with the BM3D method
%   Inputs:
%       object - complex object
%       phase - object phase
%       sigma - denoise parameter
%   Output:
%       denoised_object - denoised complex object
%       phase_denoised - denoised object phase

f = waitbar(0,'BM3D denoising');

object = gather(object);
mina = min(min(abs(object)));
objectamp = abs(object)-mina;
maxa = max(max(objectamp));
objectamp = objectamp./maxa;    % Normalized amplitude

minp = min(min(phase));
objectphase = phase - minp;
maxp = max(max(objectphase));
objectphase = objectphase./maxp;    % Normalized phase

[~, oA] = BM3D(1, objectamp, sigma,'lc');    % denoising amplitude
try
    waitbar(0.5,f,'BM3D denoising');
catch
    f = waitbar(1,'finish');
end
[~, oP] = BM3D(1, objectphase, sigma,'lc');  % denoising phase


denoised_object = (oA.*maxa+mina).*exp(1i.*(oP.*maxp+minp));
phase_denoised = oP.*maxp+minp;
try
    waitbar(1,f,'finish');
catch
    f = waitbar(1,'finish');
end

try
    close(f)
end

end
```
