## Supplementary material for "FPM app: an open-source application for simple and intuitive Fourier ptychographic reconstruction": FPM app: GDUpdate_Multiplication_rank1.m

```
function [ O,P ] = GDUpdate_Multiplication_rank1(O,P,dpsi,Omax,cen,Ps,alpha,beta)
%GDUPDATE_MULTIPLICATION update estimate of O and P according to gradient
%descent method, where psi = O*P
%   Inputs:
%   O0: object estimate, n1xn2
%   P0: pupil function estimate: m1xm2
%   psi: update estimate field estimate
%   psi0: previous field estimate
%   cen: location of pupil function
%   alpha: gradient descent step size for O
%   betta: gradient descent step size for P
%   Ps: support constraint for P0, e.g. spatially confined probe or
%   objective with known NA
%   iters: # of iterations to run on updates
%
% last modified by Lei Tian,, 3/1/2014

% size of P, Np<=No
Np = size(P); %Np = Np(:);

% operator to put P at proper location at the O plane
n1 = cen-floor(Np/2);
n2 = n1+Np-1;
% operator to crop region of O from proper location at the O plane
downsampl = @(x) x(n1(1):n2(1),n1(2):n2(2));

O1 = downsampl(O);

O(n1(1):n2(1),n1(2):n2(2)) = O(n1(1):n2(1),n1(2):n2(2))...
    + 1/max(max(abs(P)))*abs(P).*conj(P).*dpsi./(abs(P).^2+alpha);
P = P+1/Omax*(abs(O1).*conj(O1)).*dpsi./(abs(O1).^2+beta).*Ps;
end
```
