## Supplementary material for "FPM app: an open-source application for simple and intuitive Fourier ptychographic reconstruction": FPM app: img_order.m

```
function [order] = img_order(LEDs, imageColOrder,  LEDsUsed, type, bck)
% Function that creates an array of image numbers in the order they are 
% going to be updating reconstructing object
%   Inputs:
%       LEDs - LED matrix used to collect images
%           0 in place where there is no LED
%           1 in place where there is LED
%           2 in place where there is central LED
%           Example:
%               LEDs = [0,1,1,1,0
%                       1,1,2,1,1
%                       0,1,1,1,0];
%       imageColOrder - matrix that shows in which order the images were 
%                       collected
%           1 - first image; 2 - second image; etc
%           Example:
%               imageColOrder = [0,1,2,3,0
%                                4,5,6,7,8
%                               0,9,10,11,0];
%       LEDsUSED - matrix that shows which images you want to reconstruct
%           1 - LED used in reconstruction
%           0 - LED not used in reconstruction
%           Example:
%               LEDsUsed = [0,1,1,1,0
%                           1,1,0,1,1
%                           0,1,1,1,0];
%       type - type of the reconstruction order
%           1 - by NA (from the lower to the higher ilumination angle)
%           2 - by intensity (from the brightest to the darkest image)
%       bck - background values of the images (array)
%   Output:
%       order - image reconstruction order (array)

% bck = gather(bck);
nImgs = sum(sum(LEDs))-1;
order = zeros(sum(sum(LEDsUsed)),1);

if type == '1'  % NA order
    spiral = createSpiral(LEDs,-1,0);
    n = 0;
    for m = 1:nImgs
        [y,x] = find(spiral == m);
        if LEDsUsed(y,x) == 1
            n = n+1;
            order(n) = imageColOrder(y,x);
        end
    end
end

if type == '2'  % intensity order
%     n = 0;
    for m = 1:sum(sum(LEDsUsed))
        a = find(bck == max(bck));
%         [y,x] = find(imageColOrder == a(1));
%         if LEDsUsed(y,x) == 1
%             n = n+1;
            order(m) = a(1);
%         end
        bck(a(1)) = -1;
    end
end

end
```
