## Supplementary material for "FPM app: an open-source application for simple and intuitive Fourier ptychographic reconstruction": FPM app: initialPupil.m

```
function [pupil] = initialPupil(pupil_0,type)
% Function that transforms initial pupil function shape in z direction
%   Inputs:
%       pupil_0 - initial pupil function
%       type - type of the transformed pupil
%           '1' - ones
%           '2' - tukey
%           '3' - gauss
%   Output:
%       pupil - transformed pupil function

type = '2';
[ny,nx] = size(pupil_0);
dx = sum(sum(pupil_0(ceil(end/2),:)));
dy = sum(sum(pupil_0(:,ceil(end/2))));
switch type
    case '1'
        pupil = pupil_0;
    case '2'
        xx = -nx/2:nx/2-1;
        [x,y] = meshgrid(xx,xx);
        calc_radius0 = sqrt(x.^2+y.^2);
        pupil = zeros(nx);
        tt0 = tukeywin(1000,0.25);
        tt1 = tt0(501:1000);
        tt2 = imresize(tt1, [round(dx/2*7/6) 1]);
        for m = 0:length(tt2)-1
           mask = zeros(nx);
           mask(calc_radius0>=m) = 1;
           mask(calc_radius0>m+1) = 0;
           pupil(mask == 1) = tt2(m+1);
        end
        pupil = imresize(pupil, [ny,nx]);
%         figure, imagesc(pupil,[min(min(pupil(pupil>0))) 1]); title('tukey')
%             colorbar
%         figure, imagesc(pupil_0); title('ones') colorbar
    case '3'
        pupila = gausswin(dy,0.25)*gausswin(dx,0.25)';
        pupila = padarray(pupila,[round((ny-dy)/2),round((nx-dx)/2)]);
        pupil = (pupila(1:ny,1:nx)).*pupil_0;
%         figure, imagesc(pupila,[min(min(pupila(pupila>0))) 1]); title('gauss')
%         colorbar
%         figure, imagesc(pupil_0); title('ones')
%         colorbar
end

end
```
