## Supplementary material for "FPM app: an open-source application for simple and intuitive Fourier ptychographic reconstruction": FPM app: InputImagesCrop.m

```
function [I2,ICO2] = InputImagesCrop(I,ICO,LEDsUsed,ROI)
% Function that crops input data to given ROI size and given number of
% images
%   Inputs:
%       I - vector of input images
%       ICO - image collecting order
%       LEDsUsed - LEDs used in reconstruction
%       ROI - Region Of Interest
%   Outputs:
%       I2 - I consisting of images specified by LEDsUsed matrix and in ROI
%            size
%       ICO2 - ICO consisting of images specified by LEDsUsed matrix

% Crop to ROI size
Io = I(ROI(2):ROI(2)+ROI(4)-1,ROI(1):ROI(1)+ROI(3)-1,:);

% Only Used LEDs and new ICO
[ny,nx,nImgs] = size(Io);
% I2 = uint8(zeros(ny,nx,sum(sum(LEDsUsed))));
ICO2 = zeros(size(ICO));
kk = 1;
for nn = 1:nImgs
    [ky,kx] = find(ICO == nn);
    if LEDsUsed(ky,kx) == 1
        I2(:,:,kk) = Io(:,:,nn);
        ICO2(ky,kx) = kk;
        kk = kk + 1;
    end
end

end
```
