## Supplementary material for "FPM app: an open-source application for simple and intuitive Fourier ptychographic reconstruction": FPM app: LEDsPosCorrection.m

```
if options.LEDcorrection ~= '0'
    
    switch options.LEDcorrection
        case '2'    % SA correction
%             O2 = gather(O); P2 = gather(P); I_mea2 = gather(I_mea);
            poscost = @(ss) sum(sum((abs(IFT(downsamp(O,ss).*P)).^2-I_mea).^2));
%             calbratetol = 0.1;
            optsanneal = saoptimset('Display','off','StallIterLimit',150);
            cen_correct = round(simulannealbnd(poscost,...
                cen,cen-mx,cen+mx,optsanneal));
            idx_Y(ledY,ledX) = cen0(1)-cen_correct(1);
            idx_X(ledY,ledX) = cen0(2)-cen_correct(2);
        case '3'    % GA correction
            O2 = gather(O); P2 = gather(P); I_mea2 = gather(I_mea);
            poscost = @(ss) sum(sum((abs(IFT(downsamp(O2,ss).*P2)).^2-I_mea2).^2));
            optsanneal = saoptimset('Display','off');
            cen_correct = ga(poscost,2,[],[],[],[],...
                cen-sp0/3,cen+sp0/3,[],[1,2],optsanneal);
            idx_Y(ledY,ledX) = cen0(1)-cen_correct(1);
            idx_X(ledY,ledX) = cen0(2)-cen_correct(2);
            
        case '4'    % Authors method [accurate] correction
            if options.useGPU == 1
                Err = gpuArray(zeros(2*mx+1));
            else
                Err = zeros(2*mx+1);
            end
            psi_mea = sqrt(I_mea); % amplitude of measured image
            for xx = -mx:mx
                for yy = -mx:mx
                    cen_tmp(2) = cen(2) + xx;
                    cen_tmp(1) = cen(1) + yy;
                    Psi0 = downsamp(O,cen_tmp).*P;
                    psi0 = abs(IFT(Psi0));   % estimated image (amplitude)        
                    Err(yy+mx+1,xx+mx+1) = rms(rms(psi0-psi_mea));
                end
            end
           [ym,xm] = find(Err==min(min(Err)));
           cen_corr = cen + [ym-mx-1,xm-mx-1];

           % current LED position in Fourier domain
           nYc = cen_corr(1) - cen0(1);
           nXc = cen_corr(2) - cen0(2);
           idx_X(ledY,ledX) = gather(nXc);
           idx_Y(ledY,ledX) = gather(nYc);
            
            
        case '5'  % Authors method [fast] correction     
            
            %         tic
            r = 2;  % subregion radius
            if options.useGPU == 1
                Err = gpuArray(zeros(2*mx+1));
            else
                Err = zeros(2*mx+1);
            end
            % stop criterion
            cond = 0;
            
            % initial LED position in Err matrix
            xcorr0 = mx+1;
            ycorr0 = mx+1;
            
            % corrected LED position in Err matrix
            xcorr = xcorr0;
            ycorr = ycorr0;
            % corrected LED position
            cen_corr = cen;
            
            % amplitude of measured image
            psi_mea = sqrt(I_mea);
            
            while cond ~=1
                for x = 1:2*r+1
                    for y = 1:2*r+1
                        xx = xcorr - r - 1 + x;
                        yy = ycorr - r - 1 + y;
                        if xx < 1 || yy < 1 || xx > 2*mx+1 || yy > 2*mx+1
                            cond = 1;
                            break
                        end
                        if Err(yy,xx)==0
                            cen_tmp(2) = cen_corr(2) - r - 1 + x;
                            cen_tmp(1) = cen_corr(1) - r - 1 + y;
                            Psi0 = downsamp(O,cen_tmp).*P;
                            psi0 = abs(IFT(Psi0));   % estimated image (amplitude)
                            
                            Err(yy,xx) = rms(rms(psi0-psi_mea));
                        end
                    end
                    if cond == 1
                        break
                    end
                end
                [ym,xm] = find(Err==min(Err(Err>0)));
                cen_corr2 = gather(cen + [ym,xm] - [ycorr0,xcorr0]);
                
                if cen_corr == cen_corr2
                    cond = 1;
                    % figure(9); imagesc(Err, [mini max(max(Err))]);
                    % figure(10); imagesc(log(1+abs((Obj))));
                    % pause(0.1)
                end
                
                cen_corr = cen_corr2;
                xcorr = xm;
                ycorr = ym;
            end
            
            % current LED position in Fourier domain
            nYc = cen_corr(1) - cen0(1);
            nXc = cen_corr(2) - cen0(2);
            idx_X(ledY,ledX) = nXc;
            idx_Y(ledY,ledX) = nYc;
    end
    
end
```
