## Supplementary material for "FPM app: an open-source application for simple and intuitive Fourier ptychographic reconstruction": FPM app: LoadImages.m

```
function [I,imageList] = LoadImages(loadDirectory)
% Function that is loading input images
%   Inputs:
%       loadDirectory - directory where are input FPM images
%   Outputs:
%       I - loaded images (3D matrix)   
%       imageList - list of loaded images

imageList = dir(loadDirectory);
imageList(2) = []; imageList(1) = [];
a = string(zeros(length(imageList),1));
for k = 1:length(imageList)
    a(k) = imageList(k).name;
end
imageList = sort_nat(a);
clear k a;

% [n1,n2] = size(imread(strcat(loadDirectory,'/',imageList(1))));
% I = zeros(n1,n2,length(imageList),'uint8');
I = imread(strcat(loadDirectory,'/',imageList(1)));
I(:,:,2:length(imageList)) = 0;
f = waitbar(0,strcat('Loading images ... ',imageList(1)));
s = length(imageList);
for mm = 2:s
    I(:,:,mm) = imread(strcat(loadDirectory,'/',imageList(mm)));
    progress = mm/s;
    try
        waitbar(progress,f,strcat('Loading images ... ',imageList(mm)));
    catch
        f = waitbar(progress,strcat('Loading images ... ',imageList(mm)));
    end
end
try
    close(f)
end
end
```
