## Supplementary material for "FPM app: an open-source application for simple and intuitive Fourier ptychographic reconstruction": FPM app: Proj_Fourier_v2.m

```
function [ psi] = Proj_Fourier_v2( psi0, Imea, Iest, c, F )
%PROJ_FOURIER projection based on intensity measurement in the fourier
%domain, replacing the amplitude of the Fourier transform by measured
%amplitude, sqrt(I)
% last modified by Lei Tian,, 3/1/2014

[n1,n2,r] = size(psi0);

if r == 1
    psi = F(sqrt(Imea/c).*psi0./(sqrt(Iest)+eps));
else
    psi = zeros(n1,n2,r);
    for m = 1:r
        psi(:,:,m) = F(sqrt(Imea).*psi0(:,:,m)./sqrt(Iest+eps));
    end
end

end
```
