## Supplementary material for "FPM app: an open-source application for simple and intuitive Fourier ptychographic reconstruction": FPM app: license.txt

Copyright (c) 2008, Douglas M. Schwarz
All rights reserved.

Redistribution and use in source and binary forms, with or without
modification, are permitted provided that the following conditions are
met:

    * Redistributions of source code must retain the above copyright
      notice, this list of conditions and the following disclaimer.
    * Redistributions in binary form must reproduce the above copyright
      notice, this list of conditions and the following disclaimer in
      the documentation and/or other materials provided with the distribution

THIS SOFTWARE IS PROVIDED BY THE COPYRIGHT HOLDERS AND CONTRIBUTORS "AS IS"
AND ANY EXPRESS OR IMPLIED WARRANTIES, INCLUDING, BUT NOT LIMITED TO, THE
IMPLIED WARRANTIES OF MERCHANTABILITY AND FITNESS FOR A PARTICULAR PURPOSE
ARE DISCLAIMED. IN NO EVENT SHALL THE COPYRIGHT OWNER OR CONTRIBUTORS BE
LIABLE FOR ANY DIRECT, INDIRECT, INCIDENTAL, SPECIAL, EXEMPLARY, OR
CONSEQUENTIAL DAMAGES (INCLUDING, BUT NOT LIMITED TO, PROCUREMENT OF
SUBSTITUTE GOODS OR SERVICES; LOSS OF USE, DATA, OR PROFITS; OR BUSINESS
INTERRUPTION) HOWEVER CAUSED AND ON ANY THEORY OF LIABILITY, WHETHER IN
CONTRACT, STRICT LIABILITY, OR TORT (INCLUDING NEGLIGENCE OR OTHERWISE)
ARISING IN ANY WAY OUT OF THE USE OF THIS SOFTWARE, EVEN IF ADVISED OF THE
POSSIBILITY OF SUCH DAMAGE.
