## Supplementary material for "FPM app: an open-source application for simple and intuitive Fourier ptychographic reconstruction": FPM app: sort_nat.m

```
function [cs,index] = sort_nat(c,mode)
%sort_nat: Natural order sort of cell array of strings.
% usage:  [S,INDEX] = sort_nat(C)
%
% where,
%    C is a cell array (vector) of strings to be sorted.
%    S is C, sorted in natural order.
%    INDEX is the sort order such that S = C(INDEX);
%
% Natural order sorting sorts strings containing digits in a way such that
% the numerical value of the digits is taken into account.  It is
% especially useful for sorting file names containing index numbers with
% different numbers of digits.  Often, people will use leading zeros to get
% the right sort order, but with this function you don't have to do that.
% For example, if C = {'file1.txt','file2.txt','file10.txt'}, a normal sort
% will give you
%
%       {'file1.txt'  'file10.txt'  'file2.txt'}
%
% whereas, sort_nat will give you
%
%       {'file1.txt'  'file2.txt'  'file10.txt'}
%
% See also: sort

% Version: 1.4, 22 January 2011
% Author:  Douglas M. Schwarz
% Email:   dmschwarz=ieee*org, dmschwarz=urgrad*rochester*edu
% Real_email = regexprep(Email,{'=','*'},{'@','.'})


% Set default value for mode if necessary.
if nargin < 2
	mode = 'ascend';
end

% Make sure mode is either 'ascend' or 'descend'.
modes = strcmpi(mode,{'ascend','descend'});
is_descend = modes(2);
if ~any(modes)
	error('sort_nat:sortDirection',...
		'sorting direction must be ''ascend'' or ''descend''.')
end

% Replace runs of digits with '0'.
c2 = regexprep(c,'\d+','0');

% Compute char version of c2 and locations of zeros.
s1 = char(c2);
z = s1 == '0';

% Extract the runs of digits and their start and end indices.
[digruns,first,last] = regexp(c,'\d+','match','start','end');

% Create matrix of numerical values of runs of digits and a matrix of the
% number of digits in each run.
num_str = length(c);
max_len = size(s1,2);
num_val = NaN(num_str,max_len);
num_dig = NaN(num_str,max_len);
for i = 1:num_str
	num_val(i,z(i,:)) = sscanf(sprintf('%s ',digruns{i}{:}),'%f');
	num_dig(i,z(i,:)) = last{i} - first{i} + 1;
end

% Find columns that have at least one non-NaN.  Make sure activecols is a
% 1-by-n vector even if n = 0.
activecols = reshape(find(~all(isnan(num_val))),1,[]);
n = length(activecols);

% Compute which columns in the composite matrix get the numbers.
numcols = activecols + (1:2:2*n);

% Compute which columns in the composite matrix get the number of digits.
ndigcols = numcols + 1;

% Compute which columns in the composite matrix get chars.
charcols = true(1,max_len + 2*n);
charcols(numcols) = false;
charcols(ndigcols) = false;

% Create and fill composite matrix, comp.
comp = zeros(num_str,max_len + 2*n);
comp(:,charcols) = double(s1);
comp(:,numcols) = num_val(:,activecols);
comp(:,ndigcols) = num_dig(:,activecols);

% Sort rows of composite matrix and use index to sort c in ascending or
% descending order, depending on mode.
[unused,index] = sortrows(comp);
if is_descend
	index = index(end:-1:1);
end
index = reshape(index,size(c));
cs = c(index);
```
