## Supplementary material for "FPM app: an open-source application for simple and intuitive Fourier ptychographic reconstruction": FPM app: createSpiral.m

```
function [spiral] = createSpiral(LEDs,type,secLED)
%Function that creates spiral in matrix
%   Inputs:
%       LEDs - LED matrix
%           0 in place where there is no LED
%           1 in place where there is LED
%           2 in place where there is central LED
%           Example:
%               LEDs = [0,1,1,1,0
%                       1,1,2,1,1
%                       0,1,1,1,0];
%       type - type of the spiral
%           type = -1 - anticlockwise
%           type = 1 - clockwise
%       secLED - which LED is second
%           secLED = 0 - second LED - down
%           secLED = 1 - second LED - right
%           secLED = 2 - second LED - up
%           secLED = 3 - second LED - left
%   Output:
%       spiral - matrix that contains created spiral
%           0 in place where there is no LED
%           1 in place where there is first LED
%           2 in place where there is second LED
%           etc.
%           Example:
%               spiral = [0,7,8,9,0
%                        11,6,1,2,10
%                         0,5,4,3,0];

%%
maxCounter = sum(sum(LEDs))-1; % number of LEDs
[ny,nx] = size(LEDs);
ns = max(ny,nx);
LEDs0 = padarray(LEDs,[ns,ns]);
[ny0,nx0] = size(LEDs0);

% LED order but with no LEDs positions (0) treated as LED positions
order2 = zeros(ny0,nx0);    

[ym,xm] = find(LEDs0 == 2); % first LED position
order2(ym,xm) = 1;
order = order2; % LED order
% direct = 0; % 0-down, 1-right, 2-up, 3-left

counter = 2;    % next LED number in order matrix
counter2 = 2;   % next LED number in order2 matrix
% tt = 0;
switch secLED   % second LED position and direction
    case 0
        ym = ym+1;
        tmp_direct = 0;
    case 1
        xm = xm+1;
        tmp_direct = 1;
    case 2
        ym = ym-1;
        tmp_direct = 2;
    case 3
        xm = xm-1;
        tmp_direct = 3;
end

for m = 1:ny0*nx0-1
    direct = tmp_direct;    % direction where there is next LED
    switch direct
        case 0  % going down
            order2(ym,xm) = counter2;
            if LEDs0(ym,xm) == 1 % checking if this LED exist
                order(ym,xm) = counter;
                counter = counter + 1;
                if counter>maxCounter   % if it was last LED
                    break
                end
            end
            if order2(ym,xm-1*type)==0   % checking if the LED on the right/left is not in order2
                tmp_direct = 1; % if is not (on the right) than going right
                if type == 1    % clockwise
                    tmp_direct = 3; % if is not (on the left) than going left
                end
                xm = xm-1*type; % next LED x position 
            else    % if it is going futher down
                ym = ym+1;  % next LED y position 
            end
        case 1  % going right
            order2(ym,xm) = counter2;
            if LEDs0(ym,xm) == 1
                order(ym,xm) = counter;
                counter = counter + 1;
                if counter>maxCounter
                    break
                end
            end
            if order2(ym+1*type,xm)==0
                tmp_direct = 2;
                if type == 1    % clockwise
                    tmp_direct = 0;
                end
                ym = ym+1*type;
            else
                xm = xm+1;
            end
        case 2  % going up
            order2(ym,xm) = counter2;
            if LEDs0(ym,xm) == 1
                order(ym,xm) = counter;
                counter = counter + 1;
                if counter>maxCounter
                    break
                end
            end
            if order2(ym,xm+1*type)==0
                tmp_direct = 3;
                if type == 1    % clockwise
                    tmp_direct = 1;
                end
                xm = xm+1*type;
            else
                ym = ym-1;
            end
        case 3  % going left
            order2(ym,xm) = counter2;
            if LEDs0(ym,xm) == 1
                order(ym,xm) = counter;
                counter = counter + 1;
                if counter>maxCounter
                    break
                end
            end
            if order2(ym-1*type,xm)==0
                tmp_direct = 0;
                if type == 1    % clockwise
                    tmp_direct = 2;
                end
                ym = ym-1*type;
            else
                xm = xm-1;
            end
    end
%     figure(1);
%     imagesc(order)
%     figure(2);
%     imagesc(order2)
%     pause(0.05)
    counter2 = counter2+1;

end

spiral = order(ns+1:ns+ny,ns+1:ns+nx);
% figure; imagesc(spiral)

end
```
