## Supplementary material for "FPM app: an open-source application for simple and intuitive Fourier ptychographic reconstruction": FPM app: CutPathName.m

```
function [path_out] = CutPathName(path,nos)
% Function that cuts the path name to the last nos number parts of it
%   Example:
%       nos = 2;
%       path = 'C:/Users/User1/Documents/MATLAB/measurments/USAF target/data'
%       path_out = '/USAF target/data'
if nargin < 2
    nos = 2;
end

    tmp = uint8(path);
    p = 0;
    path_out = [];
    for m = length(tmp):-1:1
        if tmp(m) == 47
            p = p+1;
        end
        sgn = tmp(m);
        path_out = [sgn,path_out];        
        if p == nos
            break
        end        
    end
    path_out = char(path_out);
    
end
```
