## Supplementary material for "FPM app: an open-source application for simple and intuitive Fourier ptychographic reconstruction": FPM app: ErrorMessages.m

```
function [NoError] = ErrorMessages(options, LEDs, I, ROI, LEDsUsed)
% Function that checks at the beginning of the reconstruction if there s no
% error. If there is an error, it returns corresponding message
%   Inputs:
%       options - reconstruction options
%           options.alpha - regularization parameter for object reconstruction
%           options.beta - regularization parameter for pupil reconstruction
%           options.maxIter - maximum number of iterations
%           options.algorithm - select reconstruction algorithm
%               1 - Quasi-Newton algorithm
%               2 - Gerchberg-Saxton
%           options.recorder - reconstruction order
%               1 - from lowest to highest NA
%               2 - from the lightest to the darkest images
%           options.LEDcorrection - LED position correction
%               1 - Angle Self-Calibration
%               2 - simulated annealing
%               3 - genetic algorithm
%           options.initialPupil - initial pupil
%               1 - ones
%               2 - tukey window
%               3 - gauss window
%           options.useGPU
%               1 - use GPU acceleration
%               0 - don't use
%       LEDs - LED matrix used to collect images
%           0 in place where there is no LED
%           1 in place where there is LED
%           2 in place where there is central LED
%           Example:
%               LEDs = [0,1,1,1,0
%                       1,1,2,1,1
%                       0,1,1,1,0];
%       I - input FPM images
%       ROI - Region Of Interest
%           ROI = [x0,y0,xSize,ySize];
%       LEDsUSED - matrix that shows which images you want to reconstruct
%           1 - LED used in reconstruction
%           0 - LED not used in reconstruction
%           Example:
%               LEDsUsed = [0,1,1,1,0
%                           1,1,0,1,1
%                           0,1,1,1,0];
%   Outputs:
%       NoError
%           1 - there is no error
%           0 - there is some error


tmp1 = 1;
tmp2 = 1;

% Check if there is GPU and enough available memory
if ~isempty(options) && ~isempty(ROI) 
    if options.useGPU == 1  
        try tt = gpuDevice;
            
        catch
            tmp1 = -1;   % No GPU
        end
        if tmp1 == 1
            Isize = sum(sum(LEDsUsed));
            Isize = 8*Isize*ROI(3)*ROI(4);
            if tt.AvailableMemory - Isize < 1e9   
                tmp1 = 0;    % No enough GPU memory
            end
        end
    end
    switch tmp1
        case 0
            f1 = errordlg(['Not enough GPU memory. AvailableMemory = ',...
                num2str(tt.AvailableMemory), '; Requested memory = ', ...
                num2str(Isize+1e9), '; Try smaller ROI']);
        case -1
            f1 = errordlg('No NVidia GPU detected. Try to actualize CUDA driver.');
            tmp1 = 0;

    end
end

if ~isempty(LEDs) && ~isempty(I)    % Check if there is LEDs matrix error
    [~,~,n] = size(I);
    m = sum(sum(LEDs))-1;
    t = max(max(LEDs));
    l = find(LEDs == t);
    if t == 1
        tmp2 = 0;
        f2 = errordlg('There is no central LED selected (it should have value 2)');
    elseif t ~= 2
        tmp2 = 0;
        f2 = errordlg('Central LED must have value 2');
    else
        if length(l) > 1
            tmp2 = 0;
            f2 = errordlg('There can be only one central LED');
        else
            if m ~= n
                tmp2 = 0;
                f2 = errordlg('Number of LEDs is not equal to the number of images');
            end
        end
    end
end


NoError = tmp1*tmp2;
end
```
