## Supplementary material for "FPM app: an open-source application for simple and intuitive Fourier ptychographic reconstruction": FPM app: Fourier_position.m

```
function [imgSize, overlap, synthNA] = Fourier_position(amplitude,phase,systemSetup,LEDs)
% Function that returns the positions of images centres in Fourier domain
% along with generated image size, overlap of the system and synthetic 
% numerical aperture
%   Inputs:
%       amplitude - amplitude of the synthetic object (just to get it size)
%       systemSetup - system setup
%           systemSetup.NA - NA
%           systemSetup.lambda - wavelength (um)
%           systemSetup.magnification - magnification
%           systemSetup.LEDspacing - spacing between neighbour LEDs in LED
%                                    matrix (mm)
%           systemSetup.LEDheight - distance between LED matrix and a 
%                                   sample (mm)
%           systemSetup.camPizSize - camera pixel size (um)
%       LEDs - LED matrix
%           0 in place where there is no LED
%           1 in place where there is LED
%           2 in place where there is central LED
%           Example:
%               LEDs = [0,1,1,1,0
%                       1,1,2,1,1
%                       0,1,1,1,0];
%   Outputs
%       imgSize - size of the low resolution imaages that will be created
%       overlap - overlap in Fourier space between neighbouring LEDs 
%                 (calculated for the central LEDs)
%       synthNA - synthetic NA (objective NA + max(illumination NA))
%       figure(12) - visualize overlap in fourier domain

%% Synthetic NA
[ny0,nx0] = size(amplitude);

lambda = systemSetup.lambda;
mag = systemSetup.magnification;
pixSizeCam = systemSetup.camPizSize;    % pixel size of the camera
pixSizeObj = pixSizeCam/mag;    % pixel size at the object plane
NA = systemSetup.NA;
um_m = NA/lambda;   % frequencies transported through the system

if max(max(LEDs)) == 1  % central LED position
    [cledY,cledX] = size(LEDs);
    cledY = ceil(cledY/2); cledX = ceil(cledX/2);
else
    [cledY,cledX] = find(LEDs==2);
end
LEDs(cledY,cledX) = 1;
[ys,xs] = size(LEDs);
% LEDnr = sum(sum(LEDs));
%centerLED = [8,8];
LEDspacing = systemSetup.LEDspacing; % spacing between adjacent LEDs
LEDheight = systemSetup.LEDheight; % distance bewteen the LED matrix and the sample
xx = 1:xs; xx = (xx - cledX).*LEDspacing;
yy = 1:ys; yy = (yy - cledY).*LEDspacing;
[LEDsPosX,LEDsPosY] = meshgrid(xx,yy);  % LED spacing in X and Y

dist = sqrt(LEDsPosX.^2+LEDsPosY.^2+LEDheight.^2);    % distances LED-sample
% sin(angle between LEDs and line perpendicular to the sample)
sin_thetaX = LEDsPosX./dist;  
sin_thetaY = LEDsPosY./dist;

illuminationNA = sqrt(sin_thetaX.^2+sin_thetaY.^2);
illuminationNA = illuminationNA.*LEDs;
% frequencies transported through the synthetic system
um_p = max(max(illuminationNA))/lambda+um_m;
synthNA = num2str(um_p*lambda);

%% Calculating generated images size

% How many times object resolution is bigger than collected image 
% resolution
% This value is set to be the same size as the value determining how many
% times the reconstructed image resolution will be bigger than collected 
% image resolution
par = 1;    

nx1 = nx0+1;
while nx1>nx0
    par = par + 1;
    nxIm = round(nx0./par./2).*2;
    FoVx = nxIm.*pixSizeObj;
    dux = 1./FoVx;
    nx1 = round(2.*um_p./dux).*2;
end
% Set par as constant value:
% par = 3;

nyIm = round(ny0/par/2)*2;
nxIm = round(nx0/par/2)*2;
imgSize = strcat(num2str(nxIm),'x',num2str(nyIm));

%% 
% FOV in x and y direction [um]
FoVx = nxIm*pixSizeObj;
FoVy = nyIm*pixSizeObj;
% sampling size at Fourier plane set by the image size (FoV)
% sampling size at Fourier plane is always = 1/FoV
if mod(nxIm,2) == 1
    dux = 1/pixSizeObj/(ROI(3)-1);
else
    dux = 1/FoVx;
end
if mod(nyIm,2) == 1
    duy = 1/pixSizeObj/(ROI(4)-1);
else
    duy = 1/FoVy;
end

m = 1:nxIm;
n = 1:nyIm;
[mm,nn] = meshgrid(m-round((nxIm+1)/2),n-round((nyIm+1)/2));
if nxIm>nyIm
    nn = nn.*max(max(mm))./max(max(nn));
else
    mm = mm.*max(max(nn))./max(max(mm));
end
ridx = sqrt(mm.^2+nn.^2);
um_idx = um_m/min(dux,duy);
% % assume a circular initial pupil function due to finite NA
pupil0 = double(ridx<um_idx);

% corresponding spatial freq for each LEDs
xFreq = sin_thetaX/lambda;
yFreq = sin_thetaY/lambda;
% spatial freq index for each plane wave relative to the center
idx_Y = round(yFreq/duy).*LEDs;
idx_X = round(xFreq/dux).*LEDs;

%% overlap
matrix = zeros(ny0,nx0);
yy = ceil(ny0/2)-ceil(nyIm)/2:ceil(ny0/2)+ceil(nyIm)/2-1;
xx = ceil(nx0/2)-ceil(nxIm)/2:ceil(nx0/2)+ceil(nxIm)/2-1; 
matrix(yy,xx) = pupil0;

overlap = 'No overlap';
ove = 0;
for m = -1:1
    for n = -1:1
        if cledY + m <= ys && cledY + m >= 1 && cledX + n <= xs && cledX + n >= 1
            if LEDs(cledY + m,cledX + n) == 1 && abs(m) + abs(n) == 1
                ove = ove+1;
                y = idx_Y(cledY + m,cledX + n);
                x = idx_X(cledY + m,cledX + n);
                thisover = matrix;
                thisover(yy+y,xx+x) = thisover(yy+y,xx+x) + pupil0;
                A(ove) = sum(sum(thisover(thisover == 2)))/2;
            end
        end
    end
end

if ove > 0
    overlap = strcat(num2str(round(mean(A)/sum(sum(matrix))*100)),'%');
end

%% Displaying
object = amplitude.*exp(1i.* phase);
radius = ceil(sum(pupil0(ceil(end/2),:))/2);
idx_Y2 = round(idx_Y*nx0/ny0);
idx_Y2 = reshape(idx_Y2,[],1);
idx_X2 = reshape(idx_X,[],1);
idx_Y2(LEDs==0)=[];
idx_X2(LEDs==0)=[];
c = [idx_X2,idx_Y2] + ceil((nx0+1)/2);
r(1:length(c)) = radius;
object = imresize(object,[nx0,nx0]);
figure(12); 
set(gcf,'Name','Positions of the image centers')
set(gcf,'NumberTitle','off')
imagesc(log(1+abs(fftshift(fft2(object)))));
colormap gray; title('Positions of the image centers in Fourier domain');
xlabel X; ylabel('Y scaled to the X size');
viscircles(c,r,'LineWidth',0.2);
r(1:length(c)) = 1;
viscircles(c,r,'Color','b','LineWidth',3);
um_pMax = 1/lambda+um_m;
um_pMax_idx = um_pMax/min(dux,duy);

m = 1:nx0;
n = 1:ny0;
[mm,nn] = meshgrid(m-round((nx0+1)/2),n-round((ny0+1)/2));
if nxIm>nyIm
    nn = nn.*max(max(mm))./max(max(nn));
else
    mm = mm.*max(max(nn))./max(max(mm));
end
ridx = sqrt(mm.^2+nn.^2);
pupil0_2 = (ridx<um_pMax_idx);
rMax = ceil(sum(pupil0_2(ceil(end/2),:))/2);
viscircles([ceil((nx0+1)/2),ceil((nx0+1)/2)],radius,'Color','k','LineStyle','--');
viscircles([ceil((nx0+1)/2),ceil((nx0+1)/2)],rMax,'Color','g','LineStyle','--');

end
```
