## Supplementary material for "FPM app: an open-source application for simple and intuitive Fourier ptychographic reconstruction": FPM app: ImageGenerationWork.m

```
function imagesIn = ImageGenerationWork(output_path,amplitude,phase,systemSetup,LEDs,showIm)
% Function that generates synthetic FPM data
%   Inputs:
%       output_path - directory where the created images will be saved
%                     (optional)
%           0 - no directory
%       amplitude - synthetic object amplitude
%       phase - synthetic object phase
%       systemSetup - system setup
%           systemSetup.NA - NA
%           systemSetup.lambda - wavelength (um)
%           systemSetup.magnification - magnification
%           systemSetup.LEDspacing - spacing between neighbour LEDs in LED
%                                    matrix (mm)
%           systemSetup.LEDheight - distance between LED matrix and a 
%                                   sample (mm)
%           systemSetup.camPizSize - camera pixel size (um)
%       LEDs - LED matrix
%           0 in place where there is no LED
%           1 in place where there is LED
%           2 in place where there is central LED
%           Example:
%               LEDs = [0,1,1,1,0
%                       1,1,2,1,1
%                       0,1,1,1,0];
%       showIm - show images during generation
%           1 - yes; 0 - no
%   Output:
%       imagesIn - generated synthetic low resolution FPM images

%%
tic
% Displaying amplitude and phase
figure(11);
set(gcf,'Name','Simulated object');
set(gcf,'NumberTitle','off');
subplot(1,2,1), imagesc(amplitude);title('Simulated object amplitude');
colormap gray; axis image
subplot(1,2,2), imagesc(phase);title('Simulated object phase');
colormap gray; axis image

% synthetic object
object = amplitude.*exp(1i.* phase);
[ny0,nx0] = size(object);

lambda = systemSetup.lambda;
mag = systemSetup.magnification;
pixSizeCam = systemSetup.camPizSize;    % pixel size of the camera
pixSizeObj = pixSizeCam/mag;    % pixel size at the object plane
NA = systemSetup.NA;
um_m = NA/lambda;   % frequencies transported through the system

if max(max(LEDs)) == 1  % central LED position
    [cledY,cledX] = size(LEDs);
    cledY = ceil(cledY/2); cledX = ceil(cledX/2);
else
    [cledY,cledX] = find(LEDs==2);
end

LEDs(cledY,cledX) = 1;
[ys,xs] = size(LEDs);
LEDnr = sum(sum(LEDs)); % number of LEDs
%centerLED = [8,8];
LEDspacing = systemSetup.LEDspacing; % spacing between adjacent LEDs
LEDheight = systemSetup.LEDheight; % distance bewteen the LED matrix and the sample
xx = 1:xs; xx = (xx - cledX).*LEDspacing;
yy = 1:ys; yy = (yy - cledY).*LEDspacing;
[LEDsPosX,LEDsPosY] = meshgrid(xx,yy);  % LED spacing in X and Y

dist = sqrt(LEDsPosX.^2+LEDsPosY.^2+LEDheight.^2);    % distance LEDs-sample
% sin(angle between LEDs and line perpendicular to the sample)
sin_thetaX = LEDsPosX./dist;
sin_thetaY = LEDsPosY./dist;

illuminationNA = sqrt(sin_thetaX.^2+sin_thetaY.^2);
illuminationNA = illuminationNA.*LEDs;
% frequencies transported through the synthetic system
um_p = max(max(illuminationNA))/lambda+um_m;

% How many times object resolution is bigger than collected image 
% resolution
% This value is set to be the same size as the value determining how many
% times the reconstructed image resolution will be bigger than collected 
% image resolution
par = 1;

nx1 = nx0+1;
while nx1>nx0
    par = par + 1;
    nxIm = round(nx0./par./2).*2;
    FoVx = nxIm.*pixSizeObj;
    dux = 1./FoVx;
    nx1 = round(2.*um_p./dux).*2;
end
% Set par as constant value:
% par = 3;

nyIm = round(ny0/par/2)*2;
nxIm = round(nx0/par/2)*2;

FoVx = nxIm*pixSizeObj;
FoVy = nyIm*pixSizeObj;
% sampling size at Fourier plane set by the image size (FoV)
% sampling size at Fourier plane is always = 1/FoV
if mod(nxIm,2) == 1
    dux = 1/pixSizeObj/(ROI(3)-1);
else
    dux = 1/FoVx;
end
if mod(nyIm,2) == 1
    duy = 1/pixSizeObj/(ROI(4)-1);
else
    duy = 1/FoVy;
end

m = 1:nxIm;
n = 1:nyIm;
[mm,nn] = meshgrid(m-round((nxIm+1)/2),n-round((nyIm+1)/2));
if nxIm>nyIm
    nn = nn.*max(max(mm))./max(max(nn));
else
    mm = mm.*max(max(nn))./max(max(mm));
end
ridx = sqrt(mm.^2+nn.^2);
um_idx = um_m/min(dux,duy);

pupil0 = double(ridx<um_idx);

% corresponding spatial freq for each LEDs
xFreq = sin_thetaX/lambda;
yFreq = sin_thetaY/lambda;
% spatial freq index for each plane wave relative to the center
idx_Y = round(yFreq/duy);
idx_X = round(xFreq/dux);

% figure(13); imagesc(pupil0); title('Pupil function'); colormap gray;

% Adding misaligment noise
% load('noise6.mat')
% idx_X = idx_X + round(2*noisex);
% idx_Y = idx_Y + round(2*noisey);

%% Generating images

imagesIn = zeros(nyIm, nxIm, LEDnr); % the final low resolution image sequence
objectFT = fftshift(fft2(object));
tt = 0;
if showIm == 0
    f = waitbar(0,'Image no...1');
end
radius = ceil(sum(pupil0(ceil(end/2),:))/2);
for y = 1:ys
    for x = 1:xs   
        kxc = idx_X(y,x) + ceil((nx0+1)/2);
        kyc = idx_Y(y,x) + ceil((ny0+1)/2);
        kyl=round(kyc-(nyIm-1)/2);kyh=round(kyc+(nyIm-1)/2);
        kxl=round(kxc-(nxIm-1)/2);kxh=round(kxc+(nxIm-1)/2);
        if LEDs(y,x) == 1
            tt = tt+1;
            imSeqLowFT = (1/par)^2 *objectFT(kyl:kyh,kxl:kxh).*pupil0;
            imagesIn(:,:,tt) = abs(ifft2(ifftshift(imSeqLowFT))).^2;

%             % adding synthetic (20%) noise
%             noim = max(max(imagesIn(:,:,tt)));
%             imagesIn(:,:,tt) = imagesIn(:,:,tt) + noim*0.2*rand(size(imagesIn(:,:,tt)));
%             % 
            
%             if x == cledX && y == cledY
%                 centerImage = tt;
%             end
%             tmp = zeros(size(object));
%             tmp(kyl:kyh,kxl:kxh) = tmp(kyl:kyh,kxl:kxh) + w_NA;
            if showIm == 1
                figure(14); 
                set(gcf,'Name','Generated images');
                set(gcf,'NumberTitle','off');
                subplot(1,2,1); imagesc(imagesIn(:,:,tt));title(strcat('image number: ', num2str(tt)));          
                subplot(1,2,2); imagesc(log(1+abs(objectFT)));title('Image position in Fourier space');
                viscircles([kxc,kyc], radius);
                colormap gray
                pause(0.1);
            else
                f = ActualizeWaitbar(tt,LEDnr,f);
            end
            if output_path ~= 0
                output_path = ChangeSlash(output_path);
                fileName = strcat(output_path, '/Img_',num2str(tt),'.png');
                imwrite(uint16(imagesIn(:,:,tt)),fileName);
            end
        end
    end
end
imagesIn = uint16(imagesIn);
try
    close(14)
end
try
    close(f)
end
% figure(15);
% imagesc(imagesIn(:,:,centerImage));
% title('Generated image (central diode)'); colormap gray;
toc
end

function f = ActualizeWaitbar(imgnr,nrofimgs,f)
if imgnr < nrofimgs
    progress = (imgnr)/nrofimgs;
    try
        waitbar(progress,f,strcat('Image no...',num2str(imgnr+1)));
    catch
        f = waitbar(progress,strcat('Image no...',num2str(imgnr+1)));
    end
else
    try
        waitbar(1,f,'finish');
    catch
        f = waitbar(1,'finish');
    end
end
end
```
