## Supplementary material for "FPM app: an open-source application for simple and intuitive Fourier ptychographic reconstruction": FPM app: ImgColOrder.m

```
function [imageColOrder,displayingOrder,nx,ny,colorMap] = ImgColOrder(LEDs, imStart, type)
% Function that generates image collecting order
%   Inputs:
%       LEDs - LED matrix
%           0 in place where there is no LED
%           1 in place where there is LED
%           2 in place where there is central LED
%           or LEDs = image collecting order; if just want to have nice
%           displaying matrix
%           Example:
%               LEDs = [0,1,1,1,0
%                       1,1,2,1,1
%                       0,1,1,1,0];
%       imStart - first collected image
%           imStart = 1 - top left
%           imStart = 2 - top right
%           imStart = 3 - bottom left
%           imStart = 4 - bottom right
%       type - collecting order type
%           type = 0 - just want to display order
%           type = 1 - line by line
%           type = 2 - row by row
%           type = 3 - snake line by line
%           type = 4 - snake row by row
%   Outputs:
%       imageColOrder - matrix of image collecting order
%           0 - no image
%           1 - first image
%           2 - second image
%           etc.
%           Example:
%               imageColOrder = [0,1,2,3,0
%                                4,5,6,7,8
%                               0,9,10,11,0];
%       displayingOrder - imageColOrder in displaying mode
%       nx,ny - size of the LEDs matrix
%       colorMap - color map of the displayingOrder
%   How to display it nicely:
%       imagesc(0.5:ny+1,0.5:nx+1,displayingOrder); colormap(colorMap)

[ny,nx] = size(LEDs);
if type~=0
    LEDs(LEDs==2) = 1;

    imageColOrder = zeros(ny,nx);
    imageColOrder = imageColOrder - 10;
    
%     LEDs no in x and y directions
    my = 1:ny;
    mx = 1:nx;
    
    if imStart == 2 || imStart == 4
        mx = nx:-1:1; 
    end
    if imStart == 3 || imStart == 4
        my = ny:-1:1; 
    end
    
    if type == 2 || type == 4   % swap x and y LED no to move row by row
        tmp = mx;
        mx = my;
        my = tmp;
    end
    
    nr = 0;     % LED number
    par = 0;    % parametr to make snake turn
    for ty = my
        par = par + 1;  
        for tx = mx
            if type == 3 || type == 4   % snake order
                 if mod(par,2) == 0
                     txs = abs(tx - nx - 1);
                     if type == 4
                         txs = abs(tx - ny - 1);
                     end
                 else   
                     txs = tx;
                 end
            else    % line by line/row by row order
                txs = tx;
            end
            if type == 1 || type == 3
                if LEDs(ty,txs) == 1
                    nr = nr + 1;
                    imageColOrder(ty,txs) = nr;
                end
            end
            if type == 2 || type == 4
                if LEDs(txs,ty) == 1
                    nr = nr + 1;
                    imageColOrder(txs,ty) = nr;
                end
            end
        end
    end
else
    % only displaying
    imageColOrder = LEDs;
    imageColOrder(imageColOrder == 0) = -10;
end

%% displaying
% colors: 
%   -16 - red
%   -15:nImgs - as in parula color map
% values
%   -16 - red line + F and L letters marking first and last LED
%   -15 - lines separating squares that represent LEDs
%   -10 - LEDs not used to collect images
%   1:nimgs - first LED, second LED, third LED, ... , last LED
for my = 0:8:8*ny-8
    for mx = 0:8:8*nx-8
        displayingOrder(my+1,mx+1:mx+8) = -15;
        displayingOrder(my+1:my+8,mx+1) = -15;
        displayingOrder(my+2:my+8,mx+2:mx+8) = imageColOrder(my/8+1,mx/8+1);
    end
end
displayingOrder(end+1,1:end) = -15;
displayingOrder(1:end,end+1) = -15;

imageColOrder(imageColOrder == -10) = 0;

% displaying v2 - adding lines

nImgs = max(max(imageColOrder));

for m = 2:nImgs-1
    [yb,xb] = find(imageColOrder == m-1);
    [y,x] = find(imageColOrder == m);
    [ya,xa] = find(imageColOrder == m+1);
    if abs(yb-y) + abs(xb-x) == 1
        if yb == y - 1
            displayingOrder(8*(y-1)+2:8*(y-1)+5,8*(x-1)+5) = -16;
        end
        if yb == y + 1
            displayingOrder(8*(y-1)+8:-1:8*(y-1)+5,8*(x-1)+5) = -16;
        end
        if xb == x - 1
            displayingOrder(8*(y-1)+5,8*(x-1)+2:8*(x-1)+5) = -16;
        end
        if xb == x + 1
            displayingOrder(8*(y-1)+5,8*(x-1)+8:-1:8*(x-1)+5) = -16;
        end
    else
        if yb < y && xb < x
            for n = 2:5
                displayingOrder(8*(y-1)+n,8*(x-1)+n) = -16;
            end
        end
        if yb > y && xb < x
            for n = 2:5
                displayingOrder(8*(y-1)+10-n,8*(x-1)+n) = -16;
            end
        end
        if yb < y && xb > x
            for n = 2:5
                displayingOrder(8*(y-1)+n,8*(x-1)+10-n) = -16;
            end
        end
        if yb > y && xb > x
            for n = 2:5
                displayingOrder(8*(y-1)+10-n,8*(x-1)+10-n) = -16;
            end
        end
    end
    
    if abs(ya-y) + abs(xa-x) == 1
        if ya == y -1
            displayingOrder(8*(y-1)+2:8*(y-1)+5,8*(x-1)+5) = -16;
        end
        if ya == y + 1
            displayingOrder(8*(y-1)+8:-1:8*(y-1)+5,8*(x-1)+5) = -16;
        end
        if xa == x - 1
            displayingOrder(8*(y-1)+5,8*(x-1)+2:8*(x-1)+5) = -16;
        end
        if xa == x + 1
            displayingOrder(8*(y-1)+5,8*(x-1)+8:-1:8*(x-1)+5) = -16;
        end
    else
        if ya < y && xa < x
            for n = 2:5
                displayingOrder(8*(y-1)+n,8*(x-1)+n) = -16;
            end
        end
        if ya > y && xa < x
            for n = 2:5
                displayingOrder(8*(y-1)+10-n,8*(x-1)+n) = -16;
            end
        end
        if ya < y && xa > x
            for n = 2:5
                displayingOrder(8*(y-1)+n,8*(x-1)+10-n) = -16;
            end
        end
        if ya > y && xa > x
            for n = 2:5
                displayingOrder(8*(y-1)+10-n,8*(x-1)+10-n) = -16;
            end
        end
    end
end

% mask First
maskF = [
    0 0 0 0 0
    0 1 1 1 1
    0 0 0 0 0
    0 1 1 1 1
    0 1 1 1 1];

% mask Last
maskL = [
    0 1 1 1 1
    0 1 1 1 1
    0 1 1 1 1
    0 1 1 1 1
    0 0 0 0 1];

[y,x] = find(imageColOrder == 1);
sy = 8*(y-1)+3:8*(y-1)+7;
sx = 8*(x-1)+3:8*(x-1)+7;
displayingOrder(sy,sx) = displayingOrder(sy,sx).*maskF;
[y,x] = find(imageColOrder == nImgs);
sy = 8*(y-1)+3:8*(y-1)+7;
sx = 8*(x-1)+3:8*(x-1)+7;
displayingOrder(sy,sx) = displayingOrder(sy,sx).*maskL;

displayingOrder(displayingOrder == 0) = -16;

colorMap=parula(nImgs+16);
colorMap = [1,0,0;colorMap];
end
```
