## Supplementary material for "FPM app: an open-source application for simple and intuitive Fourier ptychographic reconstruction": FPM app: LEDMatrix.m

```
function [LEDs, LEDs2, nx, ny, colorMap] = LEDMatrix(type, a, b, xcen, ycen, LEDsIn, iCO, prevN)
% Fuction that creates LED matrix (LEDs). Also returns matrix (LEDs2) that
% nicely visualize LED layout, marks the central LED and LED that was used
% to collect currently previewed image (in main app window)
%   Inputs:
%       type - type of the LED matrix/action to do
%           type = 1 - circular LED layout
%           type = 2 - rectangular LED layout
%           type = 3 - custom layout
%           type - 4 - only set center LED
%           type = 5 - only create LEDs2 (visualization) matrix
%       a - diamter/width (if needed)
%       b - height (if needed)
%       xcen, ycen - center LED position
%       LEDsIn - LED matrix (only needed if want to visualize already
%                existing matrix)
%       iCO - image collecting order (only needed to show preview image
%             position)
%       prevN - preview image number (only needed to show preview image
%             position)
%   Outputs:
%       LEDs - LED matrix
%           0 in place where there is no LED
%           1 in place where there is LED
%           2 in place where there is central LED
%           Example:
%               LEDs = [0,1,1,1,0
%                       1,1,2,1,1
%                       0,1,1,1,0];
%       LEDs2 - LED matrix in displaying mode
%       ny,nx - size of the LEDs
%   How to display LEDs2 nicely:
%       imagesc(0.5:ny+1,0.5:nx+1,LEDs2); colormap(colorMap)

visualize = 0;  % allow to visualize LED matrix
if type == 1 && a > 0   % circular LED layout
    D = round(a);
    LEDs = zeros(D);
    [x,y] = meshgrid(-(D-1)/2:(D-1)/2);
    r = sqrt(x.^2+y.^2);
    LEDs(r<=a/2) = 1;
    
    LEDs(round(D./2),round(D./2)) = 2;
    visualize = 1;
end

if type == 2 && a > 0 && b > 0  % rectangular LED layout
    a = round(a);
    b = round(b);
    LEDs = ones(a,b);
    LEDs(round(a./2),round(b./2)) = 2;
    visualize = 1;
end

if type == 3    % custom layout
    LEDs = cell2mat(a);
    LEDs = LEDs - 48;
    visualize = 1;
end

if type == 4    % only set center LED
    LEDs = LEDsIn;
    visualize = 1;
    if xcen > 0 && ycen > 0
        LEDs(LEDs == 2) = 1;
        LEDs(xcen,ycen) = 2;    % center LED
    end
end

if type == 5 % only create LEDs2 (visualization) matrix
    LEDs = LEDsIn;
    visualize = 1;
end

%% visualize
% colors:
%   -1:2 - as parula color map
%   3 - red
% values:
%   -1 - lines separating squares that represent LEDs
%   0 - LEDs not used to collect images
%   1 - LEDs not used to collect images
%   2 - center LED
%   3 - previewed images LED border
if visualize == 1
    [ny,nx] = size(LEDs);
    for m1 = 0:7:7*nx-7
        for m2 = 0:7:7*ny-7
            LEDs2(m2+1,m1+1:m1+7) = -1;
            LEDs2(m2+1:m2+7,m1+1) = -1;
            LEDs2(m2+2:m2+7,m1+2:m1+7) = LEDs(m2/7+1,m1/7+1);
        end
    end
    LEDs2(end+1,1:end) = -1;
    LEDs2(1:end,end+1) = -1;
    colorMap = parula(4);
    % mark preview image
    if nargin > 6 && prevN > 0
        [pLy,pLx] = find(iCO == prevN); % preview image LED [y,x] position
        if ~isnan(pLy)
            pLy = 7*(pLy-1); pLx = 7*(pLx-1);
            LEDs2(pLy+1,pLx+1:pLx+8) = 3;
            LEDs2(pLy+1:pLy+8,pLx+1) = 3;
            LEDs2(pLy+8,pLx+1:pLx+8) = 3;
            LEDs2(pLy+1:pLy+8,pLx+8) = 3;
            
            colorMap = [colorMap;1,0,0];
        end
    end
    
    
end

%     imagesc(0.5:ny+1,0.5:nx+1,LEDs2); colormap(colorMap)
end
```
