## Supplementary material for "FPM app: an open-source application for simple and intuitive Fourier ptychographic reconstruction": FPM app: ROISelection.m

```
function ROI = ROISelection(in)
% Function that selects Region Of Interest
%   Input:
%       in - image
%   Output:
%       ROI - Region Of Interest       
%           ROI = [x0,y0,xSize,ySize];
figure(100);
set(gcf,'Name','ROI selection');
set(gcf,'NumberTitle','off');
img = imagesc(in); colormap gray; title('ROI selection');
[~,~,~,ROI] = imcrop(img);
ROI = round(ROI);
if ~isempty(ROI)
    if ( mod(ROI(3),2) == 1 ), ROI(3) = ROI(3)+1; end 
    if ( mod(ROI(4),2) == 1 ), ROI(4) = ROI(4)+1; end
    delete(gcf)
end

end
```
