## Supplementary material for "FPM app: an open-source application for simple and intuitive Fourier ptychographic reconstruction": FPM app: Used_LEDs.m

```
function [LEDsUsed, LEDsUsedDispl, nx, ny] = Used_LEDs(LEDs,systemSetup,type,rmin,rmax,used)
% Function that selects with LEDs are going to be used in reconstruction
%   Inputs:
%       LEDs - LED matrix
%           0 in place where there is no LED
%           1 in place where there is LED
%           2 in place where there is central LED
%           Example:
%               LEDs = [0,1,1,1,0
%                       1,1,2,1,1
%                       0,1,1,1,0];
%       systemSetup - system setup
%           systemSetup.NA - NA
%           systemSetup.LEDspacing - spacing between neighbour LEDs in LED
%                                    matrix (mm)
%           systemSetup.LEDheight - distance between LED matrix and a 
%                                   sample (mm)
%       type - which LEDs are going to be used
%           type = 0 - only displaying (creating LEDsUsedDisp matrix)
%           type = 1 - all LEDs
%           type = 2 - only brightfield LEDs
%           type = 3 - only darkfield LEDs
%           type = 4 - custom LEDs
%       rmin,rmax - min and max distance from central LED (if type = 4)
%       used - used LEDs matrix (if type = 0)
%   Outputs:
%       LEDsUsed - used LEDs
%           0 in place where there is no LED or no used LED
%           1 in place where there is used LED
%           Example:
%               LEDsUsed = [0,1,1,1,0
%                           1,1,0,1,1
%                           0,1,1,1,0];
%       LEDsUsedDispl - used LEDs (display mode)
%       nx,ny - LEDsUsed size
%   How to display it nicely:
%       imagesc(0.5:ny+1,0.5:nx+1,LEDsUsedDispl, [-1 2]);

%% central LED position
    [cledY,cledX] = find(LEDs==2);
    LEDs(LEDs>1) = 1;

%% calculating LEDs illumination NA
if type >= 2 && type <= 4
    
    [ys,xs] = size(LEDs);

    LEDspacing = systemSetup.LEDspacing; % spacing between adjacent LEDs
    LEDheight = systemSetup.LEDheight; % distance bewteen the LED matrix and the sample
    NA = systemSetup.NA;
    xx = 1:xs; xx = (xx - cledX).*LEDspacing;
    yy = 1:ys; yy = (yy - cledY).*LEDspacing;
    [LEDsPosX,LEDsPosY] = meshgrid(xx,yy);

    dist = sqrt(LEDsPosX.^2+LEDsPosY.^2+LEDheight.^2);
    sin_thetaX = LEDsPosX./dist;
    sin_thetaY = LEDsPosY./dist;

    illuminationNA = sqrt(sin_thetaX.^2+sin_thetaY.^2);
    illuminationNA = illuminationNA.*LEDs;
end

%% Used LEDs
switch type 
    case 0
        LEDsUsed = used;
    case 1
        LEDsUsed = LEDs;
    case 2
        LEDsUsed = zeros(size(LEDs));
        LEDsUsed(illuminationNA<NA & LEDs > 0) = 1;
    case 3
        LEDsUsed = zeros(size(LEDs));
        LEDsUsed(illuminationNA>NA & LEDs > 0) = 1;
    case 4
        rr = sqrt(LEDsPosX.^2+LEDsPosY.^2)/LEDspacing;
        LEDsUsed = zeros(size(LEDs));
        LEDsUsed(rr>rmin & rr<=rmax) = 1;
        LEDsUsed(LEDs==0) = 0;
        if rmin == 0
            LEDsUsed(rr == 0) = 1;
        end
end

%% displaying

[ny,nx] = size(LEDs);
for m1 = 0:7:7*nx-7
    for m2 = 0:7:7*ny-7
        LEDsUsedDispl(m2+1,m1+1:m1+7) = -1;
        LEDsUsedDispl(m2+1:m2+7,m1+1) = -1;
        LEDsUsedDispl(m2+2:m2+7,m1+2:m1+7) = LEDs(m2/7+1,m1/7+1) + ...
            (LEDsUsed(m2/7+1,m1/7+1)-1)/2;
    end
end
LEDsUsedDispl(end+1,1:end) = -1;
LEDsUsedDispl(1:end,end+1) = -1;

end
```
