## Supplementary material for "FPM app: an open-source application for simple and intuitive Fourier ptychographic reconstruction": FPM app: info.txt

%%%%%%%%%%%%%%%%%%%%%%%%%%%%%%%%%
% FPM app
% version 1.0
% 
% Application for performing Fourier ptychography reconstruction.
% Run main.m script with MATLAB to launch FPM app
% 
% This app was created as a part of master's thesis at Faculty of 
% Mechatronics, Warsaw University of Technology, Warsaw, Poland
% 
% Created by:
%   Mikolaj Rogalski;
% 
% Last modified:
%   30.11.2020
% 
% In case of any problem with FPM app, try to find a soluction in
% FPM_app_documentation.pdf or contact author by e-mail
%%%%%%%%%%%%%%%%%%%%%%%%%%%%%%%%%
% 
% FPM app reconstruction code was based on Lei Tian implementation of: 
% "Fourier ptychography reconstruction algorithm with Quasi-Newton’s
% method"
% which can be downloaded here:
% http://sites.bu.edu/tianlab/open-source/
% Article:
% "L. Tian, X. Li, K. Ramchandran, and L. Waller, “Multiplexed coded 
% illumination for Fourier Ptychography with an LED array microscope,” 
% Biomed. Opt. Express, vol. 5, no. 7, p. 2376, Jul.2014, 
% doi: 10.1364/BOE.5.002376."
% 
% FPM app Synthetic data generation was inspired by codes attached to:
% "G. Zheng, Fourier Ptychographic Imaging A MATLAB tutorial. 
% IOP Publishing, 2016" 
% These codes may be downloaded here:
% https://smartimaging.uconn.edu/fourier-ptychtography/#
% 
% FPM app uses  block-matching and 3D filtering (BM3D) digital denoising 
% algorithm, which may be downloaded here:
% http://www.cs.tut.fi/~foi/GCF-BM3D/
% Article:
% "K. Dabov, A. Foi, and K. Egiazarian, “Video denoising by sparse 3D 
% transform-domain collaborative filtering,” Eur. Signal Process. Conf., 
% vol. 16, no. 8, pp. 145–149, 2007."
% 
% FPM app uses Douglas M. Schwarz sort_nat sorting algorithm to sort 
% strings in natural order. This algorithm may be downloaded here:
% https://uk.mathworks.com/matlabcentral/fileexchange/10959-sort_nat-natural-order-sort
% 
% All licenses may be found in LICENSES directory. BM3D and sort_nat
% licenses are also respectively in:
%       Algorithm functions/BM3D
% or in:
%       Algorithm functions/sort_nat
% directories, where those algorithms are located
% 
%%%%%%%%%%%%%%%%%%%%%%%%%%%%%%%%%
