## Supplementary material for "FPM app: an open-source application for simple and intuitive Fourier ptychographic reconstruction": FPM app: LICENSE.txt

---------------------------------------------------------------------
Copyright (2020): Mikołaj Rogalski

This code is distributed under the terms of the GNU General Public
License 2.0.

Permission to use, copy, modify, and distribute this software excluding 
BM3D and sort_nat algorithms for any purpose without fee is hereby 
granted, provided that this entire notice is included in all copies of 
any software which is or includes a copy or modification of this 
software and in all copies of the supporting documentation for such 
software.
This software is being provided "as is", without any express or
implied warranty.  In particular, the authors do not make any
representation or warranty of any kind concerning the merchantability
of this software or its fitness for any particular purpose.

Using, copying, modifying, and distributing of BM3D and sort_nat 
algorithms is allowed according to licenses attached to these 
algorithms.
---------------------------------------------------------------------
